# Substitutions at the C-8 position of quinazolin-4-ones improve the potency of nicotinamide site binding tankyrase inhibitors

**DOI:** 10.1101/2024.06.23.600314

**Authors:** Chiara Bosetti, Dionisis Kampasis, Albert Galera-Prat, Shoshy A. Brinch, Maria Karelou, Saurabh S. Dhakar, Juho Alaviuhkola, Jo Waaler, Lari Lehtiö, Ioannis Kostakis

## Abstract

Human diphtheria toxin-like ADP-ribosyltransferases, PARPs and tankyrases, transfer ADP-ribosyl groups to other macromolecules, thereby controlling various signaling events in cells. They are considered promising drug targets, especially in oncology, and some small molecule inhibitors have already been developed. These inhibitors typically interact with the nicotinamide binding site and extend along the NAD^+^ binding groove of the catalytic domain. Quinazolin-4-ones have been explored as promising scaffolds for such inhibitors and we have identified a new position within the catalytic domain that has not been extensively studied yet. In this study, we investigate larger substituents at the C-8 position and, using X-ray crystallography, we demonstrate that nitro- and diol-substituents engage in new interactions with TNKS2, improving both affinity and selectivity. Both nitro- and diol-substituents exhibit intriguing inhibition of TNKS2, with compound **49** displaying an IC_50_ of 65 nM, while compound **40**’s IC_50_ value is 14 nM. Both analogues show efficacy in cell assays and attenuate the tankyrase-controlled Wnt/β-catenin signaling with sub-micromolar IC_50_. When tested against a wider panel of enzymes, compound **40** displayed high selectivity towards tankyrases, whereas **49** also inhibited other PARPs. The results offer new insights for inhibitor development targeting tankyrases and PARPs by focusing on the subsite between a mobile active site loop and the canonical nicotinamide binding site.

## Introduction

The enzyme family of diphtheria toxin-like ADP-ribosyltransferases (ARTDs) encompasses seventeen proteins in human characterized by a structurally conserved catalytic domain that performs the ADP-ribosylation reaction on proteins and on nucleic acids [1]. Using β-nicotinamide adenine dinucleotide (NAD^+^) as a substrate, these enzymes transfer single or multiple ADP-ribose units onto macromolecules, resulting in mono-ADP-ribosylation (MARylation) or poly-ADP-ribosylation (PARylation), respectively. While the majority of ARTDs are MARylating enzymes, PARP1, PARP2, and tankyrases (TNKS1 and TNKS2) can produce PAR [2]. The PARylating representatives of the family share a triad of residues (H-Y-E) responsible for the coordination of NAD^+^ in the catalytic pocket. Although the glutamate residue in the triad appears to play a fundamental role in the elongation of the PAR chain, it is also present in the MARylating enzymes PARP3 and PARP4. Moreover, this residue alone is not sufficient to confer the PARylation activity in mutated enzymes [2–5] and differences in the NAD^+^ acceptor site may also limit the modification to linear PAR chains in tankyrases [6] or to MARylation in PARP3 and PARP4 [7].

PARP1 and PARP2 are renowned for their ability to sense DNA strand breakage and to catalytically act on the detected damage to stimulate restorative cellular responses [8]. These tasks are achieved by PARP1 and PARP2 multidomain structures, which allow precise regulation of the PARylation process. In PARP1, three zinc-finger domains (Zn1, Zn2, Zn3) and a WGR domain are engaged in DNA binding, while a helical domain (HD) undergoes structural reorganization, which results in an open conformation that fully exposes the active site to the substrate [9–11]. In addition, PARP1 can bind intact nucleosomal DNA through its BRCT domain [12]. PARP2 has a structurally similar C-terminus with WGR, HD and ART domains like PARP1, but it lacks the N-terminal Zinc fingers. In PARP2, the WGR domain is responsible for DNA binding and has been shown to bridge over double-strand DNA gaps [13]. Interaction of the WGR domain with DNA triggers structural changes in the HD domain, leading to a shift of the catalytic domain and opening of the active site [14,15]. The role of PARP1/2 in DNA lesion detection has gained extensive interest upon the discovery of sensitivity of cells defective in homologous recombination to their inhibition. Notably, BRCA mutated cells suffer from a defective DNA repair system, and further depletion of DNA break sensing processes through PARP1 inhibition leads to persistent damage and apoptosis [16,17]. Consequently, PARP1/2 inhibition has been studied as a valid therapeutic approach for specific treatment of BRCA mutated cancers.

Tankyrases range of action in cellular processes is wide due to their ability to modulate the turnover of a vast diversity of proteins. Notably, tankyrases modify protein substrates through their catalytic domain (ART), whose activity depends on helical polymerimerization mediated by their sterile alpha motif (SAM) domain [18–22]. Consequentially, PARylated proteins are ubiquitylated by the PAR-dependent E3 ligase RNF146 and undergo proteasomal degradation [23–26]. In addition to their catalytic functions, tankyrases establish protein-protein interactions by employing five conserved ankyrin repeat cluster (ARC) domains, enhancing their contribution to different biological pathways [27].

The orchestrating activity of tankyrases in the Wnt/β-catenin signaling pathway has been extensively demonstrated. Tankyrases bind and PARylate AXIN (AXIN1/AXIN2), the concentration-limiting and scaffolding component of the β-catenin destruction complex, leading to stabilization of β-catenin and transcriptional activation of target genes [19,22,28–30]. Since deregulation of Wnt/β-catenin signaling serves as driving force in a variety of diseases [31], including several types of cancer [32] and fibrotic disorders [33–35], inhibition of tankyrases has emerged as a compelling therapeutic strategy [6,36]. The therapeutic importance of tankyrases inhibition is further corroborated by its impact on a plethora of cellular processes, encompassing the Hippo/YAP pathway [36,37], telomere maintenance [38–41], mitotic spindle assembly [39,40,42,43], insulin sensitivity [44–46], and LKB1-AMPK signaling [47].

Given the paramount roles of PARylating ARTDs enzymes, the discovery of PARP inhibitors has been greatly incentivized since the ‘90s [48] and culminated in 2014 with the approval of Olaparib as treatment for advanced ovarian cancer with BRCA germline mutation [49]. Over the years, multiple compound scaffolds have been developed [50,51] and other clinical approvals have followed for the treatment of ovarian, breast and prostate cancers [52–54]. All the approved drugs and most of the developed PARP inhibitors bind to the conserved nicotinamide (NI) pocket of the ART domain, interacting with the protein in the same fashion as the nicotinamide moiety of NAD^+^. The strategy to achieve higher affinity and selectivity of these NI-site binders towards defined ARTDs includes extending the compounds beyond the NI pocket, where it is possible to tailor the binding interactions for different ARTD enzymes [55].

The journey towards the discovery of tankyrases inhibitors began with XAV939, which binds with nanomolar potency to the NI site [28] (**Table 1**). Although not exclusively selective towards tankyrases, this chemical probe has proven to be a powerful tool for investigating their involvement in multiple cellular processes. At the same time, the compound IWR-1 was identified to impact the Wnt-signaling pathway through tankyrase inhibition [56]. Notably, IWR-1 was the first discovered to uniquely target the adenosine pocket of the NAD^+^ binding cleft [57]. Subsequent studies led to the development of tankyrase inhibitors that bind either to the NI site [58–65], the adenosine site [66–73] or both sub-pockets [74–76]. Some of these inhibitors have demonstrated a favorable therapeutic window in cancer models [77] and a few have even entered clinical trials [78–80].

**Table 1.**
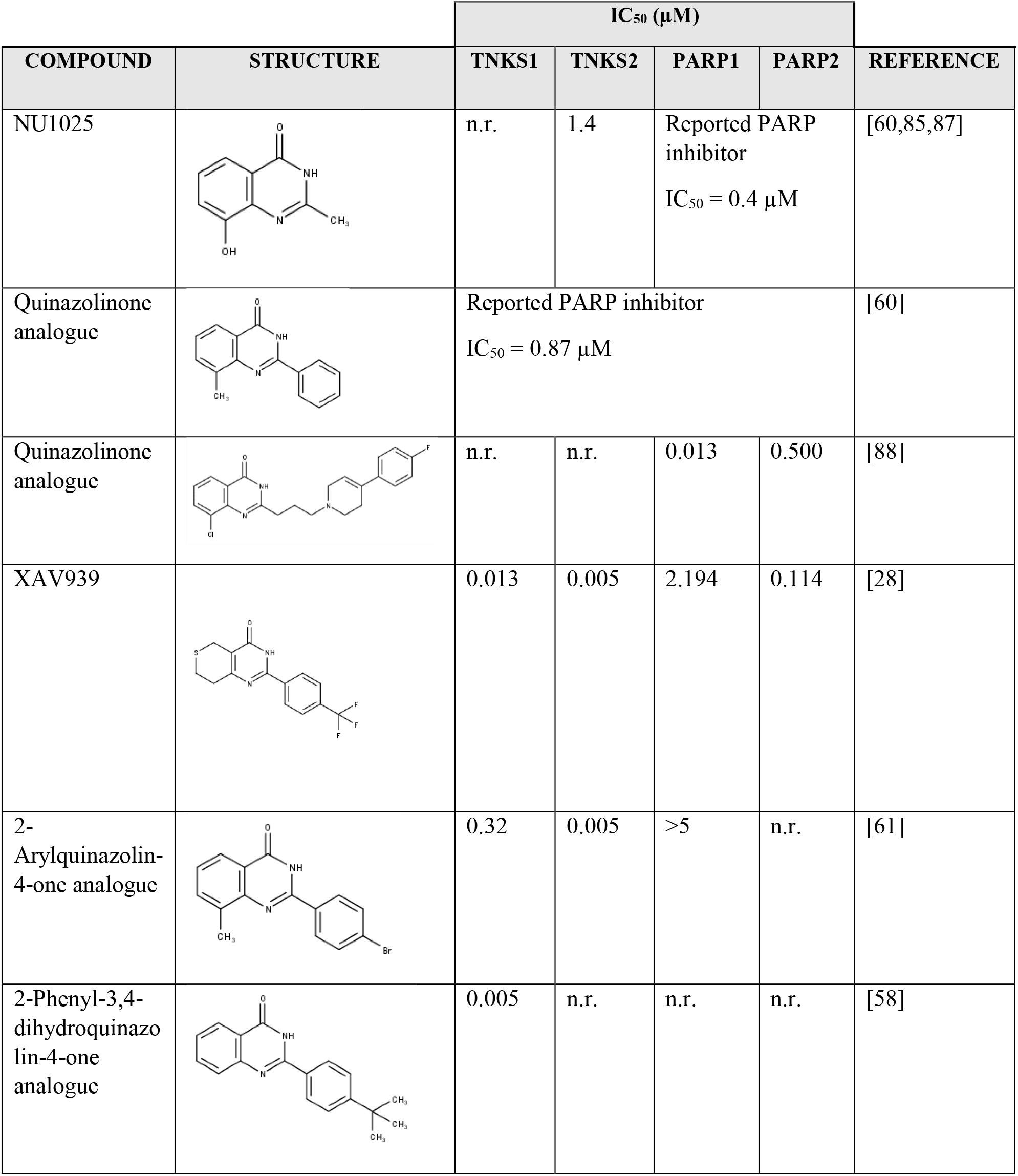
Compounds sharing the quinazolin-4-one scaffold and binding to the NI site. IC_50_ values for TNKS1/2 and PARP1/2 are indicated (n.r. = not reported). The 2-Phenyl-3,4-dihydroquinazolin-4-one analogue shown displays selectivity towards TNKS1 over six other ARTD family members [58].

|  |  | IC<sub>50</sub> (μM) |  |  |  |  |
| --- | --- | --- | --- | --- | --- | --- |
| COMPOUND | STRUCTURE | TNKS1 | TNKS2 | PARP1 | PARP2 | REFERENCE |
| NU1025 |  | n.r. | 1.4 | Reported PARP inhibitor<br>IC<sub>50</sub> = 0.4 μM |  | [60,85,87] |
| Quinazolinone analogue |  | Reported PARP inhibitor<br>IC<sub>50</sub> = 0.87 μM |  |  |  | [60] |
| Quinazolinone analogue |  | n.r. | n.r. | 0.013 | 0.500 | [88] |
| XAV939 |  | 0.013 | 0.005 | 2.194 | 0.114 | [28] |
| 2-Arylquinazolin-4-one analogue |  | 0.32 | 0.005 | >5 | n.r. | [61] |
| 2-Phenyl-3,4-dihydroquinazolin-4-one analogue |  | 0.005 | n.r. | n.r. | n.r. | [58] |

Among the numerous compound scaffolds unveiled in recent years, quinazoline derivatives have emerged as promising candidates in chemotherapy [60]. They exhibit versatile mechanisms of action, functioning as protein kinase inhibitors [81], apoptosis inducers and modulators of diverse cellular signaling pathways [82–84]. One of the first inhibitor scaffolds to target PARPs was quinazolin-4-one, and the small PARP inhibitor NU1025 was already shown in the ‘90s to potentiate the effect of chemotherapeutic agents [85]. In the quest to elaborate on structural activity relationships (SAR), it was discovered that 2-phenyl analogues significantly reduce the potency against PARPs [60]. In contrast, the 2-phenyl quinazolin-4-ones exhibit improved potency and can even provide selectivity towards tankyrases over other PARPs (**Table 1**). The quinazolin-4-one core occupies the NI site, while the phenyl moiety extends to a region lined by hydrophobic residues in tankyrases, explaining the observed selectivity [58,86] (**Fig. 1**).

**Fig. 1.**
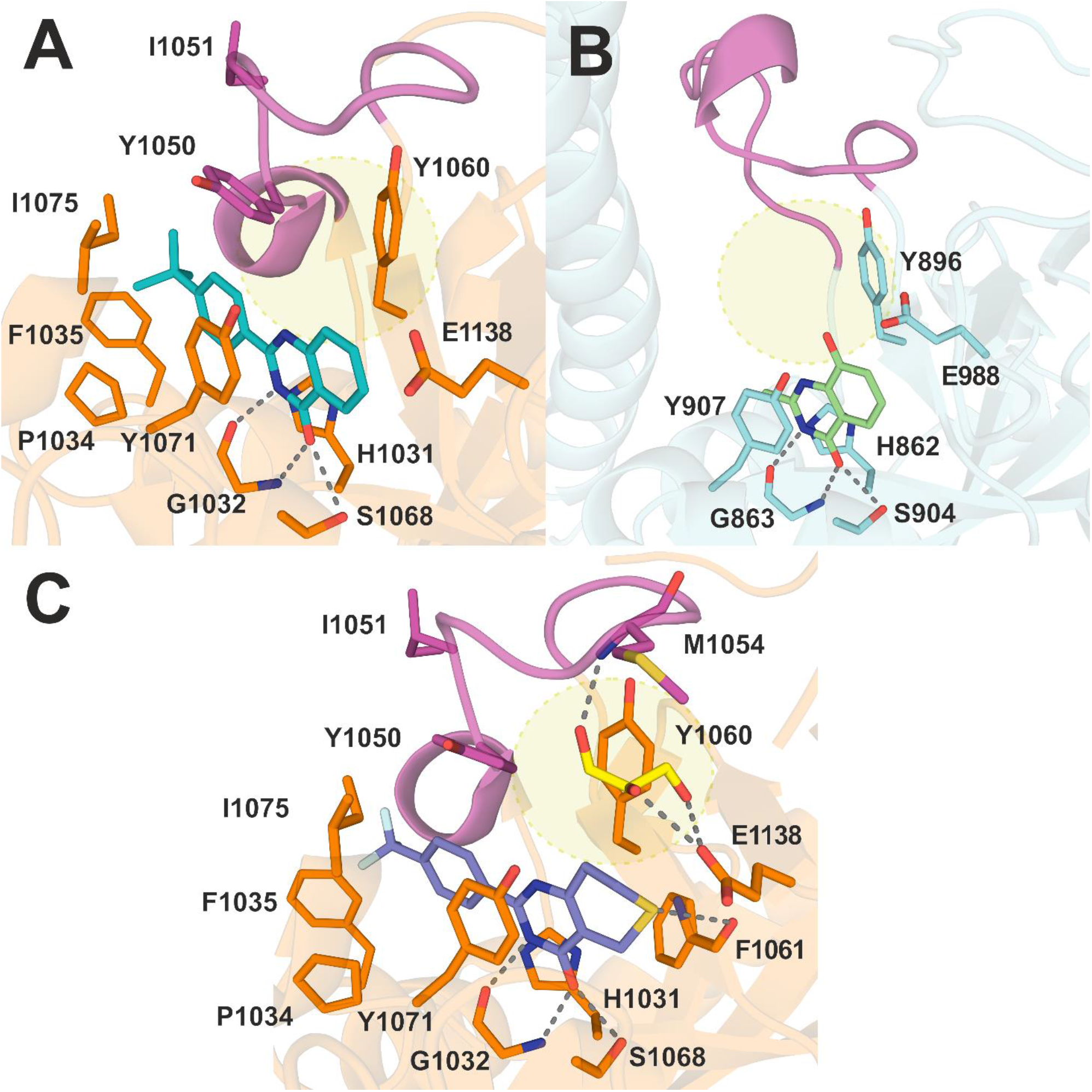
Crystal structures of compounds characterized by quinazolin-4-one scaffold and XAV939 occupying the NI pocket. **A)** Crystal structure of compound (PDB: 4BUD) with human TNKS2 [58]. Hydrogen bonds established with residues of the pocket are represented with dashed lines. The D-loop is colored in fuchsia to highlight its distance from the compound and the space that we targeted during the design of inhibitors presented in this study (in yellow). **B)** Crystal structure of the small PARP inhibitor NU1025 with *Gallus gallus* PARP1 (PDB: 4PAX) [89]. **C)** Crystal structure of XAV939 in complex with human TNKS2 shows the presence of a glycerol molecule (yellow) near positions 7 and 8 and facing the D-loop (PDB: 3KR8) [90]. This glycerol especially establishes hydrogen bonds with the main chain amide of Met1054 and with Glu1138.

While it is well established that the phenyl substituent increases the inhibitor potency towards tankyrases, the study of modifications at the C-7 and C-8 positions has been limited to small substituents [62]. Nevertheless, these compounds display an improved binding in the region of the active site facing the donor site loop (D-loop), a loop often flexible in crystal structures and characterized by high B-factors or even lacking electron density despite the high affinity of the ligand (**Fig. 1**) [91]. Crystal structure data from previous studies revealed that this region is accessible in TNKS2 and could be occupied by substituents at the C-7 and C-8 positions of bound inhibitors (**Fig. 1**) [90]. Consequently, we aimed to explore the SAR of this site with larger substituents that could also establish new interactions. We synthesized a series of substituted quinazolin-4-ones and evaluated them in biochemical assays. Our efforts led to the discovery of novel tankyrase inhibitors characterized by nanomolar potency, with SAR supported by crystal structures, specificity towards tankyrases over human PARP enzymes and activity in cell assays.

## Results

### Design of the quinazolin-4-ones

Aiming to explore the structural requirements needed for PARP inhibition, as well as those that induce specificity towards tankyrases, we designed novel compounds by modifying the C-7 and C-8 positions of the quinazolin-4-one scaffold. Prior studies suggest that small, electron-rich substituents at C-8 position may enhance inhibitory potency towards tankyrases [62]. Moreover, crystal structure data of compound XAV939 in complex with TNKS2 indicates the presence of space surrounding the C-7 and C-8 positions that could be exploited by larger polar groups (**Fig. 1**) [90]. To evaluate this possibility, we synthesized a series of quinazolin-4-ones targeting this region of the binding pocket.

### Chemistry

The synthetic approach presented herein enables the formation of three classes of final derivatives through a shared route. The overall procedure is straightforward and versatile, allowing the synthesis of a multitude of end products with only slight modifications of the reaction conditions. For the 8-nitro substituted analogues **11-13** and **39-41**, we used 2-amino-3-nitrobenzoic acid (**1**) as the starting material, which was stirred with the appropriate chloride under inert conditions to yield the intermediate amides **3-4** and **32** (Schemes 1 and 2). For the synthesis of derivatives **30** and **31**, a slightly different approach was employed, by firstly treating benzoic acid analogues **25** and **26** with ethyl chloroformate, followed by the reaction of intermediates **27** and **28** with 2-amino-3-nitrobenzoic acid. The next step concerns the formation of benzoxazinone derivatives **6-7** and **33-35**, which was achieved by heating amides **3-4** and **30-32** respectively in acetic anhydride. Analogue **5** was obtained in one step, by stirring 2-amino-3-nitrobenzoic acid in acetic anhydride for 20 hours. Treatment of these benzoxazinone derivatives with NH_3_, yielded amides **8-10** and **36**-**38**, and further ring closure under alkaline conditions provided the nitro-quinazolinones **11-13** and **39-41** respectively. The above-mentioned methodology was extended to the synthesis of the 7-substituted analogues **64-66** originating from 2-amino-4-nitrobenzoic acid (***53***) (Scheme 3).

**Scheme 1.**
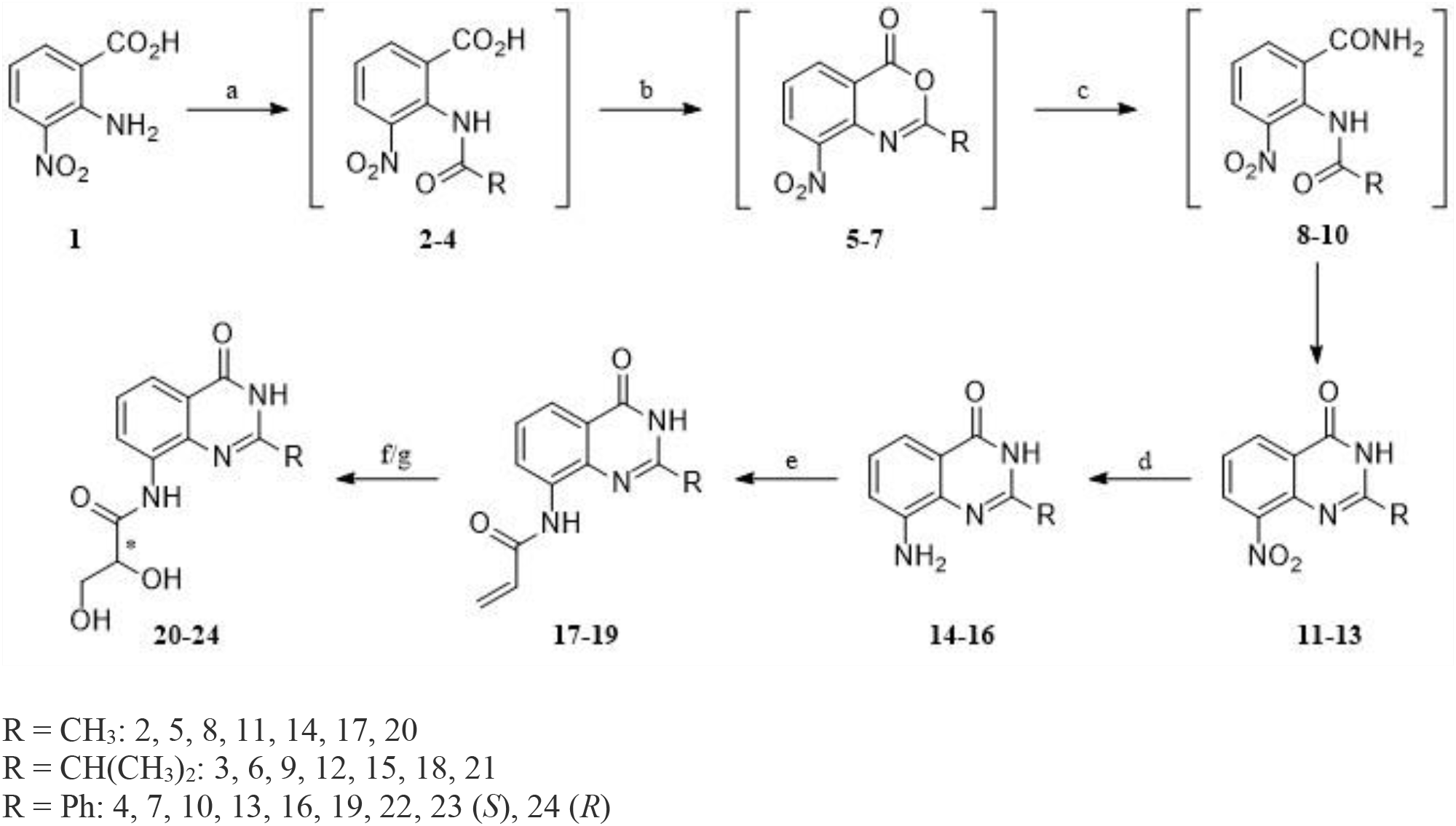
Synthesis of derivatives **11-13** and **17-24**. **Reactants and conditions:** (a) Ac_2_O (*for 2*)/ Appropriate chloride, Et_3_N, THF anh., 0 °C, 24h (*for 3 and 4*); (b) Ac_2_O, reflux, 2h; (c) NH_3_, THF, 12h; (d) H_2,_ Pd/C, EtOH abs., 50 psi, 6h; (e) acryloyl chloride, Et_3_N, THF anh., 16h; (f) OsO_4_, NMO, THF, 72h; (g) AD-mix-α (*for 23*)/ AD-mix-β (*for 24*), tBuOH/H_2_O 1:1, 36h.

**Scheme 2.**
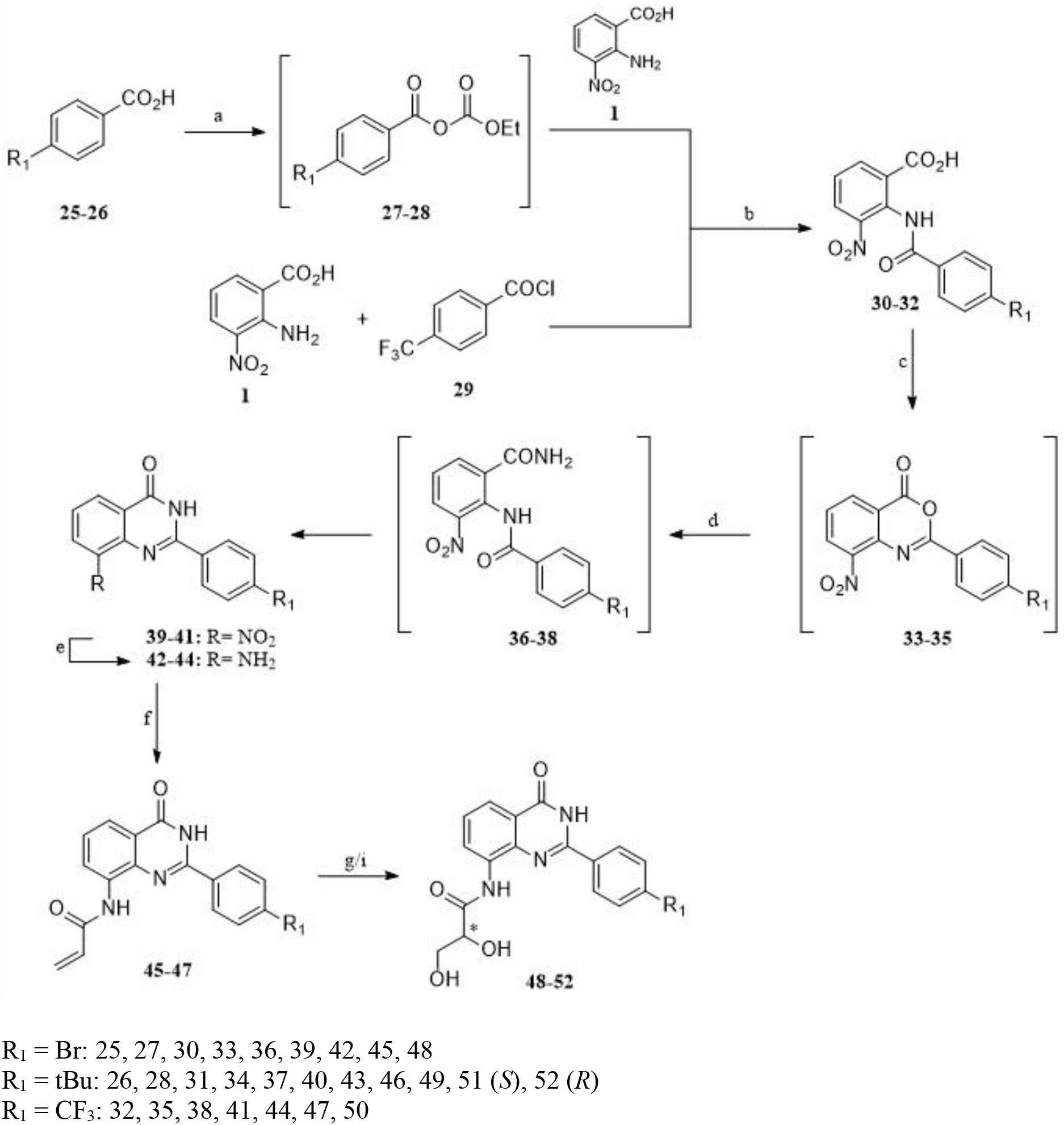
Synthesis of derivatives **39-41** and **45-52**. **Reactants and conditions:** (a) ClCOOEt, Et_3_N, THF anh., 0 °C, 30 min; (b) THF anh., 0 °C, suitable base, 24h, K_2_CO_3_, 12h; (c) Ac_2_O, reflux, 2h; (d) NH_3_, THF, 12h; (e) H_2_-Ni Raney, EtOH abs., 20h (*for 42*), H_2_-Pd/C, EtOH abs., 50 psi, 6h (*for 43-44*); (f) acryloyl chloride, Et_3_N, THF anh., 16h; (g) OsO_4_, NMO, THF, 24h; (i) AD-mix-α (*for 51*)/ AD-mix-β (*for 52*), tBuOH/H_2_O 1:1, 36h.

**Scheme 3.**
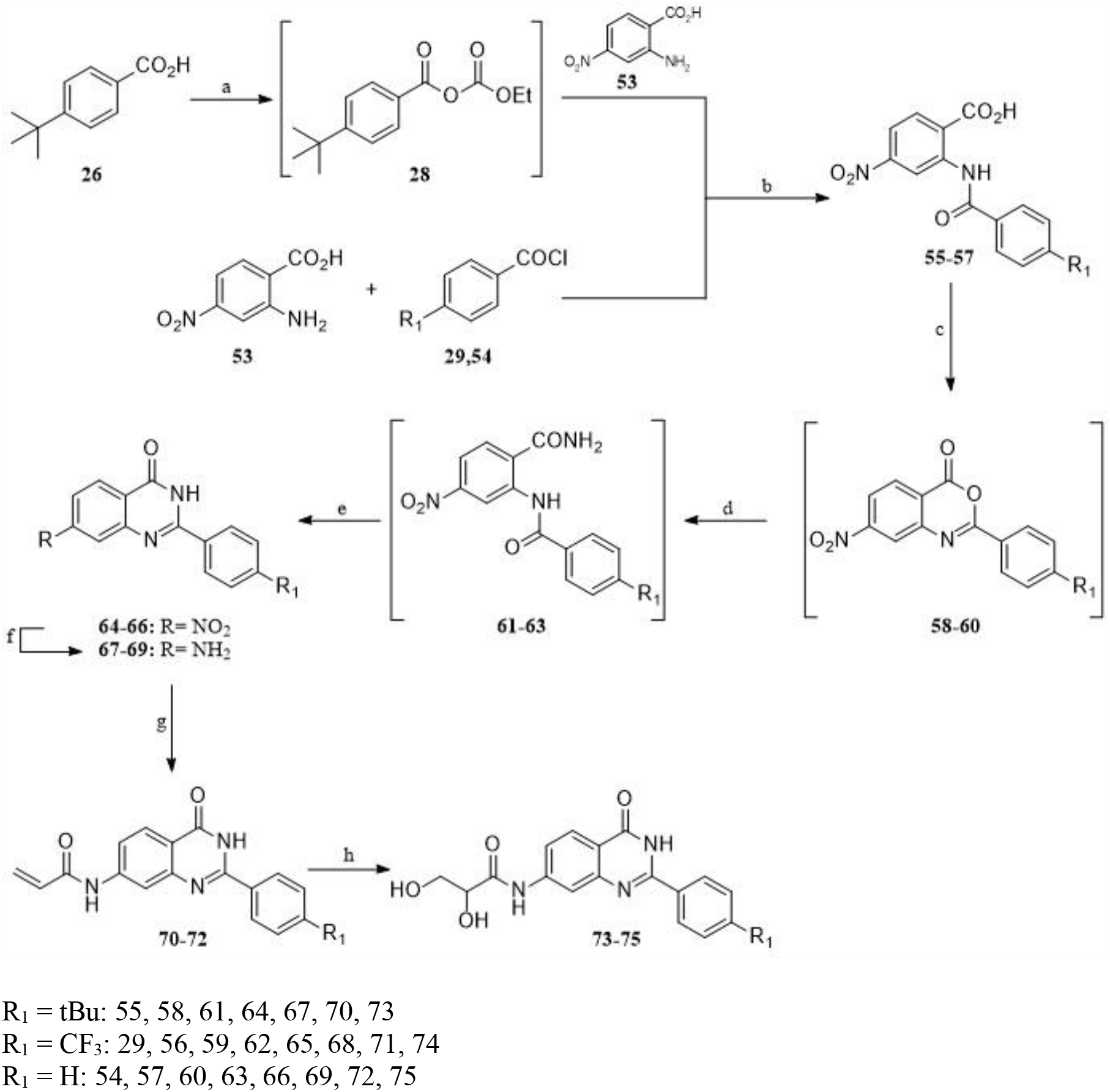
Synthesis of derivatives **64-66** and **70-75**. **Reactants and conditions:** (a) ClCOOEt, Et_3_N, THF anh., 0 °C, 30 min; (b) THF anh., 0 °C, suitable base, 24h, K_2_CO_3_, 12h; (c) Ac_2_O, reflux, 2h; (d) NH_3_, THF, 12h; (e) aq. NaOH 5%, reflux, 1h; (f) H_2_-Pd/C, EtOH abs., 50 psi, 4h; (g) acryloyl chloride, Et_3_N, THF anh., 16h; (h) OsO_4_, NMO, THF, 24h.

Subsequent reduction of the nitro group by catalytic hydrogenation under pressure, afforded amines **14-16**, **42-44** and **67-69** respectively. Under these conditions, the bromo derivative **39** underwent dehalogenation, providing amine **16** instead of the desirable **42**. Thus, in this instance, we performed the reduction under atmospheric pressure, using Ni Raney as the catalyst. These amines were then converted to the corresponding olefins **17-19**, **45-47**, and **70-72** by treatment with the appropriate chloride under inert conditions.

These unsaturated derivatives underwent catalytic syn-hydroxylation with osmium tetroxide and *N*-methylmorpholine-*N*-oxide as the oxidizing agent, to yield cis-diols **20-22**, **48-50**, and **73-75** respectively.

Finally, through an enantioselective process, we were able to obtain enantiomers **23** and **51** by treating acrylamides **19** and **46** respectively with AD-mix-α in a mixture of *t*-BuOH/H_2_O 1:1. Utilizing the same procedure, (R)-diols **24** and **52** were afforded upon treatment with AD-mix-β (Scheme 2).

### Biochemical evaluation and structural studies

To investigate the limits of the NI pocket area facing the D-loop, we initially evaluated the potency of three compounds featuring a bulky diol substituent on C-8 position against PARP1/2 and TNKS1/2 (**Table 2**). These three molecules are distinguished by different hydrophobic groups at the C-2 position, which are analogous to the previously characterized inhibitors of PARP1 and tankyrases (**Fig. 1**). The substituents vary in size and consist of a methyl (compound **20**), an isobutyl (compound **21**) and a phenyl group (compound **22**).

**Table 2.**
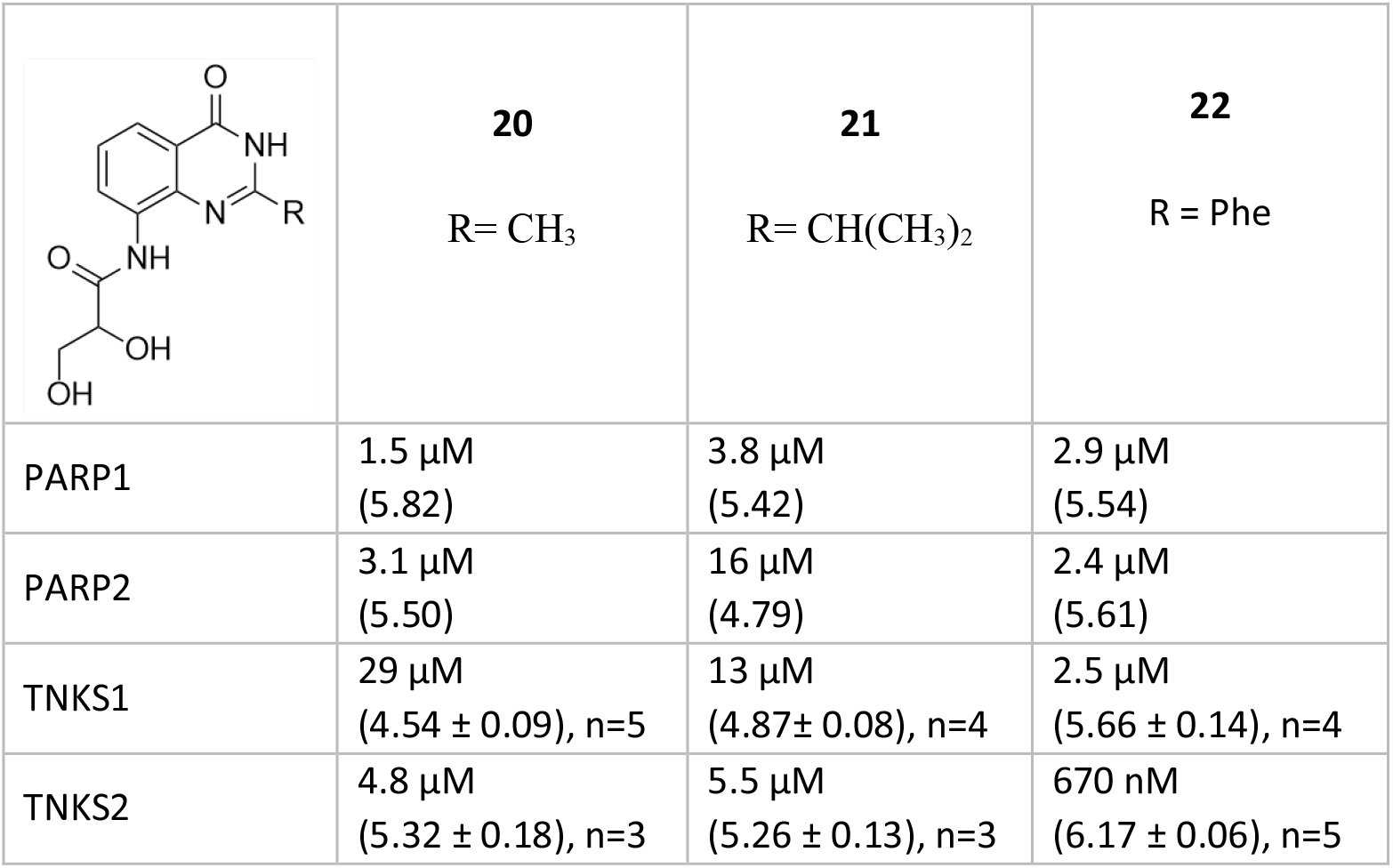
Potencies of the diol substituted compounds against PARP1/2 and TNKS1/2. IC_50_ values (pIC_50_ ± SEM, number of repetitions are indicated) are reported.

|  |  |  |  |
| --- | --- | --- | --- |
|  | **20**<br>R = CH<sub>3</sub> | **21**<br>R = CH(CH<sub>3</sub>)<sub>2</sub> | **22**<br>R = Phe |
| PARP1 | 1.5 μM<br>(5.82) | 3.8 μM<br>(5.42) | 2.9 μM<br>(5.54) |
| PARP2 | 3.1 μM<br>(5.50) | 16 μM<br>(4.79) | 2.4 μM<br>(5.61) |
| TNKS1 | 29 μM<br>(4.54 ± 0.09), n=5 | 13 μM<br>(4.87 ± 0.08), n=4 | 2.5 μM<br>(5.66 ± 0.14), n=4 |
| TNKS2 | 4.8 μM<br>(5.32 ± 0.18), n=3 | 5.5 μM<br>(5.26 ± 0.13), n=3 | 670 nM<br>(6.17 ± 0.06), n=5 |

Interestingly compound **22**, despite being a weak PARP1/2 inhibitor, displayed good potency towards TNKS2. This observation led us to speculate that the combination of the bulky polar diol on C-8 and the phenyl moiety on C-2 could drive selectivity of quinazolin-4-one inhibitors towards tankyrases. Consequently, we focused our efforts on exploring the potential of diol-based compounds as tankyrase inhibitors by introducing various substituents at the *para-*position of the phenyl ring at C-2. Additionally, we synthesized the (R)- and (S)-enantiomers of selected diols to investigate potential disparities in binding affinity. Moreover, to expand our understanding of the chemical space around the quinazolin-4-one core, we designed and synthesized a series of analogues having the diol substituent on C-7 position. Finally, to obtain more robust SAR data, we evaluated the inhibitory potency of the nitro and acrylamido substituted synthetic intermediates, since they possess substantially different electrochemical and steric properties in comparison to our final diols.

### SAR of compounds with C-8 substituents

Since the initial analysis of the diol-analogues highlighted that tankyrases are especially inhibited by the C-2 phenyl quinazolin-4-one scaffold (**Table 2**), we aimed to enhance the potency of compound **22** and we investigated the SAR of analogues by initially assessing their potencies with TNKS2 (**Table 3**). While most of the substituents were well tolerated, it was evident and in agreement with previous studies (**Table 1**), that the C-2 phenyl group was crucial for achieving high potency. Analogues lacking this group exhibited modest affinity at best (**11**,**12**,**17**). Subsequently, we solved co-crystal structures of key analogues with TNKS2 to confirm that the compounds establish common hydrogen bonds with Gly1032 and Ser1068 (**Fig. 2A-D**) in the same fashion as the previously characterized quinazolin-4-ones [58]. Notably, the quinazolin-4-one scaffold forms two hydrogen bonds with the backbone of Gly1032 and the carbonyl group establishes a hydrogen bond with the hydroxyl group of Ser1068. In addition, inhibitors display a π-π stacking interaction with Tyr1071.

**Fig. 2.**
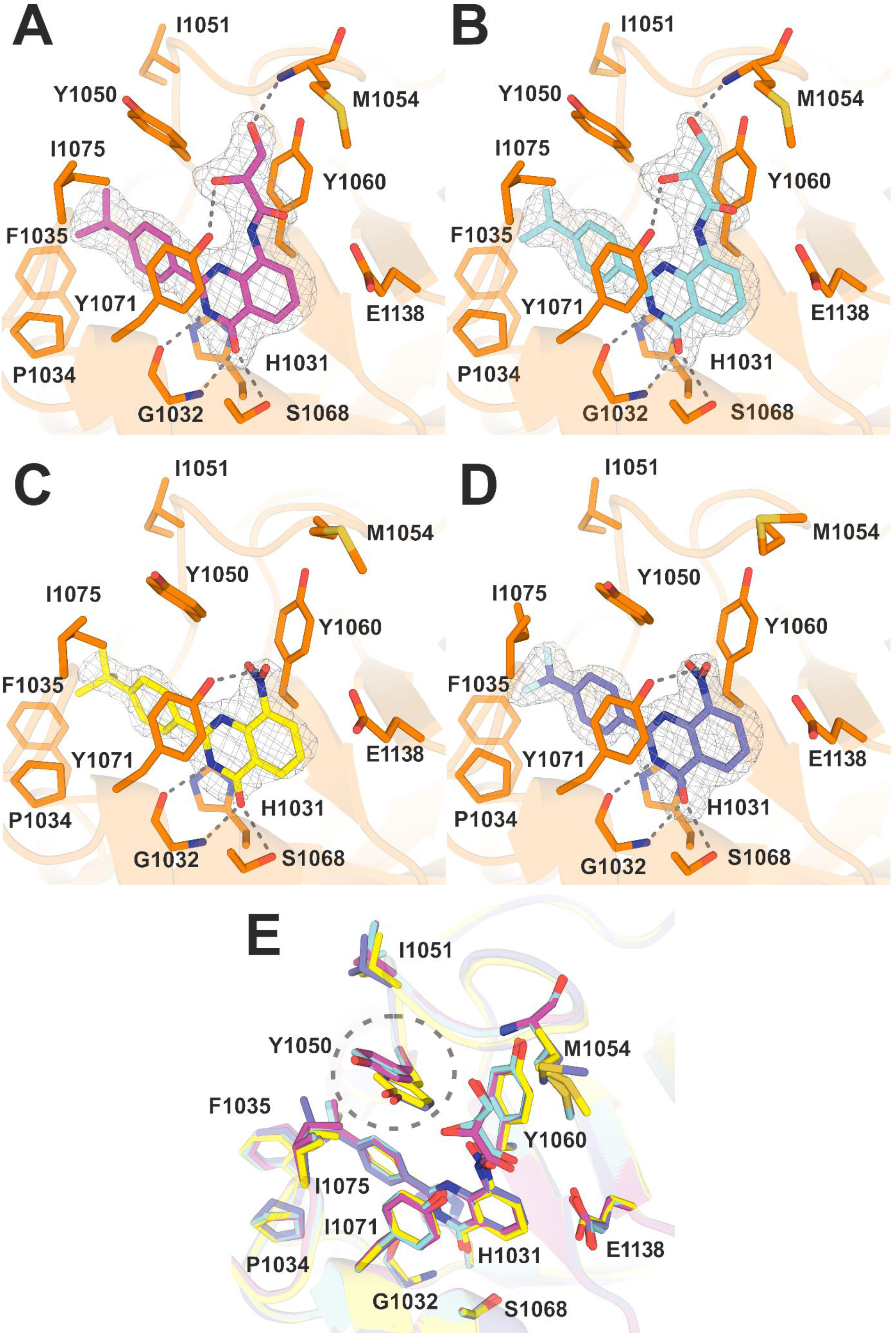
Crystal structures of compounds **51**, **52**, **40**, **41** with TNKS2. The sigma-A weighted 2Fo-Fc electron density maps are shown in gray and contoured at 1.0 σ. Hydrogen bonds established with residues of the pocket are represented with dashed lines. **A)** Crystal structure of compound **51** with TNKS2 (PDB: 8S4W**). B)** Crystal structure of compound **52** (PDB: 8S4V). **C)** Crystal structure of compound **40** (PDB: 8S4X) **D)** Crystal structure of compound **41** (PDB: 8S60). **E)** Superimposed crystal structures of TNKS2 with compounds **51** (fuchsia), **52** (light blue), **40** (yellow), **41** (periwinkle). The residue Tyr1050 is indicated with a dashed circle to highlight its shift in crystal structures with diol-based compounds **51** and **52** in comparison to structures with compounds **40** and **41**.

**Table 3.**
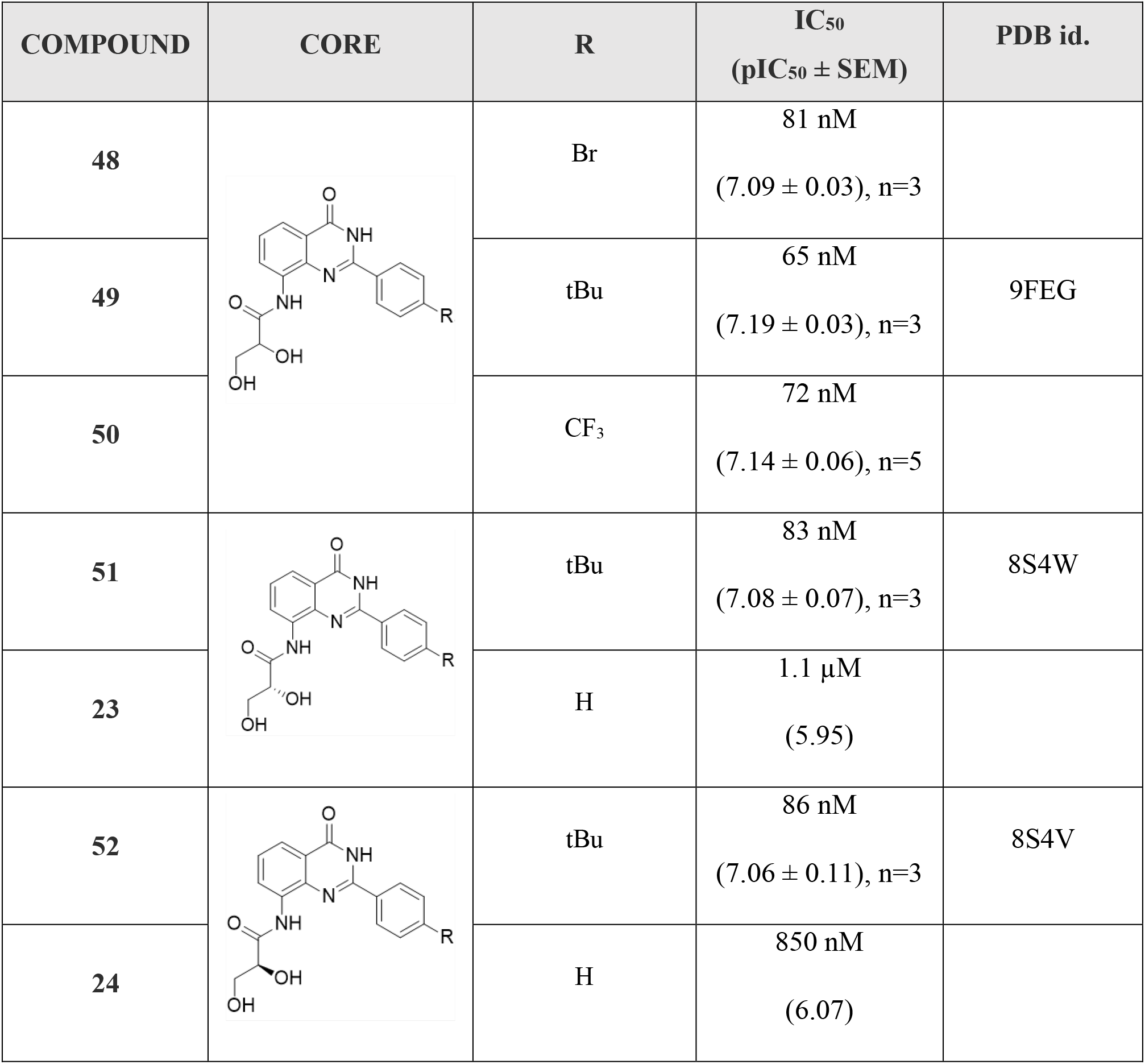
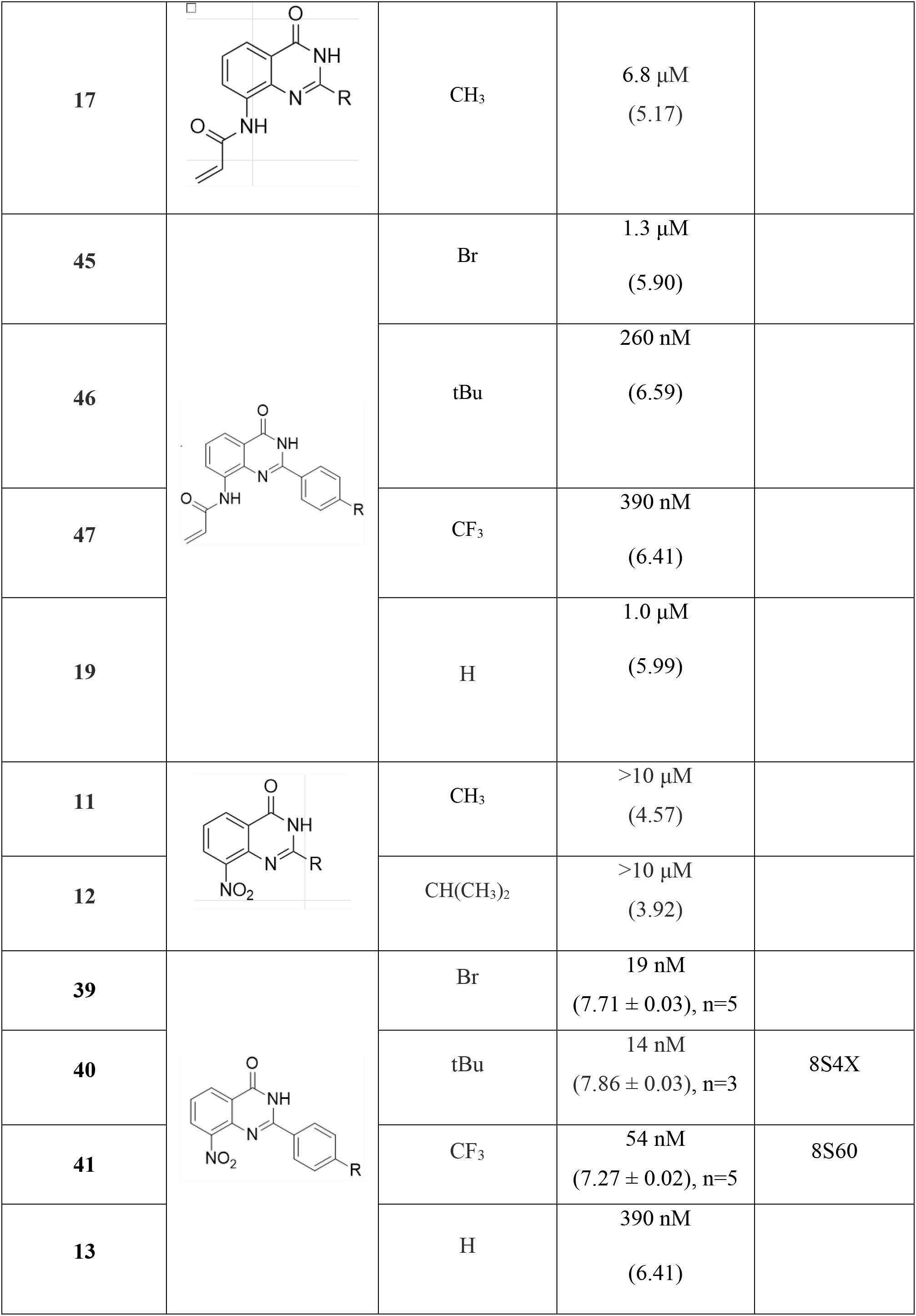
Structural analogues of 1. IC_50_ (pIC_50_ ± SEM, n=1-5) values with TNKS2 are reported.

| COMPOUND | CORE | R | IC<sub>50</sub><br>(pIC<sub>50</sub> ± SEM) | PDB id. |
| --- | --- | --- | --- | --- |
| 48 | | Br | 81 nM<br>(7.09 ± 0.03), n=3 | |
| 49 | | tBu | 65 nM<br>(7.19 ± 0.03), n=3 | 9FEG |
| 50 | | CF<sub>3</sub> | 72 nM<br>(7.14 ± 0.06), n=5 | |
| 51 | | tBu | 83 nM<br>(7.08 ± 0.07), n=3 | 8S4W |
| 23 | | H | 1.1 μM<br>(5.95) | |
| 52 | | tBu | 86 nM<br>(7.06 ± 0.11), n=3 | 8S4V |
| 24 |  | H | 850 nM<br>(6.07) |  |
| 17 |  | CH<sub>3</sub> | 6.8 μM<br>(5.17) |  |
| 45 |  | Br | 1.3 μM<br>(5.90) |  |
| 46 |  | tBu | 260 nM<br>(6.59) |  |
| 47 |  | CF<sub>3</sub> | 390 nM<br>(6.41) |  |
| 19 |  | H | 1.0 μM<br>(5.99) |  |
| 11 |  | CH<sub>3</sub> | >10 μM<br>(4.57) |  |
| 12 |  | CH(CH<sub>3</sub>)<sub>2</sub> | >10 μM<br>(3.92) |  |
| 39 |  | Br | 19 nM<br>(7.71 ± 0.03), n=5 |  |
| 40 |  | tBu | 14 nM<br>(7.86 ± 0.03), n=3 | 8S4X |
| 41 |  | CF<sub>3</sub> | 54 nM<br>(7.27 ± 0.02), n=5 | 8S60 |
| 13 |  | H | 390 nM<br>(6.41) |  |

Compounds **48**, **49** and **50** are diol-based analogues of compound **22** and exhibit up to a 10-fold increase in potency due to the *para* substitution of the phenyl ring at C-2 position. Notably, compound **49**, characterized by a *t*-Bu group, demonstrates the highest IC_50_ value (IC_50_ 65 nM) among this triad of inhibitors, followed by compound **50** (CF_3_ substituted, IC_50_ 72 nM) and compound **48** (Br substituted, 81 nM). Given the success of these inhibitors, we proceeded with the analysis of their enantiomeric analogues by studying their potencies and binding to TNKS2 NI pocket.

Crystal structures of compounds **51** and **52** with TNKS2 reveal that the polar diol substituent on C-8 extends towards Met1054 and packs against Tyr1060. The bulky diol causes a shift of Tyr1050 compared to its position in crystal structures of compounds **40** and **41** (**Fig. 2E**). The hydroxyl group of Tyr1050 moves away from the diol by 2.2 Å and 2.6 Å in comparison to structures of compounds **40** and **41**, respectively. The distal hydroxyl group of the diol establishes a new hydrogen bond with the main chain amide of Met1054. The proximal hydroxyl group of both enantiomers forms a hydrogen bond with Tyr1071. For compound **51** this hydrogen bond is 3.2 Å, while for compound **52** it measures 3.0 Å. Moreover, crystal structures show that the hydrophobic substituent in para-position of compounds **51** and **52** (*t*-Bu) positions itself in the hydrophobic pocket formed by Pro1034, Phe1035, Ile1051, Ile1075. IC_50_ measurements indicate that this functional group drastically increases the potency of compounds **51** (IC_50_ 83 nM) and **52** (IC_50_ 86 nM) in comparison to compounds **23** (IC_50_ 1.1 µM) and **24** (IC_50_ 850 nM). Structural data indicate that compounds **51** and **52** establish similar interactions with TNKS2, as confirmed by their matching potency measurements. In addition, the potency values of these compounds are comparable to the potency of compounds **48**, **49** and **50**, which include both enantiomers.

Crystal structures of compounds **40** and **41** show that the nitro group in position C-8 establishes a hydrogen bond with Tyr1071. The nitro group also faces Tyr1050 and Tyr1060. Both the hydrophobic *t*-Bu (compound **40**) and CF_3_ (compound **41**) substituents occupy the hydrophobic binding site and extend towards the solvent. The efficiency of interaction appears to be higher for compound **40**, as indicated by the potency measurements (IC_50_ 14 nM for compound **40**, IC_50_ 54 nM for compound **41**). Compound **39** is characterized by a single bromine atom in the *para* position and displays a potency (IC_50_ 19 nM) comparable to compound **40**. On the other hand, the absence of substituent on the phenyl ring in compound **13** hinders the potency (IC_50_ 390 nM) in comparison to compounds **39**, **40** and **41**. Compounds **11** and **12** are devoid of the phenyl ring and display potency values above 10 µM due to the absence of the T-shaped π-π stacking with Tyr1050.

Compounds **45**, **46**, **47** and **19** are characterized by an acrylamide substituent in the C-8 position and display lower potency compared to nitro-substituted compounds **39**, **40**, **41** and **13**, respectively, due to the absence of the hydrogen bond with Tyr1071. In addition, they show decreased IC_50_ values in comparison to the diol-based compounds. Notably, the acrylamide substituent contains an alkene group unable to form hydrogen bonds with Met1054 and Tyr1071 as the hydroxyl groups in the diol substituent. In line with other compounds potency tendencies, compound **46** exhibits the best potency (IC_50_ 260 nM) among acrylamide-based compounds, thanks to the *t*-Bu group in the *para*-position. Compound **46** is followed, in order of potency, by compound **47** (CF_3_ substituted, IC_50_ 390 nM), **45** (Br substituted, IC_50_ 1.3 µM) and **19** (no substitution, 1.0 µM).

### SAR of compounds with C-7 substituents

In general, the substituents at the C-7 position were not well tolerated, except for compounds **73**, **74** and **75**, which are diol-based. Compound **73** displays the best potency (IC_50_ 160 nM) due to its *t*-Bu substituent in *para*-position. Compound **74** is the second best tolerated compound (IC_50_ 420 nM), while compound **75** has mediocre potency (IC_50_ 2.8 µM) (**Table 4**). The reason behind this exception can be found in the flexibility of the diol moiety, which can adopt multiple conformations and potentially establish a hydrogen bond with Glu1138. In agreement with this analysis, compounds **64**, **65** and **66**, which are characterized by a nitro-group in C-7 position, are the least tolerated among all the studied compounds, possibly due to unfavorable interactions with the catalytic glutamate. Compounds **70**, **71**, **72** have an acrylamide-based substituent in the C-7 position and display potencies in the micromolar range. Compound **70** is the most tolerated compound (IC_50_ 1.0 µM), followed by compound **71** (IC_50_ 2.0 µM) and compound **72** (IC_50_ 5.1 µM). Despite being smaller, the acrylamide substituent is likely unable to form similar interactions with the protein as the diol group, explaining the drop in the potency.

**Table 4.**
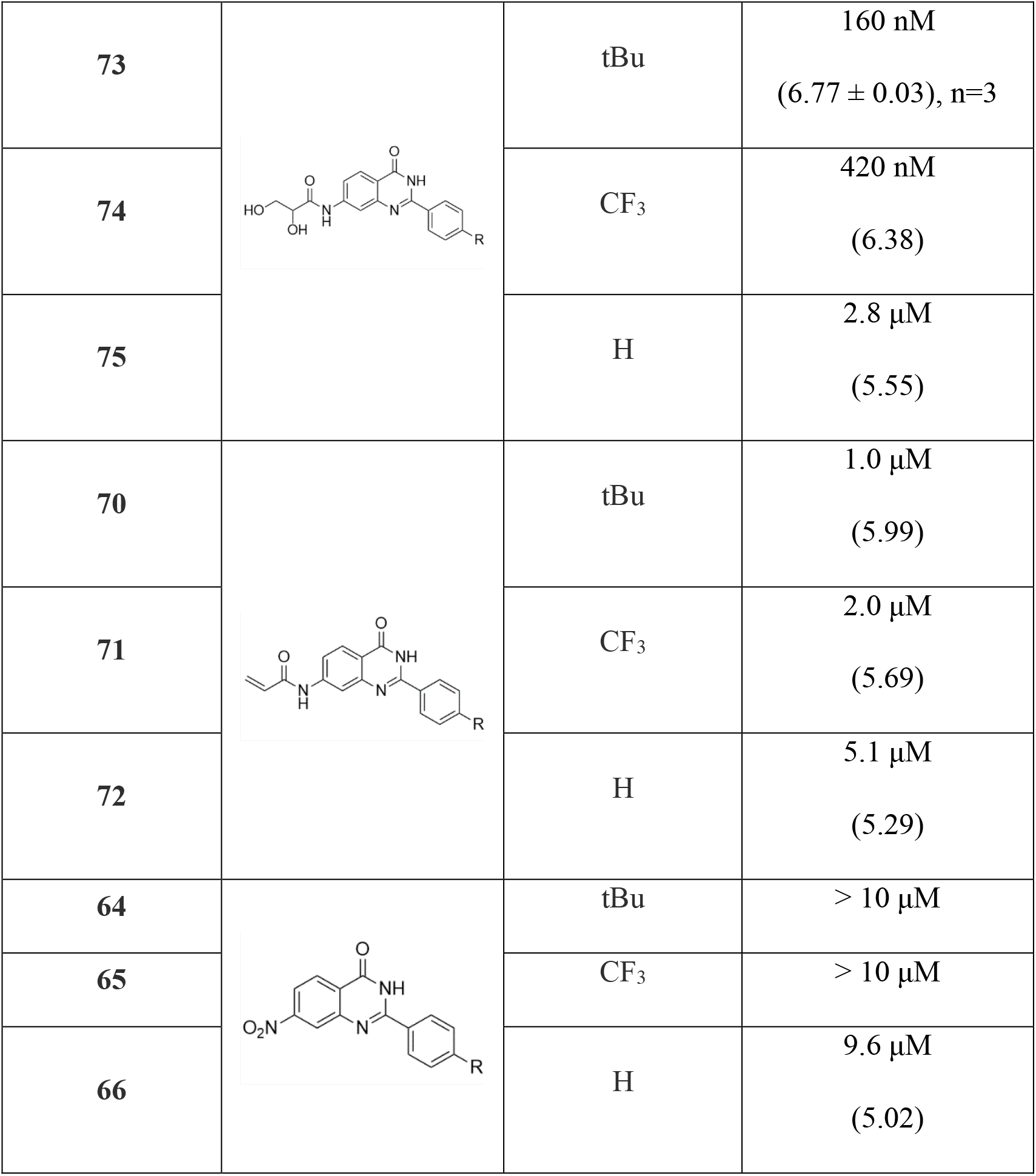
C-7 substituted structural analogues of 1. IC_50_ (pIC_50_ ± SEM, n=1 or 3) values with TNKS2 are reported.

|  |  |  |  |
| --- | --- | --- | --- |
| **73** |  | tBu | 160 nM<br>(6.77 ± 0.03), n=3 |
| **74** |  | CF<sub>3</sub> | 420 nM<br>(6.38) |
| **75** |  | H | 2.8 μM<br>(5.55) |
| **70** |  | tBu | 1.0 μM<br>(5.99) |
| **71** |  | CF<sub>3</sub> | 2.0 μM<br>(5.69) |
| **72** |  | H | 5.1 μM<br>(5.29) |
| **64** |  | tBu | > 10 μM |
| **65** |  | CF<sub>3</sub> | > 10 μM |
| **66** |  | H | 9.6 μM<br>(5.02) |

### Study of selectivity towards tankyrases

To confirm the selectivity of the compounds displaying the highest potency, we measured their inhibition against a panel of PARPs encompassing the PARylating enzymes (PARP1/2, TNKS1/2), as well as representative MARylating PARPs (PARP3/4/10/14/15). Interestingly, the diol substituent of **49** is tolerated and multiple PARPs are inhibited. There is clear selectivity over the PARP1-3, which display IC_50_ values in the micromolar range (**Table 5**), but **49** is also a potent inhibitor of PARP4, PARP10, PARP14 and PARP15. In contrast, **40** is extremely selective, with > 188-fold selectivity towards tankyrases over all the tested enzymes. Comparison of the available protein structures provides possible explanations for this behavior, since the D-loop of tankyrases includes a unique tyrosine (Tyr1050 in TNKS2) that provides the π-hole interaction with the nitro-group (**Fig. S1**). It is important to note that the D-loop shows sequence, length and flexibility variability across the ARTD family [55,92] (**Fig. S1**). In PARP1/2 the D-loop is more rigid and packs between the HD and ART domains, PARP4 has an extended loop and many of the MARylating enzymes vary in D-loop length and plasticity, but still lack a residue analogous to Tyr1050 of TNKS2 (**Fig. S1**). Overall, the D-loop can adopt multiple conformations, making the diol moiety of **49** tolerable.

**Table 5.** Profile of the selected compounds against the PARP enzymes with IC_50_ (pIC_50_ ± SEM, number of repetitions n indicated) and Wnt reporter assay activities reported as IC_50_ (n=3).

|  | **49** | **40** |
| --- | --- | --- |
| **PARP/TNKS profiling** |  |  |
| **PARP1** | 4.4 µM | >> 10 µM |
| **PARP2** | 3.6 µM | >> 10 µM |
| **PARP3** | 2.7 µM | >> 10 µM |
| **PARP4** | 230 nM<br>(6.64 ± 0.02), n=3 | >> 10 µM |
| **TNKS1** | 58 nM<br>(7.30 ± 0.13), n=4 | 53 nM<br>(7.31 ± 0.1), n=4 |
| **TNKS2** | 65 nM<br>(7.19 ± 0.03), n=3 | 14 nM<br>(7.86 ± 0.03), n=3 |
| **PARP10** | 220 nM<br>(6.65 ± 0.03), n=3 | >> 10 µM |
| **PARP14** | 250 nM<br>(6.60 ± 0.06), n=5 | >> 10 µM |
| **PARP15** | 340 nM<br>(6.47 ± 0.03), n=3 | >> 10 µM |
| **Activity in cellular context** |  |  |
| **Wnt/β-catenin reporter assay** | 600 nM | 520 nM |

To gain insight into compound **49** tolerability, we solved the co-crystal structure of PARP15 catalytic domain with the compound (**Fig. 3**). The crystal structure reveals that one enantiomer of the racemic mixture is selected in the crystal form, even if IC_50_ measurements of the two enantiomers **51** and **52** are equivalent with IC_50_ (pIC_50_ ± SEM) of 230 nM (6.65 ± 0.08) and 220 nM (6.67 ± 0.05), respectively. The diol substituent of the compound extends towards the D-loop and packs against Tyr591. The distal hydroxyl group of the diol forms hydrogen bonds with residues Ala583 and Ser585. At the same time, the compound is anchored to the NI site by establishing hydrogen bonds with Ser599 and Gly560. Interestingly, the proximal hydroxyl group of the diol is not engaged in any hydrogen bond with Tyr604 due to its distance from the residue (3.9 Å). This is a clear difference with TNKS2 crystal structures with compound **51** (PDB: 8S4W) and **52** (PDB: 8S4V), where the vicinity of Tyr1071 allows the participation of the proximal hydroxyl in a hydrogen bond (**Fig. 2**). With this crystal structure we confirm the absence of a residue in the D-loop that firmly packs against the compound in the same fashion as Tyr1050 in TNKS2. We also observed 2-Methyl-2,4-pentanediol (MPD), organic compound used for cryo-protection of the crystal, in the crystal structure with hydroxyl groups facing the compound (**Fig. 3**). In conclusion, this crystal structure attests the compatibility of the diol substituent at C-8 position with D-loops of different members of the ARTD family.

**Fig. 3.**
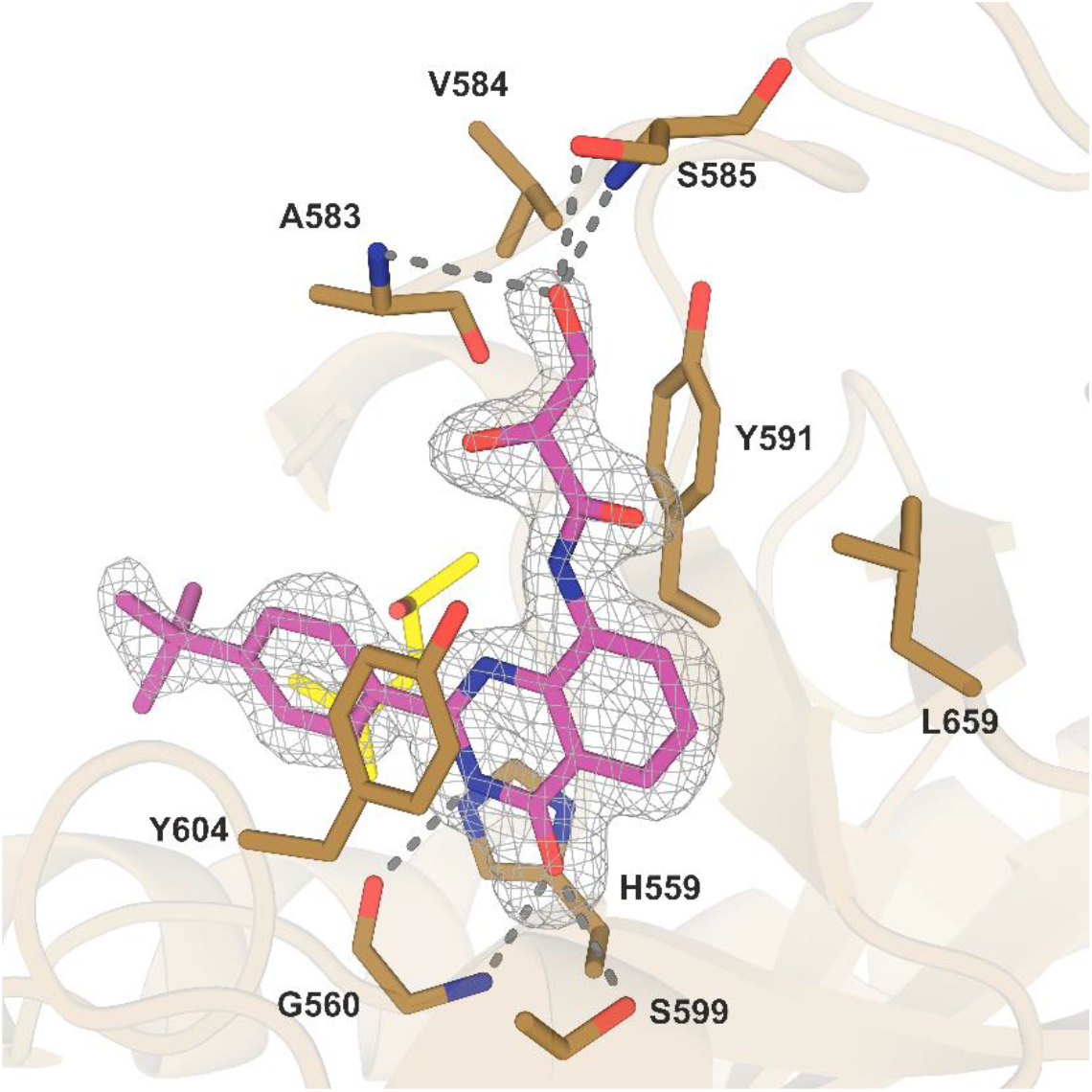
Crystal structures of compounds **49** with PARP15 (PDB: 9FEG). The sigma-A weighted 2Fo-Fc electron density map is contoured at 1.0 σ and colored gray. The dashed lines show the hydrogen bonds established with residues of the pocket. MPD is colored yellow, while the compound is depicted in color fuchsia.

### Efficacy in cell assays

To conclude the analysis of compounds **40** and **49**, we tested their inhibition in cells by using a luciferase-based Wnt/β-catenin signaling pathway reporter assay. In this assay, SuperTOP-Luciferase/Renilla HEK293 cells are employed [69]. Upon activation of the canonical Wnt signaling by presence of WNT3a in the medium, the expression of luciferase, which is under the control of a promoter with 7 TCF binding sites, is increased. This occurs since β-catenin binds to TCF/LEF transcription factors and activates transcription of downstream genes of the WNT pathway [93]. However, inhibition of tankyrases leads to increased levels of the β-catenin destruction complex, consequently to lower levels of β-catenin and to decreased expression of firefly luciferase [94]. Our experiment demonstrates that compounds **40** and **49** effectively impact the Wnt/β-catenin signaling pathway in a cellular context by inducing a reduction of firefly luciferase expression that translates into IC_50_ values of 520 nM and 600 nM, respectively (**Table 5**).

## Conclusions

In the present study, we explored the structural requirements for the inhibition of human TNKS1/2, focusing on the C-2, C-7 and C-8 positions of the quinazolin-4-one scaffold. Our findings offer valuable insights into the SAR in the context of tankyrase inhibition. At the C-2 position, our results indicate that the incorporation of a phenyl ring significantly enhances activity in comparison to the simple aliphatic substituents seen in compounds **8-15**, as it snuggly fits inside the hydrophobic pocket. Potency is further increased by introducing hydrophobic moieties at the *para*-position, with size appearing to be more crucial than electronic properties (*t*-Bu > Br > CF_3_ > H). Substituents at C-7 are not well-tolerated, possibly due to their proximity to the catalytic glutamate. In contrast, the C-8 position of the quinazolin-4-one is evidently more versatile and can accommodate a wide range of substituents with various physicochemical and electronic properties. Among the three series of compounds assessed, nitro-derivatives possess optimal inhibitory potency, possibly due to their size. Intriguingly, the bulkier diols display activity both in biochemical and cellular assays. Out of the 34 compounds tested, **40** and **49** exhibited significant inhibitory potency against tankyrases. Compound **40** showed high selectivity towards tankyrases over other PARP enzymes, while **49** also inhibited several PARPs at nanomolar potencies. Since both of analogues were active in cellular assays, they could provide further inspiration for optimization of the quinazolin-4-one scaffold as tankyrase and PARP inhibitors. These results may also be adopted for the optimization of other nicotinamide mimetics when carrying out hit/lead optimization of potential candidates for cancer treatment.

## Materials and Methods

### Chemistry

All commercially available reagents and solvents were purchased from Alfa Aesar and used without any further purification. Melting points were determined on a Büchi apparatus and were uncorrected. All NMR spectra were recorded on 400 or 600 MHz Bruker spectrometers respectively Avance™ DRX and III instruments (Bruker BioSpin GmbH – Rheinstetten, Germany). ^1^H NMR (400 and 600 MHz) and ^13^C NMR (101 and 151 MHz, recorded with complete proton decoupling) spectra were obtained with samples dissolved in CDCl_3_ or DMSO-*d_6_* with the residual solvent signals used as internal references: 7.26 ppm for CHCl_3_, and 2.50 ppm for (CD_3_)(CD_2_H)S(O) regarding ^1^H NMR experiments; 77.2 ppm for CDCl_3_ and 39.4 ppm for (CD_3_)_2_S(O) concerning ^13^C NMR experiments. Chemical shifts (δ) are given in ppm to the nearest 0.01 (^1^H) or 0.1 ppm (^13^C). The coupling constants (J) are given in Hertz (Hz). The signals are reported as follows: (s = singlet, d = doublet, t = triplet, m = multiplet, br = broad). Assignments of ^1^H and ^13^C NMR signals were unambiguously achieved with the help of D/H exchange and 2D techniques: COSY, NOESY, HMQC, and HMBC experiments. Flash chromatography was performed on Merck silica gel (40–63 μm) with the indicated solvent system using gradients of increasing polarity in most cases (Merck KGaA – Darmstadt, Germany). The reactions were monitored by analytical thin-layer chromatography (Merck pre-coated silica gel 60 F254 TLC plates, 0.25-mm layer thickness). Compounds were visualized on TLC plates by both UV radiation (254 and 365 nm). All solvents for absorption and fluorescence experiments were of spectroscopic grade.

### Synthesis of derivatives 11-75

#### 2-Methyl-8-nitroquinazolin-4(3H)-one (11)

A suspension of 2-amino-3-nitrobenzoic acid (546 mg, 3 mmol, ***1***) in (CH_3_CO)_2_O (3 mL, 31.74 mmol) was stirred for 20 h. The volatiles were then vacuum evaporated, and the residue was treated NH_3_ (0.5M solution in THF, 15 mL). After completion of the reaction, the solvent was vacuum evaporated, the resulting solid was dissolved in 5% aq. NaOH solution (10 mL) and refluxed for 10 minutes. After cooling, the reaction mixture was diluted with water and acidified with 9% aq. HCl solution (pH≈3). The precipitate was filtered, washed with water, and air-dried to give crude **11**, which was purified by column chromatography (silica gel) using a mixture of *c*Hex/EtOAc 1/2 as the eluent, to afford 400 mg (65%) of the title compound. Mp: 206-208 °C. ^1^H NMR (600 MHz, DMSO-*d_6_*) δ (ppm) 12.55 (brs, 1H, D_2_O exch., N*H*), 8.29 (dd, *J* = 7.9, 2.3 Hz, 1H, H-5), 8.21 (dd, *J* = 7.9, 2.2 Hz, 1H, H-7), 7.58 (t, *J* = 7.8 Hz, 1H, H-6), 2.37 (s, 3H, CH_3_); ^13^C NMR (151 MHz, DMSO-*d_6_*) δ (ppm) 160.3 (*C*O), 157.6 (C-2), 146.3 (C-8), 140.6 (C-8a),129.3 (C-5), 127.6 (C-7), 125.4 (C-6), 122.2 (C-4a), 21.8 (*C*H_3_); HRMS (ESI^+^) m/z 206.0565 (calcd for C_9_H_7_N_3_O_3_^+^, 206.0560).

#### 2-Isobutyl-8-nitroquinazolin-4(3H)-one (12)

To a solution of 2-amino-3-nitrobenzoic acid (992 mg, 5.45 mmol, ***1***) in anh. THF (20 mL) at 0 °C, Et_3_N (1.52 mL, 10.9 mmol) and 3-methylbutanoyl chloride (1.07 mL, 8.18 mmol) were added under argon. The resulting solution was stirred at rt for 36 h, after which the volatiles were vacuum evaporated, and the mixture was diluted with water and acidified with 9% aq. HCl solution (pH≈3). The precipitate was filtered and air-dried to give compound **3**. Without further purification, the solid was suspended in (CH_3_CO)_2_O (4 mL, 42.32 mmol) and the mixture was refluxed for 2 h. The volatiles were then vacuum evaporated, and the residue was treated with NH_3_ (0.5M solution in THF, 20 mL). Upon completion, the solvent was evaporated under reduced pressure, the residue was dissolved in 5% aq. NaOH solution (10 mL) and refluxed for 30 minutes. After cooling, the reaction mixture was diluted with water and acidified with 9% aq. HCl solution (pH≈3). The precipitate was filtered, washed with water, and air-dried to afford 950 mg (70.3 %) of the title compound **12**, practically pure, which was used for the next step without further purification. Mp: 222-224 °C. ^1^H NMR (600 MHz, DMSO-*d_6_*) δ (ppm) 12.56 (brs, 1H, D_2_O exch., N*H*), 8.30 (dd, *J* = 8.0, 1.4 Hz, 1H, H-5), 8.22 (dd, *J* = 7.8, 1.5 Hz, 1H, H-7), 7.59 (t, *J* = 7.9 Hz, 1H, H-6), 2.48 (d, *J* = 7.0 Hz, 2H, C*H*_2_), 2.13 (m, 1H, C*H*), 0.93 (d, *J* = 6.7 Hz, 6H, ((C*H*_3_)_2_); ^13^C NMR (151 MHz, DMSO-*d_6_*) δ (ppm) 160.3 (*C*O), 160.0 (C-2), 146.4 (C-8), 140.5 (C-8a), 129.3 (C-5), 127.6 (C-7), 125.5 (C-6), 122.3 (C-4a), 43.2 (*C*H_2_), 26.9 (*C*H), 22.0 ((*C*H_3_)_2_); HRMS (ESI^+^) m/z 248.1032 (calcd for C_12_H_13_N_3_O_3_^+^, 248.1030).

#### 8-Nitro-2-phenylquinazolin-4(3H)-one (13)

This compound was synthesized by an analogous procedure as described for the preparation of compound **12.** Yield: 92.7%. Mp: 180-182 °C. ^1^H NMR (600 MHz, DMSO-*d*_6_) δ (ppm) 12.93 (brs, 1H, D_2_O exch., N*H*), 8.39 (dd, *J* = 7.2, 1.2 Hz, 1H, H-5), 8.31 (dd, *J* = 8.1, 1.5 Hz, 1H, H-7), 8.16 (dd, *J* = 7.3, 1.8 Hz, 2H, H-2’, H-6’), 7.73 – 7.60 (m, 2H, H-6, H-4’), 7.58 (t, *J* = 7.6 Hz, 2H, H-3’, H-5’); ^13^C NMR (151 MHz, DMSO-*d*_6_) δ (ppm) 160.9 (*C*O), 154.6 (C-2), 146.7 (C-8), 140.5 (C-8a), 132.2 (C-4’), 131.92 (C-1’), 129.7 (C-5), 128.7 (C-3’, C-5’), 128.2 (C-2’, C-6’), 128.1 (C-7), 126.0, (C-6), 122.5 (C-4a); HRMS (ESI^+^) m/z 268.0720 (calcd for C_14_H_9_N_3_O_3_^+^, 268.0717).

#### N-(2-Methyl-4-oxo-3,4-dihydroquinazolin-8-yl)acrylamide (17)

A solution of **11** (520 mg, 1.31 mmol) in abs. EtOH (10 mL) was hydrogenated in the presence of Pd/C (15 mg), under pressure (50 psi), at rt, for 6 h. Upon completion, the mixture was filtered through a Celite pad, and the filtrate was evaporated to dryness to afford the amino derivative **14**. Without further purification, the residue was dissolved in anh. THF (10 mL) and to this solution were added Et_3_N (0.56 mL, 3.98 mmol) and 3-chloropropionyl chloride (190 μL, 1.98 mmol) and the resulting suspension was stirred for 15 minutes at rt. The volatiles were then vacuum evaporated, the residue was diluted with water and acidified with 9% aq. HCl solution (pH≈3), to give 95 mg (56%) of the title compound **17**, practically pure, which was used for the next step without further purification. Mp: 206-208 °C. ^1^H NMR (600 MHz, DMSO-*d_6_*) δ (ppm) 12.34 (brs, 1H, D_2_O exch., N*H*CO), 9.79 (brs, 1H, D_2_O exch., N*H*), 8.66 (dd, *J* = 7.9, 1.4 Hz, 1H, H-7), 7.77 (dd, *J* = 7.9, 1.4 Hz, 1H, H-5), 7.42 (t, *J* = 7.9 Hz, 1H, H-6), 6.79 (dd, *J* = 10.1, 1.7 Hz, 1H, COC*H*=CHH), 6.30 (dd, *J* = 16.9, 1.7 Hz, 1H, COCH=C*H*H), 5.80 (dd, *J* = 10.1, 1.7 Hz, 1H, COCH=CH*H*), 2.44 (s, 3H, C*H*_3_); ^13^C NMR (151 MHz, DMSO-*d_6_*) δ (ppm) 163.4 (NH*C*O), 161.4 (*C*O), 154.0 (C-2), 138.9 (C-8a), 133.3 (C-4a), 132.1 (CO*C*H=CHH), 127.3 (COCH=*C*HH), 125.7 (C-6), 123.1 (C-7), 120.5 (C-8), 119.8 (C-5), 21.6 (*C*H_3_); HRMS (ESI^+^) m/z 230.0930 (calcd for C_12_H_11_N_3_O_2_^+^, 230.0924).

#### N-(4-Oxo-2-phenyl-3,4-dihydroquinazolin-8-yl)acrylamide (19)

This compound was synthesized by an analogous procedure as described for the preparation of compound **17**. Yield: 40.8%. Mp: >230 °C. ^1^H NMR (400 MHz, DMSO-*d_6_*) δ (ppm) 12.63 (brs, 1H, D_2_O exch., N*H*), 9.92 (brs, 1H, D_2_O exch., N*H*CO), 8.71 (dd, *J* = 8.0, 1.4 Hz, 1H, H-7), 8.39 (dd, *J* = 7.3, 1.8 Hz, 2H, H-2’, H-6’), 7.86 (dd, *J* = 8.0, 1.4 Hz, 1H, H-5), 7.67 – 7.54 (m, 2H, H-3’, H-4’, H-5’), 7.50 (t, *J* = 8.0 Hz, 1H, H-6), 6.88 (ddd, *J* = 17.0, 10.2, 1.2 Hz, 1H, COC*H*=CHH), 6.34 (dd, *J* = 16.9, 1.8 Hz, 1H, COCH=C*H*H), 5.84 (dd, *J* = 10.2, 1.9 Hz, 1H, COCH=CH*H*); ^13^C NMR (151 MHz, DMSO-*d_6_*) δ (ppm) 163.4 (NH*C*O), 161.9 (*C*O), 151.7 (C-2), 138.9 (C-8a), 134.0 (C-4a), 132.3 (C-1’), 132.2 (CO*C*H=CHH), 131.6 (C-4’), 128.6 (C-3’, C-5’), 128.3, (C-2’, C-6’), 127.4 (COCH=*C*HH), 126.5 (C-6), 123.64 (C-7), 120.9 (C-8), 120.1 (C-5); HRMS (ESI^+^) m/z 292.1084 (calcd for C_17_H_13_N_3_O_2_^+^, 292.1081).

#### 2,3-Dihydroxy-N-(2-methyl-4-oxo-3,4-dihydroquinazolin-8-yl)propanamide (20)

To a solution of compound **17** (54 mg, 0.24 mmol) in THF (10 mL) were added osmium tetroxide (2.5% sol. in isopropanol) (90 μL, 0.008 mmol) and *N*-methyl-morpholine *N*-oxide (140 mg, 1.2 mmol), and the resulting mixture was stirred at rt for 3 days. Subsequently, a saturated NaHSO_3_ solution (0.5 mL) was added, and the mixture was stirred at rt for 60 minutes. Afterwards, the volatiles were vacuum evaporated, and the residue was extracted with EtOAc. The organic layer was washed with a saturated NaHCO_3_ solution, water, and brine, dried over anh. Na_2_SO_4_ and concentrated to dryness, to afford 30 mg (51.7%) of the title compound. Mp: 205-207 °C (EtOH). ^1^H NMR (600 MHz, DMSO-*d_6_*) δ (ppm) 12.36 (brs, 1H, D_2_O exch., N*H*), 10.49 (brs, 1H, D_2_O exch., N*H*CO), 8.75 (dd, *J* = 8.0, 1.4 Hz, 1H, H-7), 7.73 (dd, *J* = 8.0, 1.4 Hz, 1H, H-5), 7.41 (t, *J* = 8.0 Hz, 1H, H-6), 6.28 (d, *J* = 5.0 Hz, 1H, CHO*H*), 4.86 (t, *J* = 5.8 Hz, 1H, CH_2_O*H*), 4.13 (td, *J* = 5.1, 3.1 Hz, 1H, C*H*), 3.74 – 3.62 (m, 2H, C*H*_2_), 2.40 (s, 3H, C*H*_3_); ^13^C NMR (151 MHz, DMSO-*d_6_*) δ (ppm) 171.2 (NH*C*O), 161.4 (*C*O), 154.1 (C-2), 138.2 (C-8a), 132.9 (C-4a), 126.0 (C-6), 120.8 (C-7), 120.4 (C-8), 119.3 (C-5), 73.6 (*C*H), 63.7 (*C*H_2_), 21.8 (*C*H_3_); HRMS (ESI^+^) m/z 264.0985 (calcd for C_12_H_13_N_3_O_4_^+^, 264.0979).

#### 2,3-Dihydroxy-N-(2-isobutyl-4-oxo-3,4-dihydroquinazolin-8-yl)propanamide (21)

A solution of **12** (215 mg, 0.87 mmol) in abs. EtOH (10 mL) was hydrogenated in the presence of Pd/C (15 mg), under pressure (50 psi), at rt, for 6 h. Upon completion, the mixture was filtered through a Celite pad, and the filtrate was evaporated to dryness to afford the amino derivative **15**. Without further purification, the residue was dissolved in anh. THF (10 mL) and to this solution were added Et_3_N (0.36 mL, 2.61 mmol) and 3-chloropropionyl chloride (125 μL, 1.31 mmol) and the resulting suspension was stirred for 15 minutes at rt. The volatiles were then vacuum evaporated, the residue was diluted with water and acidified with 9% aq. HCl solution (pH≈3). The precipitate was filtered and air-dried to provide compound **18** practically pure. The solid was dissolved in THF (15 mL), followed by the addition of osmium tetroxide (2.5% sol. in isopropanol) (360 μL, 0.032 mmol) and *N*-methyl-morpholine *N*-oxide (560 mg, 4.8 mmol), and the resulting mixture was stirred at rt for 3 days. Afterwards, a saturated NaHCO_3_ solution (1 mL) was added, and the mixture was stirred at rt for 1 h. The volatiles were then vacuum evaporated, and the residue was extracted with EtOAc. The organic phase was washed with water and brine, dried over anh. Na_2_SO_4_ and concentrated to dryness, to afford 167 mg (62.9%) of the title compound. Mp: >230 °C (EtOAc). ^1^H NMR (600 MHz, DMSO-*d_6_*) δ (ppm) 12.32 (brs, 1H, D_2_O exch., N*H*), 10.56 (brs, 1H, D_2_O exch., N*H*CO), 8.75 (dd, *J* = 8.0, 1.4 Hz, 1H, H-7), 7.74 (dd, *J* = 8.0, 1.4 Hz, 1H, H-5), 7.42 (t, *J* = 8.0 Hz, 1H, H-6), 6.23 (d, *J* = 5.0 Hz, 1H, CHO*H*), 4.83 (t, *J* = 5.8 Hz, 1H, CH_2_O*H*), 4.12 (td, *J* = 5.2, 3.2 Hz, 1H, C*H*OH), 3.74 – 3.62 (m, 2H, C*H*_2_OH), 2.53 (d, *J* = 7.2, 2H, C*H*_2_CH(CH_3_)_2_), 2.28 (dp, *J* = 13.6, 6.8 Hz, 1H, CH_2_C*H*(CH_3_)_2_), 0.97 ( d, *J* = 6.7 Hz, 6H, (C*H*_3_)_2_); ^13^C NMR (151 MHz, DMSO-*d*_6_) δ (ppm) 171.1 (NH*C*O), 161.4 (*C*O), 156.4 (C-2), 138.1 (C-8a), 133.0 (C-4a), 125.9 (C-6), 120.6 (C-7), 120.5 (C-8), 119.1 (C-5), 73.5 (*C*HOH), 63.6 (*C*H_2_OH), 43.1 (*C*H_2_CH(CH_3_)_2_), 26.5 (CH_2_*C*H(CH_3_)_2_)), 22.1 ((*C*H_3_)_2_); HRMS (ESI^+^) m/z 306.1453 (calcd for C_15_H_19_N_3_O_4_^+^, 306.1448).

#### 2,3-Dihydroxy-N-(4-oxo-2-phenyl-3,4-dihydroquinazolin-8-yl)propanamide (22)

This compound was synthesized by an analogous procedure as described for the preparation of compound **20**. Yield: 72.3%. Mp: 214-216 °C. ^1^H NMR (600 MHz, DMSO-*d_6_*) δ (ppm) 12.68 (brs, 1H, D_2_O exch., N*H*), 10.78 (brs, 1H, D_2_O exch., N*H*CO), 8.78 (dd, *J* = 7.9, 1.4 Hz, 1H, Η-7), 8.29 (dd, *J* = 7.1, 1.9 Hz, 2H, Η-2΄, Η-6’), 7.81 (dd, *J* = 7.9, 1.3 Hz, 1H, Η-5), 7.63 (t, *J* = 7.3 Hz, 1H, Η-4΄), 7.58 (t, *J* = 7.5 Hz, 2H, Η-3’, Η-5’), 7.49 (t, *J* = 8.0 Hz, 1H, Η-6), 6.42 (d, *J* = 5.1 Hz, 1H CHO*H*), 4.93 (t, *J* = 5.8 Hz, 1H, CH_2_O*H*), 4.17 (td, *J* = 3.3 Hz, 1H, C*H*), 3.68-3.75 (m, 2H, C*H*_2_); ^13^C NMR (151 MHz, DMSO-*d_6_*) δ (ppm) 169.5 (NH*C*O), 161.8 (*C*O), 152.1 (C-2), 138.1 (C-8a), 132.9 (C-4a), 132.2 (C-1’), 131.8 (C-4’), 128.6 (C-3’, C-5’) 127.8 (C-2’, C-6’), 126.7 (C-6), 121.1 (C-7), 120.8 (C-8), 119.9 (C-5), 74.9 (*C*H), 67.1 (*C*H_2_); HRMS (ESI^+^) m/z 326.1140 (calcd for C_17_H_15_N_3_O_4_^+^, 326.1135).

#### (S)-2,3-Dihydroxy-N-(4-oxo-2-phenyl-3,4-dihydroquinazolin-8-yl)propanamide (23)

AD-mix-α (160 mg) was added to a mixture of *t*-BuOH/H_2_O (3:3 mL) and the suspension was stirred at rt for 1 h. Compound **19** (30 mg, 0.12 mmol) was then added at 0 °C, and the reaction mixture was stirred for 24 h. Upon completion, a saturated NaHSO_3_ solution (0.5 mL) was added, and the mixture was stirred at rt for 60 minutes. Afterwards, the volatiles were vacuum evaporated, and the residue was extracted with EtOAc. The organic layer was washed water and brine, dried over anh. Na_2_SO_4_ and concentrated to dryness. Flash chromatography on silica gel using a mixture of *n*-Hex/THF 3/1 – 1/2 as the eluent, to afford 29 mg (74.4%) of the title compound. Mp: 292 °C. ^1^H NMR (600 MHz, DMSO-*d_6_*) δ (ppm) 12.68 (brs, 1H, D_2_O exch., N*H*), 10.78 (brs, 1H, D_2_O exch., N*H*CO), 8.78 (dd, *J* = 7.9, 1.4 Hz, 1H, Η-7), 8.29 (dd, *J* = 7.1, 1.9 Hz, 2H, Η-2΄, Η-6’), 7.81 (dd, *J* = 7.9, 1.3 Hz, 1H, Η-5), 7.63 (t, *J* = 7.3 Hz, 1H, Η-4΄), 7.58 (t, *J* = 7.5 Hz, 2H, Η-3’, Η-5’), 7.49 (t, *J* = 8.0 Hz, 1H, Η-6), 6.42 (d, *J* = 5.1 Hz, 1H CHO*H*), 4.93 (t, *J* = 5.8 Hz, 1H, CH_2_O*H*), 4.17 (d, *J* = 3.3 Hz, 1H, C*H*), 3.68-3.75 (m, 2H, C*H*_2_); ^13^C NMR (151 MHz, DMSO-*d_6_*) δ (ppm) 169.5 (NH*C*O), 161.8 (*C*O), 152.1 (C-2), 138.1 (C-8a), 132.9 (C-4a), 132.2 (C-1’), 131.8 (C-4’), 128.6 (C-3’, C-5’) 127.8 (C-2’, C-6’), 126.7 (C-6), 121.1 (C-7), 120.8 (C-8), 119.9 (C-5), 74.9 (*C*H), 67.1 (*C*H_2_); HRMS (ESI^+^) m/z 326.1140 (calcd for C_17_H_15_N_3_O_4_^+^, 326.1135).

#### (R)-2,3-Dihydroxy-N-(4-oxo-2-phenyl-3,4-dihydroquinazolin-8-yl)propanamide (24)

This compound was synthesized by an analogous procedure as described for the preparation of compound **23**. Yield: 79.5%. Mp: 291 °C. ^1^H NMR (600 MHz, DMSO-*d_6_*) δ (ppm) 12.68 (brs, 1H, D_2_O exch., N*H*), 10.78 (brs, 1H, D_2_O exch., N*H*CO), 8.78 (dd, *J* = 7.9, 1.4 Hz, 1H, Η-7), 8.29 (dd, *J* = 7.1, 1.9 Hz, 2H, Η-2΄, Η-6’), 7.81 (dd, *J* = 7.9, 1.3 Hz, 1H, Η-5), 7.63 (t, *J* = 7.3 Hz, 1H, Η-4΄), 7.58 (t, *J* = 7.5 Hz, 2H, Η-3’, Η-5’), 7.49 (t, *J* = 8.0 Hz, 1H, Η-6), 6.42 (d, *J* = 5.1 Hz, 1H CHO*H*), 4.93 (t, *J* = 5.8 Hz, 1H, CH_2_O*H*), 4.17 (d, *J* = 3.3 Hz, 1H, C*H*), 3.68-3.75 (m, 2H, C*H*_2_); ^13^C NMR (151 MHz, DMSO-*d_6_*) δ (ppm) 169.5 (NH*C*O), 161.8 (*C*O), 152.1 (C-2), 138.1 (C-8a), 132.9 (C-4a), 132.2 (C-1’), 131.8 (C-4’), 128.6 (C-3’, C-5’) 127.8 (C-2’, C-6’), 126.7 (C-6), 121.1 (C-7), 120.8 (C-8), 119.9 (C-5), 74.9 (*C*H), 67.1 (*C*H_2_); HRMS (ESI^+^) m/z 326.1142 (calcd for C_17_H_15_N_3_O_4_^+^, 326.1135).

#### 2-(4-Bromobenzamido)-3-nitrobenzoic acid (30)

To a solution of 4-bromobenzoic acid (4.42 g, 22 mmol, ***25***) in anh. THF (20 mL) at 0 °C, Et_3_N (4.2 mL, 30 mmol) and ClCO_2_Et (2.1 mL, 22 mmol) were added under argon. The resulting mixture was stirred for 30 minutes at rt, after which Na_2_CO_3_ (3.18 g, 30 mmol) and 2-amino-3-nitrobenzoic acid (1.82 g, 10 mmol, ***1***) were added at rt, and the reaction mixture was stirred for 24 h. Upon completion, the volatiles were vacuum evaporated, the residue was diluted with water and acidified with 9% aq. HCl solution (pH≈3). The mixture was extracted with EtOAc and the organic layer was washed with water and brine, dried over anh. Na_2_SO_4_ and evaporated to dryness. The crude solid was dissolved in THF (20 mL), followed by the addition of an aq. K_2_CO_3_ solution (6g, 44 mmol, in 25 mL H_2_O). The mixture was stirred at rt for 12 h, after which it was diluted with water and acidified with 9% aq. HCl solution (pH≈4). The precipitate was filtered and air-dried to give crude **30**, which was purified by column chromatography (silica gel) using a mixture of *c*Hex/CH_2_Cl_2_/CH_3_COOH 90/10/0.1 - CH_2_Cl_2_/MeOH/CH_3_COOH 98/2/0.1 as the eluent, to afford 2.1 g (57.6%) of the title compound. Mp: 194 °C. ^1^H NMR (400 MHz, DMSO-*d*_6_) δ 11.69 (brs, D_2_O exch., 1H, COO*H*), 8.26 (dd, *J* = 7.8, 1.6 Hz, 1H, H-6), 7.97 (d, *J* = 7.8 Hz, 1H, H-4), 7.90 (d, *J* = 8.7 Hz, 2H, H-2’, H-6’), 7.79 (d, *J* = 8.4 Hz, 2H, H-3’, H-5’), 7.36 (t, *J* = 8.2, 7.4 Hz, 1H, H-5); ^13^C NMR (101 MHz, DMSO-*d*_6_) δ 167.6 (COOH), 164.4 (NHCO), 144.6 (C-3), 135.5 (C-6), 133.3 (C-2), 132.5 (C-1’), 132.3 (C-3’, C-5’), 130.1 (C-2’, C-6’), 129.7 (C-1), 127.2 (C-4), 126.6 (C-4’), 124.2 (C-5).

#### 2-(4-(tert-Butyl)benzamido)-3-nitrobenzoic acid (31)

This compound was synthesized by an analogous procedure as described for the preparation of compound **30**. Yield: 35.4%. Μp: 211 °C. ^1^H NMR (600 MHz, Acetone-*d*_6_) δ 11.58 (brs, D_2_O exch., 1H, COO*H*), 8.46 (dd, *J* = 7.9, 1.6 Hz, 1H, H-6), 8.24 (dd, *J* = 8.2, 1.6 Hz, 1H, H-4), 7.98 (d, *J* = 8.8 Hz, 2H, H-2’, H-6’), 7.65 (d, *J* = 8.8 Hz, 2H, H-3’, H-5’), 7.55 (t, *J* = 8.0 Hz, 1H, H-5), 1.39 (s, 9H, (C*H_3_*)_3_); ^13^C NMR (151 MHz, Acetone-*d*_6_) δ 167.9 (*C*OOH), 164.1 (NH*C*O), 156.2 (C-4’), 144.8 (C-3), 135.3 (C-6), 133.3 (C-2), 130.7 (C-1’), 129.7 (C-4), 127.5 (C-2’, C-6’), 125.9 (C-3’, C-5’), 123.9 (C-5), 122.3 (C-1), 34.8 (*C*(CH_3_)_3_), 30.5 ((*C*H_3_)_3_).

#### 3-Nitro-2-(4-(trifluoromethyl)benzamido)benzoic acid (32)

To a solution of 2-amino-3-nitrobenzoic acid (350 mg, 1.92 mmol, ***1***) in anh. THF (15 mL) at 0 °C, Et_3_N (0.6 mL, 4.22 mmol) and 4-(trifluoromethyl)benzoyl chloride (320 μL, 2.16 mmol) were added under argon. The resulting mixture was stirred at rt for 3 h, after which, it was diluted with water and acidified with 9% aq. HCl solution (pH≈3). The precipitate was filtered and air-dried to give crude **32**, which was purified by column chromatography (silica gel) using a mixture of *c*Hex/CH_2_Cl_2_/CH_3_COOH 50/50/0.1 - CH_2_Cl_2_/MeOH/CH_3_COOH 95/5/0.1, to afford 285 mg (41.9%) of the title compound. Mp: 191 °C. ^1^H NMR (400 MHz, DMSO-*d*_6_) δ 11.18 (brs, D_2_O exch., 1H, COO*H*), 8.23 (dd, *J* = 7.8, 1.6 Hz, 1H, H-6), 8.20 (dd, *J* = 8.2, 1.6 Hz, 1H, H-4), 8.15 (d, *J* = 8.1 Hz, 2H, H-2’, H-6’), 7.98 (d, *J* = 8.3 Hz, 2H, H-3’, H-5’), 7.61 (t, *J* = 8.0 Hz, 1H, H-5); ^13^C NMR (101 MHz, DMSO-*d*_6_) δ 167.2, 164.6, 146.3, 137.5, 135.4, 132.7, 132.4, 132.1, 130.8, 129.0, 128.8, 128.6, 126.9, 126.4, 126.3, 125.6, 122.9.

#### 2-(4-Bromophenyl)-8-nitroquinazolin-4(3H)-one (39)

A suspension of compound **30** (1.6 g, 4.4 mmol) in (CH_3_CO)_2_O (4 mL, 42.32 mmol) was refluxed for 2 h to obtain the necessary intermediate **26**. The volatiles were then vacuum evaporated and NH_3_ (0.5M solution in THF, 15 mL) was added to the flask. The mixture was stirred at rt for 12 h, after which, it was diluted with water and acidified with 9% aq. HCl solution (pH≈3). The precipitate was filtered, washed with water, and air-dried to give crude **39**, which was purified by trituration with THF (10 mL), to afford 967 mg (63.7%) of the title compound. Mp: 297.6 °C (dec.). ^1^H NMR (400 MHz, DMSO-*d*_6_) δ 13.01 (brs, D_2_O exch., 1H, Ν*Η*), 8.38 (dd, *J* = 8.0, 1.5 Hz, 1H, H-5), 8.32 (dd, *J* = 7.8, 1.5 Hz, 1H, H-7), 8.09 (d, *J* = 8.9 Hz, 2H, H-2’, H-6’), 7.80 (d, *J* = 8.8 Hz, 2H, H-3’, H-5’), 7.66 (t, *J* = 7.9 Hz, 1H, H-6); ^13^C NMR (101 MHz, DMSO-*d*_6_) δ 161.3 (*C*O), 154.3 (C-2), 147.1 (C-8), 140.9 (C-8a), 132.3 (C-3’, C-5’), 131.6 (C-1’), 130.6 (C-2’, C-6’), 130.2 (C-5), 128.8 (C-7), 126.7 (C-4’), 126.6 (C-6), 123.0 (C-4a); HRMS (ESI^+^) m/z 345.9823 (calcd for C_14_H_8_BrN_3_O_3_^+^, 345.9822).

#### 2-(4-(tert-Butyl)phenyl)-8-nitroquinazolin-4(3H)-one (40)

This compound was synthesized by an analogous procedure as described for the preparation of compound **39**. Yield: 63.5%. Mp: 289 °C. ^1^H NMR (400 MHz, DMSO-*d*_6_) δ 12.90 (brs, D_2_O exch., 1H, N*H*), 8.37 (dd, *J* = 8.0, 1.5 Hz, 1H, H-5), 8.30 (dd, *J* = 7.8, 1.5 Hz, 1H, H-7), 8.10 (d, *J* = 9.1 Hz, 2H, Η-2’, Η-6’), 7.63 (t, *J* = 7.9 Hz, 1H, Η-6), 7.59 (d, *J* = 8.7 Hz, 2H, Η-3’, Η-5’), 1.32 (s, 9H, (C*H*_3_)_3_); ^13^C NMR (101 MHz, DMSO-*d*_6_) δ 161.4 (*C*O), 155.8 (C-4’), 155.0 (C-2), 147.1 (C-8), 141.1 (C-8a), 130.2 (C-5), 129.7 (C-1’), 128.7 (C-7), 128.5 (C-2’, C-6’), 126.3 (C-6), 126.1 (C-3’, C-5’), 122.9 (C-4a), 35.3 (*C*(CH_3_)_3_), 31.3 ((*C*H_3_)_3_); HRMS (ESI^+^) m/z 324.1347 (calcd for C_18_H_17_N_3_O_3_^+^, 324.1343).

#### 8-Nitro-2-(4-(trifluoromethyl)phenyl)quinazolin-4(3H)-one (41)

This compound was synthesized by an analogous procedure as described for the preparation of compound **39**. Yield: 87.8%. Mp: 305 °C. ^1^H NMR (400 MHz, DMSO-*d*_6_) δ 13.14 (brs, D_2_O exch., 1H, Ν*Η*), 8.39 (dd, *J* = 8.0, 1.5 Hz, 1H, H-5), 8.35 – 8.29 (m, 3H, H-7, H-2’, H-6’), 7.95 (d, *J* = 8.8 Hz, 2H, H-3’, H-5’), 7.68 (t, *J* = 7.9 Hz, 1H, H-6); ^13^C NMR (151 MHz, DMSO-*d*_6_) δ 161.3, 154.0, 147.2, 140.7, 136.4, 132.5, 132.3, 132.1, 131.9, 130.2, 129.5, 128.8, 127.0, 126.1, 125.2, 123.4, 123.2; HRMS (ESI^+^) m/z 336.0599 (calcd for C_15_H_8_F_3_N_3_O_3_^+^, 336.0591).

#### N-(2-(4-Bromophenyl)-4-oxo-3,4-dihydroquinazolin-8-yl)acrylamide (45)

A suspension of **39** (150 mg, 0.44 mmol) in abs. EtOH (10 mL) was hydrogenated in the presence of Raney Ni (10 mg), under atmospheric pressure, at rt, for 20 h. Subsequently, the mixture was filtered through a Celite pad, and evaporated to dryness to give the amino derivative **42**, which was dissolved, without further purification, in anh. THF (10 mL). To this solution were then added Et_3_N (0.26 mL, 1.83 mmol) and acryloyl chloride (70 μL, 0.87 mmol) under argon, and the resulting suspension was stirred for 16 h at rt. Upon completion, the volatiles were vacuum evaporated, the residue was diluted with water and acidified with 9% aq. HCl solution (pH≈3). The precipitate was filtered, air-dried, and purified by column chromatography (silica gel) using a mixture of *n*-Hex/CH_2_Cl_2_ 3/1 - CH_2_Cl_2_/MeOH 100/0.5, to afford 72 mg (44.4%) of the title compound. Mp: 323 °C (dec.). ^1^H NMR (400 MHz, DMSO-*d*_6_) δ 12.74 (brs, D_2_O exch., 1H, N*H*), 9.94 (brs, D_2_O exch., 1H, N*H*CO), 8.72 (d, *J* = 7.9 Hz, 1H, H-7), 8.35 (d, *J* = 8.2 Hz, 2H, H-2’, H-6’), 7.86 (d, *J* = 7.9 Hz, 1H, H-5), 7.79 (d, *J* = 8.3 Hz, 2H, H-3’, H-5’), 7.51 (t, *J* = 8.0 Hz, 1H, H-6), 6.89 (dd, *J* = 17.0, 10.2 Hz, 1H, C*H*=CHH), 6.34 (dd, *J* = 16.9, 1.9 Hz, 1H, CH=C*H*H), 5.84 (dd, *J* = 10.1, 1.9 Hz, 1H, CH=CH*H*); ^13^C NMR (101 MHz, DMSO-*d*_6_) δ 163.9 (NH*C*O), 162.4 (*C*O), 151.3 (C-2), 139.2 (C-8a), 134.6 (C-4a), 132.7 (*C*H=CHH), 132.0 (C-3’, C-5’, C-1’), 130.8 (C-2’, C-6’), 127.9 (CH=*C*HH), 127.2 (C-6), 126.1 (C-4’), 124.3 (C-7), 121.5 (C-8), 120.6 (C-5); HRMS (ESI^+^) m/z 370.0184 (calcd for C_17_H_12_BrN_3_O_2_^+^, 370.0186).

#### N-(2-(4-(tert-Butyl)phenyl)-4-oxo-3,4-dihydroquinazolin-8-yl)acrylamide (46)

This compound was synthesized by an analogous procedure as described for the preparation of compound **17**. Yield: 74.5%. Mp: 298 °C. ^1^H NMR (400 MHz, DMSO-*d*_6_) δ 12.63 (brs, D_2_O exch., 1H, N*H*), 9.91 (brs, D_2_O exch., 1H, N*H*CO), 8.71 (dd, *J* = 8.0, 1.7 Hz, 1H, H-7), 8.32 (d, *J* = 8.3 Hz, 2H, H-2’, H-6’), 7.85 (dd, *J* = 7.9, 1.4 Hz, 1H, H-5), 7.59 (d, *J* = 8.3 Hz, 2H, H-3’, H-5’), 7.49 (t, *J* = 8.0 Hz, 1H, H-6), 6.87 (dd, *J* = 16.9, 10.2 Hz, 1H, C*H*=CHH), 6.34 (dd, *J* = 16.9, 1.8 Hz, 1H, CH=CH*H*), 5.84 (dd, *J* = 10.2, 1.8 Hz, 1H, CH=C*H*H), 1.35 (s, 9H, (C*H*_3_)_3_); ^13^C NMR (101 MHz, DMSO-*d*_6_) δ 163.9 (NH*C*O), 162.4 (*C*O), 155.1 (C-4’), 152.1 (C-2), 139.4 (C-8a), 134.4 (C-4a), 132.7 (*C*H=CHH), 130.1 (C-1’), 128.6 (C-2’, C-6’), 127.8 (CH=*C*HH), 126.8 (C-6), 125.9 (C-3’,C-5’), 124.0 (C-7), 121.3 (C-8), 120.5 (C-5), 35.2 (*C*(CH_3_)_3_), 31.4 ((*C*H_3_)_3_); HRMS (ESI^+^) m/z 348.1714 (calcd for C_21_H_21_N_3_O_2_^+^, 348.1707).

#### N-(4-Oxo-2-(4-(trifluoromethyl)phenyl)-3,4-dihydroquinazolin-8-yl)acrylamide (47)

This compound was synthesized by an analogous procedure as described for the preparation of compound **17**. Yield: 84.5%. Mp: 320 °C (dec.). ^1^H NMR (400 MHz, DMSO-*d*_6_) δ 12.89 (brs, D_2_O exch., 1H, N*H*), 9.97 (brs, D_2_O exch., 1H, N*H*CO), 8.73 (dd, *J* = 8.1, 1.4 Hz, 1H, H-7), 8.58 (d, *J* = 8.2 Hz, 2H, H-2’, H-6’), 7.94 (d, *J* = 8.2 Hz, 2H, H-3’, H-5’), 7.88 (dd, *J* = 8.0, 1.4 Hz, 1H, H-5), 7.54 (t, *J* = 8.0 Hz, 1H, H-6), 6.88 (dd, *J* = 17.0, 10.2 Hz, 1H, C*H*=CHH), 6.34 (dd, *J* = 17.0, 1.8 Hz, 1H, CH=CH*H*), 5.85 (dd, *J* = 10.2, 1.8 Hz, 1H, CH=C*H*H); ^13^C NMR (101 MHz, DMSO-*d*_6_) δ 164.0, 162.3, 151.0, 139.1, 136.7, 134.7, 132.6, 131.9, 131.6, 129.7, 128.0, 127.5, 125.9, 124.4, 123.1, 121.7, 120.6; HRMS (ESI^+^) m/z 360.0964 (calcd for C_18_H_12_F_3_N_3_O_2_^+^, 360.0954).

#### N-(2-(4-Bromophenyl)-4-oxo-3,4-dihydroquinazolin-8-yl)-2,3-dihydroxypropanamide (48)

This compound was synthesized by an analogous procedure as described for the preparation of compound **20**. Yield: 53.2%. Mp: 314 °C. ^1^H NMR (400 MHz, DMSO-*d*_6_) δ 12.77 (brs, D_2_O exch., 1H, N*H*), 10.74 (brs, D_2_O exch., 1H, N*H*CO), 8.78 (dd, *J* = 7.9, 1.4 Hz, 1H, H-7), 8.20 (d, *J* = 8.7 Hz, 2H, Η-2΄, Η-6’), 7.85 – 7.76 (m, 3H, H-5, H-3’, H-5’), 7.51 (t, *J* = 8.0 Hz, 1H, H-6), 6.40 (d, *J* = 5.3 Hz, 1H, CHO*H*), 4.96 (t, *J* = 5.8 Hz, 1H, CH_2_O*H*), 4.18 (td, *J* = 5.0, 3.3 Hz, 1H, C*H*), 3.78 – 3.67 (m, 2H, C*H*_2_); ^13^C NMR (101 MHz, DMSO-*d*_6_) δ 171.7 (NH*C*O), 162.4 (*C*O), 151.4 (C-2), 138.3 (C-8a), 134.1 (C-4a), 132.2 (C-3’, C-5’), 132.1 (C-1’), 130.3 (C-2’, C-6’), 127.4 (C-6), 126.1 (C-4’), 121.4 (C-7), 121.3 (C-8), 119.8 (C-5), 74.0 (*C*H), 64.0 (*C*H_2_); HRMS (ESI^+^) m/z 404.0241 (calcd for C_17_H_14_BrN_3_O_4_^+^, 404.0240).

#### N-(2-(4-(tert-Butyl)phenyl)-4-oxo-3,4-dihydroquinazolin-8-yl)-2,3-dihydroxypropanamide (49)

This compound was synthesized by an analogous procedure as described for the preparation of compound **20**. Yield: 84.3%. Mp: 218 °C. ^1^H NMR (400 MHz, DMSO-*d*_6_) δ 12.69 (brs, D_2_O exch., 1H, N*H*), 10.79 (brs, D_2_O exch., 1H, N*H*CO), 8.78 (dd, *J* = 7.9, 1.4 Hz, 1H, H-7), 8.23 (d, *J* = 8.6 Hz, 2H, Η-2΄, Η-6’), 7.81 (dd, *J* = 8.0, 1.4 Hz, 1H, H-5), 7.59 (d, *J* = 8.5 Hz, 2H, H-3’, H-5’), 7.48 (t, *J* = 8.0 Hz, 1H, H-6), 6.41 (d, *J* = 5.2 Hz, 1H, CHO*H*), 4.94 (t, *J* = 5.8 Hz, 1H, CH_2_O*H*), 4.17 (dd, *J* = 5.2, 3.4 Hz, 1H, C*H*), 3.72 (q, *J* = 6.1, 5.0 Hz, 2H, C*H*_2_), 1.35 (s, 9H (CH_3_)_3_); ^13^C NMR (101 MHz, DMSO-*d*_6_) δ 171.7 (NH*C*O), 162.5 (*C*O), 155.2 (C-4’), 152.0 (C-2), 138.5 (C-8a), 134.0 (C-4a), 129.9 (C-1’), 128.1 (C-2’, C-6’), 127.1 (C-6), 126.0 (C-3’, C-5’), 121.3 (C-7), 119.8 (C-5), 74.1 (*C*H), 64.1 (*C*H_2_), 35.2 (*C*(CH_3_)_3_), 31.4 ((*C*H_3_)_3_); HRMS (ESI^+^) m/z 382.1767 (calcd for C_21_H_23_N_3_O_4_^+^, 382.1761).

#### 2,3-Dihydroxy-N-(4-oxo-2-(4-(trifluoromethyl)phenyl)-3,4-dihydroquinazolin-8-yl)propanamide (50)

This compound was synthesized by an analogous procedure as described for the preparation of compound **20**. Yield: 69.1%. Mp: 302 °C. ^1^H NMR (400 MHz, DMSO-*d*_6_) δ 12.94 (brs, D_2_O exch., 1H, N*H*), 10.74 (brs, D_2_O exch., 1H, N*H*CO), 8.79 (dd, *J* = 8.0, 1.4 Hz, 1H, H-7), 8.44 (d, *J* = 8.2 Hz, 2H, H-2’, H-6’), 7.95 (d, *J* = 8.3 Hz, 2H, H-3’, H-5’), 7.83 (dd, *J* = 8.0, 1.4 Hz, 1H, H-5), 7.53 (t, *J* = 8.0 Hz, 1H, H-6), 6.39 (d, *J* = 5.3 Hz, 1H, CHO*H*), 4.97 (t, *J* = 5.7 Hz, 1H, CH_2_O*H*), 4.18 (td, *J* = 5.0, 3.5 Hz, 1H, C*H*), 3.72 (dt, *J* = 6.2, 4.1 Hz, 2H, C*H*_2_); ^13^C NMR (101 MHz, DMSO-*d*_6_) δ 171.7, 162.3, 151.0, 138.1, 136.7, 134.2, 132.3, 132.0, 131.7, 131.4, 129.2, 127.8, 126.1, 125.7, 123.0, 121.5, 121.5, 119.8, 74.0, 64.0; HRMS (ESI^+^) m/z 394.1007 (calcd for C_18_H_14_F_3_N_3_O_4_^+^, 394.1009).

#### (S)-N-(2-(4-(tert-Butyl)phenyl)-4-oxo-3,4-dihydroquinazolin-8-yl)-2,3-dihydroxypropanamide (51)

This compound was synthesized by an analogous procedure as described for the synthesis of compound **23**. Yield 64%. Mp: 287 °C. ^1^H NMR (400 MHz, DMSO-*d*_6_) δ 12.69 (brs, D_2_O exch., 1H, N*H*), 10.79 (brs, D_2_O exch., 1H, N*H*CO), 8.78 (dd, *J* = 7.9, 1.4 Hz, 1H, H-7), 8.23 (d, *J* = 8.6 Hz, 2H, Η-2΄, Η-6’), 7.81 (dd, *J* = 8.0, 1.4 Hz, 1H, H-5), 7.59 (d, *J* = 8.5 Hz, 2H, H-3’, H-5’), 7.48 (t, *J* = 8.0 Hz, 1H, H-6), 6.41 (d, *J* = 5.2 Hz, 1H, CHO*H*), 4.94 (t, *J* = 5.8 Hz, 1H, CH_2_O*H*), 4.17 (dd, *J* = 5.2, 3.4 Hz, 1H, C*H*), 3.72 (q, *J* = 6.1, 5.0 Hz, 2H, C*H*_2_), 1.35 (s, 9H (CH_3_)_3_); ^13^C NMR (101 MHz, DMSO-*d*_6_) δ 171.7 (NH*C*O), 162.5 (*C*O), 155.2 (C-4’), 152.0 (C-2), 138.5 (C-8a), 134.0 (C-4a), 129.9 (C-1’), 128.1 (C-2’, C-6’), 127.1 (C-6), 126.0 (C-3’, C-5’), 121.3 (C-7), 119.8 (C-5), 74.1 (*C*H), 64.1 (*C*H_2_), 35.2 (*C*(CH_3_)_3_), 31.4 ((*C*H_3_)_3_); HRMS (ESI^+^) m/z 382.1765 (calcd for C_21_H_23_N_3_O_4_^+^, 382.1761).

#### (R)-N-(2-(4-(tert-Butyl)phenyl)-4-oxo-3,4-dihydroquinazolin-8-yl)-2,3-dihydroxypropanamide (52)

This compound was synthesized by an analogous procedure as described for the preparation of compound **23**. Yield: 57.8%. Mp: 287 °C. ^1^H NMR (400 MHz, DMSO-*d*_6_) δ 12.69 (brs, D_2_O exch., 1H, N*H*), 10.79 (brs, D_2_O exch., 1H, N*H*CO), 8.78 (dd, *J* = 7.9, 1.4 Hz, 1H, H-7), 8.23 (d, *J* = 8.6 Hz, 2H, Η-2΄, Η-6’), 7.81 (dd, *J* = 8.0, 1.4 Hz, 1H, H-5), 7.59 (d, *J* = 8.5 Hz, 2H, H-3’, H-5’), 7.48 (t, *J* = 8.0 Hz, 1H, H-6), 6.41 (d, *J* = 5.2 Hz, 1H, CHO*H*), 4.94 (t, *J* = 5.8 Hz, 1H, CH_2_O*H*), 4.17 (dd, *J* = 5.2, 3.4 Hz, 1H, C*H*), 3.72 (q, *J* = 6.1, 5.0 Hz, 2H, C*H*_2_), 1.35 (s, 9H (CH_3_)_3_); ^13^C NMR (101 MHz, DMSO-*d*_6_) δ 171.7 (NH*C*O), 162.5 (*C*O), 155.2 (C-4’), 152.0 (C-2), 138.5 (C-8a), 134.0 (C-4a), 129.9 (C-1’), 128.1 (C-2’, C-6’), 127.1 (C-6), 126.0 (C-3’, C-5’), 121.3 (C-7), 119.8 (C-5), 74.1 (*C*H), 64.1 (*C*H_2_), 35.2 (*C*(CH_3_)_3_), 31.4 ((*C*H_3_)_3_); HRMS (ESI^+^) m/z 382.1765 (calcd for C_21_H_23_N_3_O_4_^+^, 382.1761).

#### 2-(4-(tert-Butyl)benzamido)-4-nitrobenzoic acid (55)

This compound was synthesized by an analogous procedure as described for the preparation of compound **30**. Yield: 32.7%. Mp: 288 °C. ^1^H NMR (400 MHz, DMSO-*d*_6_) δ 12.23 (brs, D_2_O exch., 1H, COO*H*), 9.55 (d, *J* = 2.4 Hz, 1H, H-3), 8.27 (d, *J* = 8.7 Hz, 1H, H-6), 7.99 (dd, *J* = 8.7, 2.4 Hz, 1H, H-5), 7.90 (d, *J* = 8.6 Hz, 2H, H-2’, H-6’), 7.63 (d, *J* = 8.6 Hz, 2H, H-3’, H-5’), 1.33 (s, 9H, (C*H*_3_)_3_); ^13^C NMR (101 MHz, DMSO-*d*_6_) δ 169.1 (*C*OOH), 165.5 (NH*C*O), 156.2 (C-4’), 150.7 (C-4), 142.1 (C-2), 133.2 (C-6), 131.5 (C-1’), 127.5 (C-2’, C-6’), 126.4 (C-3’, C-5’), 122.3 (C-1), 117.5 (C-5), 114.7 (C-3), 35.3 (*C*(CH_3_)_3_), 31.3 ((*C*H_3_)_3_).

#### 4-Nitro-2-(4-(trifluoromethyl)benzamido)benzoic acid (56)

This compound was synthesized by an analogous procedure as described for the preparation of compound **30**. Yield: 80.9%. Mp: 275 °C. ^1^H NMR (600 MHz, DMSO-*d*_6_) δ 11.98 (brs, D_2_O exch., 1H, COO*H*), 9.54 (s, 1H, H-3), 8.26 (d, *J* = 8.6 Hz, 1H, H-6), 8.23 (d, *J* = 8.1 Hz, 2H, H-2’, H-6’), 7.98 (d, *J* = 8.0 Hz, 2H, H-3’, H-5’), 7.90 (d, *J* = 8.3 Hz, 1H, H-5); ^13^C NMR (151 MHz, DMSO-*d*_6_) δ 167.7, 164.1, 148.8, 147.3, 141.6, 138.9, 132.8, 128.6, 126.4, 125.3, 123.46, 117.1, 113.7.

#### 2-Benzamido-4-nitrobenzoic acid (57)

This compound was synthesized by an analogous procedure as described for the preparation of compound **30**. Yield: 54%. Mp: 268 °C. ^1^H NMR (600 MHz, Acetone-*d*_6_) δ 12.32 (brs, D_2_O exch., 1H, COO*H*), 9.86 (d, J = 2.0 Hz, 1H, H-3), 8.45 (d, *J* = 8.7 Hz, 1H, H-6), 8.09 – 8.06 (m, 2H, H-2’, H-6’), 8.04 (dd, *J* = 8.7, 2.3 Hz, 1H, H-5), 7.69 (m, 1H, H-4’), 7.64 (m, 2H, H-3’, H-5’); ^13^C NMR (151 MHz, Acetone-*d*_6_) δ 168.7 (COOH), 165.4 (NHCO), 151.3 (C-4), 143.0 (C-2), 134.3 (C-1’), 132.9 (C-6), 132.5 (C-4’), 129.0 (C-3’, C-5’), 127.3 (C-2’, C-6’), 120.3 (C-1), 116.6 (C-5), 114.4 (C-3).

#### 2-(4-(tert-Butyl)phenyl)-7-nitroquinazolin-4(3H)-one (64)

This compound was synthesized by an analogous procedure as described for the preparation of compound **40**. Yield: 90.1%. Mp: 312 °C. ^1^H NMR (400 MHz, DMSO-*d_6_*) δ 12.84 (brs, D_2_O exch., 1H, N*H*), 8.42 (d, *J* = 2.3 Hz, 1H, H-8), 8.36 (d, *J* = 8.7 Hz, 1H, H-5), 8.21 (dd, *J* = 8.7, 2.2 Hz, 1H, H-6), 8.17 (d, *J* = 9.0 Hz, 2H, H-2’, H-6’), 7.60 (d, *J* = 8.4 Hz, 2H, H-3’, H-5’), 1.34 (s, 9H, (C*H*_3_)_3_); ^13^C NMR (101 MHz, DMSO-*d*_6_) δ 161.9 (CO), 155.5 (C-4’), 154.8 (C-2), 151.8 (C-7), 149.8 (C-8a), 129.8 (C-1’), 128.7 (C-5), 128.4 (C-2’, C-6’), 126.0 (C-3’, C-5’), 125.7 (C-4a), 122.8 (C-8), 120.3 (C-6), 35.2 (*C*(CH_3_)_3_), 31.4 ((*C*H_3_)_3_); HRMS (ESI^+^) m/z 324.1348 (calcd for C_18_H_17_N_3_O_3_^+^, 324.1343).

#### 7-Nitro-2-(4-(trifluoromethyl)phenyl)quinazolin-4(3H)-one (65)

This compound was synthesized by an analogous procedure as described for the preparation of compound **40**. Yield: 90.9%. Mp: 335 °C (dec.). ^1^H NMR (400 MHz, DMSO-*d*_6_) δ 13.10 (brs, D_2_O exch., 1H, N*H*), 8.43 (d, *J* = 2.2 Hz, 1H, H-8), 8.40 – 8.34 (m, 3H, H-5, H-2’, H-6’), 8.24 (dd, *J* = 8.7, 2.3 Hz, 1H, H-6), 7.94 (d, *J* = 8.3 Hz, 2H, H-3’, H-5’); ^13^C NMR (101 MHz, DMSO-*d*_6_) δ 161.7, 153.8, 151.8, 149.3, 136.5, 132.5, 132.2, 131.9, 131.6, 129.5, 128.7, 126.0, 125.7, 123.0, 121.0; HRMS (ESI^+^) m/z 336.0595 (calcd for C_15_H_8_F_3_N_3_O_3_^+^, 336.0591).

#### 7-Nitro-2-phenylquinazolin-4(3H)-one (66)

This compound was synthesized by an analogous procedure as described for the preparation of compound **40**. Yield: 93.9%. Mp: 338 °C. ^1^H NMR (600 MHz, DMSO-d_6_) δ 12.88 (brs, D_2_O exch., 1H), 8.44 (d, J = 2.2 Hz, 1H, H-8), 8.37 (d, J = 8.6 Hz, 1H, H-6), 8.25 – 8.17 (m, 3H, H-5, H-2’, H-6’), 7.64 (t, J = 7.3 Hz, 1H, H-4’), 7.58 (t, J = 7.5 Hz, 2H, H-3’, H-5’); ^13^C NMR (151 MHz, DMSO-*d*_6_) δ 161.9 (*C*O), 155.1 (C-2), 151.9 (C-7), 149.6 (C-8a), 132.6 (C-4’), 132.5 (C-1’), 129.2 (C-3’, C-5’), 128.7 (C-5), 128.5 (C-2’, C-6’), 125.8 (C-4a), 122.8 (C-8), 120.5 (C-6); HRMS (ESI^+^) m/z 268.0722 (calcd for C_14_H_9_N_3_O_3_^+^, 268.0717).

#### N-(2-(4-(tert-Butyl)phenyl)-4-oxo-3,4-dihydroquinazolin-7-yl)acrylamide (70)

This compound was synthesized by an analogous procedure as described for the preparation of compound **17**. Yield: 90.4%. Mp: 268 °C. ^1^H NMR (400 MHz, DMSO-*d*_6_) δ 12.35 (brs, D_2_O exch., 1H, N*H*), 10.57 (brs, D_2_O exch., 1H, N*H*CO), 8.21 (d, *J* = 2.0 Hz, 1H, H-8), 8.13 (d, *J* = 8.2 Hz, 2H, H-2’, H-6’), 8.10 (d, *J* = 8.6 Hz, 1H, H-5), 7.65 (dd, *J* = 8.7, 2.0 Hz, 1H, H-6), 7.57 (d, *J* = 8.5 Hz, 2H, H-3’, H-5’), 6.51 (dd, *J* = 17.0, 10.1 Hz, 1H, C*H*=CHH), 6.35 (dd, *J* = 17.0, 1.9 Hz, 1H, CH=C*H*H), 5.86 (dd, *J* = 10.0, 2.0 Hz, 1H, CH=CH*H*), 1.34 (s, 9H, (CH_3_)_3_); ^13^C NMR (101 MHz, DMSO-*d*_6_) δ 164.3 (NHCO), 162.2 (CO), 154.1 (C-4’), 153.1 (C-2), 150.4 (C-8a), 144.8 (C-4a), 132.0 (*C*H=CHH), 130.4 (C-1’), 128.5 (CH=*C*HH), 128.0 (C-2’, C-6’), 127.3 (C-5), 125.9 (C-3’, C-5’), 118.7 (C-6), 116.8 (C-7), 116.1 (C-8), 35.2 (*C*(CH_3_)_3_), 31.4 ((*C*H_3_)_3_); HRMS (ESI^+^) m/z 348.1708 (calcd for C_21_H_21_N_3_O_2_^+^, 348.1707).

#### N-(4-Oxo-2-(4-(trifluoromethyl)phenyl)-3,4-dihydroquinazolin-7-yl)acrylamide (71)

This compound was synthesized by an analogous procedure as described for the preparation of compound **17**. Yield: 66.3%. Mp: 315 °C. ^1^H NMR (400 MHz, DMSO-*d*_6_) δ 12.63 (brs, D_2_O exch., 1H, N*H*), 10.60 (brs, D_2_O exch., 1H, N*H*CO), 8.37 (d, *J* = 8.2 Hz, 2H, H-2’, H-6’), 8.26 (d, *J* = 2.0 Hz, 1H, H-8), 8.12 (d, *J* = 8.7 Hz, 1H, H-5), 7.92 (d, *J* = 8.4 Hz, 2H, H-3’, H-5’), 7.69 (dd, *J* = 8.6, 2.1 Hz, 1H, H-6), 6.50 (dd, *J* = 17.0, 10.1 Hz, 1H, C*H*=CHH), 6.35 (dd, *J* = 17.0, 2.0 Hz, 1H, CH=C*H*H), 5.86 (dd, *J* = 10.0, 2.0 Hz, 1H, CH=CH*H*); ^13^C NMR (101 MHz, DMSO-*d*_6_) δ 164.3, 162.0, 152.1, 150.0, 145.0, 137.1, 132.0, 131.8, 131.4, 129.2, 128.6, 127.4, 126.0, 123.1, 119.3, 117.1, 116.3; HRMS (ESI^+^) m/z 360.0958 (calcd for C_18_H_12_F_3_N_3_O_2_^+^, 360.0954).

#### N-(4-Oxo-2-phenyl-3,4-dihydroquinazolin-7-yl)acrylamide (72)

This compound was synthesized by an analogous procedure as described for the preparation of compound **17**. Yield: 48.6%. Mp: 313 °C (dec.). ^1^H NMR (400 MHz, DMSO-*d*_6_) δ 12.43 (brs, D_2_O exch., 1H, N*H*), 10.57 (brs, D_2_O exch., 1H, N*H*CO), 8.22 (d, *J* = 2.0 Hz, 1H, H-8), 8.18 (dt, *J* = 6.7, 1.6 Hz, 2H, H-2’, H-6’), 8.10 (d, *J* = 8.7 Hz, 1H, H-5), 7.66 (dd, *J* = 8.6, 2.1 Hz, 1H, H-6), 7.61 – 7.52 (m, 3H, H-3’, H-4’, H-5’), 6.50 (dd, *J* = 17.0, 10.0 Hz, 1H, C*H*=CHH), 6.35 (dd, *J* = 17.0, 2.0 Hz, 1H, *C*H=C*H*H), 5.85 (dd, *J* = 10.0, 2.0 Hz, 1H, *C*H=CH*H*); ^13^C NMR (101 MHz, DMSO-*d*_6_) δ 164.3 (*C*O), 162.1 (NH*C*O), 153.3 (C-2), 150.3 (C-8a), 144.9 (C-4a), 133.2 (C-4’), 131.99 (*C*H=CHH), 131.9 (C-1’), 129.1 (C-3’, C-5’), 128.5 (CH=*C*HH), 128.2 (C-2’, C-6’), 127.3 (C-5), 118.8 (C-6), 116.9 (C-7), 116.2 (C-8); HRMS (ESI^+^) m/z 292.1085 (calcd for C_17_H_13_N_3_O_2_^+^, 292.1081).

#### N-(2-(4-(tert-Butyl)phenyl)-4-oxo-3,4-dihydroquinazolin-7-yl)-2,3-dihydroxypropanamide (73)

This compound was synthesized by an analogous procedure as described for the preparation of compound **20**. Yield: 81.8%. Mp: 216 °C. ^1^H NMR (600 MHz, DMSO-*d*_6_) δ 12.35 (brs, D_2_O exch., 1H), 10.08 (brs, D_2_O exch., 1H), 8.24 (d, *J* = 2.0 Hz, 1H, H-8), 8.13 (d, *J* = 8.5 Hz, 2H, H-2’, H-6’), 8.06 (d, *J* = 8.6 Hz, 1H, H-5), 7.77 (dd, *J* = 8.7, 2.1 Hz, 1H, H-6), 7.57 (d, *J* = 8.5 Hz, 2H, H-3’, H-5’), 5.85 (d, *J* = 5.6 Hz, 1H, CHO*H*), 4.87 (t, *J* = 5.7 Hz, 1H, CH_2_O*H*), 4.13 (td, *J* = 5.6, 4.1 Hz, 1H, C*H*OH), 3.73 – 3.62 (m, 2H, C*H*_2_OH), 1.34 (s, 9H, (C*H*_3_)_3_); ^13^C NMR (101 MHz, DMSO-*d*_6_) δ 172.7 (NH*C*O), 162.2 (*C*O), 154.8 (C-4’), 153.0 (C-2), 150.3 (C-8a), 144.5 (C-4a), 130.5 (C-1’), 128.0 (C-2’, C-6’), 127.0 (C-5), 125.9 (C-3’, C-5’), 119.0 (C-6), 116.7 (C-7), 116.3 (C-8), 74.08 (*C*HOH), 64.2 (*C*H_2_OH), 35.2 (*C*(CH_3_)_3_), 31.4 ((*C*H_3_)_3_); HRMS (ESI^+^) m/z 382.1763 (calcd for C_21_H_23_N_3_O_4_^+^, 382.1761).

#### 2,3-Dihydroxy-N-(4-oxo-2-(4-(trifluoromethyl)phenyl)-3,4-dihydroquinazolin-7-yl)propanamide (74)

This compound was synthesized by an analogous procedure as described for the preparation of compound **20**. Yield: 48.8%. Mp: 321 °C. ^1^H NMR (400 MHz, DMSO-*d*_6_) δ 12.61 (brs, D_2_O exch., 1H, N*H*), 10.11 (brs, D_2_O exch., 1H, N*H*CO), 8.35 (d, *J* = 8.2 Hz, 2H, H-2’, H-6’), 8.28 (d, *J* = 2.0 Hz, 1H, H-8), 8.09 (d, *J* = 8.7 Hz, 1H, H-5), 7.92 (d, *J* = 8.3 Hz, 2H, H-3’, H-5’), 7.81 (dd, *J* = 8.7, 2.1 Hz, 1H, H-6), 5.88 (d, *J* = 5.6 Hz, 1H, CHO*H*), 4.90 (t, *J* = 5.7 Hz, 1H, CH_2_O*H*), 4.19 – 4.11 (m, 1H, C*H*OH), 3.73 – 3.62 (m, 2H, C*H*_2_OH); ^13^C NMR (101 MHz, DMSO-*d*_6_) δ 172.8, 162.1, 152.1, 149.9, 144.5, 137.1, 131.4, 129.2, 127.1, 126.0, 123.1, 119.6, 116.9, 116.5, 74.0, 64.2; HRMS (ESI^+^) m/z 394.1008 (calcd for C_18_H_14_F_3_N_3_O_4_^+^, 394.1009).

#### 2,3-Dihydroxy-N-(4-oxo-2-phenyl-3,4-dihydroquinazolin-7-yl)propanamide (75)

This compound was synthesized by an analogous procedure as described for the preparation of compound **20**. Yield: 43.6%. Mp: 275 °C. ^1^H NMR (400 MHz, DMSO-*d*_6_) δ 12.40 (brs, D_2_O exch., 1H, N*H*), 10.08 (brs, D_2_O exch., 1H, N*H*CO), 8.25 (d, *J* = 2.0 Hz, 1H, H-8), 8.17 (m, 2H, H-2’, H-6’), 8.07 (d, *J* = 8.7 Hz, 1H, H-5), 7.78 (dd, *J* = 8.7, 2.1 Hz, 1H, H-6), 7.62 – 7.51 (m, 3H, C-3’, C-4’, C-5’), 5.86 (d, *J* = 5.7 Hz, 1H, CHO*H*), 4.88 (t, *J* = 5.7 Hz, 1H, CH_2_O*H*), 4.13 (td, *J* = 5.4, 4.1 Hz, 1H, C*H*OH), 3.73 – 3.60 (m, 2H, C*H*_2_OH); ^13^C NMR (101 MHz, DMSO-*d*_6_) δ 172.8 (NH*C*O), 162.2 (*C*O), 153.2 (C-2), 150.2 (C-8a), 144.5 (C-4a), 133.2 (C-1’), 131.9 (C-4’), 129.1 (C-3’, C-5’), 128.2 (C-2’, C-6’), 127.0 (C-5), 119.2 (C-6), 116.8 (C-7), 116.3 (C-8), 74.1 (*C*HOH), 64.2 (*C*H_2_OH); HRMS (ESI^+^) m/z 326.1144 (calcd for C_17_H_15_N_3_O_4_^+^, 326.1135).

### Protein production

The expression and purification of human TNKS1 (residues 1030–1317) and TNKS2 (residues 873– 1161 construct for activity assay, residues 952–1161 for crystallization) recombinant proteins, together with other PARP enzymes constructs, were carried out as described in former studies [14,95,96]. Information on constructs employed in the biochemical activity assay is available in **Table S1**. Affinity chromatography (nickel-based IMAC, MBP-based affinity chromatography) and size exclusion chromatography were commonly used as purification methods. In addition, DNA-binding PARPs were further purified through heparin-based affinity chromatography prior to size exclusion chromatography.

### Biochemical activity assay

The enzymatic assay is based on the quantification of residual NAD^+^ through its conversion into a fluorescent compound, as extensively described by Narwal et al. [95]. The enzymatic reactions were carried out at room temperature. TNKS1/2 was incubated for 20 hours in shaking conditions (300 rpm using PST-100HL plate shaker-thermostat from Biosan) with half-logarithmic dilutions of each compound and 5 µM NAD^+^. Each reaction was executed in assay buffer 50 mM Bis-Tris propane (BTP), pH 7.0, 0.5 mM Tris(2-carboxyethyl)phosphine (TCEP), 0.01% Triton X-100. After the incubation time, treatment with 20% acetophenone in ethanol, 2 M KOH and formic acid allowed the conversion of unreacted NAD^+^ into a stable fluorescent compound. The fluorescent intensities were measured with excitation wavelength of 372 nm and emission wavelength of 444 nm (Tecan Spark multimode microplate reader). The conditions for the activity assay with other PARP enzymes were set as described in **Table S1** and key IC_50_ curves are shown in **Fig. S2**.

### Cell assays

The luciferase-based WNT/β-catenin signaling pathway reporter assay was performed in human HEK293 cells, as previously described [69]. HEK293 cells, stably transfected with plasmids expressing ST-luciferase and *Renilla* luciferase, were seeded in 96-well plates coated with poly-l-lysine. The day after, the cells were incubated for 24 hours with various compound concentrations in 50% WNT3a-conditioned medium. After 24 hours of compound exposure, the cells were lysed and the firefly luciferase and Renilla activities were measured on a GloMax Luminometer (Promega) using a Dual-Glo Luciferase Assay System (Promega).

### Crystallization, data collection, processing and refinement

TNKS2 ART domain (residues 952–1161, 7.7 mg/mL) was incubated with chymotrypsin (1:100) for 2 hours at room temperature. Protein crystals were grown at 4°C by mixing the treated protein in a 1:1 ratio with 100 mM Tris, pH 8.5, 22% PEG 3350, 200 mM Li2SO4. The crystallization was carried out with the sitting-drop vapor diffusion method in SwissCI 96-Well 3-Drop plate. 3D crystals were fully grown after two days. Selected compounds were added to the crystallization droplets with a final concentration of 300 µM. After two days of incubation at 4°C, the crystals were cryo-protected by soaking into precipitant solution added with 250 mM NaCl and 20% glycerol. PARP15 ART domain (residues 481-678, 19 mg/mL) was co-crystallized with compound **49** by incubating the protein in a 1:2 ratio with 200 mM NH_4_Cl and 20 % PEG3350. The crystals grew at room temperature by employing the the sitting-drop vapor diffusion method in SwissCI 96-Well 3-Drop plate. After two days, 3D crystals were completely grown and were cryo-protected using 30% MPD and 200 mM NH_4_Cl. Afterwards, all the crystals were frozen in liquid nitrogen. Data were collected at MAX IV Laboratory (BioMAX beamline), ESRF (ID30A-1 and ID30A-3 beamlines) and Diamond (beamline I24). Diffraction data were processed using the XDS package [97]. Molecular replacement was performed with PHASER by using TNKS2 structure (PDB: 7OM1) and PARP15 (PDB: 7Z2O) as model [98]. Protein model building was carried out with Coot, while REFMAC5 and Phenix were employed for refinement of the crystal structures [99–101]. Data collection and refinement statistics for crystal structures is provided in **Table S2**.

## Data access

The coordinates and structure factors of crystal structures are deposited to PDB with IDs 8S60, 8S4V, 8S4W, 8S4X, 9FEG. Raw diffraction data will be available at Etsin (https://doi.org/10.23729/f89486c4-d6b0-4595-a7eb-f68ae81ca9f2).

## Author contributions

I.K., L.L. conceptualization; C.B., D.K., A. G.-P., S.A.B., S.S.D, J.A., M.K., investigation; C.B., D.K. writing-original draft; C.B. D.K. J. W., L.L, I.K. writing-reviewing and editing; J.W., I.K., L.L. funding acquisition; J.W., L.L., I.K supervision.

## Acknowledgements

The use of the facilities and expertise of the Biocenter Oulu Structural Biology core facility, a member of Biocenter Finland, Instruct-ERIC Centre Finland and FINStruct, are gratefully acknowledged. We are grateful to local contacts at the MAX IV laboratory, ESRF and Diamond for helping in using beamlines BioMAX, ID30A-1, ID30A-3 and I24.

