## Supplementary information for "Substitutions at the C-8 position of quinazolin-4-ones improve the potency of nicotinamide site binding tankyrase inhibitors"

**Table S1.** Protein constructs and conditions employed for the activity assays.

| Enzyme | Residues | Vector | Activity assay conditions | Comments |
|---|---|---|---|---|
| PARP1 | 1 - 1014 | pNH-TrxT | 5 nM PARP1 and 500 nM NAD+<br>50 mM Tris pH 8.0, 10 µg/mL activated DNA, 5 mM MgCl2, 0.2 mg/ml BSA<br>30 minutes shaking at RT | |
| PARP2 | 1 - 583 | pNH-TrxT | 30 nM PARP2 and 500 nM NAD+<br>50 mM Tris pH 8.0, 10 µg/mL activated DNA, 5 mM MgCl2, 0.1 mg/ml BSA<br>30 min shaking at RT | |
| PARP3 | 1 - 533 | pNH-TrxT | 20 nM PARP3 and 500 nM NAD+<br>50 mM PIPES pH 7.0, 20 µg/mL activated DNA, 5 mM MgCl2, 0.2 mg/ml BSA<br>4 hours shaking at RT | |
| PARP4 | 250 - 565 | pNIC28-Bsa4 | 400 nM PARP4 and 5 µM NAD+<br>50mM Na-P pH 7.5, 1 mg/ml BSA, 0.5 mM TCEP<br>2.5 hours shaking at RT | |
| TNKS1 | 1030 - 1317 | pNIC-MBP (pNIC28-Bsa4 based) | 100 nM TNKS1 and 20 µM NAD+<br>50mM BisTris propane pH 7.0, 0.5 mM TCEP, 0.01% Triton X-100<br>20 hours shaking at RT | Final construct provided with His-tag and MBP-tag |
| TNKS2 | 873–1161 | pNIC-MBP (pNIC28-Bsa4 based) | 20 nM TNKS2 and 5 µM NAD+<br>50mM BisTris propane pH 7.0, 0.5 mM TCEP, 0.01% Triton X-100<br>20 hours shaking at RT | Final construct provided with His-tag and MBP-tag |
| PARP10 | 809 - 1017 | pNIC-MBP (pNIC28-Bsa4 based) | 60 nM PARP10, 2.5 µM SRPK2, C8 scaffold (1:8 molar ratio with the enzyme) and 500 nM NAD+<br>50 mM Na-P pH 7.0<br>18 hours shaking at RT | Final construct provided with His-tag, MBP-tag and dockerin domain |
| PARP14 | 1535 - 1801 | pNIC-MBP (pNIC28-Bsa4 based) | 110 nM PARP14, C8 scaffold (1:11 molar ratio with the enzyme) and 500 nM NAD+<br>50 mM Na-P pH 7.0<br>18 hours shaking at RT | Final construct provided with His-tag, MBP-tag and dockerin domain |

| | | | | |
|---|---|---|---|---|
| PARP15 | 481-678 | pNIC-MBP (pNIC28-Bsa4 based) | 130 nM PARP15, C8 scaffold (1:11 molar ratio with the enzyme) and 500 nM NAD+<br>50 mM Na-P pH 7.0<br>18 hours shaking at RT | Final construct provided with His-tag, MBP-tag and dockerin domain |

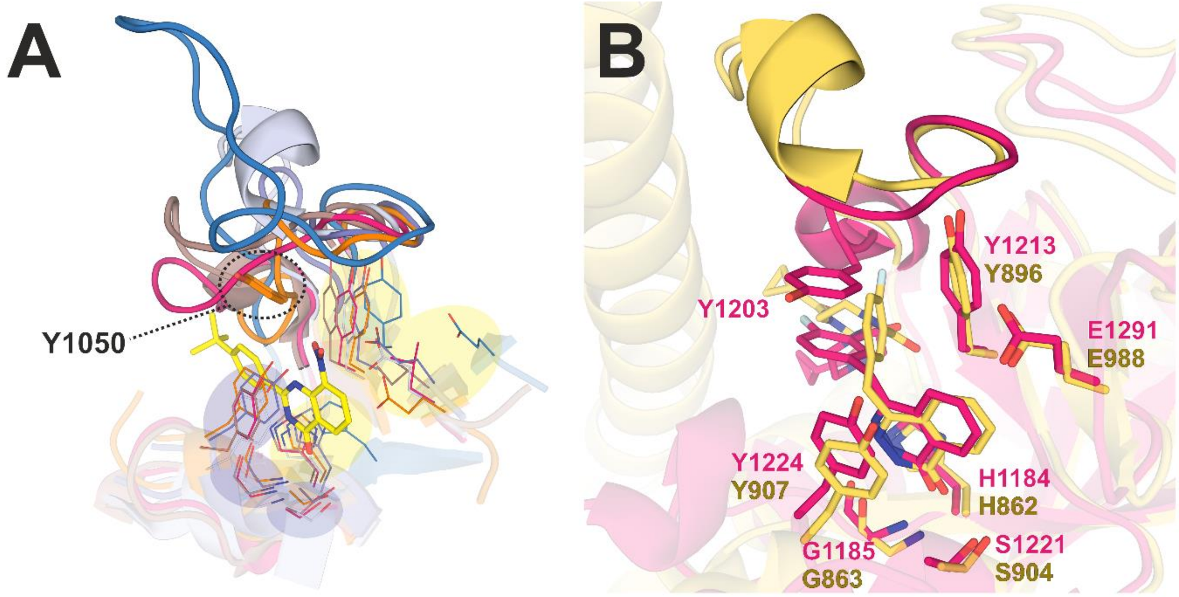

**Figure S1.**
A) Superimposition of crystal structure of TNKS2 (orange) in complex with compound 40 (yellow) with crystal structures of other PARP enzymes. PARP1 (PDB: 5LX6) is depicted in grey, PARP3 (PDB: 4L7N) in violet, PARP4 (PDB: 8SWY) in blue, PARP10 (PDB: 7ONT) in brown, PARP14 (PDB: 6RY4) in magenta. Binding compounds and part of the protein structures are hidden for representation purposes. The D-loops are in solid colors. Residues relevant for the catalysis (yellow background) and for the anchoring of the nicotinamide moiety (blue background) are shown as lines. Tyr1050 of TNKS2 is indicated since it is a non-conserved residue that establishes the π-hole interaction with the nitro moiety of compound 40. B) Superimposition of crystal structure of olaparib in complex with TNKS1 (magenta, PDB: 7KKO) and in complex with PARP1 (yellow, PDB: 7AAD).

It is well established that PARP1/2 D-loop is longer and more rigid in comparison to D-loop of tankyrases, possibly due to the presence of multiple proline residues [1]. This is evident by comparing crystal structures in which the same compound is bound to different members of the family, for example crystal structures of olaparib in complex with PARP1 [2] and TNKS1 [3]. According to this crystal structure data, the D-loop of TNKS1 and PARP1 impact differently the accomodation of olaparib in the NI pocket, since TNKS1 D-loops is shorter and its unique tyrosine residue of (Tyr1203) forms a π-π stacking interaction with the compound. D-loops of other PARPs display different length and plasticity, for example PARP3 is characterized by a shorter D-loop than PARP1 [4], while PARP4 crystal structures show a very extended loop [5], but they still lack a residue analogous to Tyr1203 of TNKS1 or Tyr1050 of TNKS2.

**Table S2.** Data collection and refinement statistics for crystal structures.

| **Protein, Inhibitor (PDB id.)** | **TNKS2, 40 (8S4X)** | **TNKS2, 41 (8S60)** | **PARP15, 49 (9FEG)** | **TNKS2, 51 (8S4W)** | **TNKS2, 52 (8S4V)** |
|---|---|---|---|---|---|
| **Data collection** | | | | | |
| Beamline | ID30A-3 (ESRF) | BioMAX (MAX IV) | I24 (Diamond) | ID30A-3 (ESRF) | ID30A-1 (ESRF) |
| Wavelength (Å) | 0.96770 | 0.920013 | 0.97950 | 0.96770 | 0.96546 |
| Space group | C 2 2 21 | C 2 2 21 | P 21 21 21 | P 41 21 2 | P 41 21 2 |
| Unit cell dimensions<br>*a*, *b*, *c* (Å)<br>α, β, γ (°) | 93.42, 95.50, 119.44<br>90, 90, 90 | 90.42, 98.16, 119.58<br>90, 90, 90 | 45.26, 68.46, 159.60<br>90, 90, 90 | 66.60, 66.60, 120.59<br>90, 90, 90 | 66.56, 66.56, 119.09<br>90, 90, 90 |
| Resolution range(Å) (Outer cell) | 50-2.50 (2.50-2.65) | 50-1.80 (1.80-1.91) | 50-1.75 (1.75-1.80) | 50-2.12 (2.12-2.25) | 50-1.93 (1.93-2.04) |
| Total n. of reflections | 96758 (15904) | 669135 (109322) | 332818 (53510) | 202891 (33204) | 210404 (34374) |
| N. of unique reflections | 18713 (2986) | 49407 (7875) | 50967 (8073) | 16027 (2463) | 20895 (3305) |
| Completeness (%) | 98.9 (99.1) | 100 (99.9) | 100 (100) | 99.5 (96.8) | 99.8 (99.3) |
| $\langle I/\sigma(I)\rangle$ | 11.74 (1.77) | 17.75 (2.11) | 11.51 (2.21) | 10.11 (1.50) | 13.17 (2.04) |
| CC1/2 (%) | 99.7 (77.0) | 99.9 (87.6) | 99.8 (78.7) | 99.7 (79.7) | 99.7 (83.9) |
| $R_{meas}$ | 0.107 (0.949) | 0.082 (0.103) | 0.122 (0.758) | 0.168 (1.67) | 0.100 (0.801) |
| R-factor observed (%) | 9.7 (85.6) | 7.9 (99.1) | 11.2 (69.9) | 16.1 (160.6) | 9.5 (76.1) |
| **Refinement** | | | | | |
| R-work/R-free | 0.197/0.246 | 0.182/0.211 | 0.182/0.213 | 0.189/0.244 | 0.197/235 |
| **N. of non-hydrogen atoms** | 3431 | 3601 | 3483 | 1802 | 1776 |
| Protein | 3340 | 3351 | 3229 | 1677 | 1675 |
| Ligands | 104 | 76 | 83 | 62 | 62 |
| Solvent | 21 | 174 | 214 | 86 | 62 |
| **RMSD** | | | | | |
| Bonds (Å) | 0.002 | 0.014 | 0.010 | 0.011 | 0.016 |
| Angles (°) | 0.44 | 1.70 | 0.84 | 1.08 | 1.06 |
| **Average *B* factors (Å$^2$)** | 56.68 | 38.93 | 23.00 | 48.37 | 43.61 |
| Protein | 56.59 | 38.76 | 22.73 | 48.57 | 43.71 |
| Ligands | 64.72 | 42.91 | 34.49 | 47.98 | 42.72 |
| Solvent | 43.97 | 40.49 | 24.95 | 44.75 | 41.24 |
| **Ramachandran plot** | | | | | |
| Favoured (%) | 97.79 | 98.29 | 98.19 | 97.07 | 97.55 |
| Allowed (%) | 2.21 | 1.71 | 1.81 | 2.93 | 2.45 |
| Outliers (%) | 0 | 0.56 | 0 | 0 | 0 |

*Values within parentheses refers to the highest resolution shell.

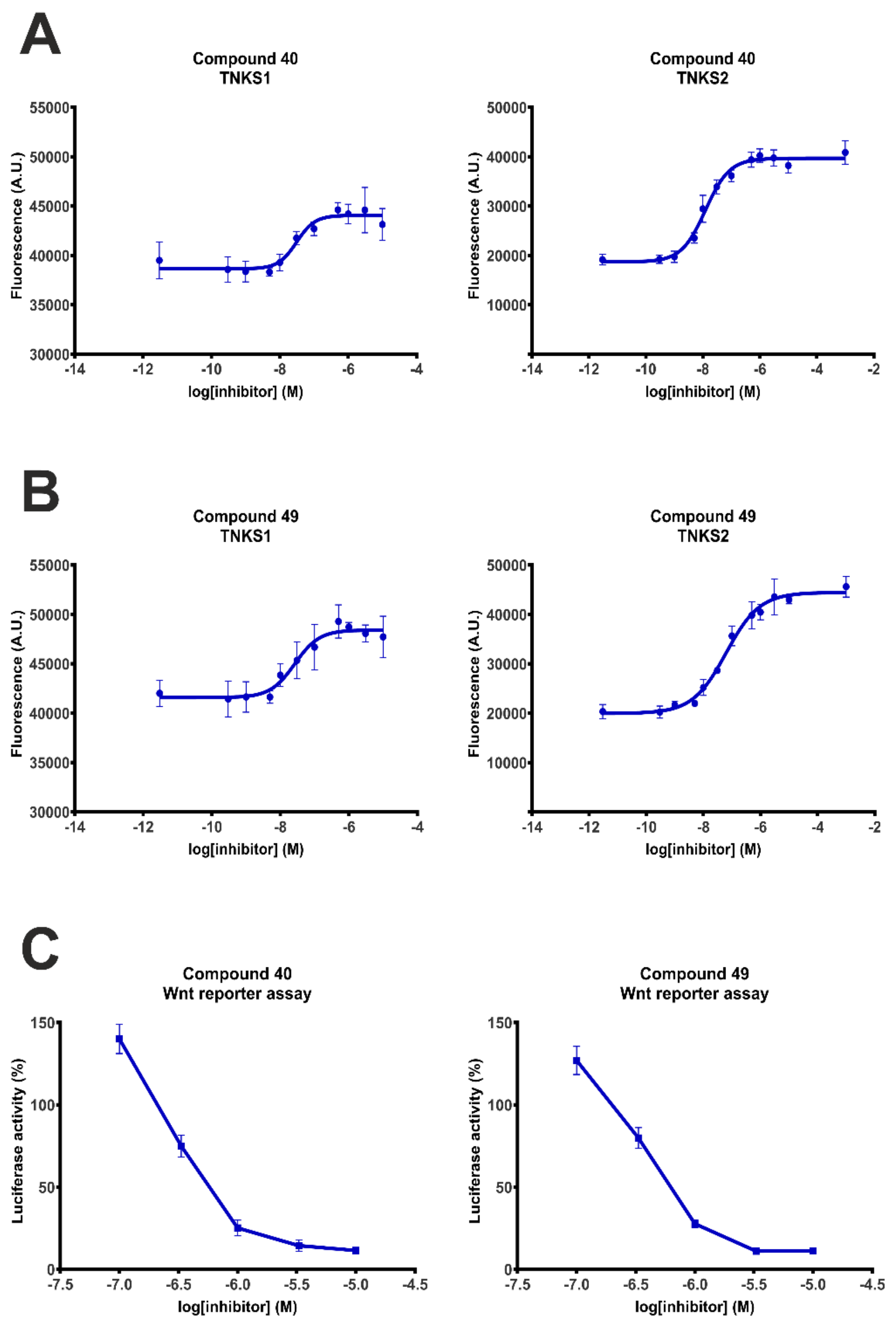

**Figure S2.**
Dose response curves for A) compound 40 and B) compound 49 resulted from the biochemical activity assay of TNKS1/2. Mean is shown and error bars indicate the SD (n=4). C) Reduction of luciferase activity (%) induced by compounds 40 and 49 in WNT/β-catenin signalling reporter assay in HEK293 cells. Mean ± SD are indicated (n=3).

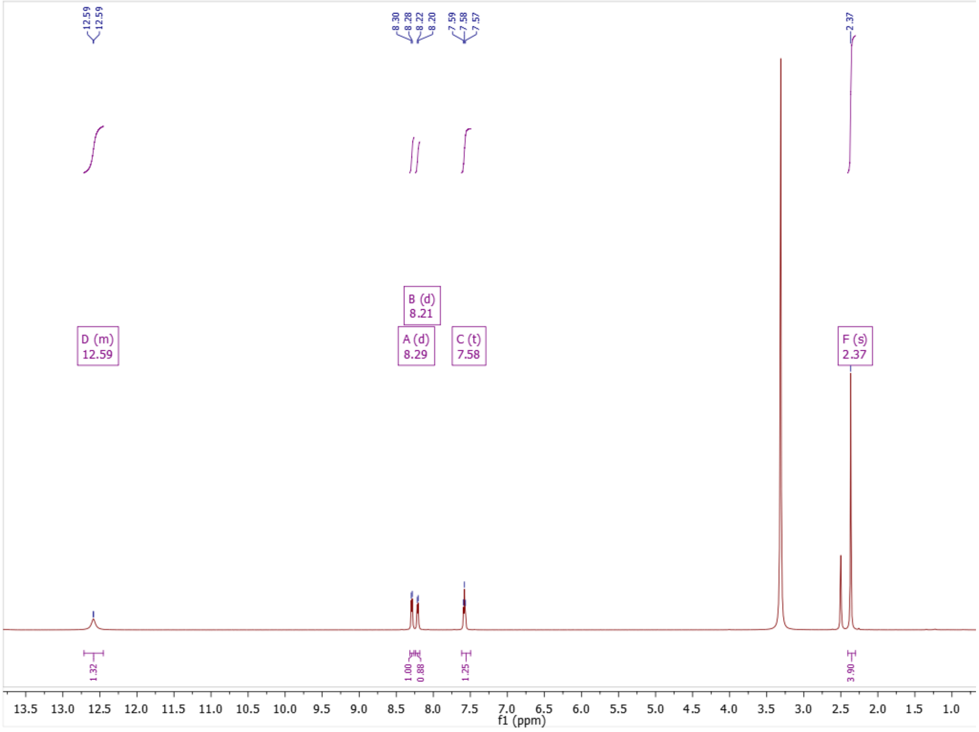

**Figure S3:** $^{1}$H NMR of compound **11**

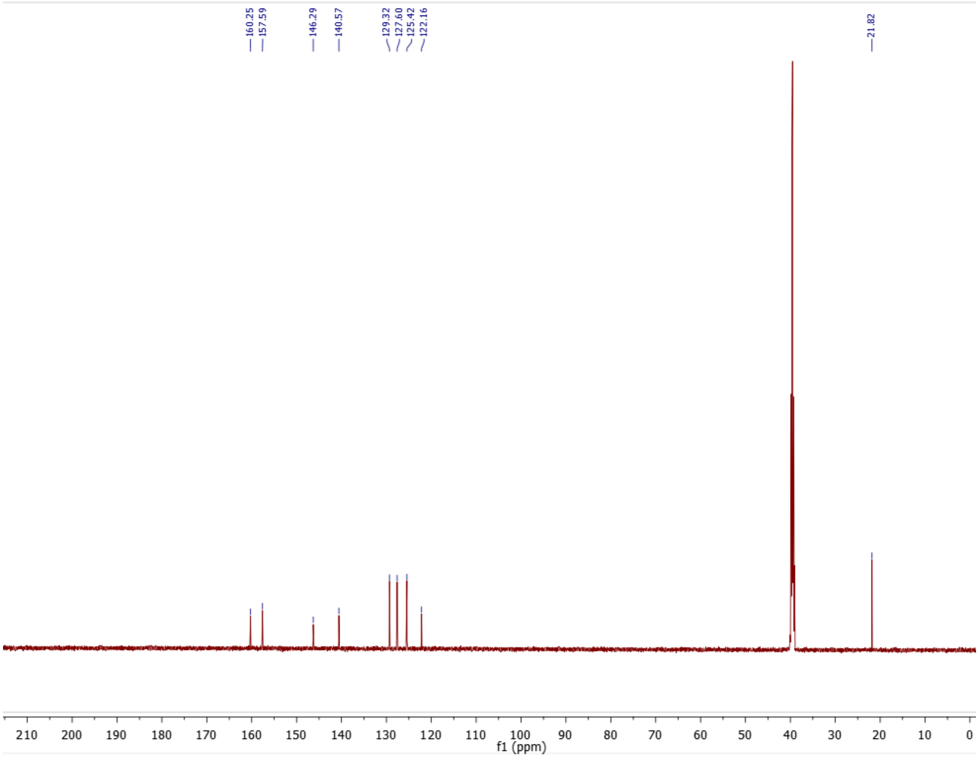

**Figure S4:** $^{13}$C NMR of compound **11**

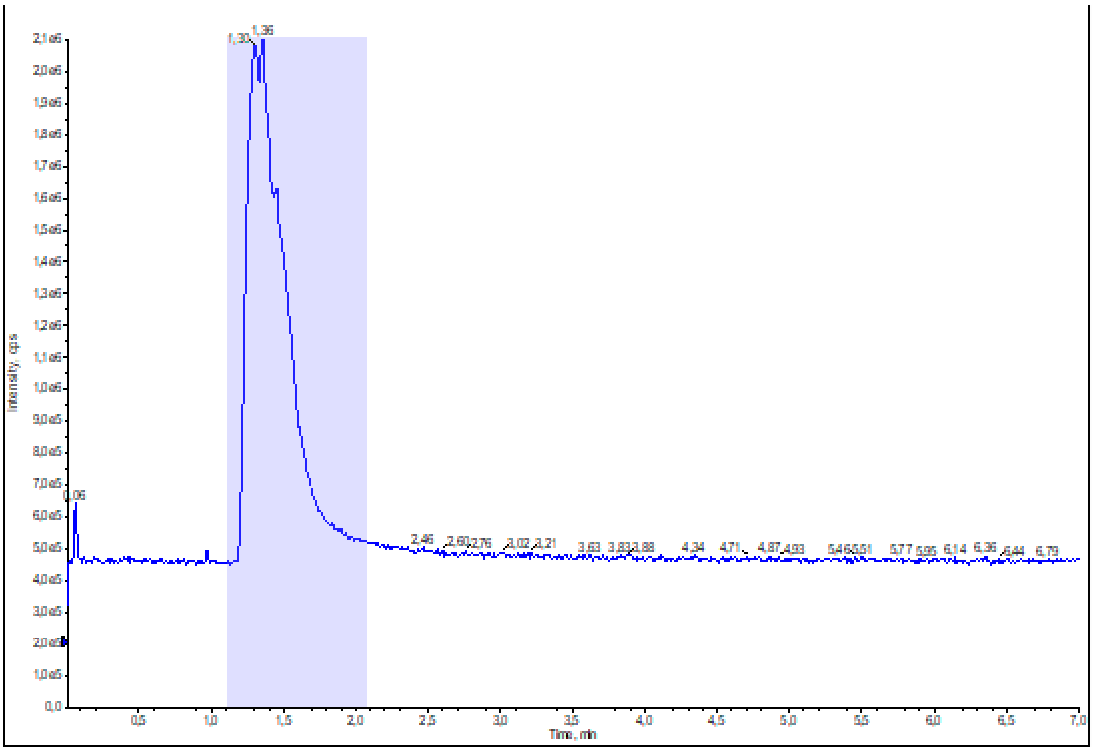

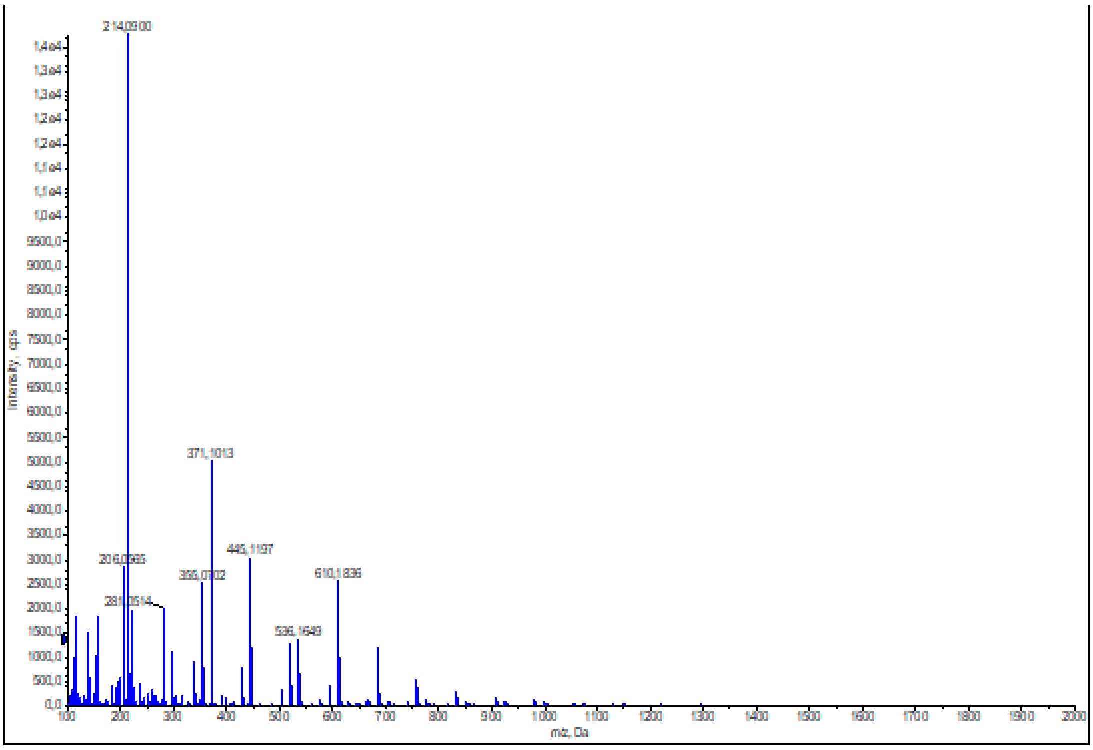

**Figure S5:** HRMS (ESI+) of compound **11**

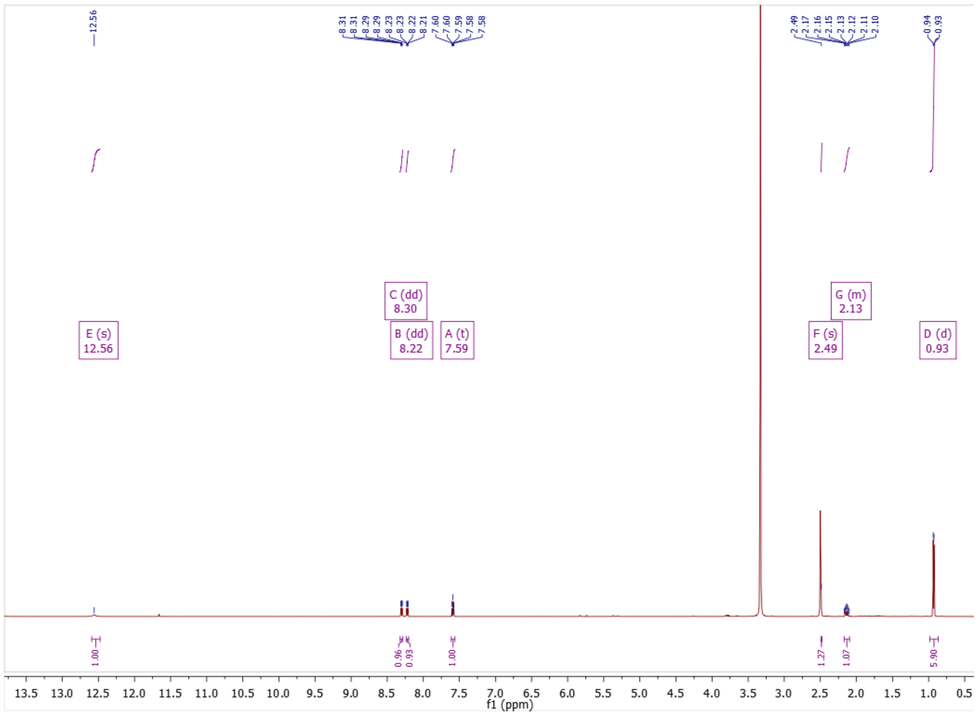

**Figure S6:** $^{1}$H NMR of compound **12**

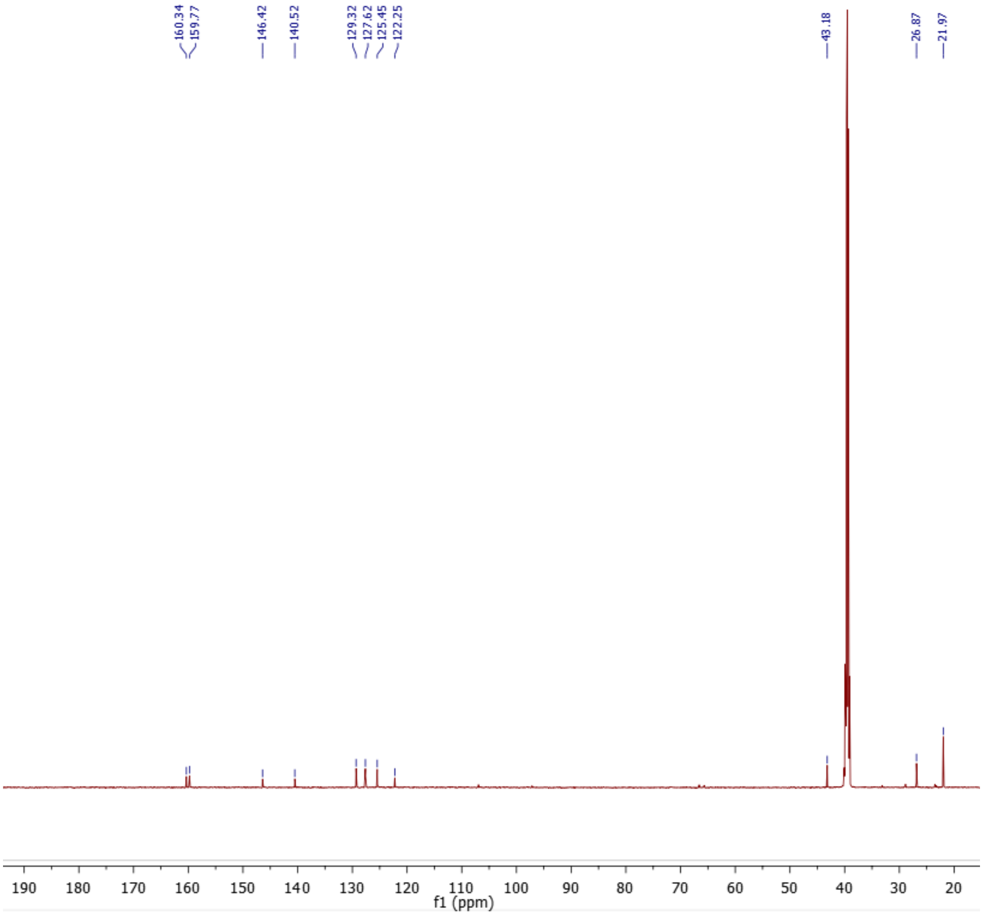

**Figure S7:** $^{13}$C NMR of compound **12**

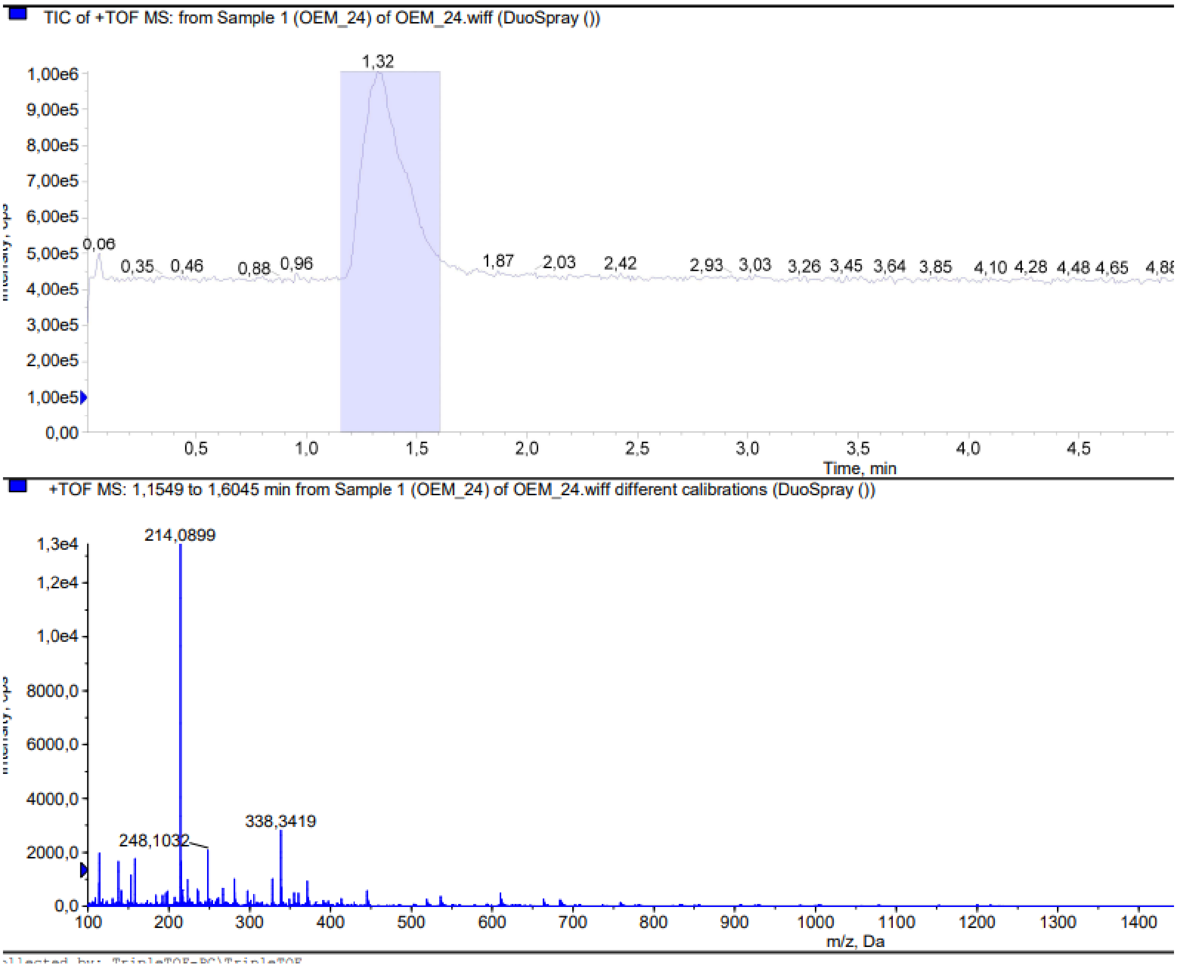

**Figure S8:** HRMS (ESI+) of compound **12**

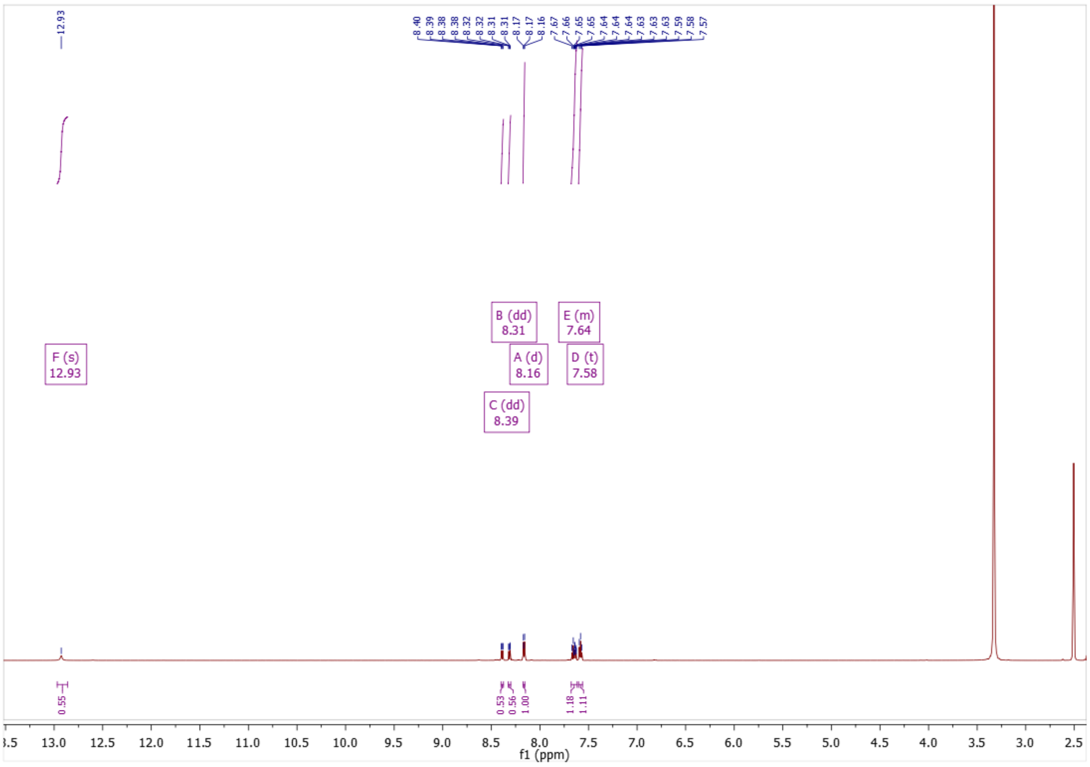

**Figure S9:** $^{1}$H NMR of compound **13**

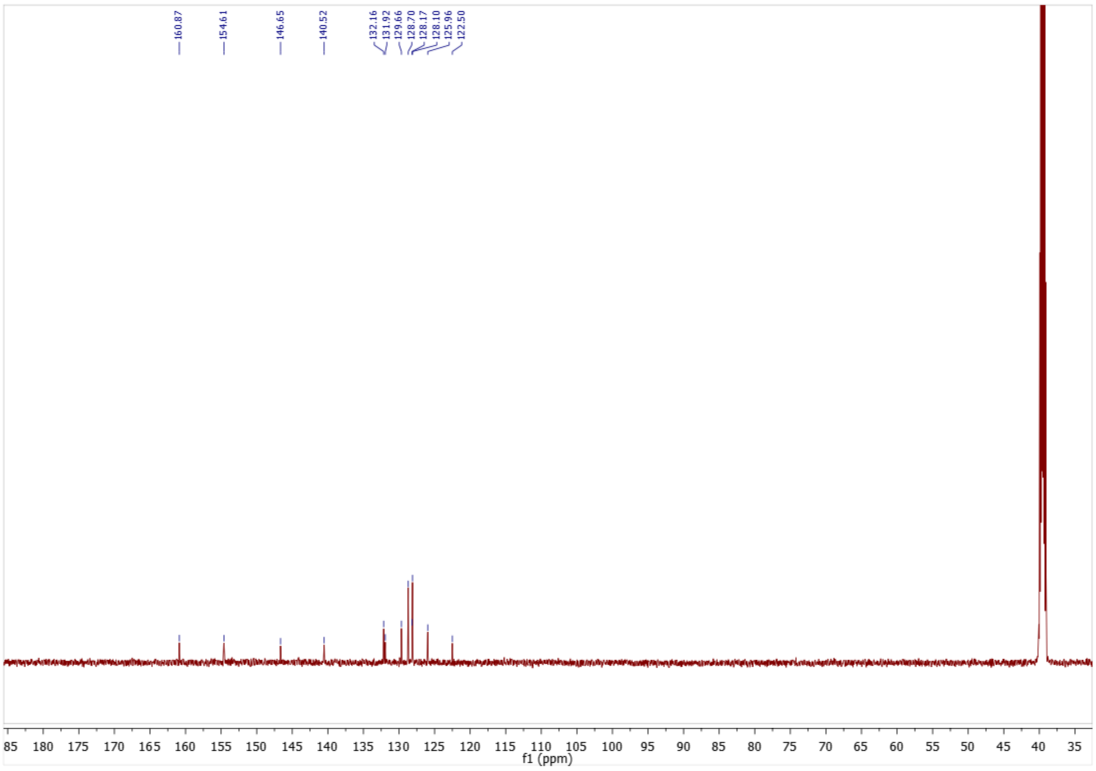

**Figure S10:** $^{13}$C NMR of compound **13**

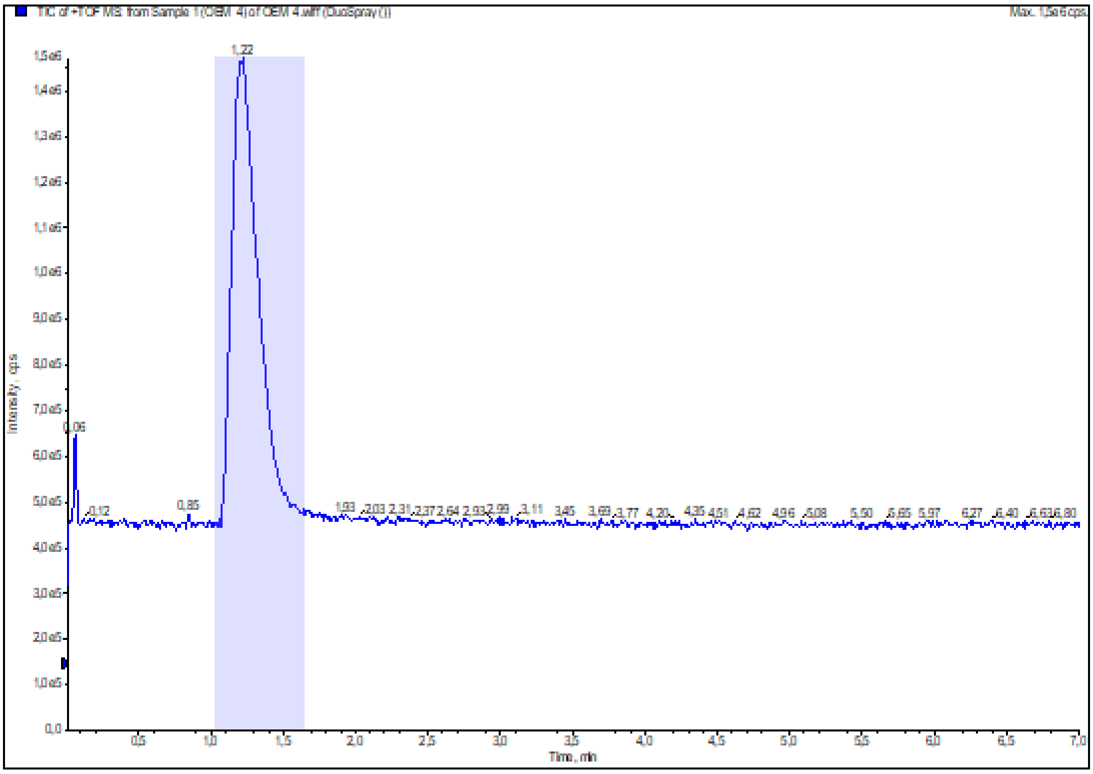

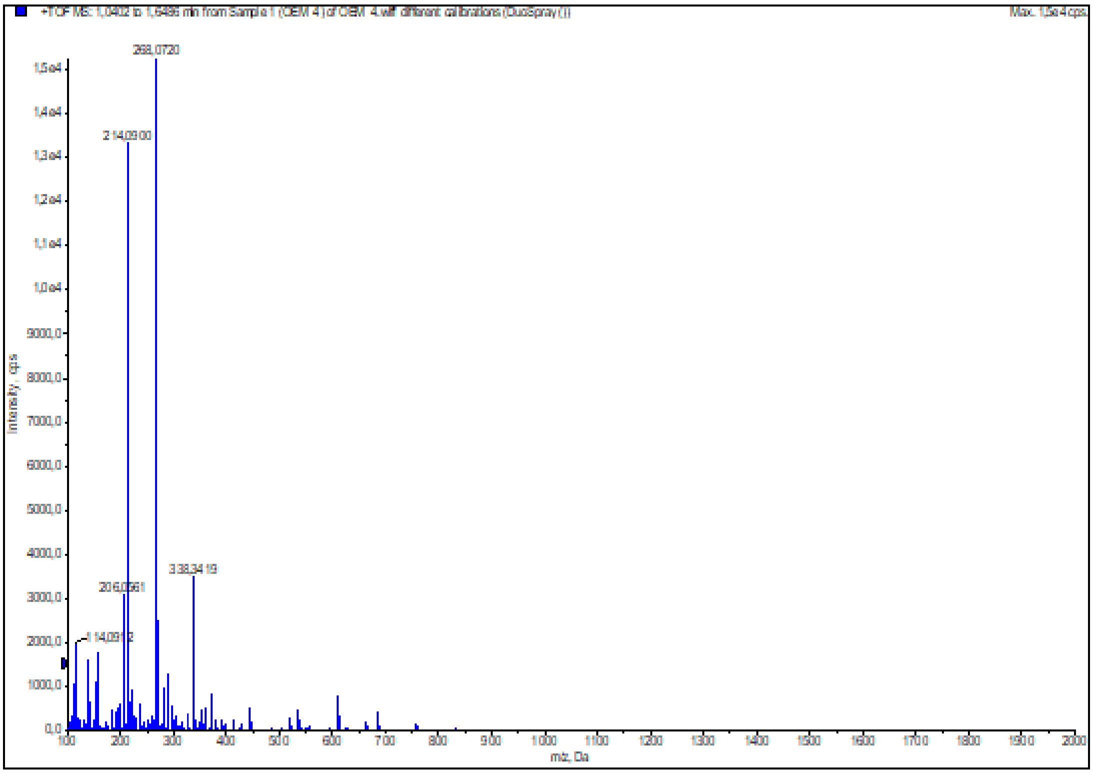

**Figure S11:** HRMS (ESI+) of compound **13**

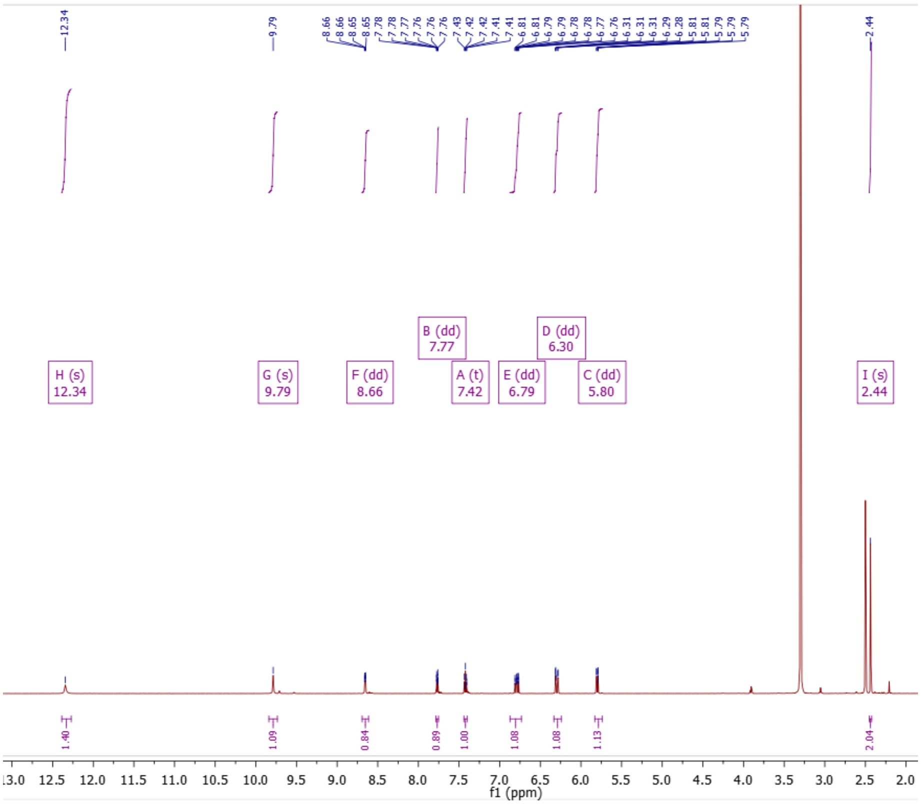

**Figure S12:** $^{1}$H NMR of compound **17**

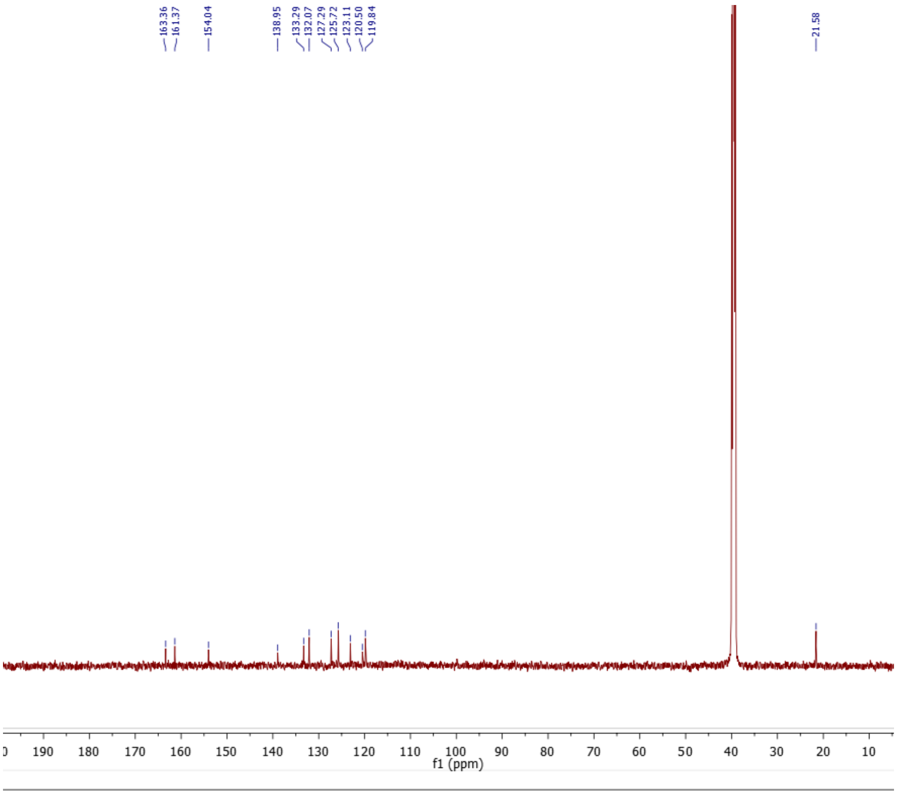

**Figure S13:** $^{13}$C NMR of compound **17**

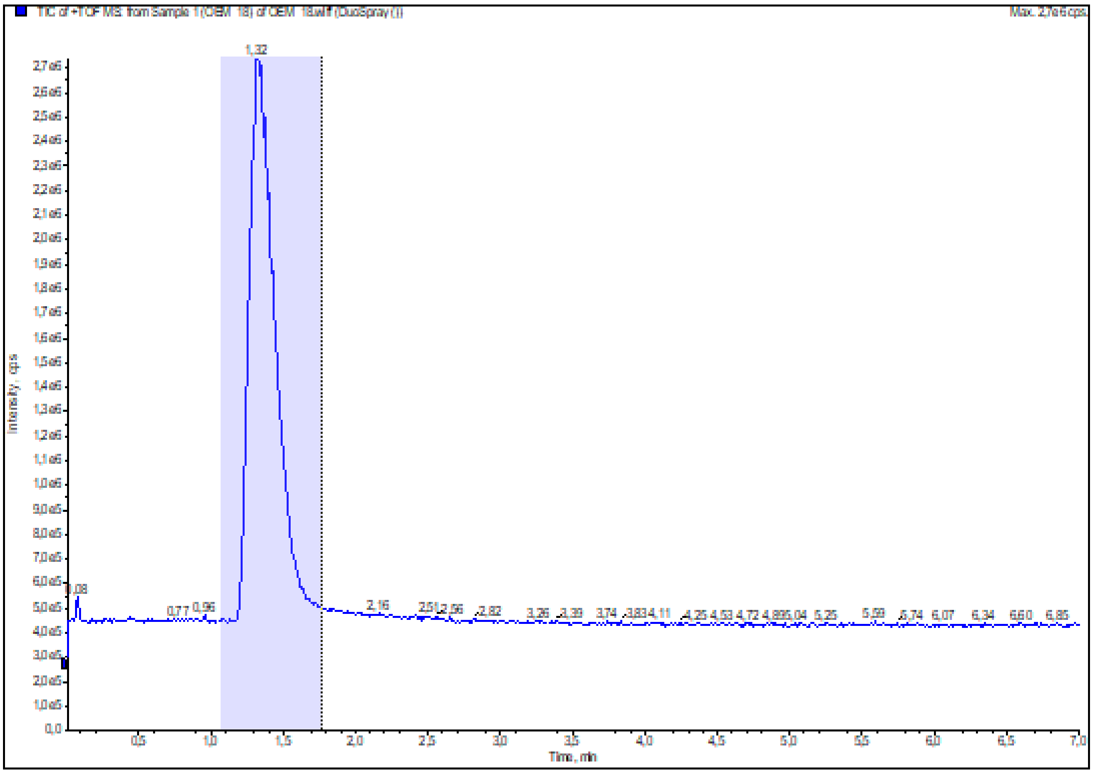

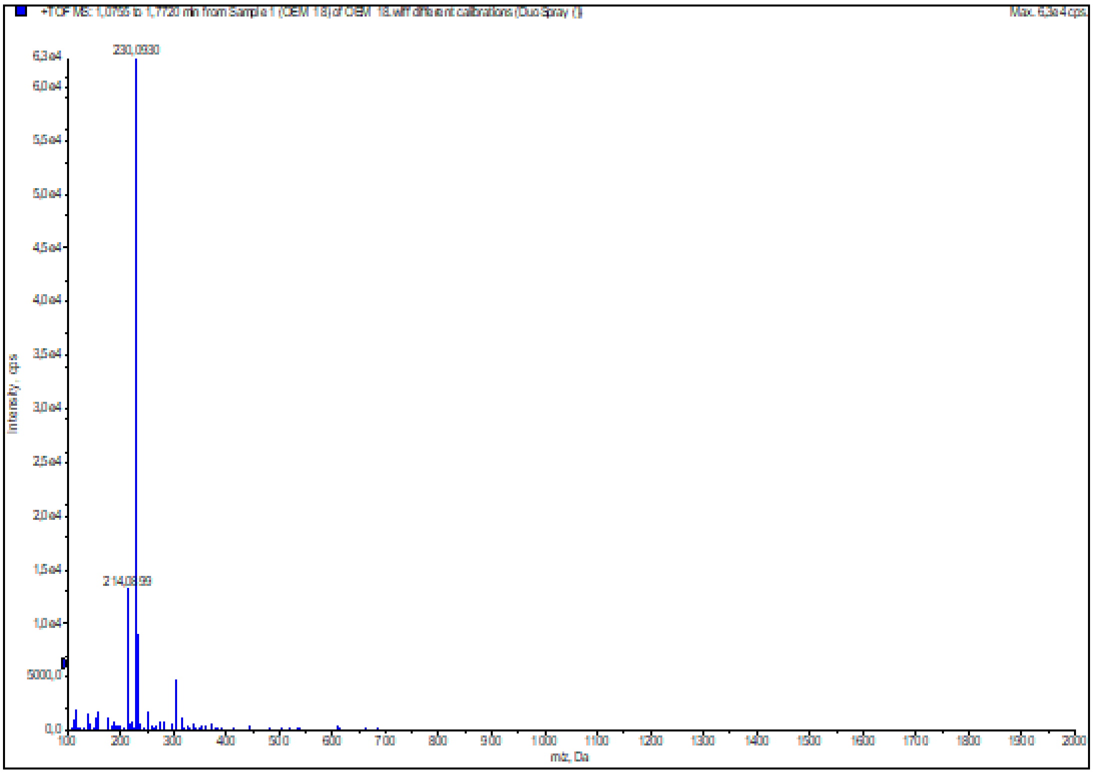

**Figure S14:** HRMS (ESI+) of compound **17**

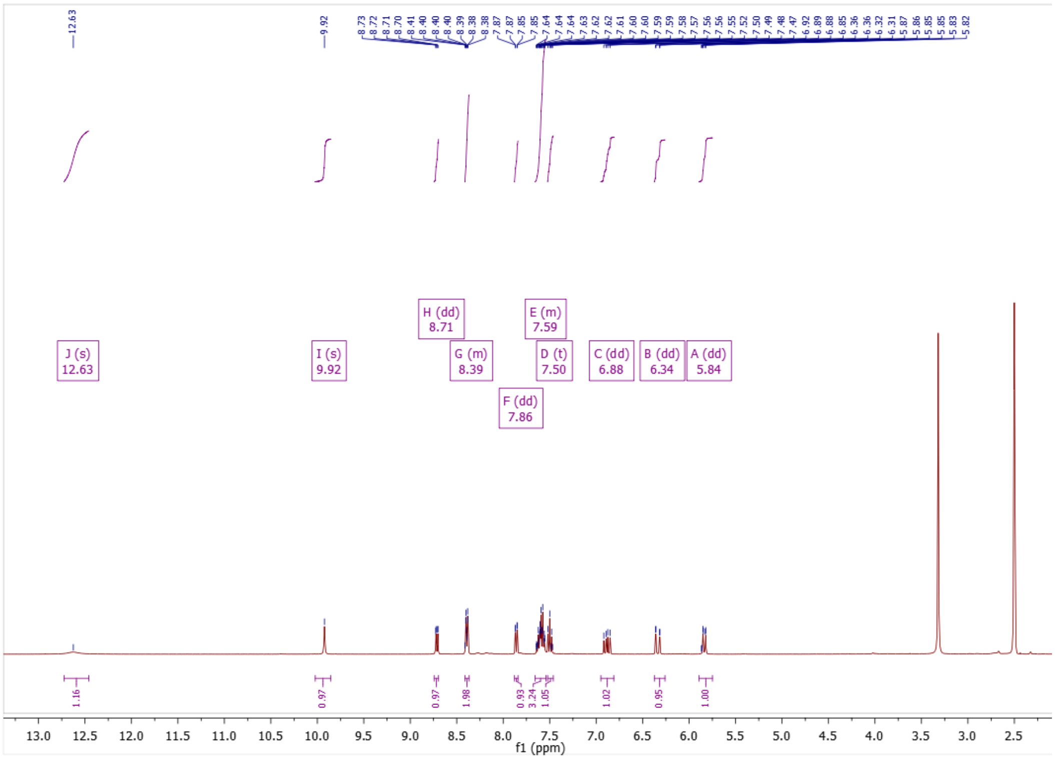

**Figure S15:** $^{1}$H NMR of compound **19**

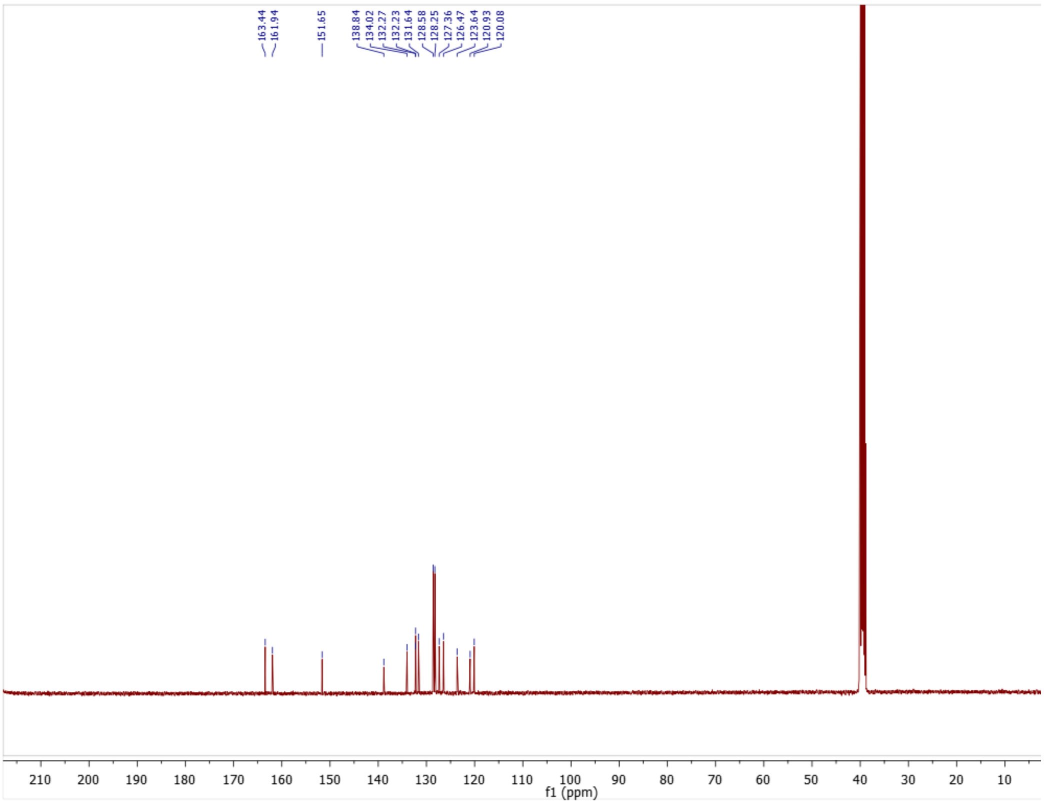

**Figure S16:** $^{13}$C NMR of compound **19**

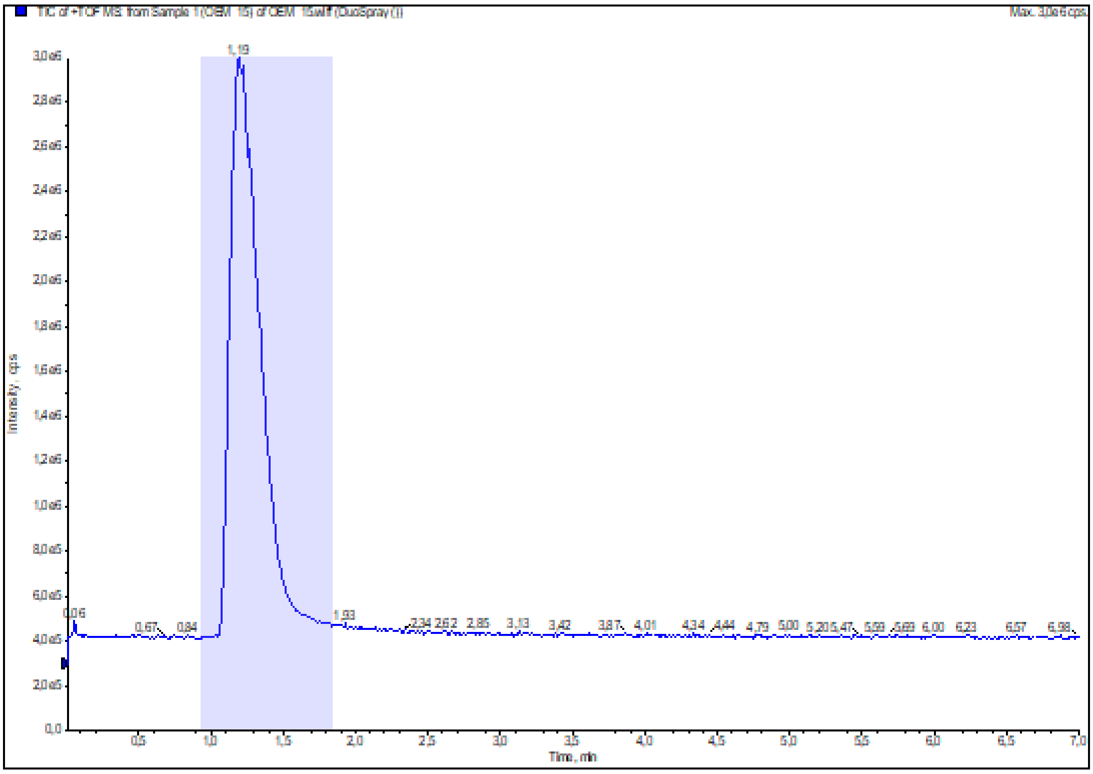

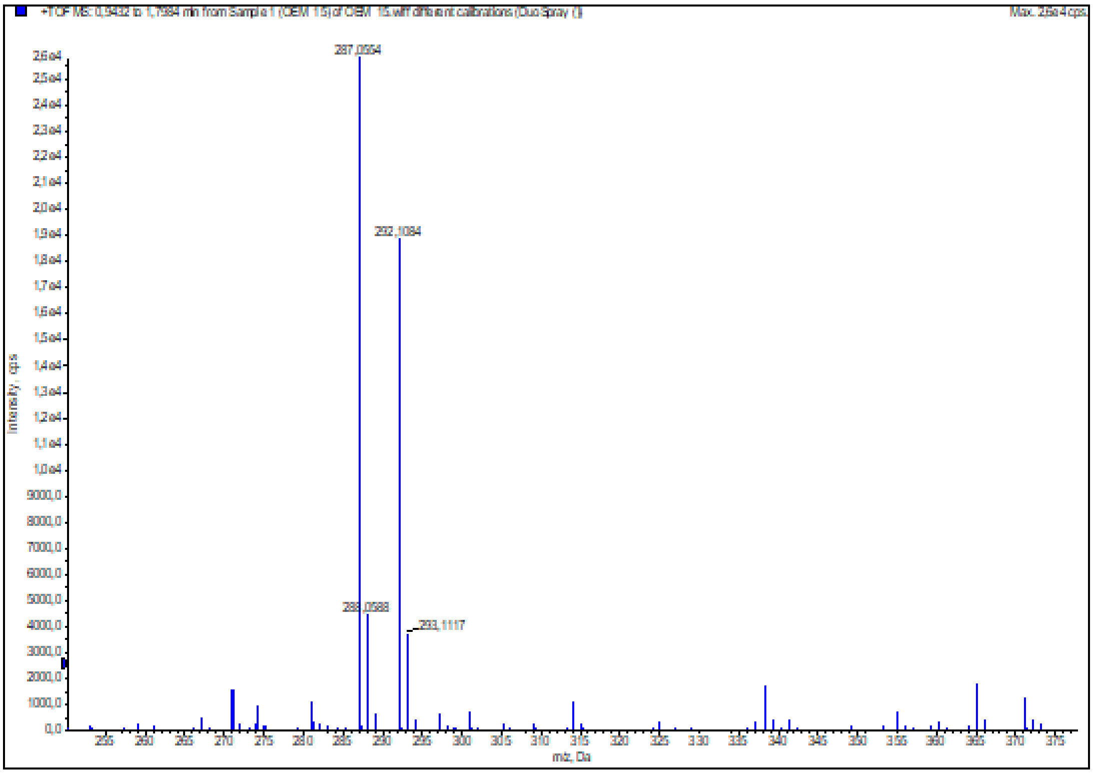

**Figure S17:** HRMS (ESI+) of compound **19**

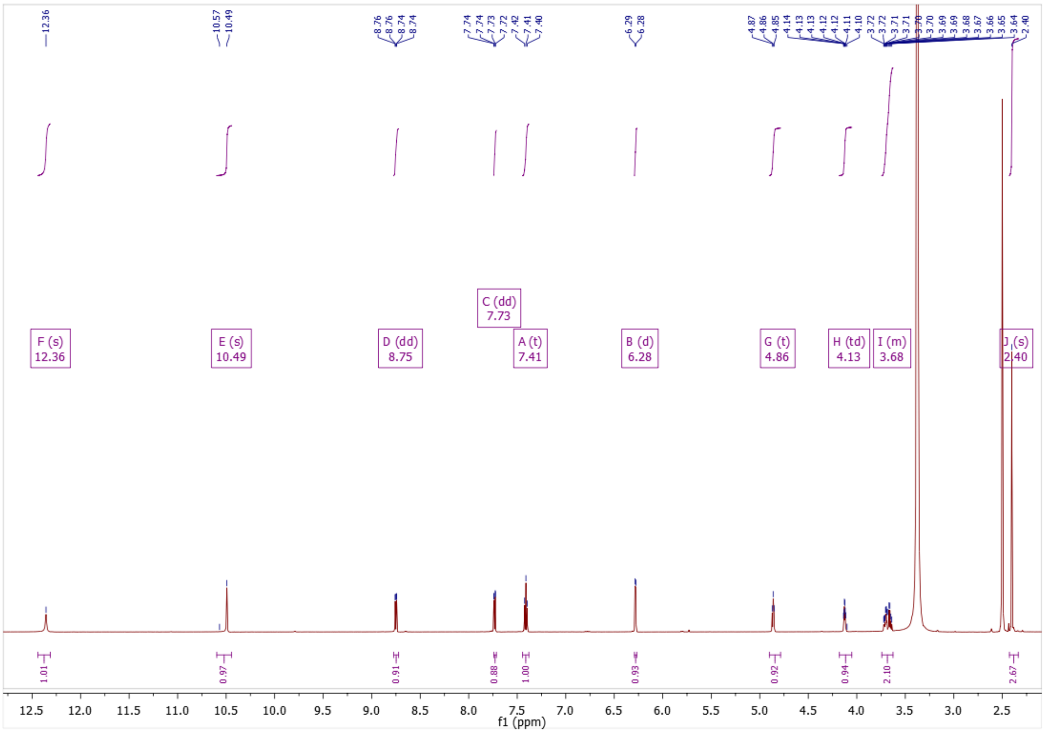

**Figure S18:** $^{1}$H NMR of compound **20**

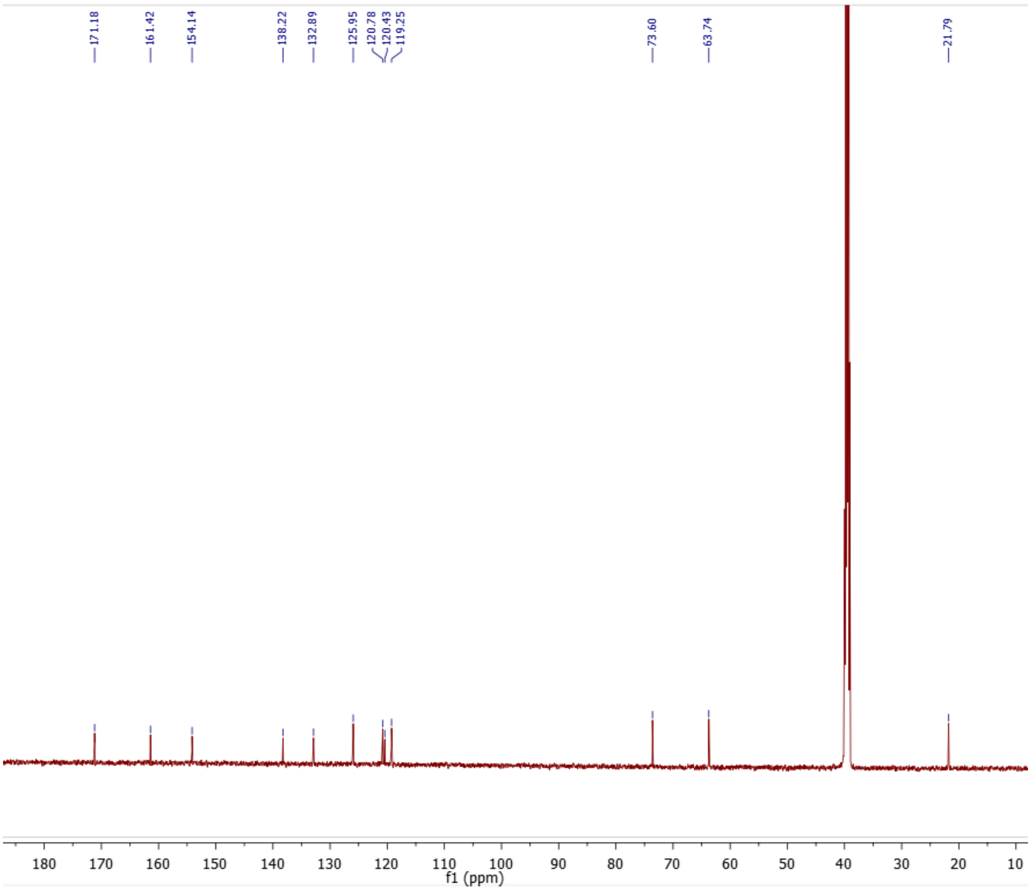

**Figure S19:** $^{13}$C NMR of compound **20**

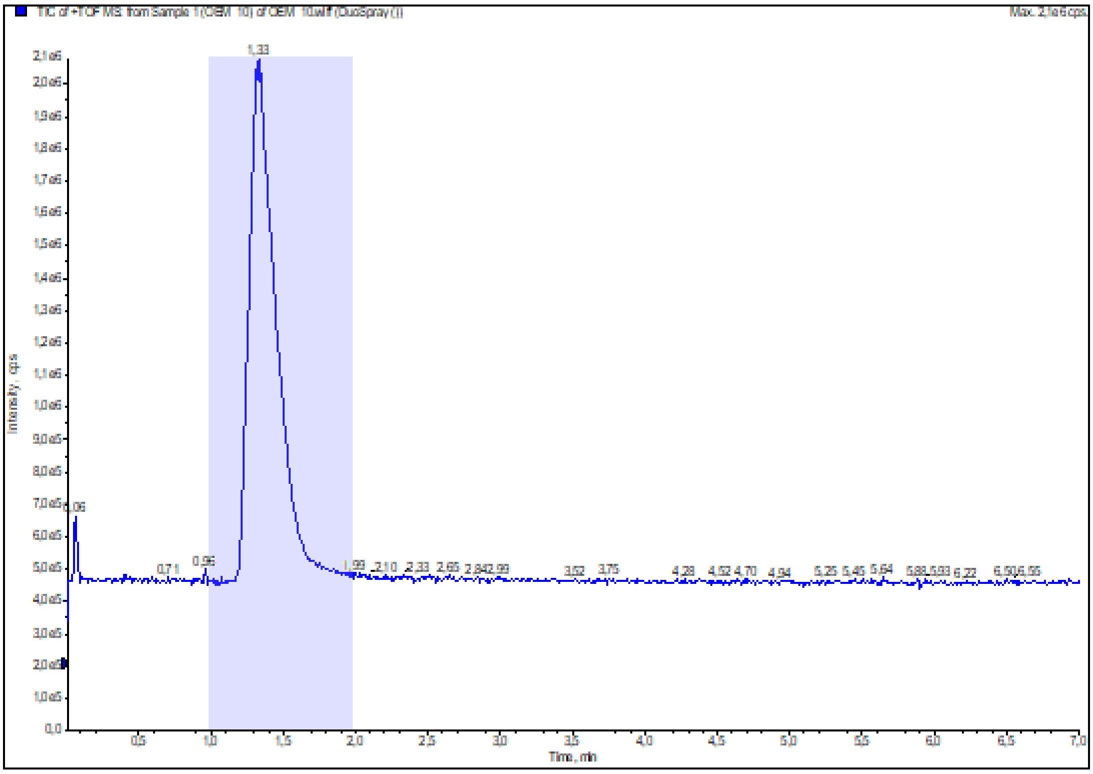

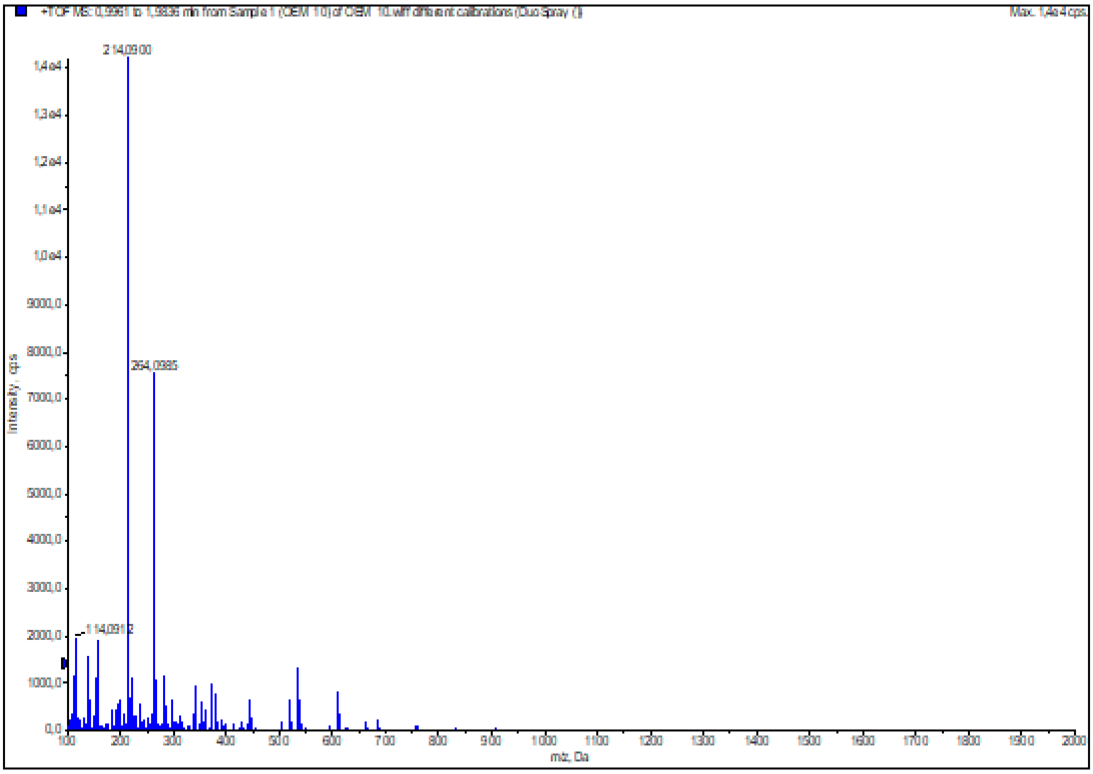

**Figure S20:** HRMS (ESI+) of compound **20**

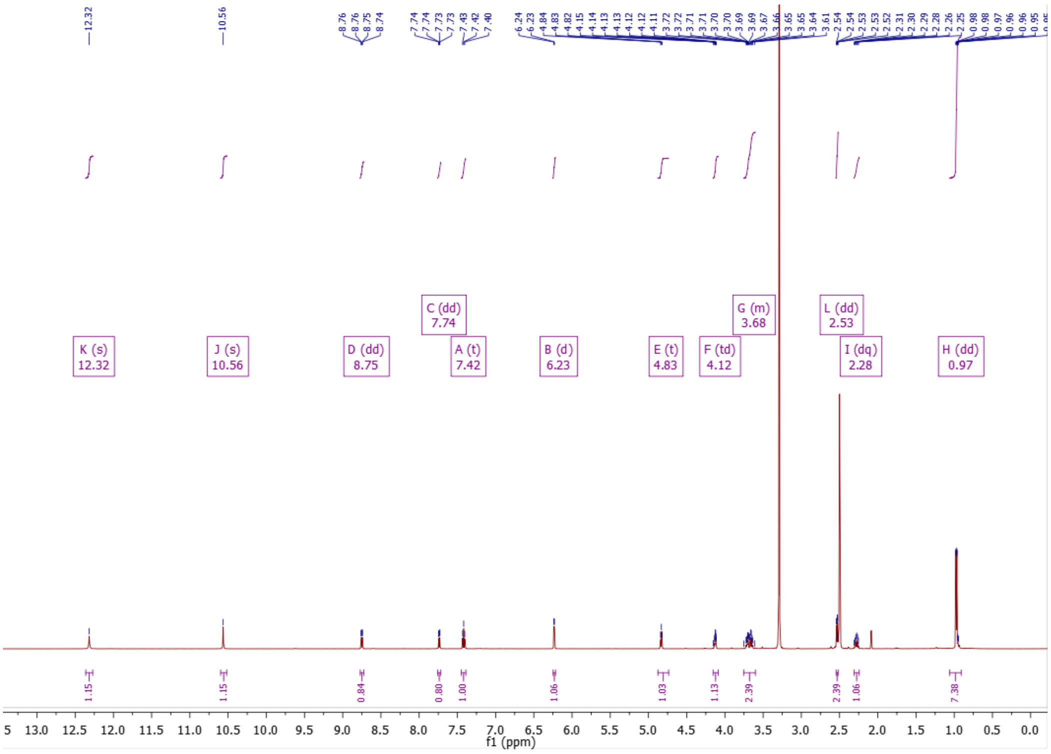

**Figure S21:** $^{1}$H NMR of compound **21**

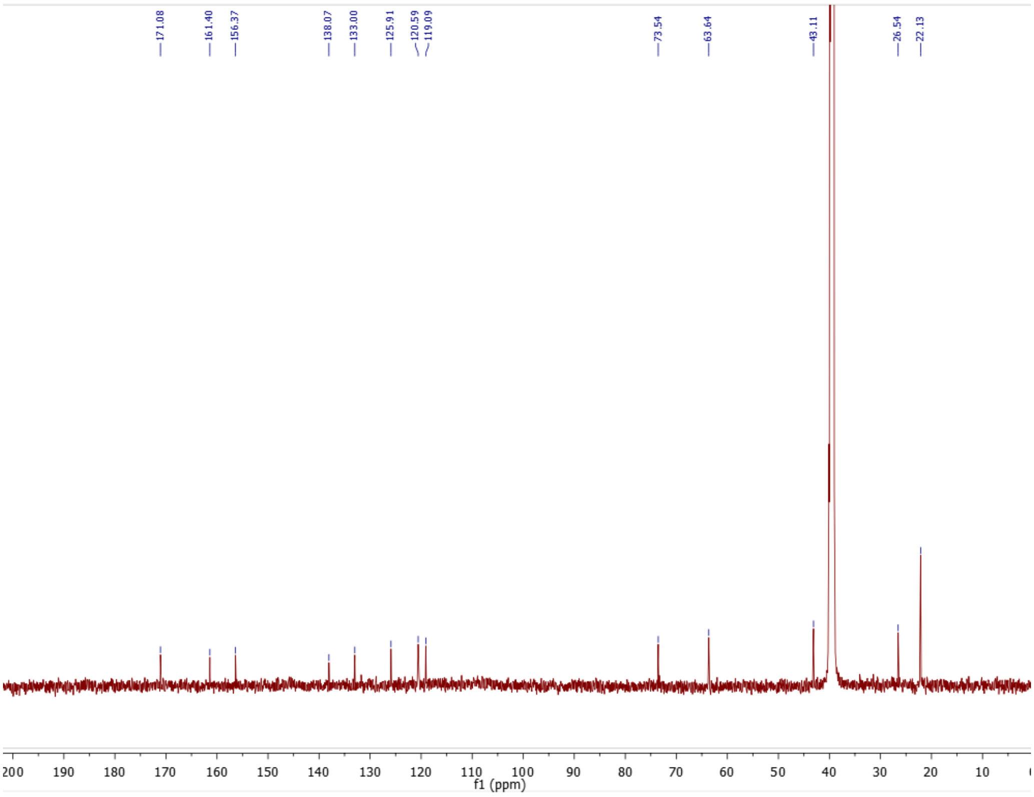

**Figure S22:** $^{13}$C NMR of compound **21**

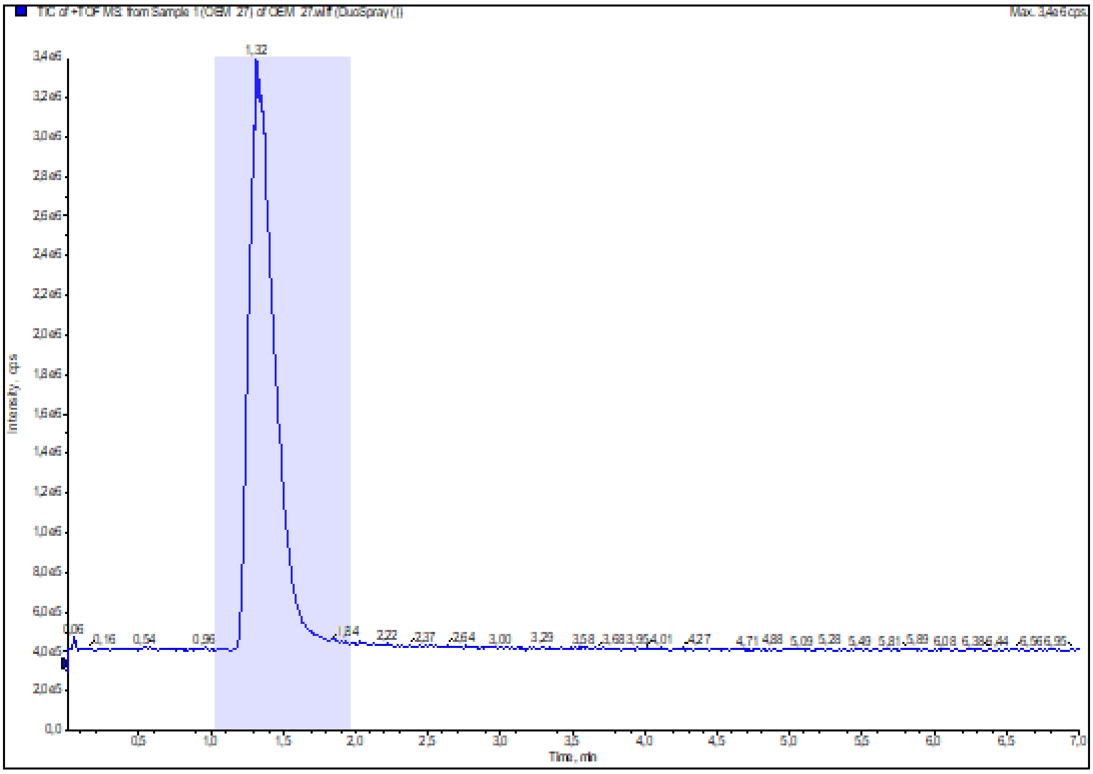

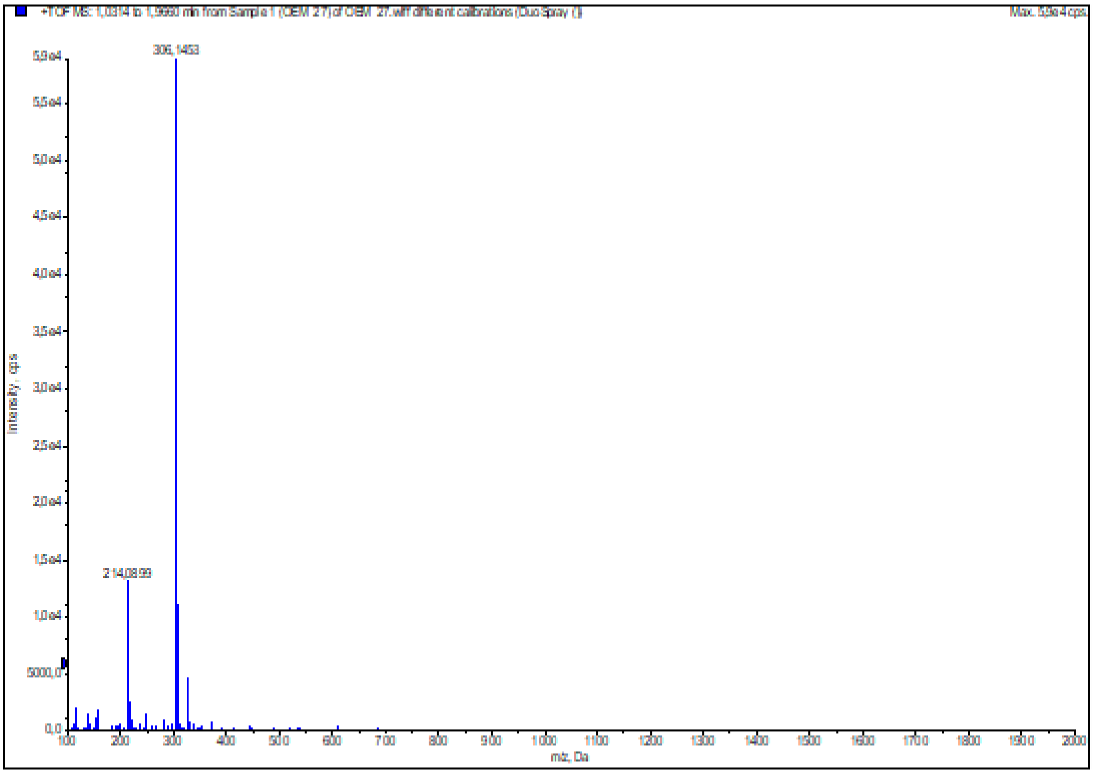

**Figure S23:** HRMS (ESI+) of compound **21**

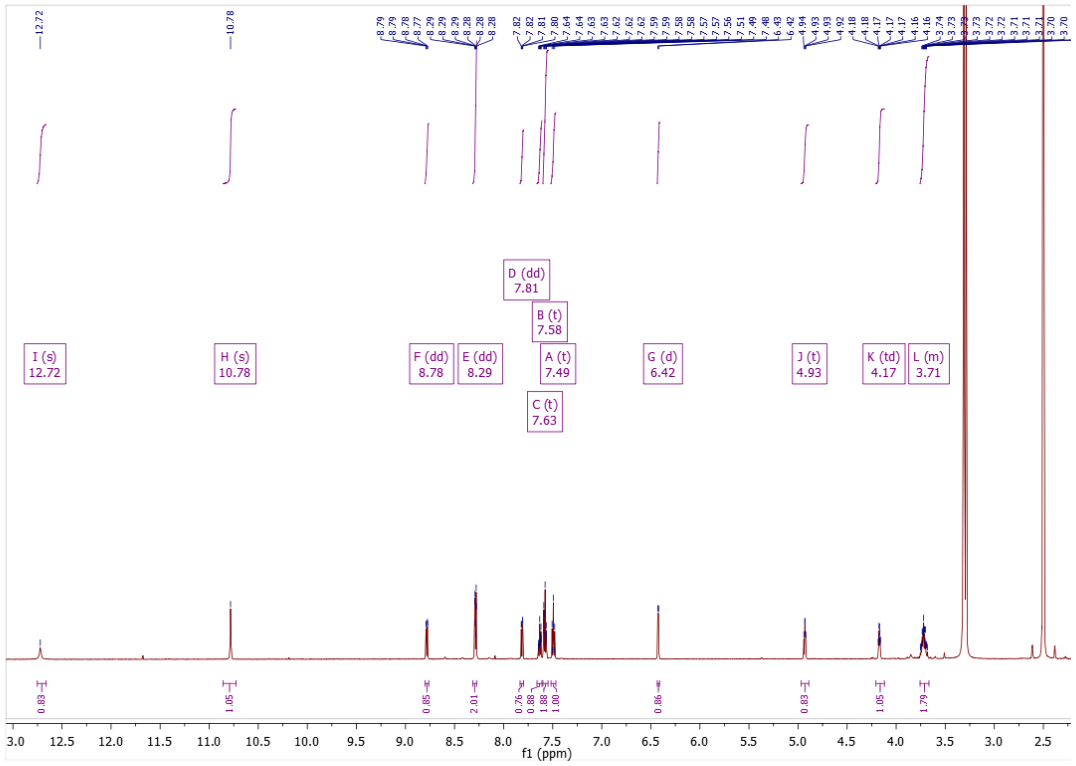

**Figure S24:** $^{1}$H NMR of compound **22**

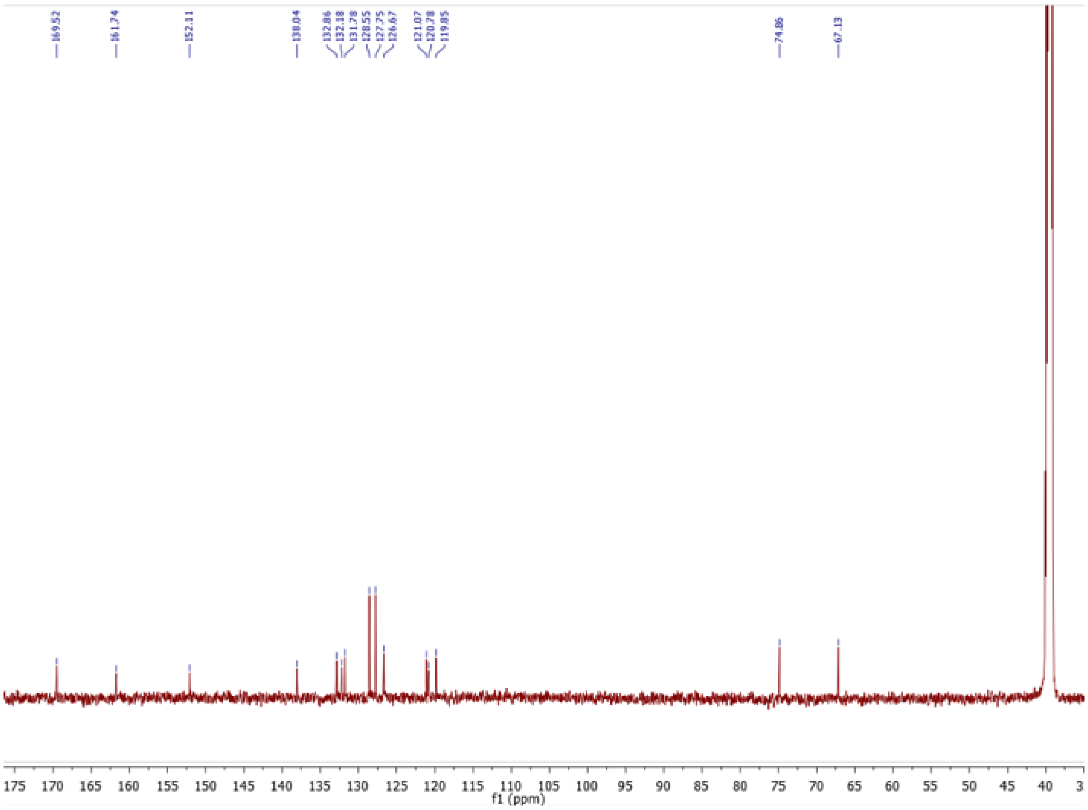

**Figure S25:** $^{13}$C NMR of compound **22**

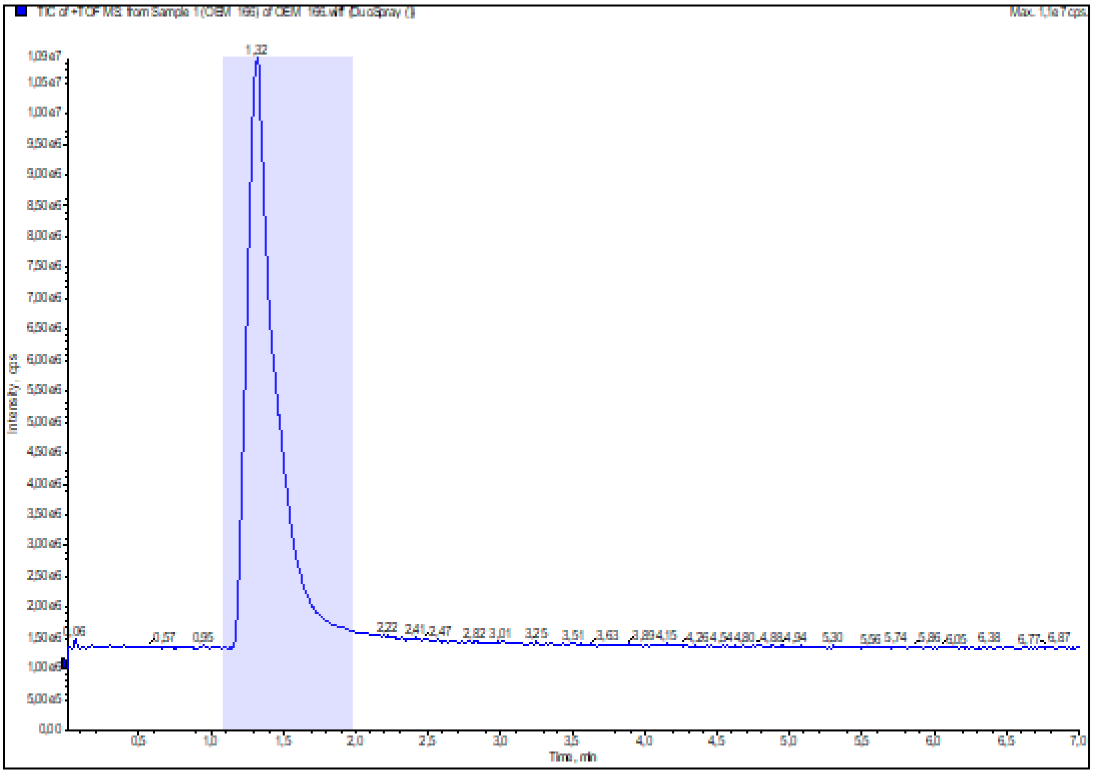

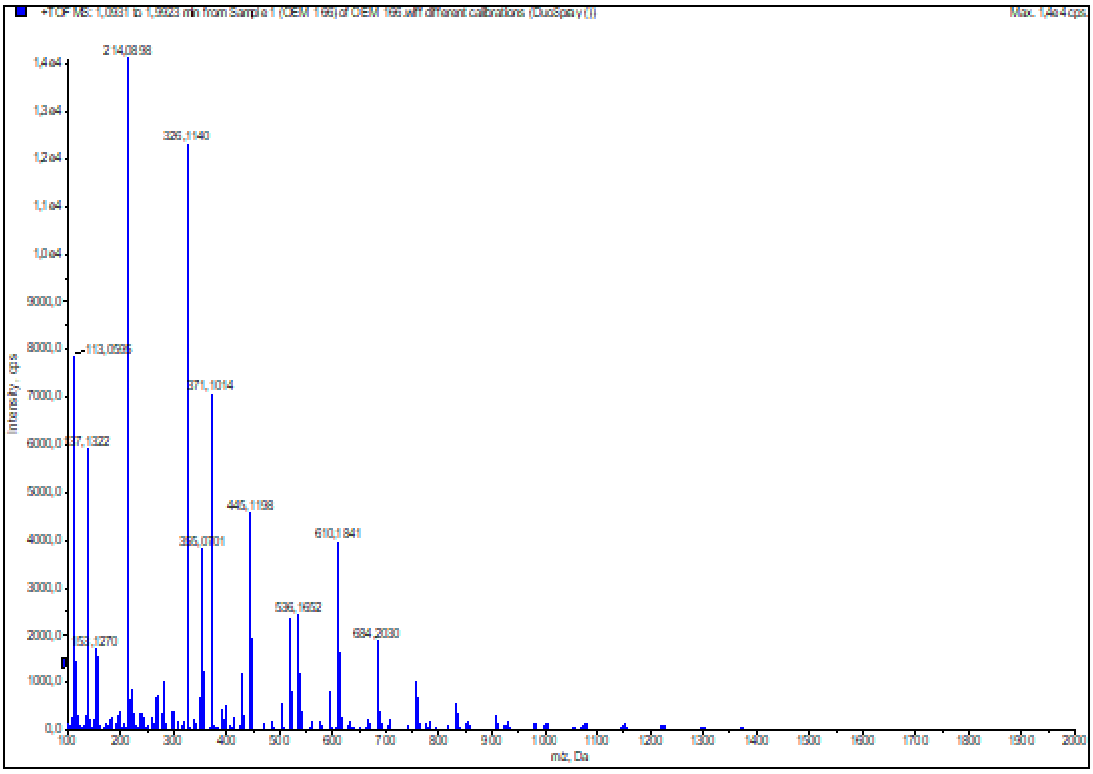

**Figure S26:** HRMS (ESI+) of compound **22**

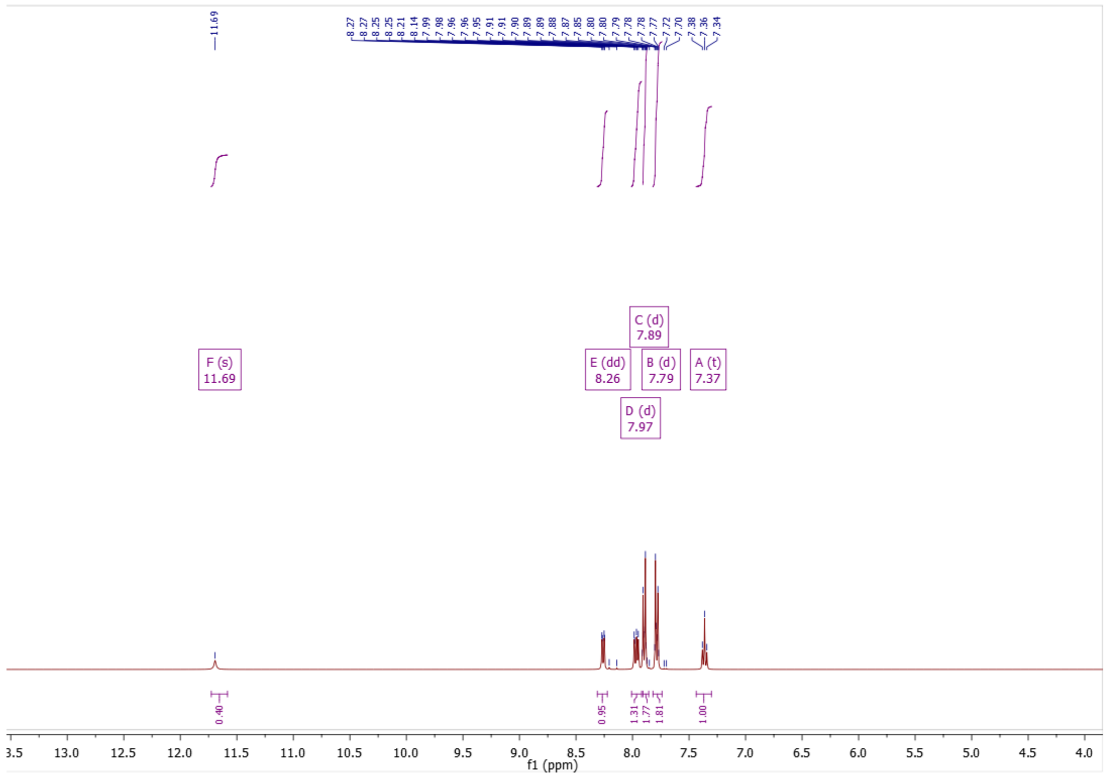

**Figure S27:** $^{1}$H NMR of compound **30**

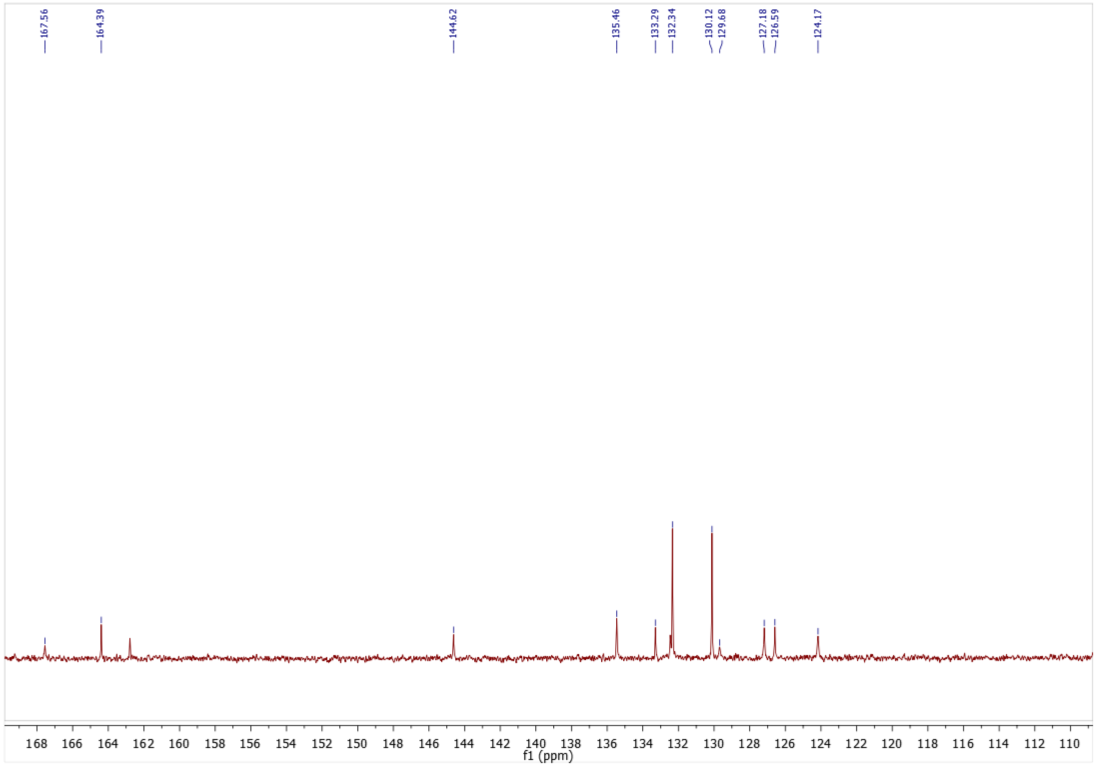

**Figure S28:** $^{13}$C NMR of compound **30**

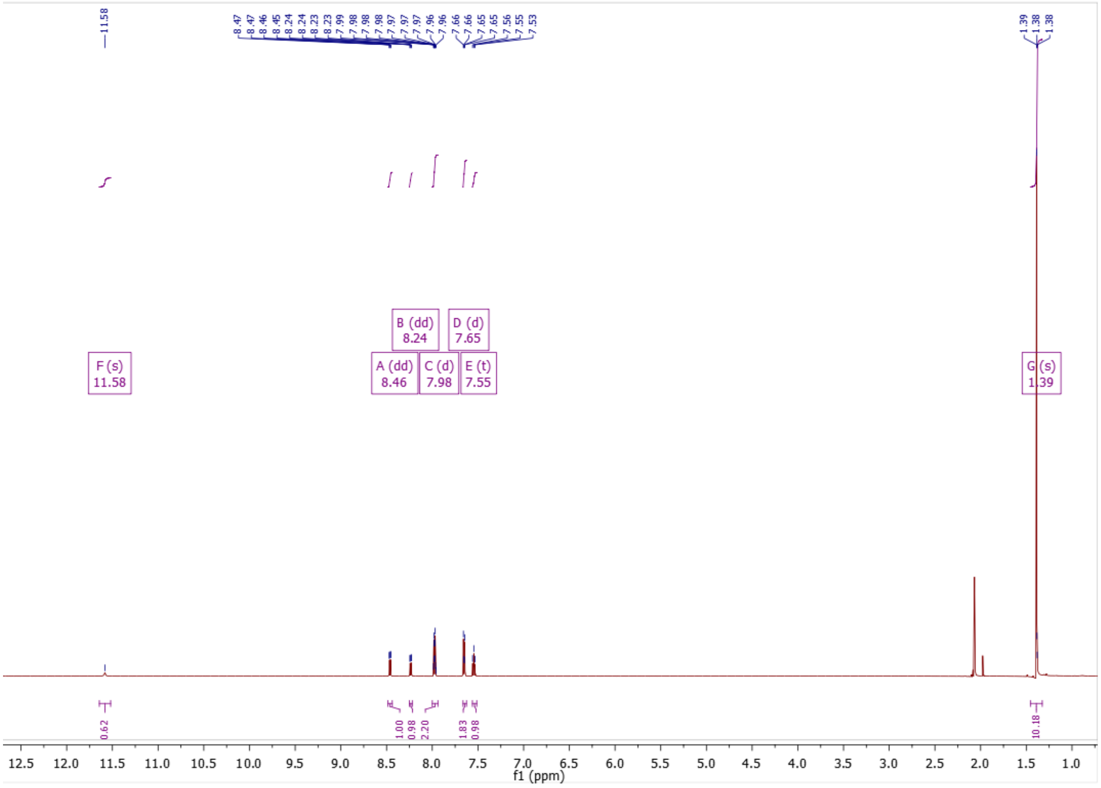

**Figure S29:** $^{1}$H NMR of compound **31**

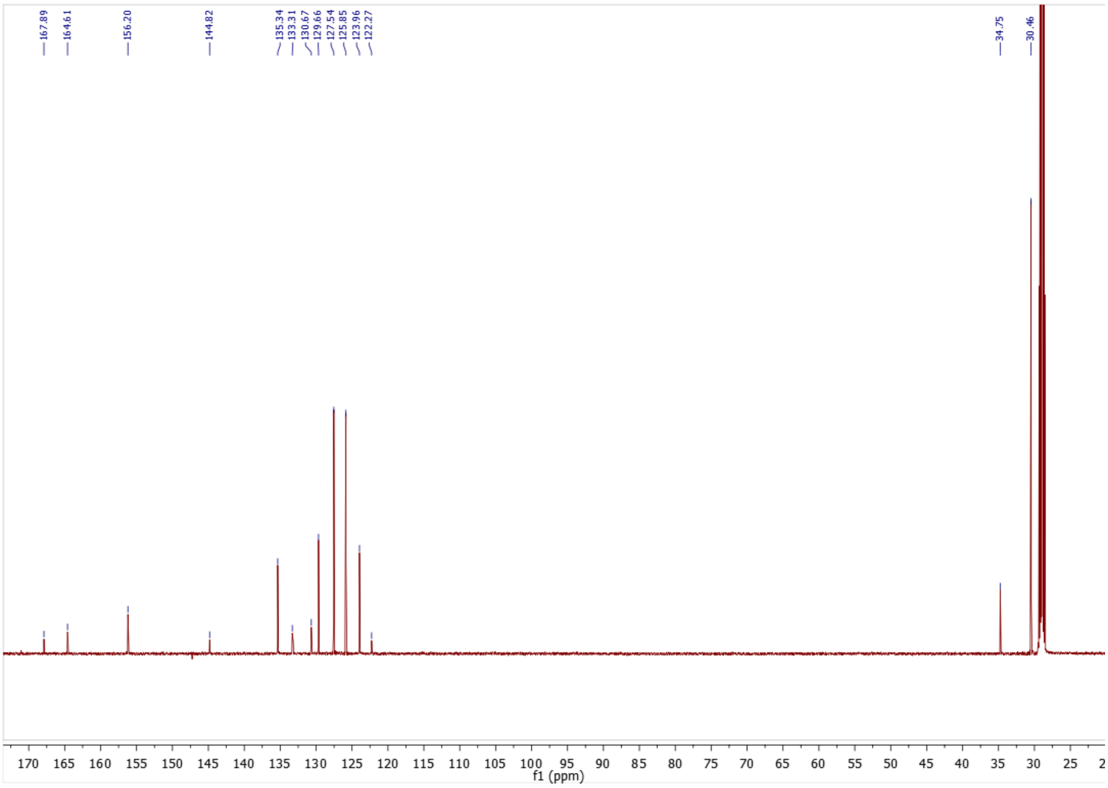

**Figure S30:** $^{13}$C NMR of compound **31**

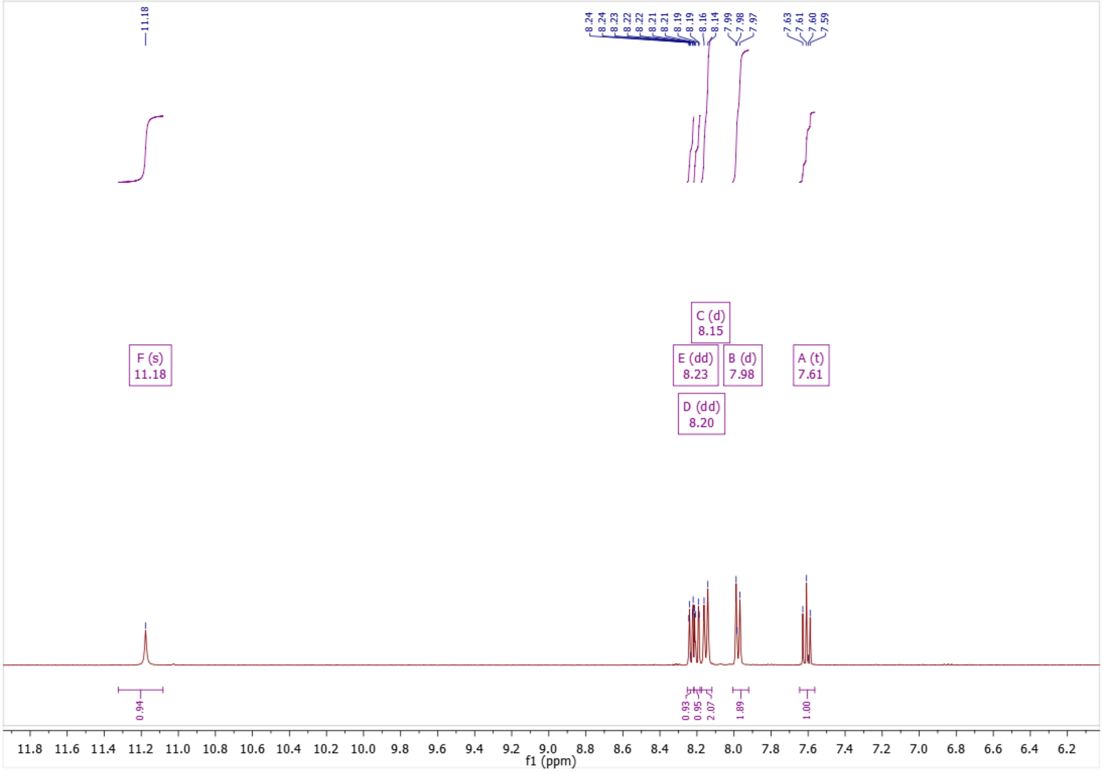

**Figure S31:** $^{1}$H NMR of compound **32**

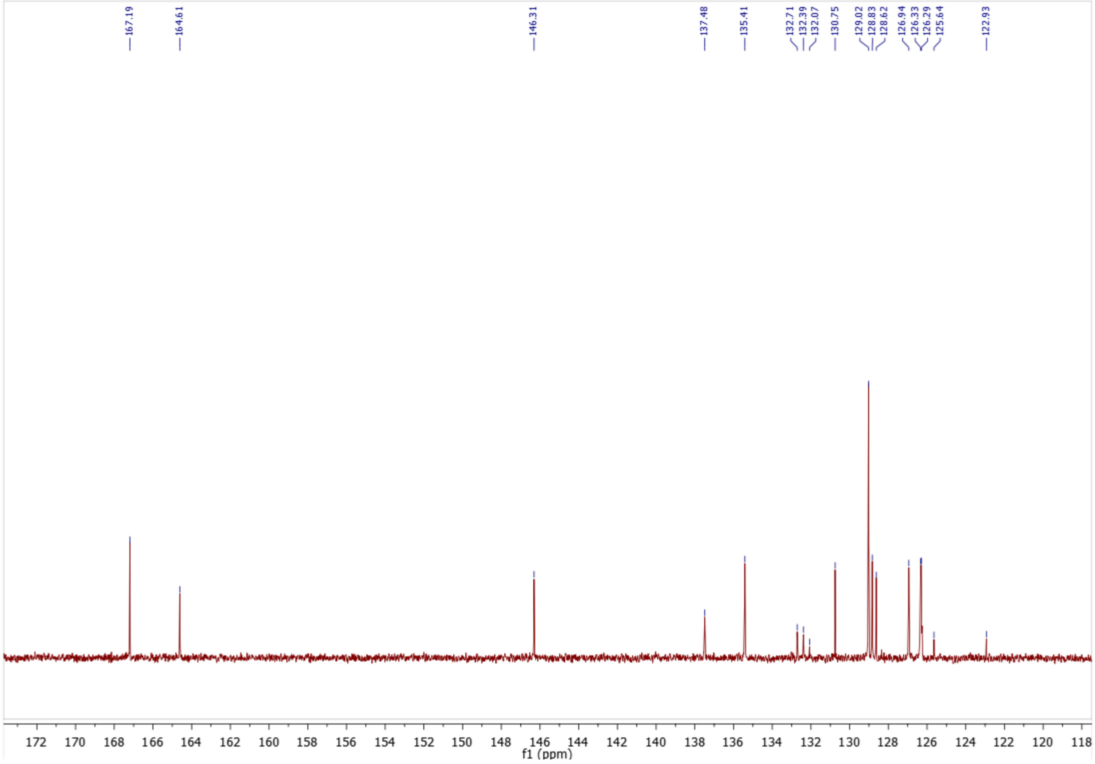

**Figure S32:** $^{13}$C NMR of compound **32**

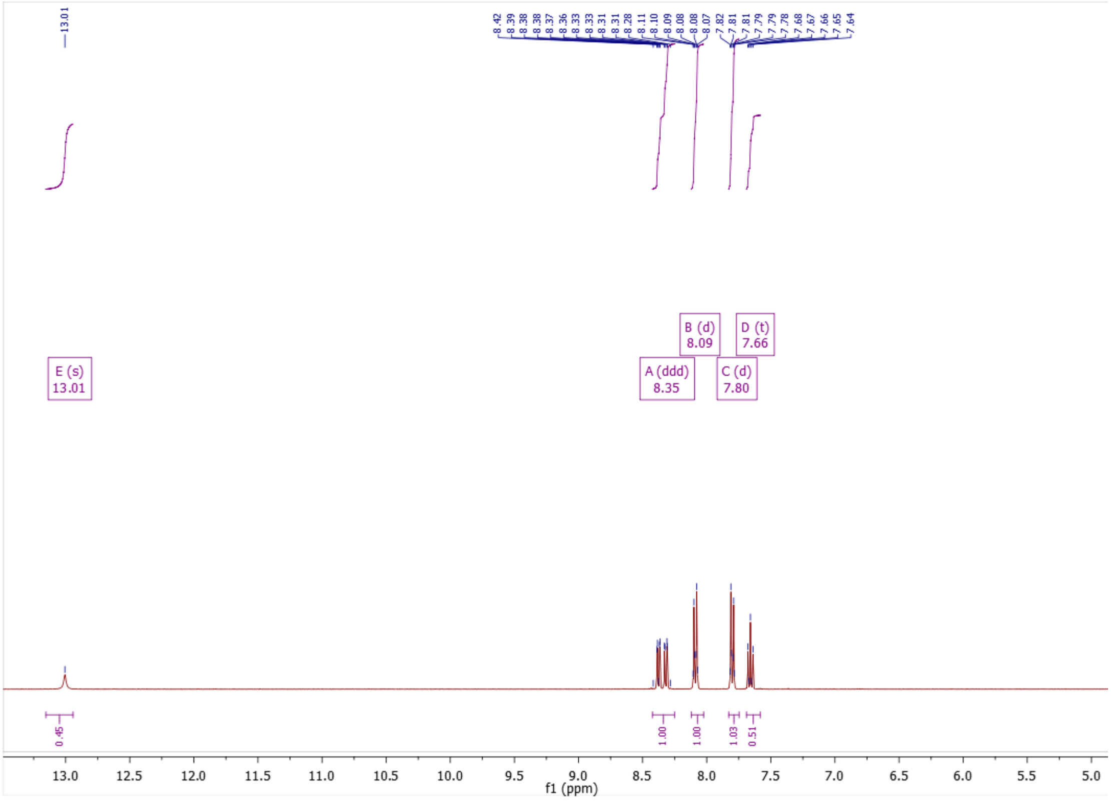

**Figure S33:** $^{1}$H NMR of compound **39**

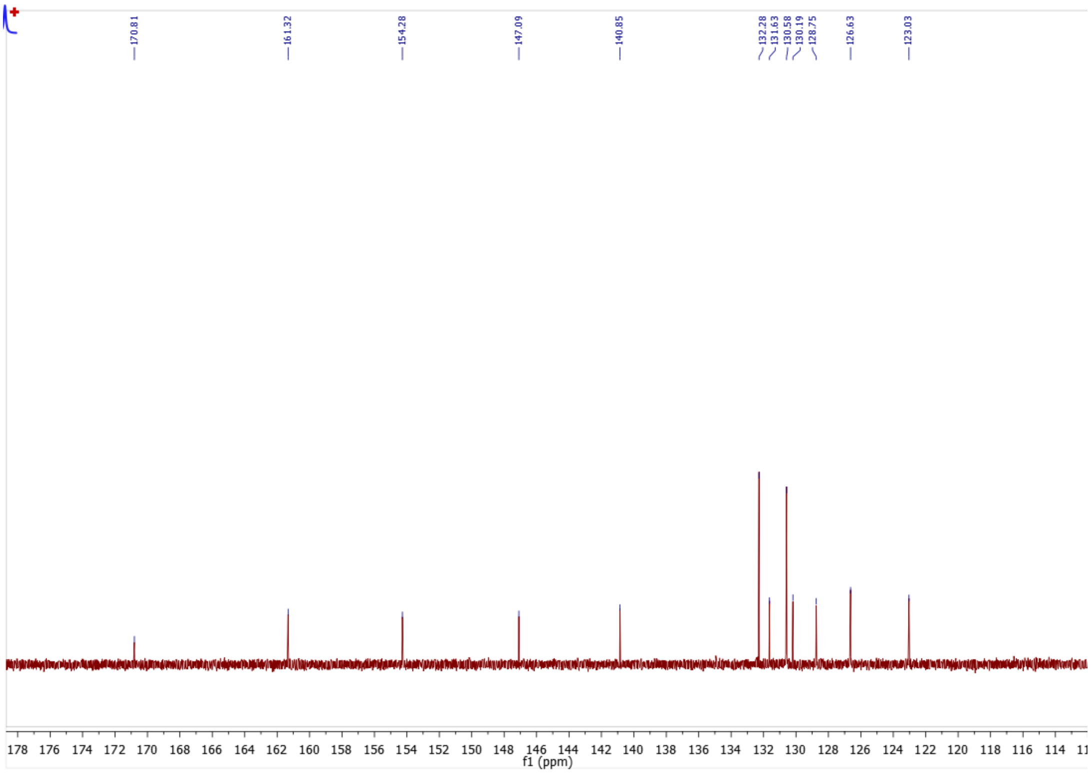

**Figure S34:** $^{13}$C NMR of compound **39**

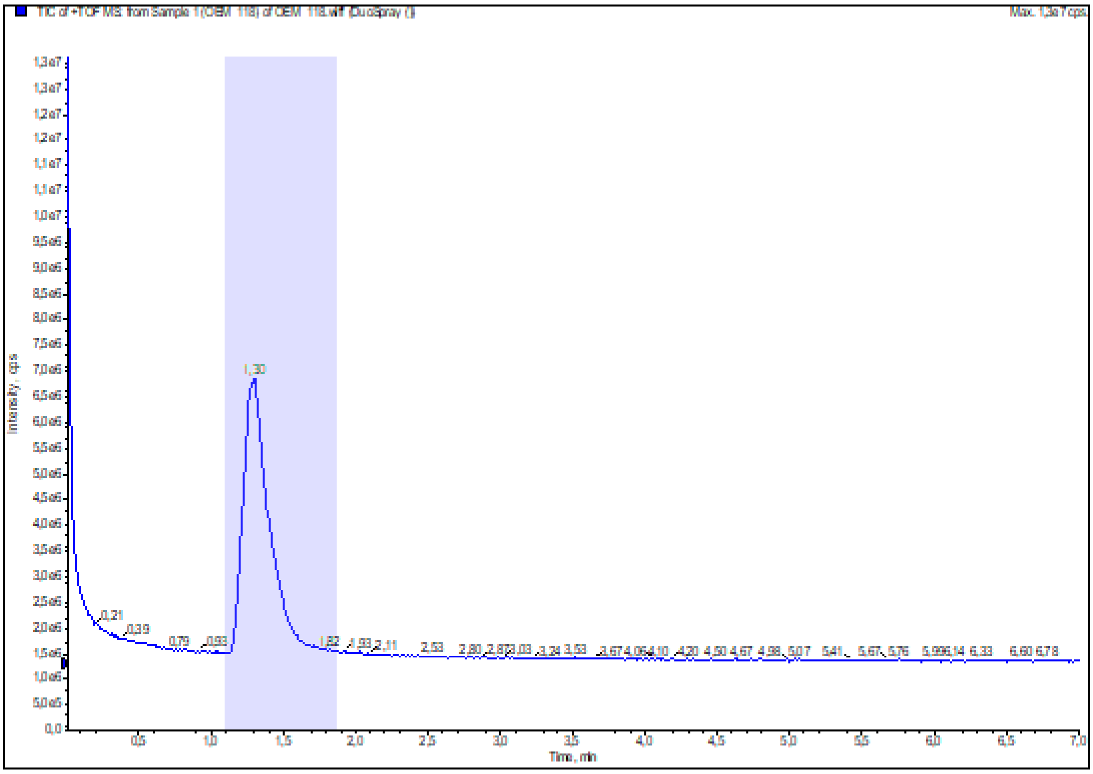

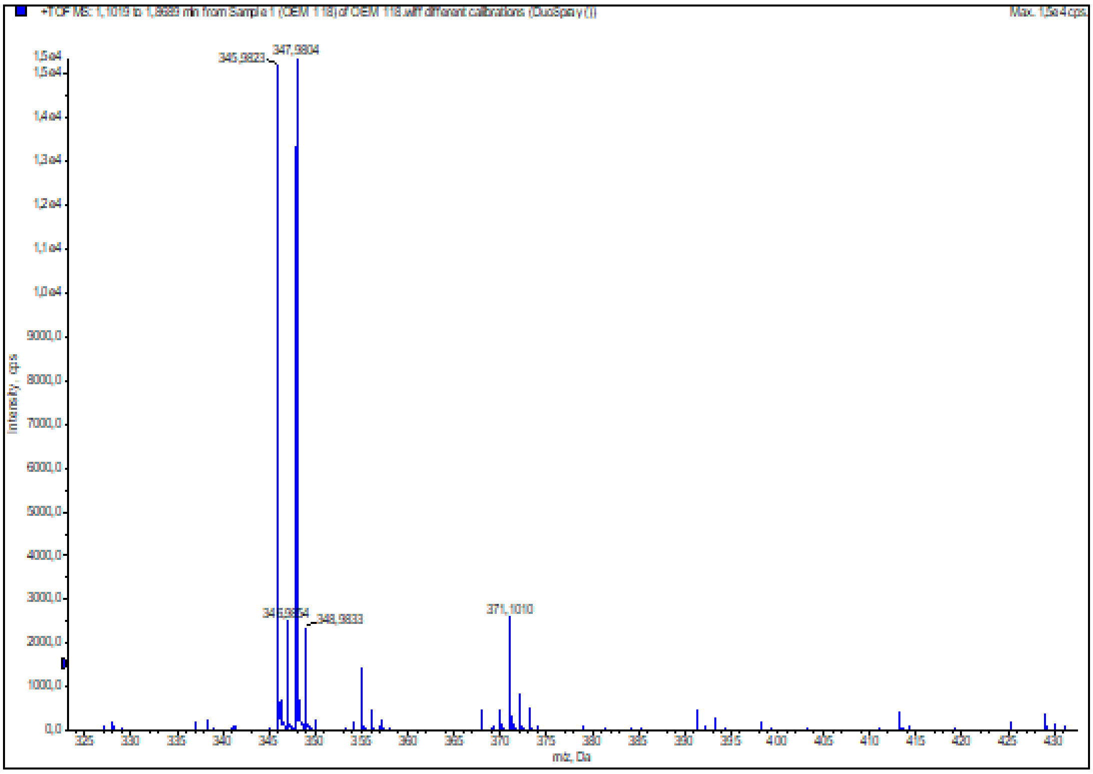

**Figure S35:** HRMS (ESI+) of compound **39**

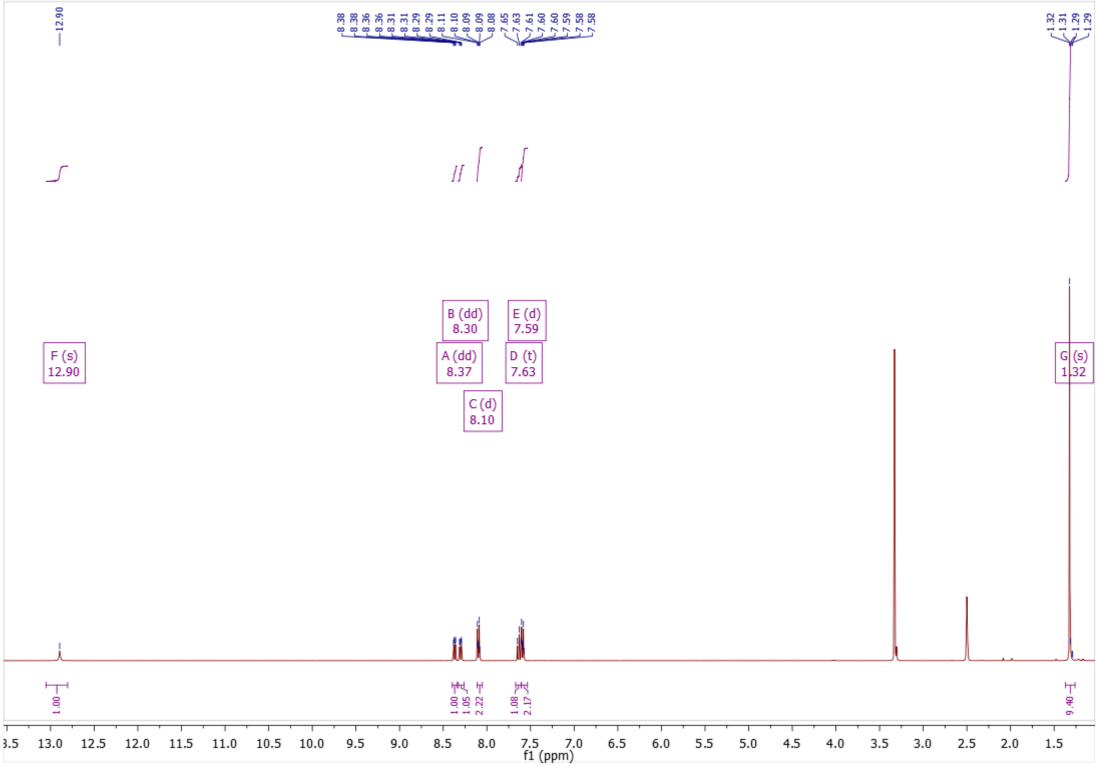

**Figure S36:** $^{1}$H NMR of compound **40**

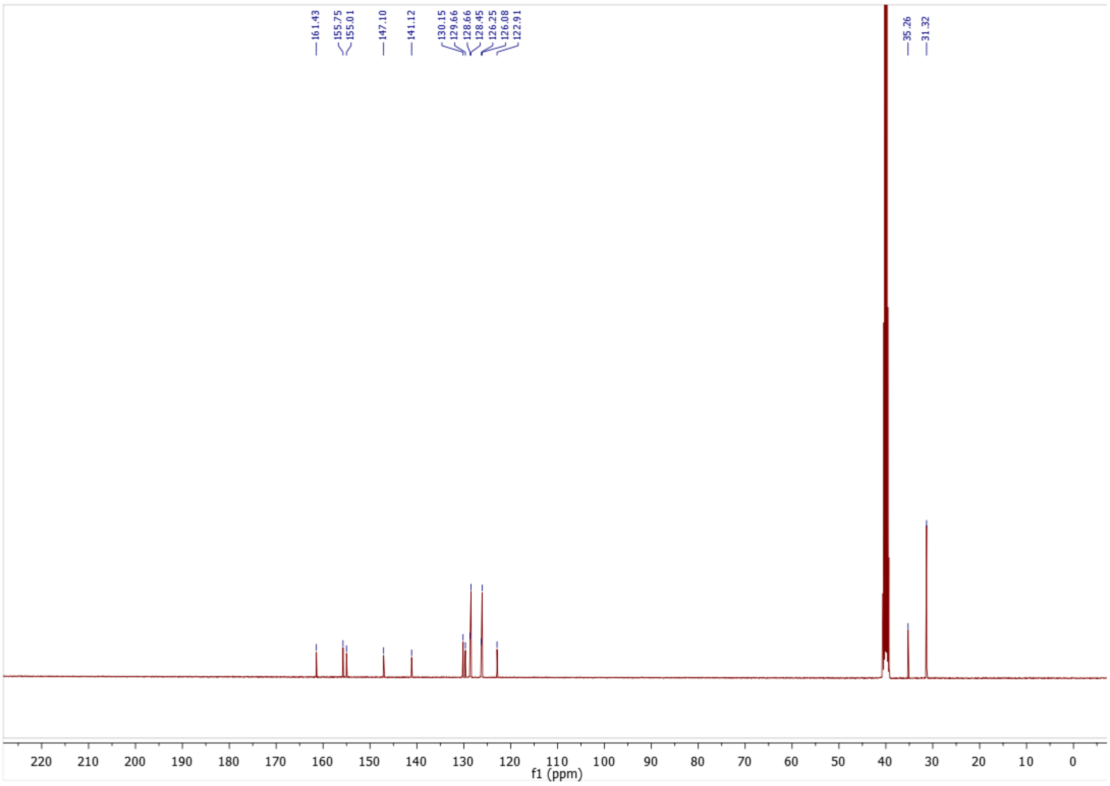

**Figure S37:** $^{13}$C NMR of compound **40**

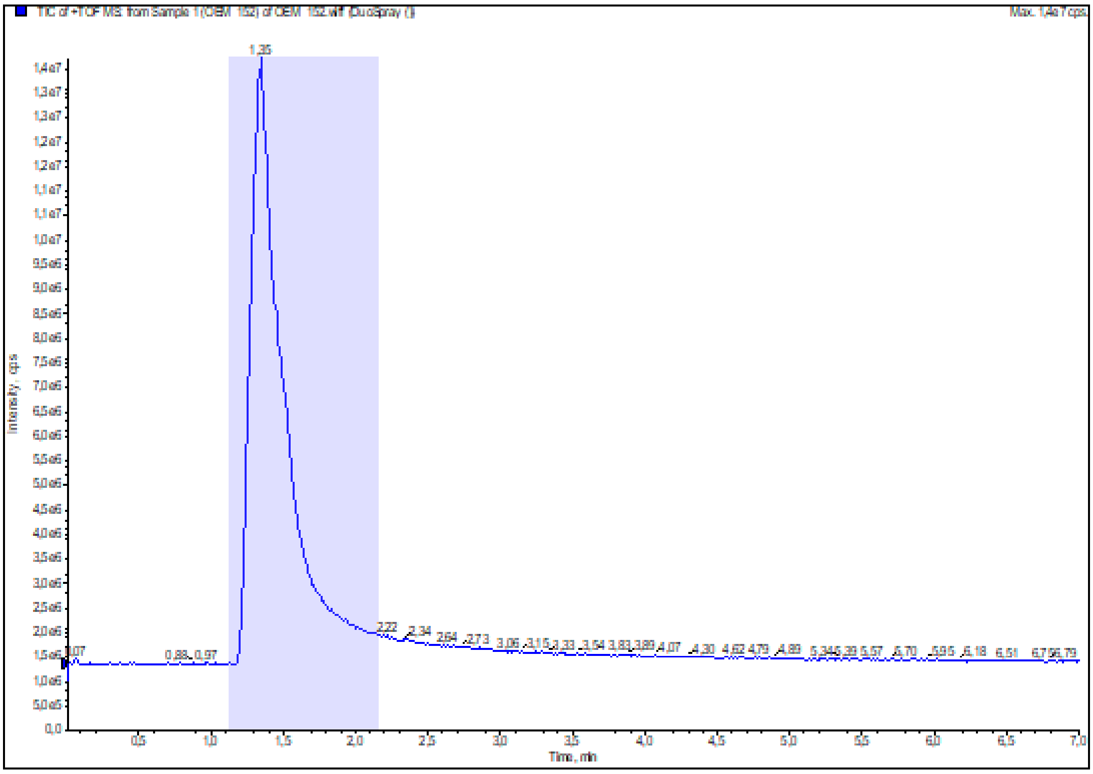

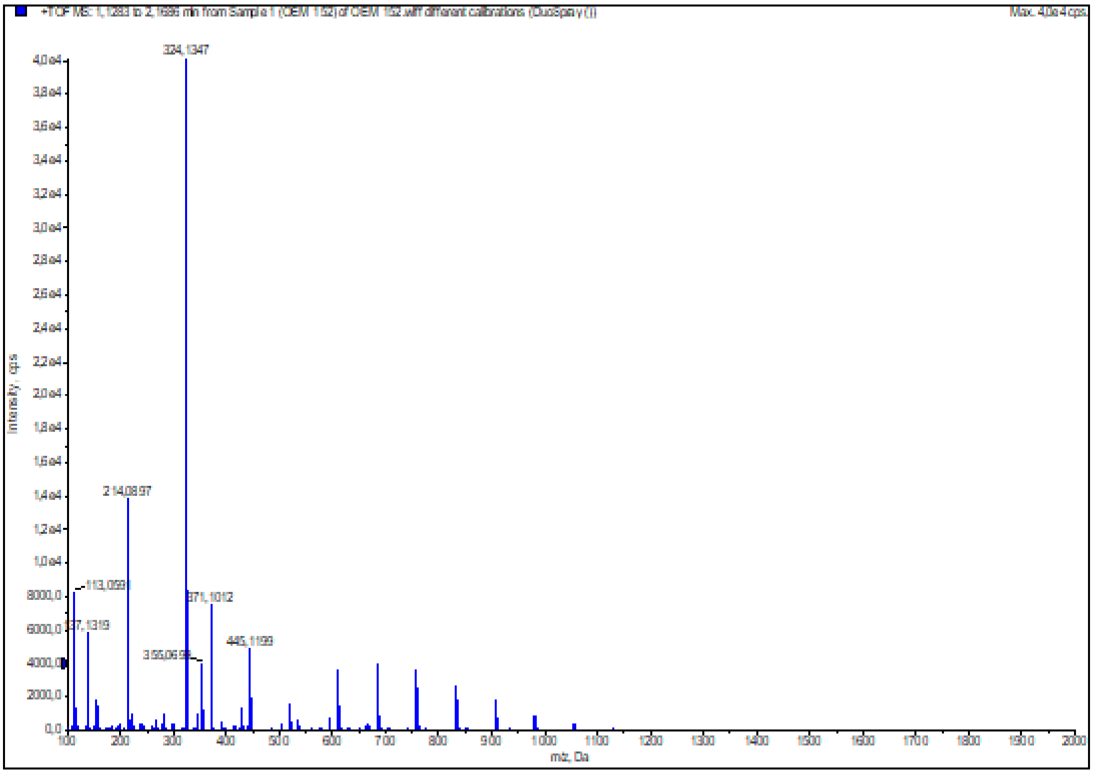

**Figure S38:** HRMS (ESI+) of compound **40**

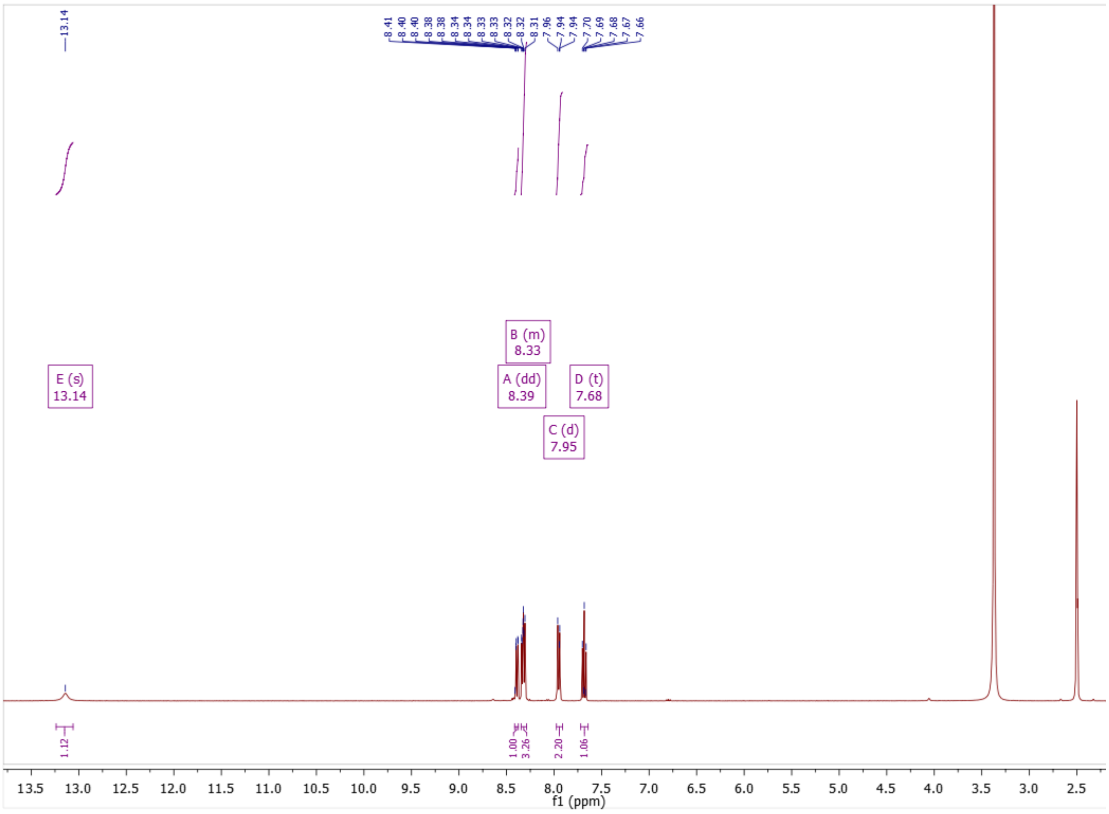

**Figure S39:** $^{1}$H NMR of compound **41**

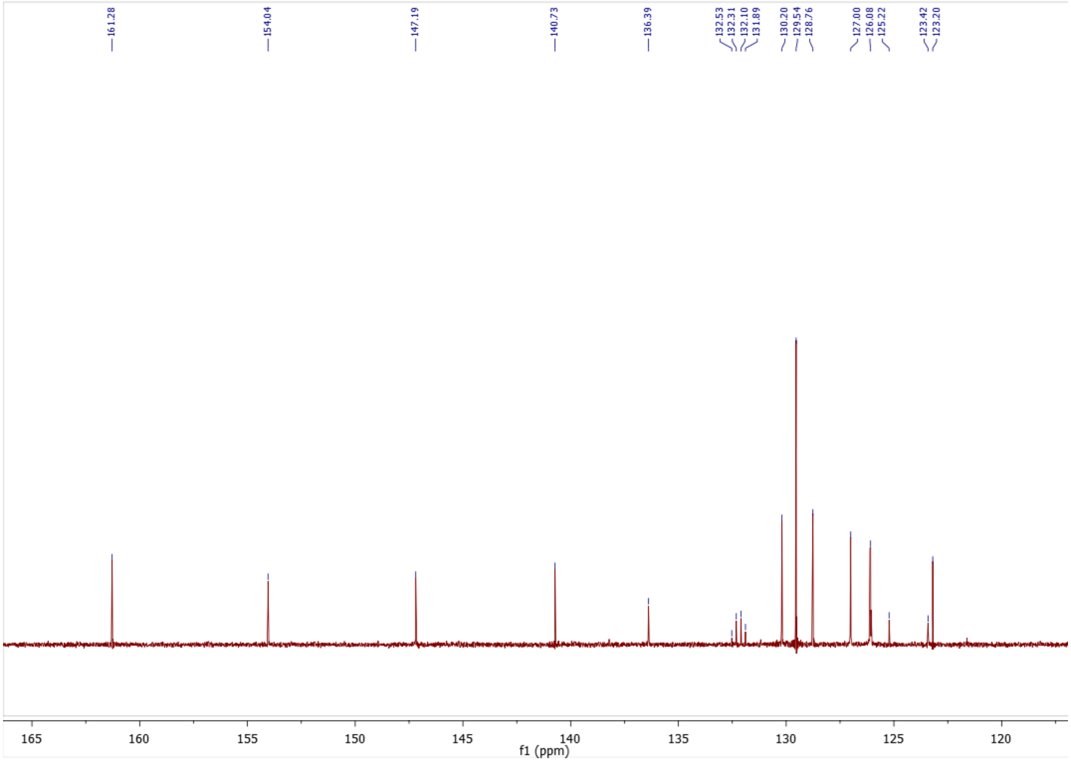

**Figure S40:** $^{13}$C NMR of compound **41**

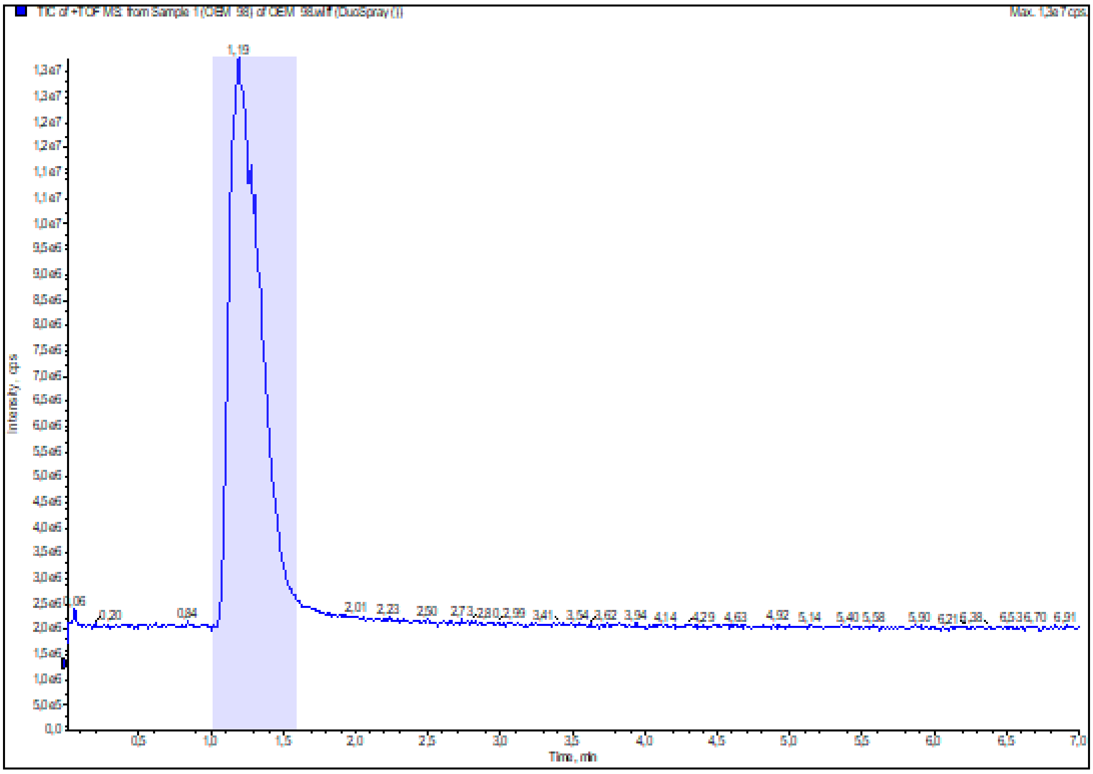

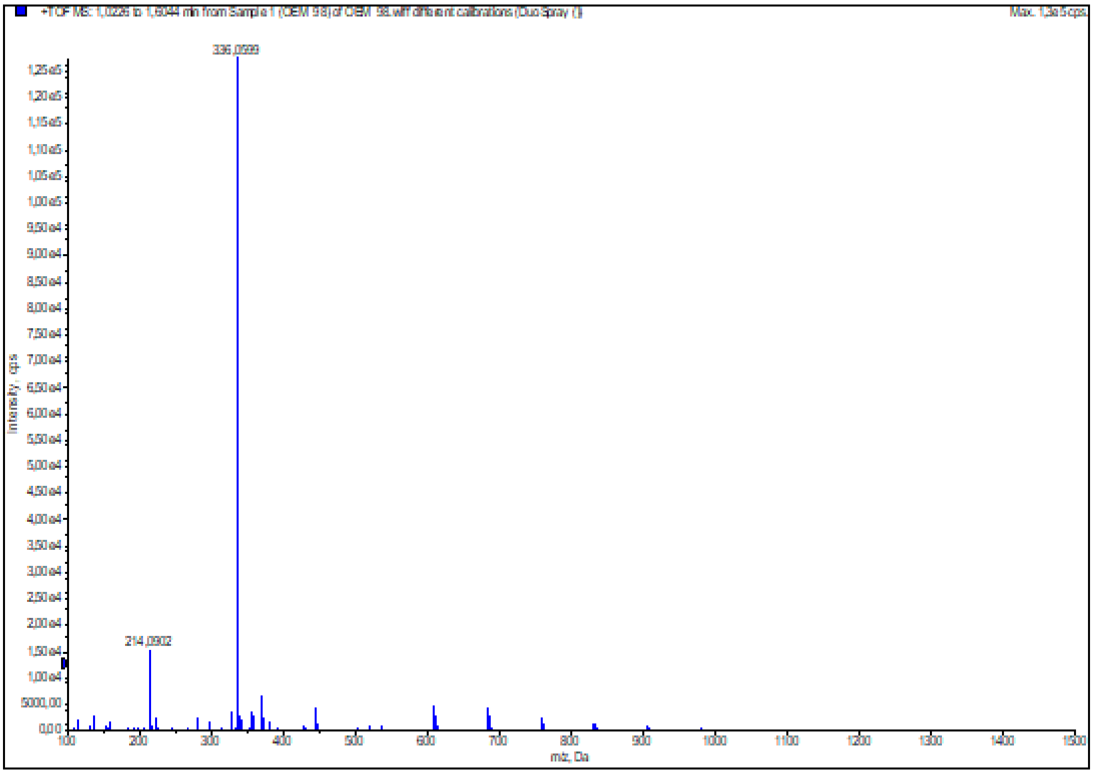

**Figure S41:** HRMS (ESI+) of compound **41**

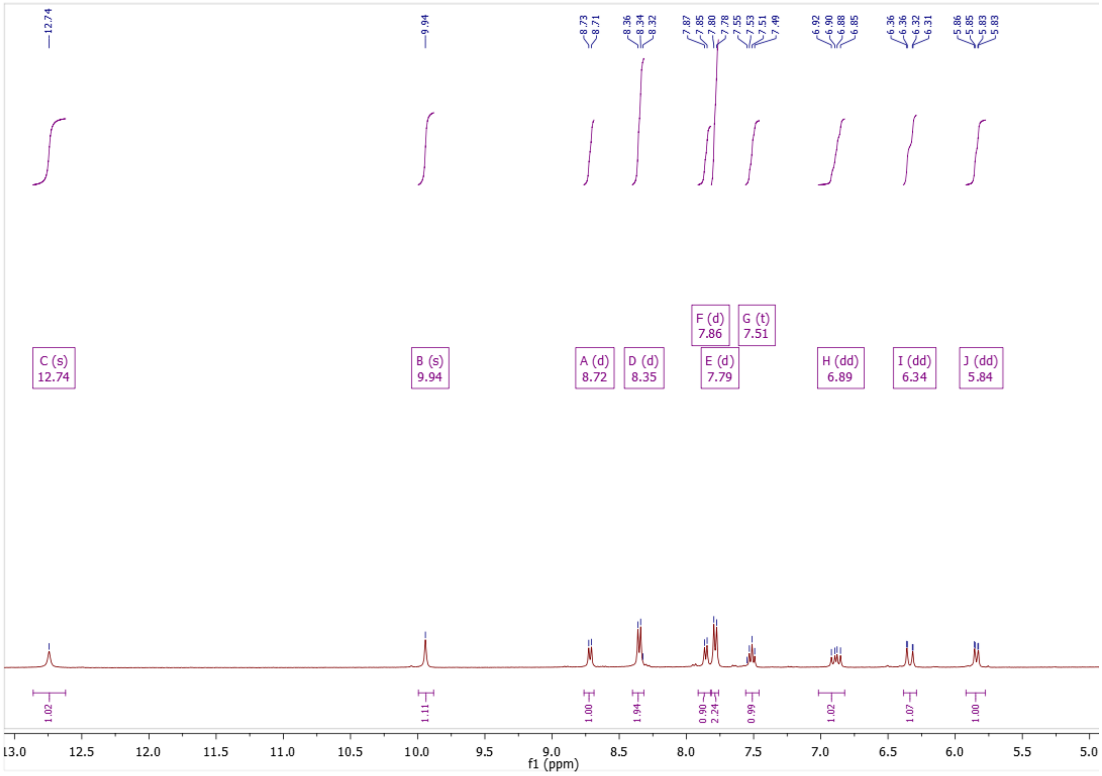

**Figure S42:** $^{1}$H NMR of compound **45**

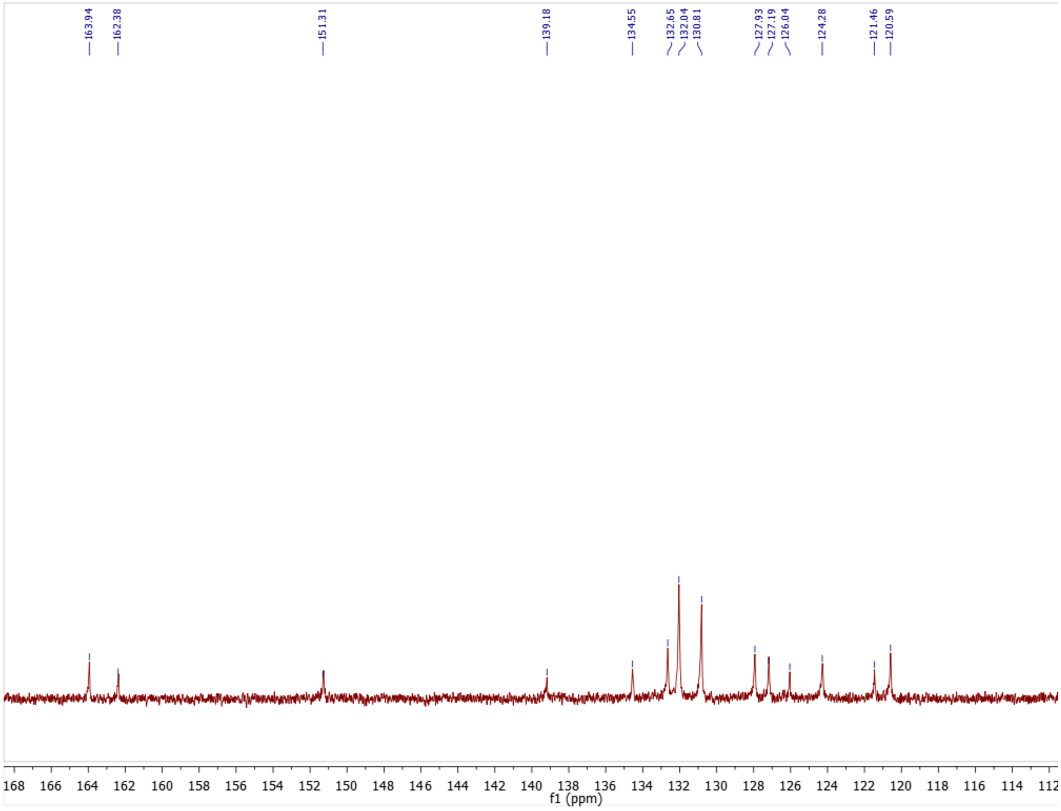

**Figure S43:** $^{13}$C NMR of compound **45**

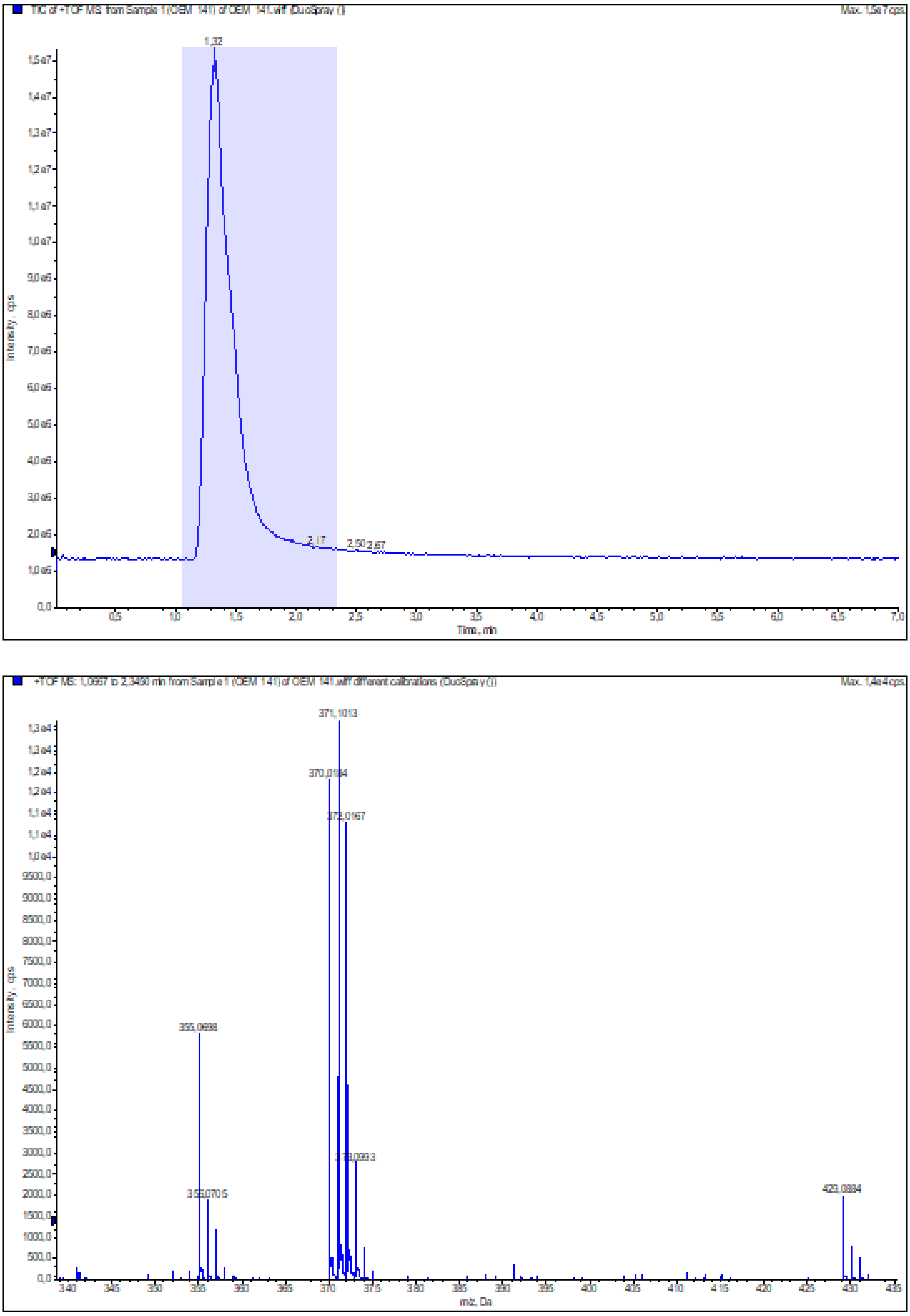

**Figure S44:** HRMS (ESI+) of compound **45**

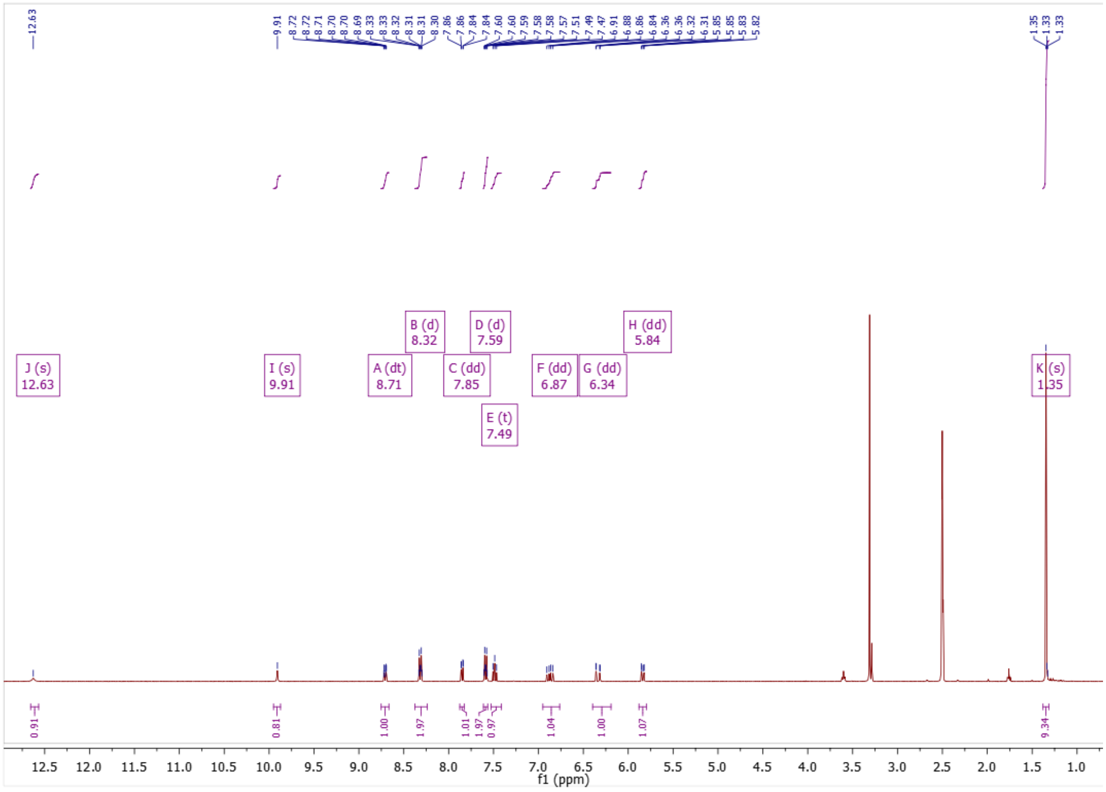

**Figure S45:** $^{1}$H NMR of compound **46**

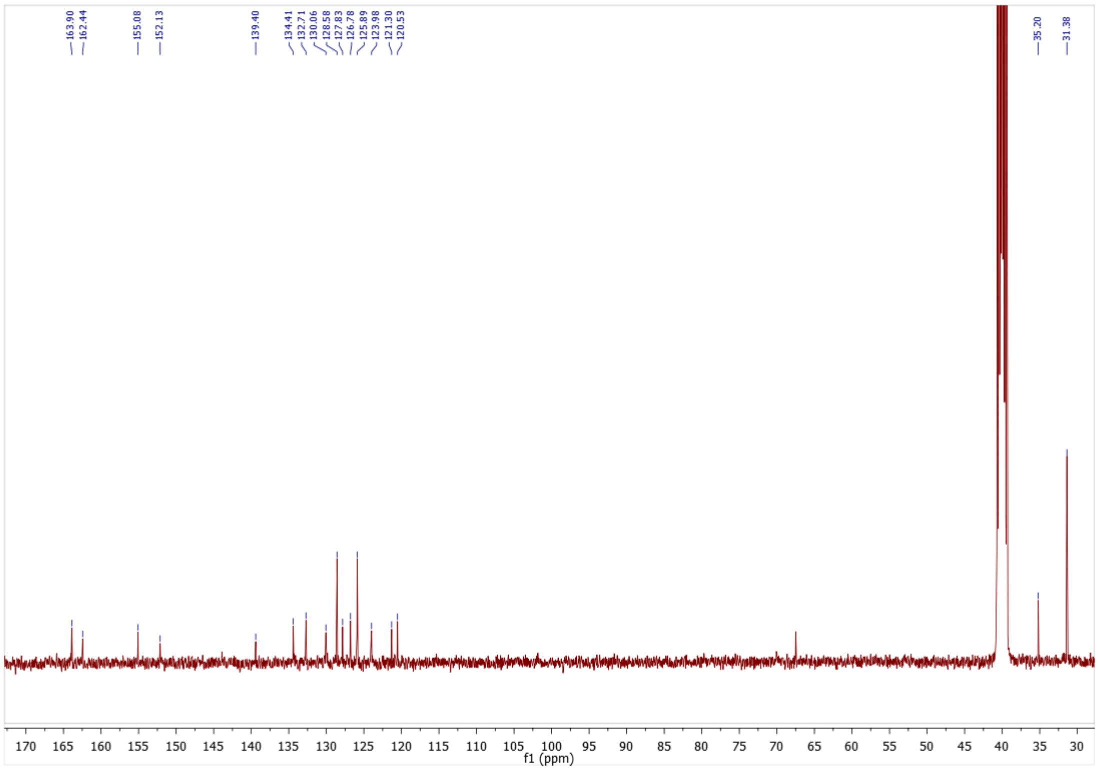

**Figure S46:** $^{13}$C NMR of compound **46**

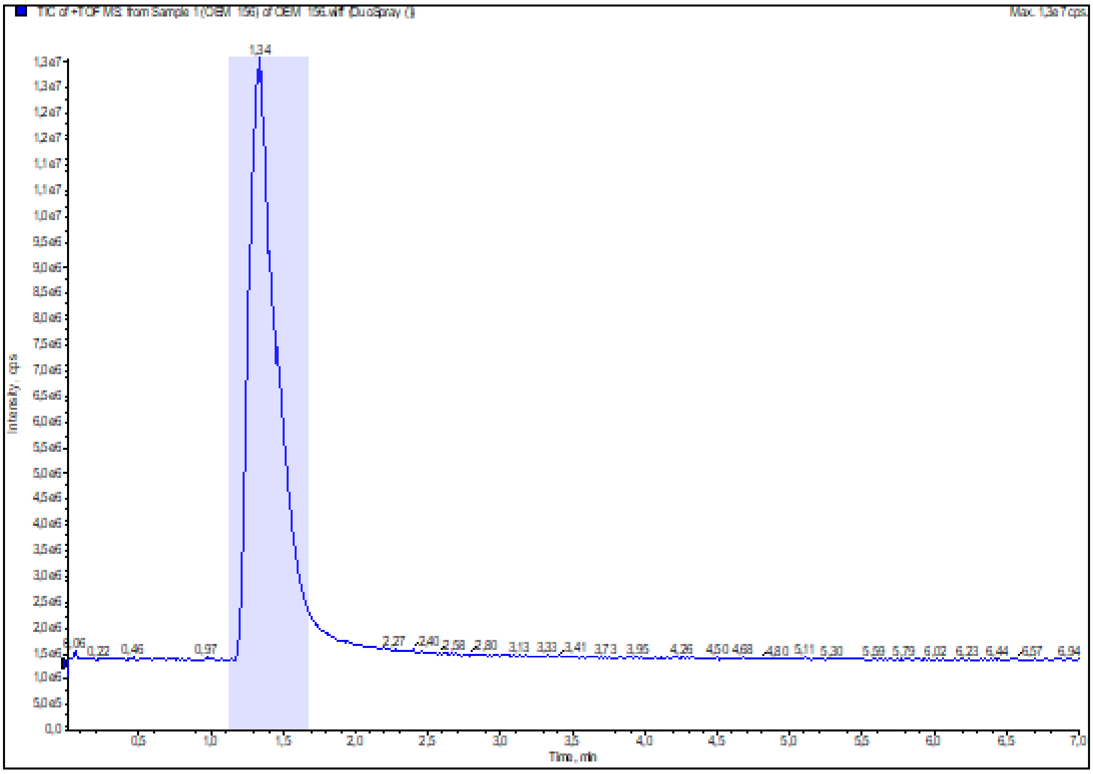

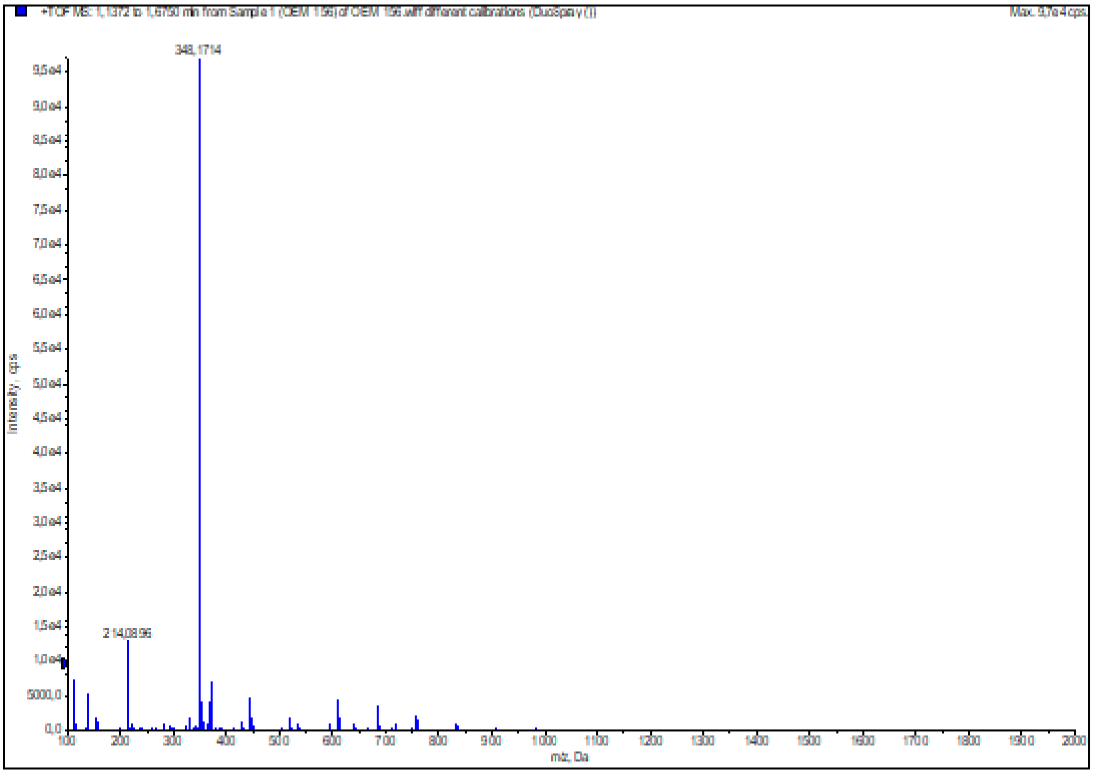

**Figure S47:** HRMS (ESI+) of compound **46**

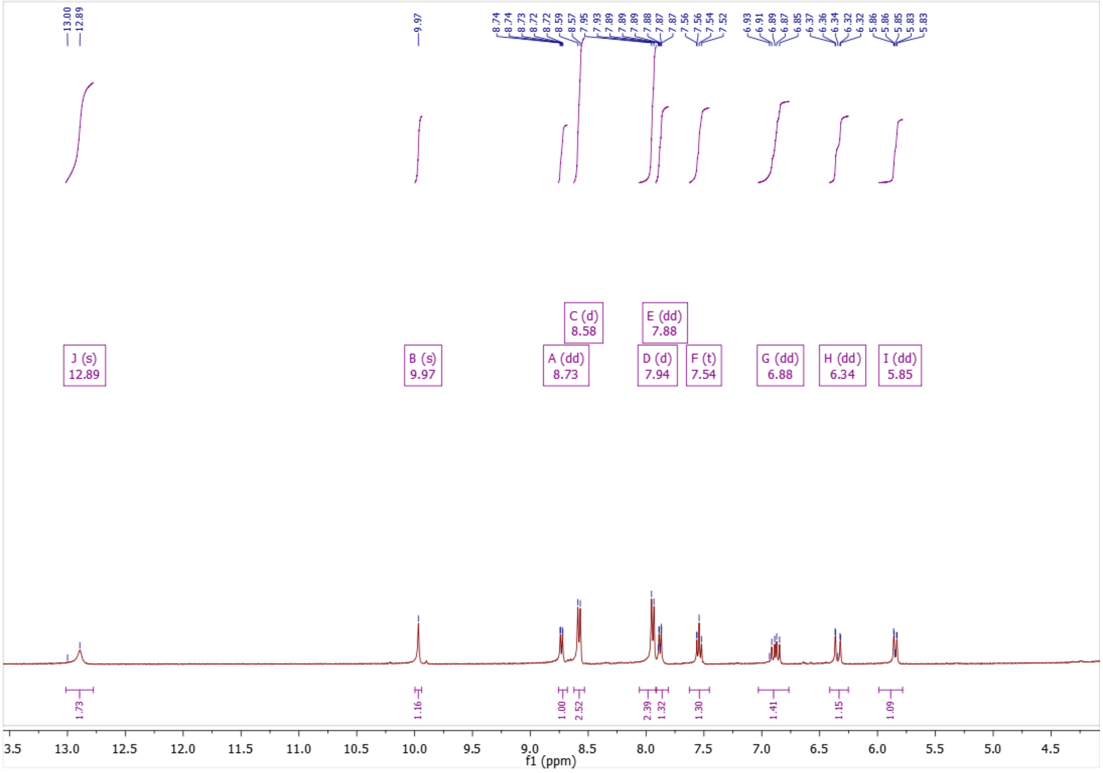

**Figure S48:** $^{1}$H NMR of compound **47**

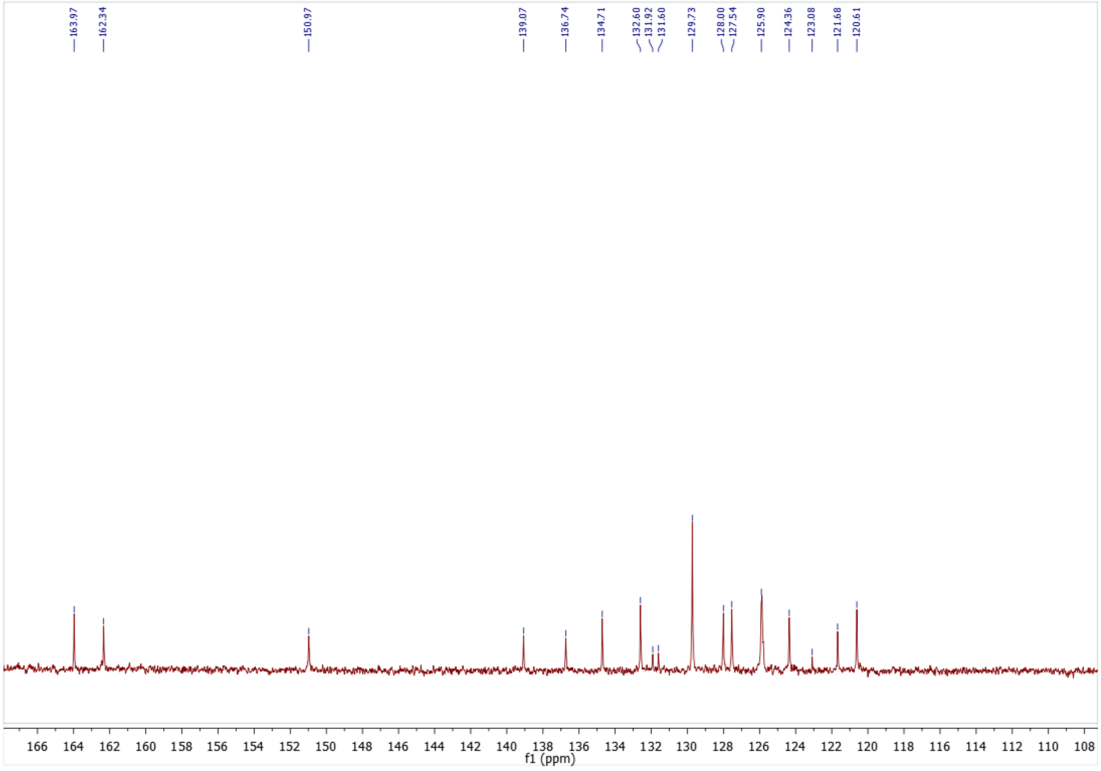

**Figure S49:** $^{13}$C NMR of compound **47**

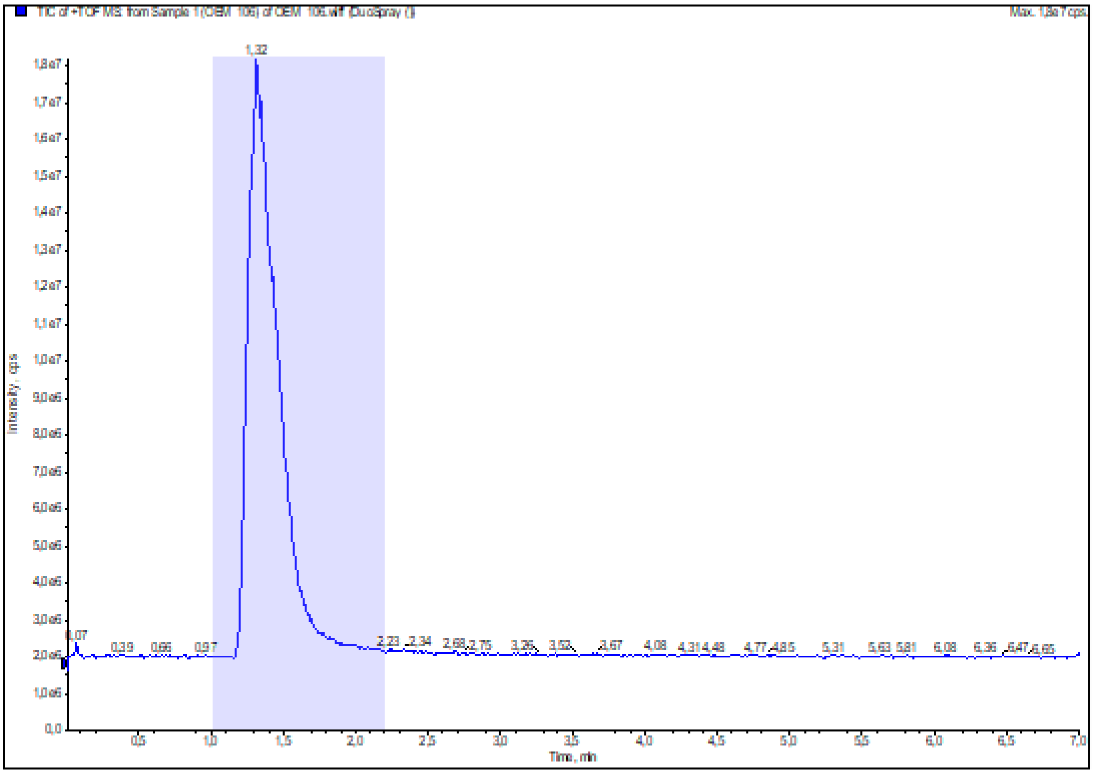

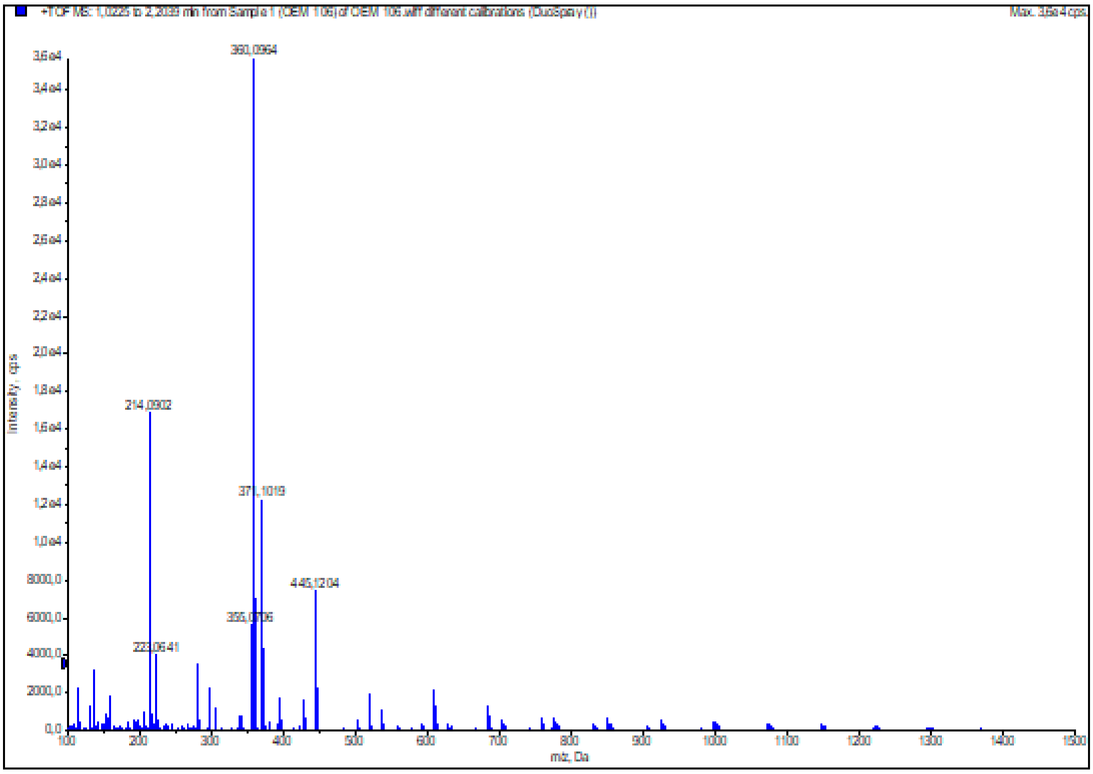

**Figure S50:** HRMS (ESI+) of compound **47**

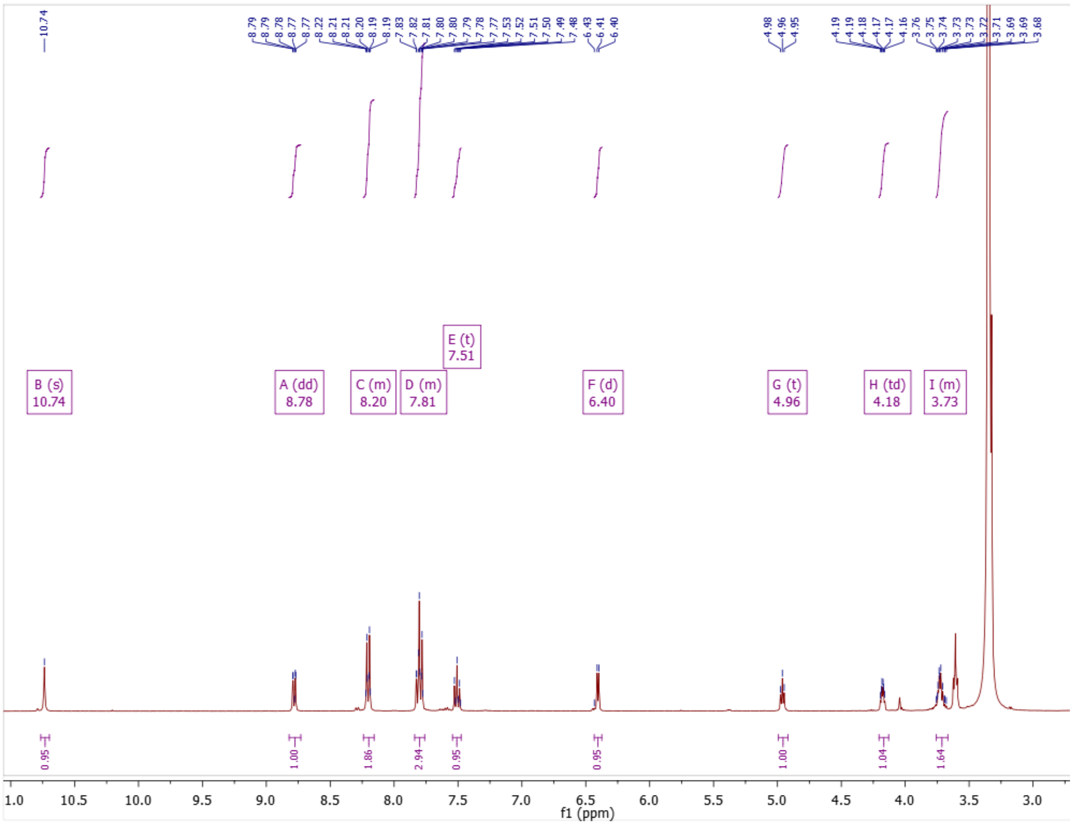

**Figure S51:** $^{1}$H NMR of compound **48**

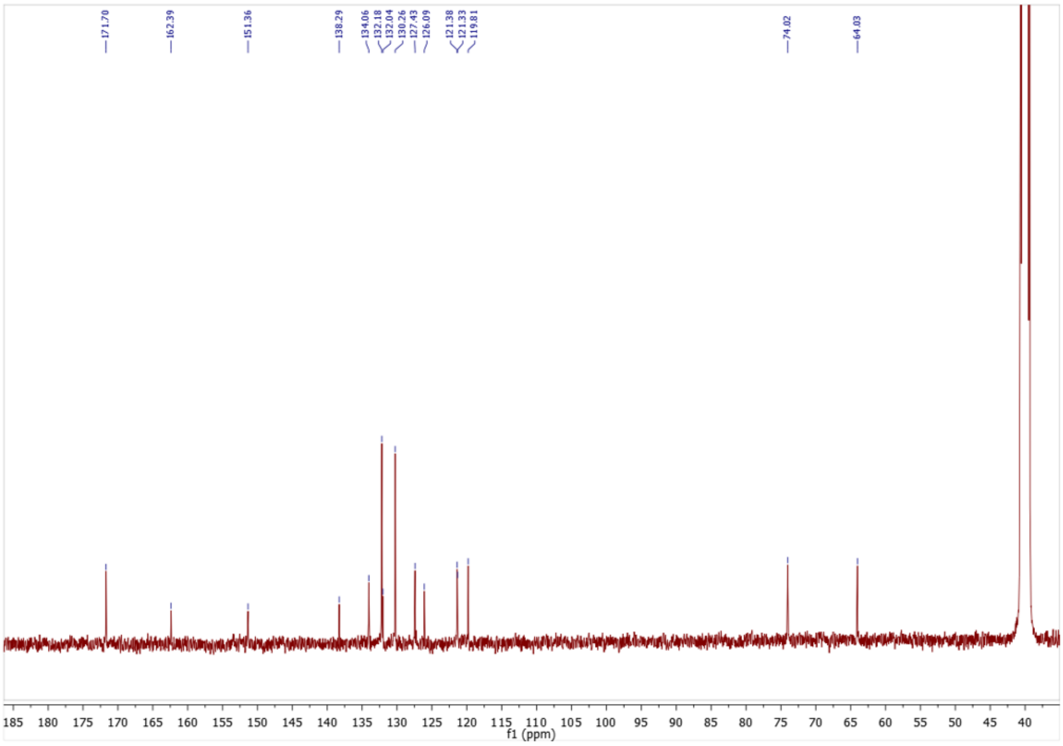

**Figure S52:** $^{13}$C NMR of compound **48**

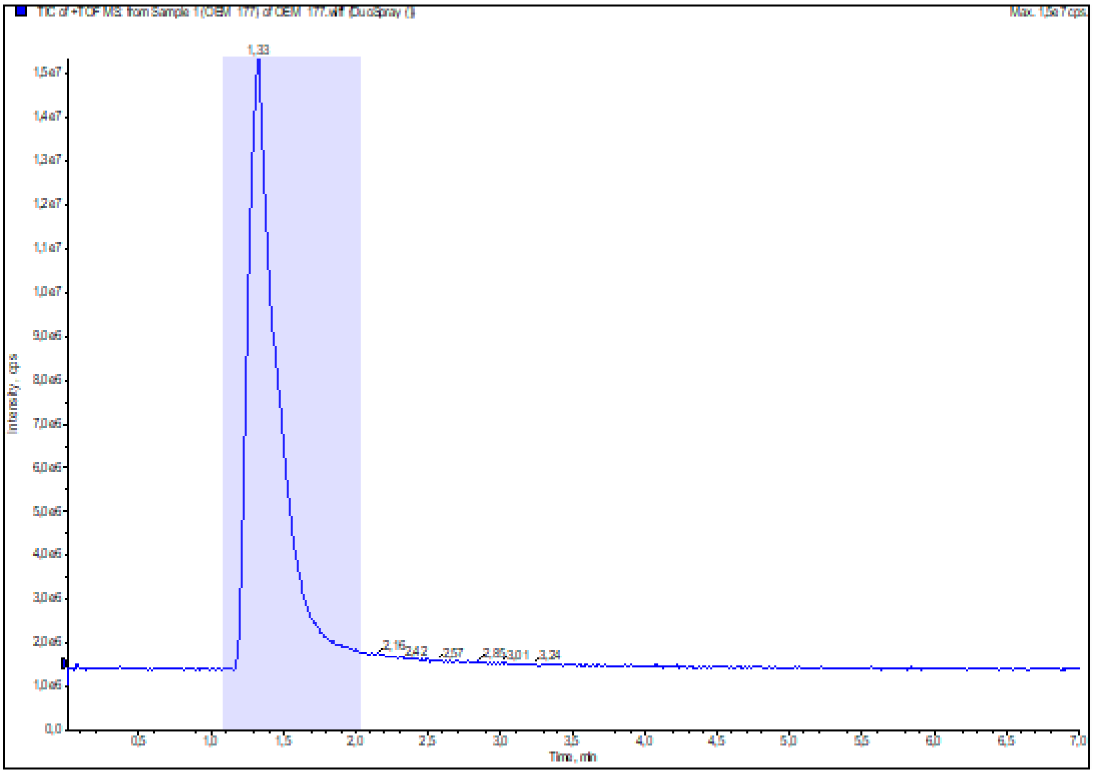

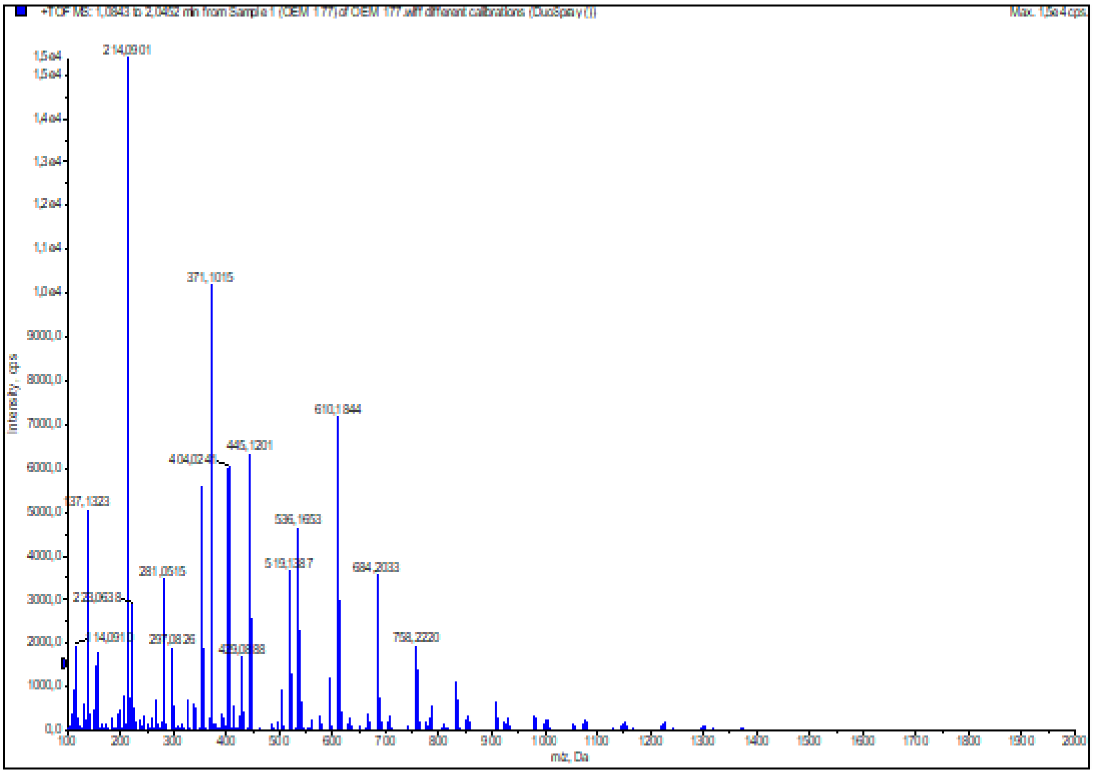

**Figure S53:** HRMS (ESI+) of compound **48**

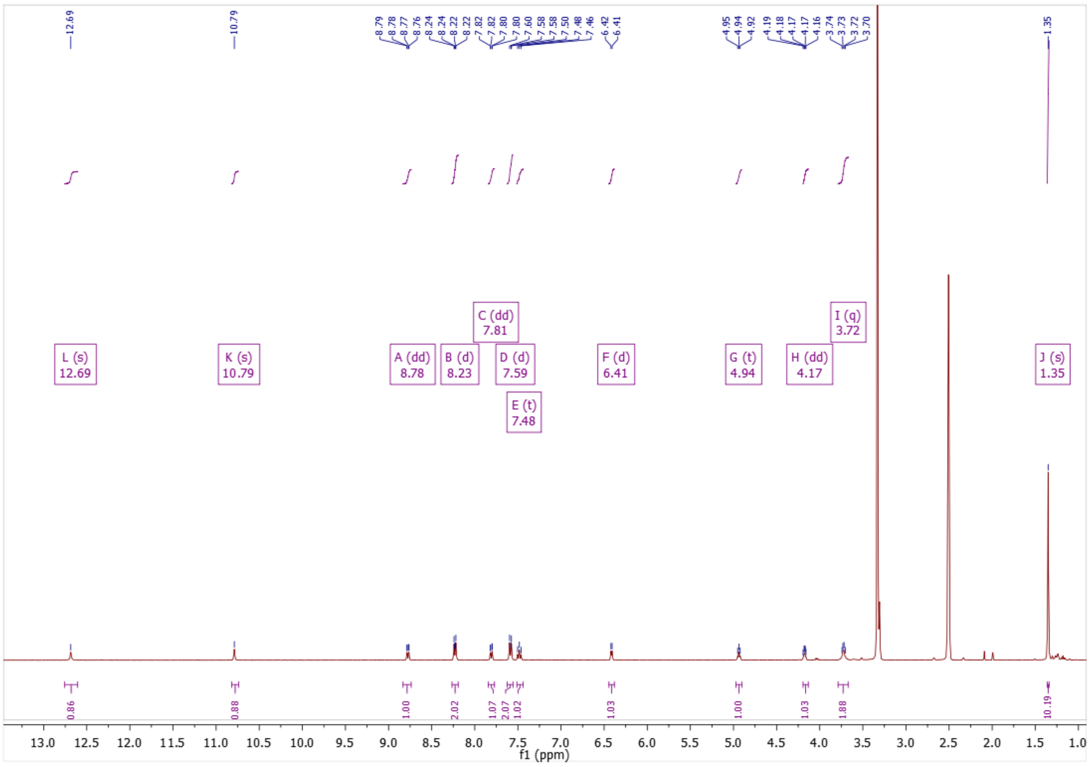

**Figure S54:** $^{1}$H NMR of compound **49**

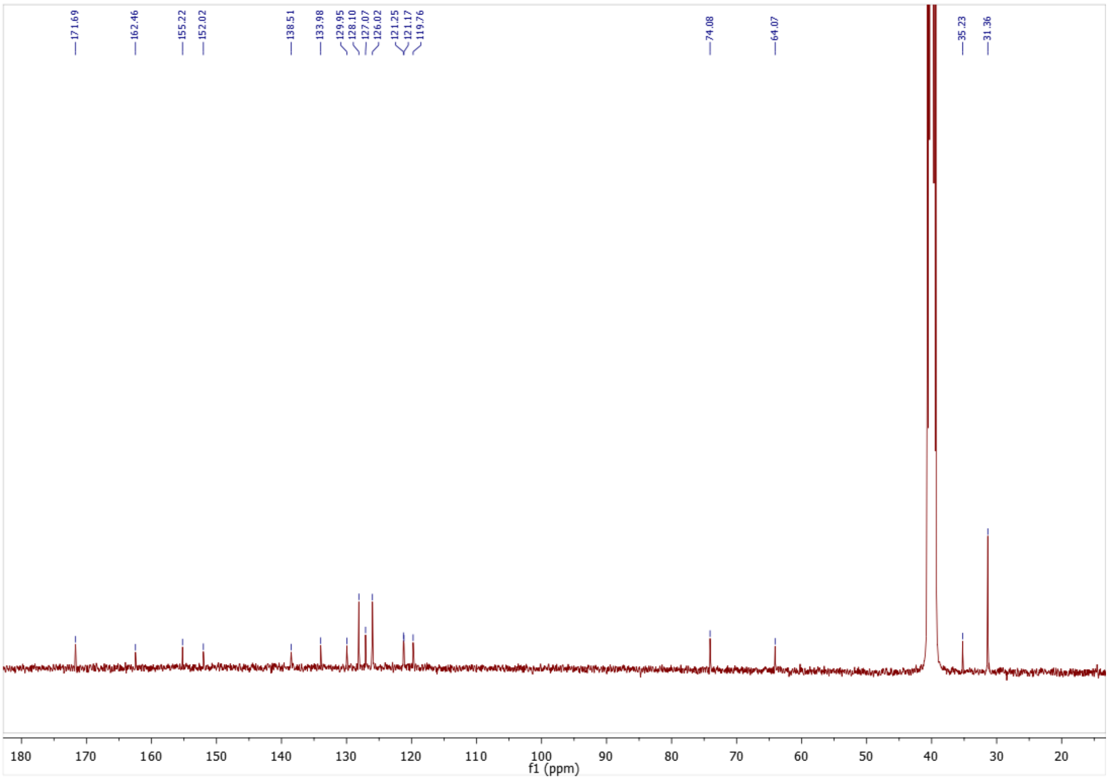

**Figure S55:** $^{13}$C NMR of compound **49**

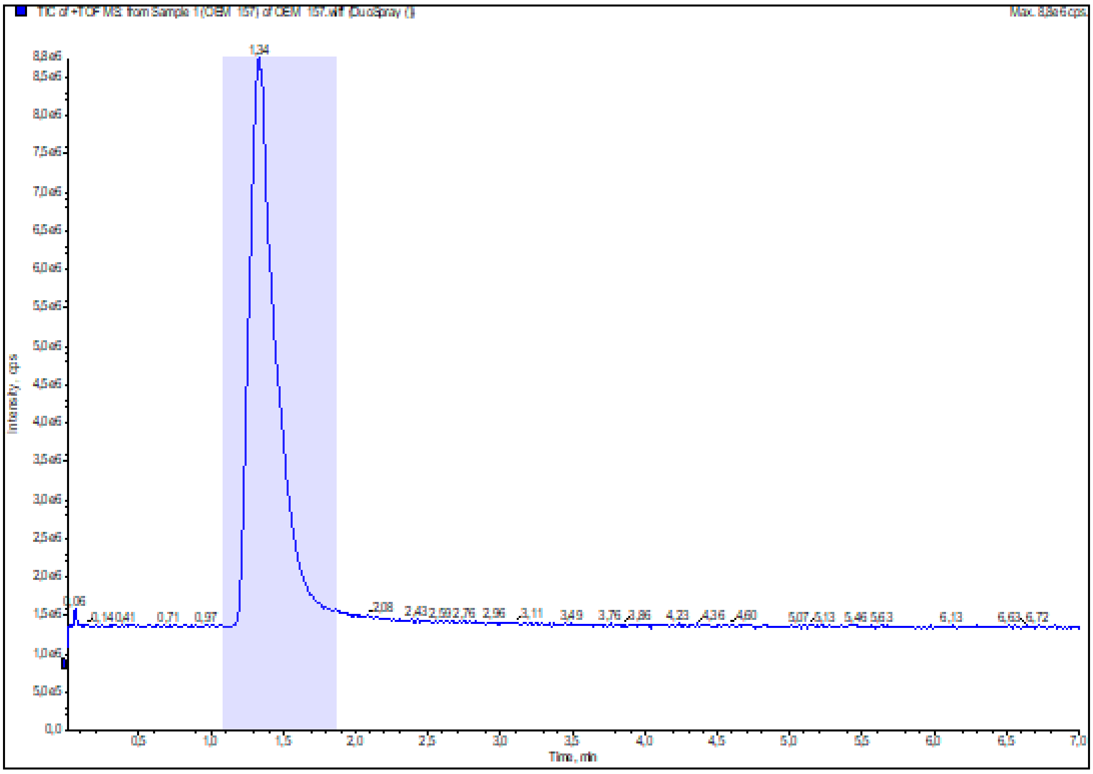

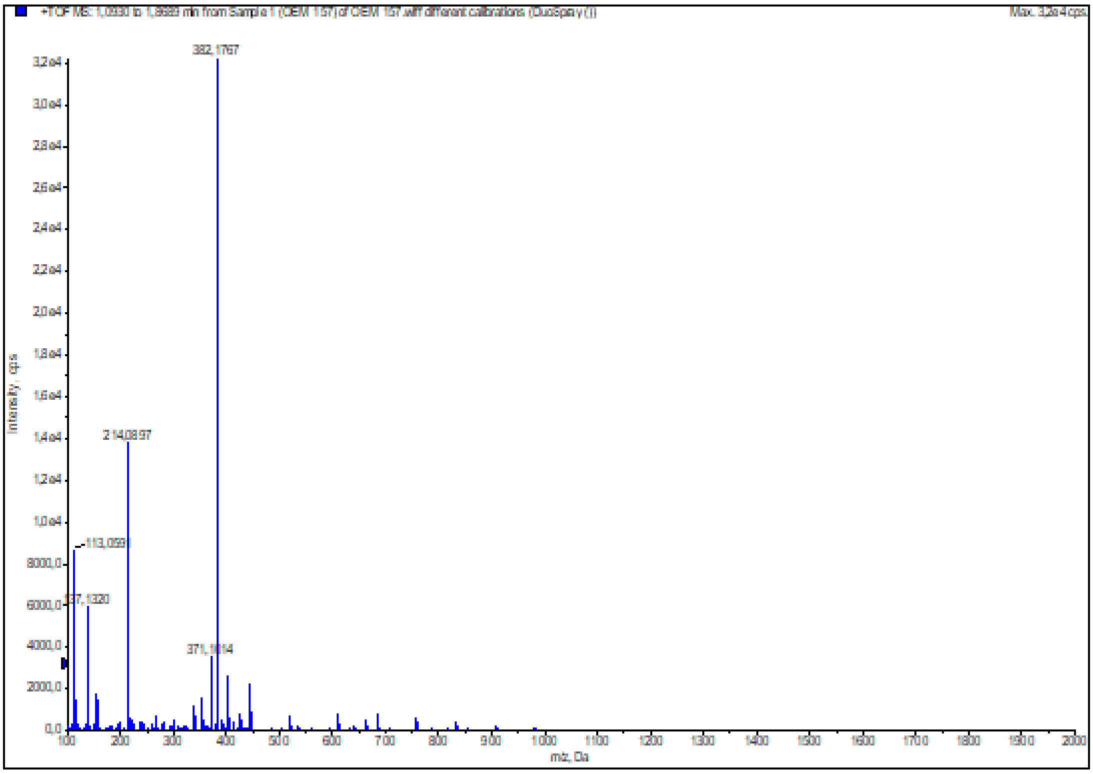

**Figure S56:** HRMS (ESI+) of compound **49**

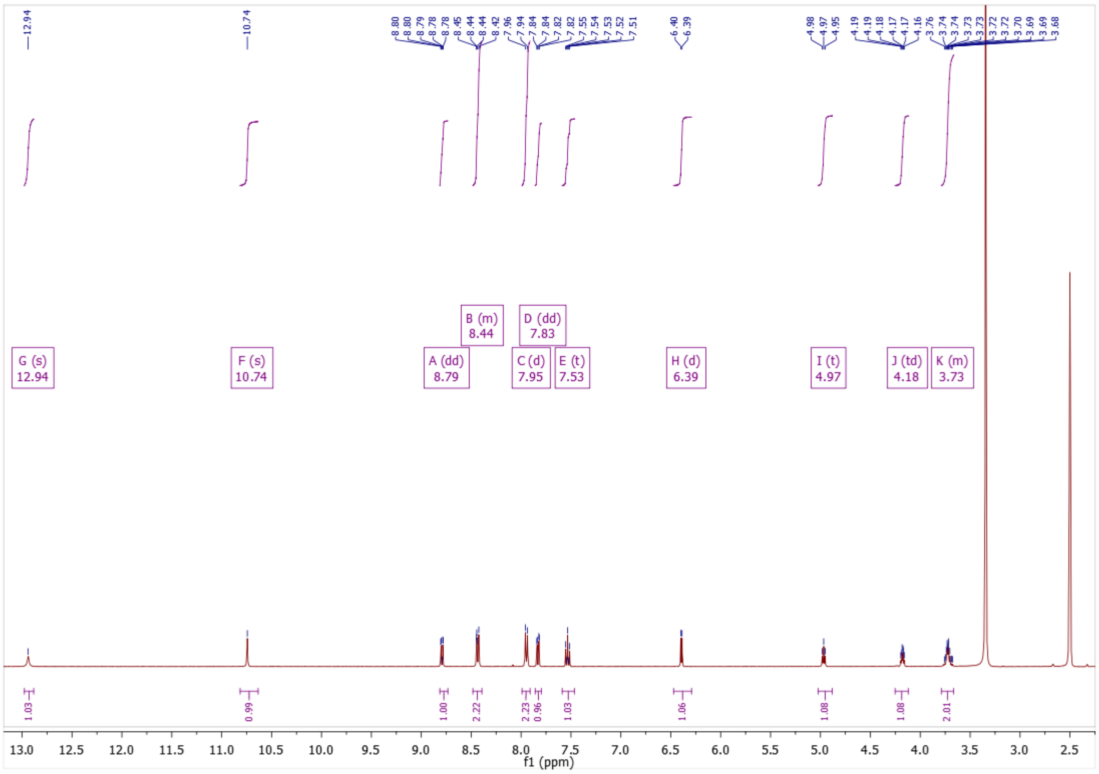

**Figure S57:** $^{1}$H NMR of compound **50**

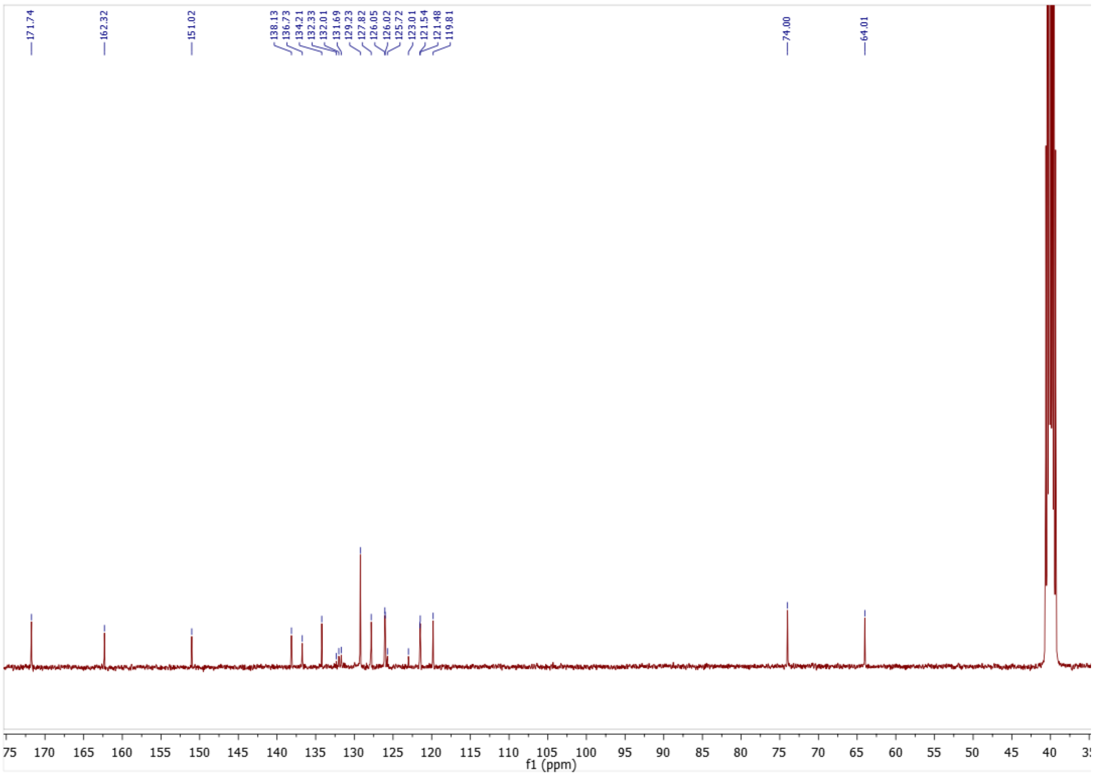

**Figure S58:** $^{13}$C NMR of compound **50**

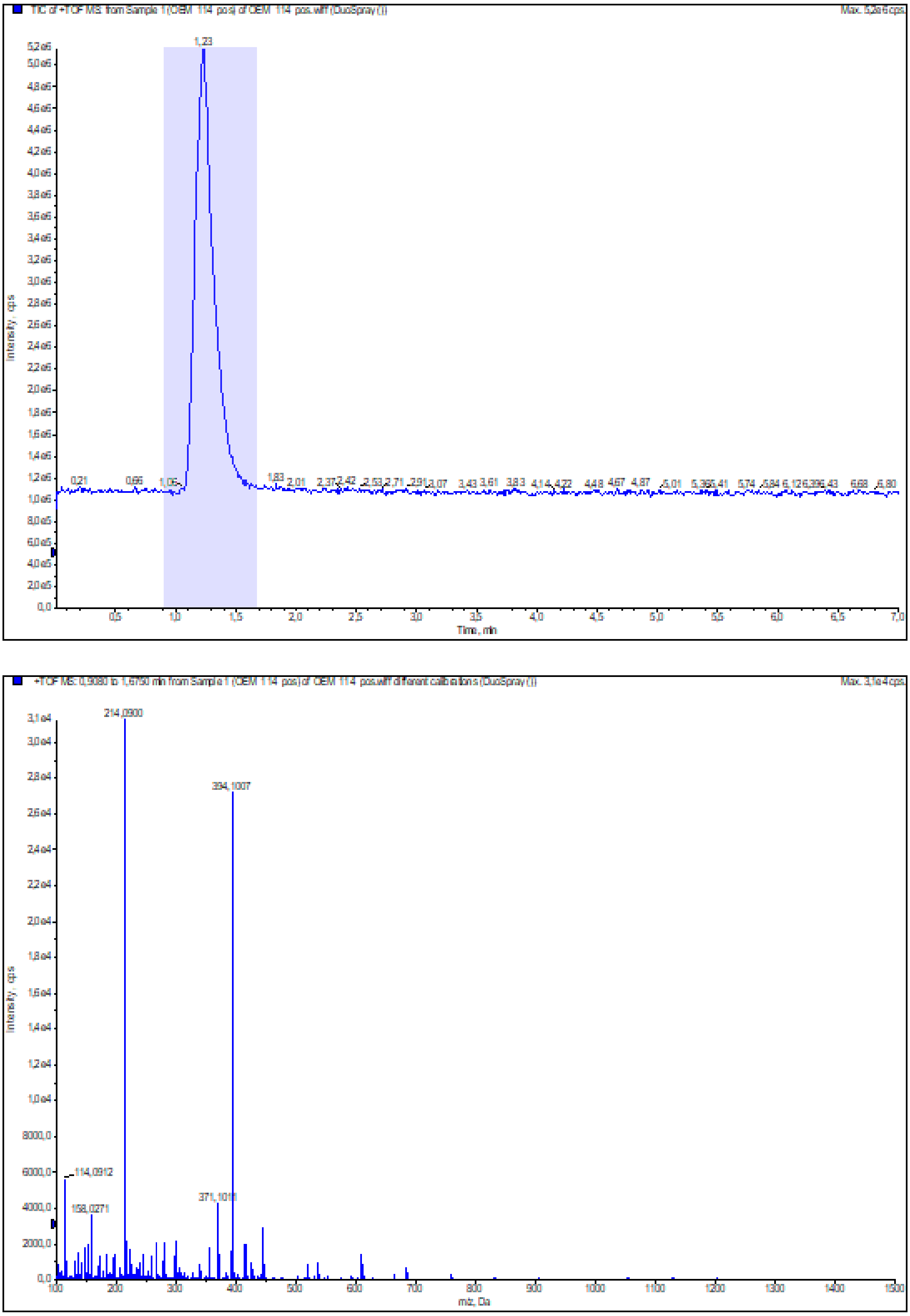

**Figure S59:** HRMS (ESI+) of compound **50**

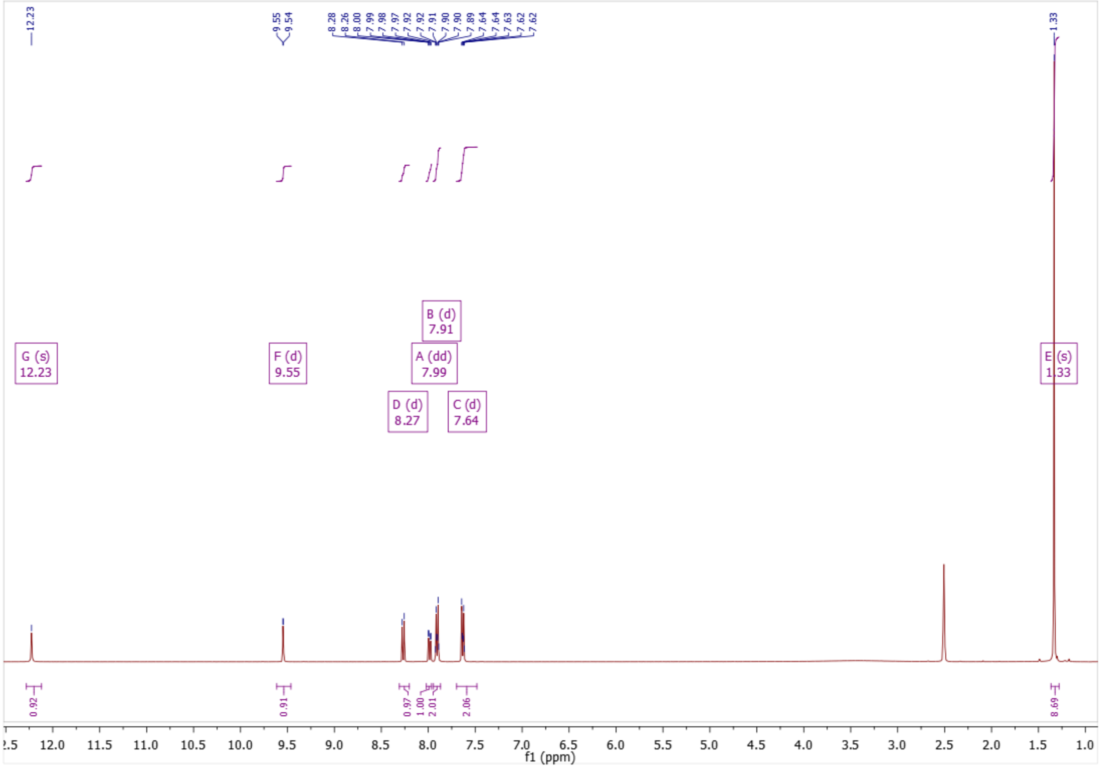

**Figure S60:** $^{1}$H NMR of compound **55**

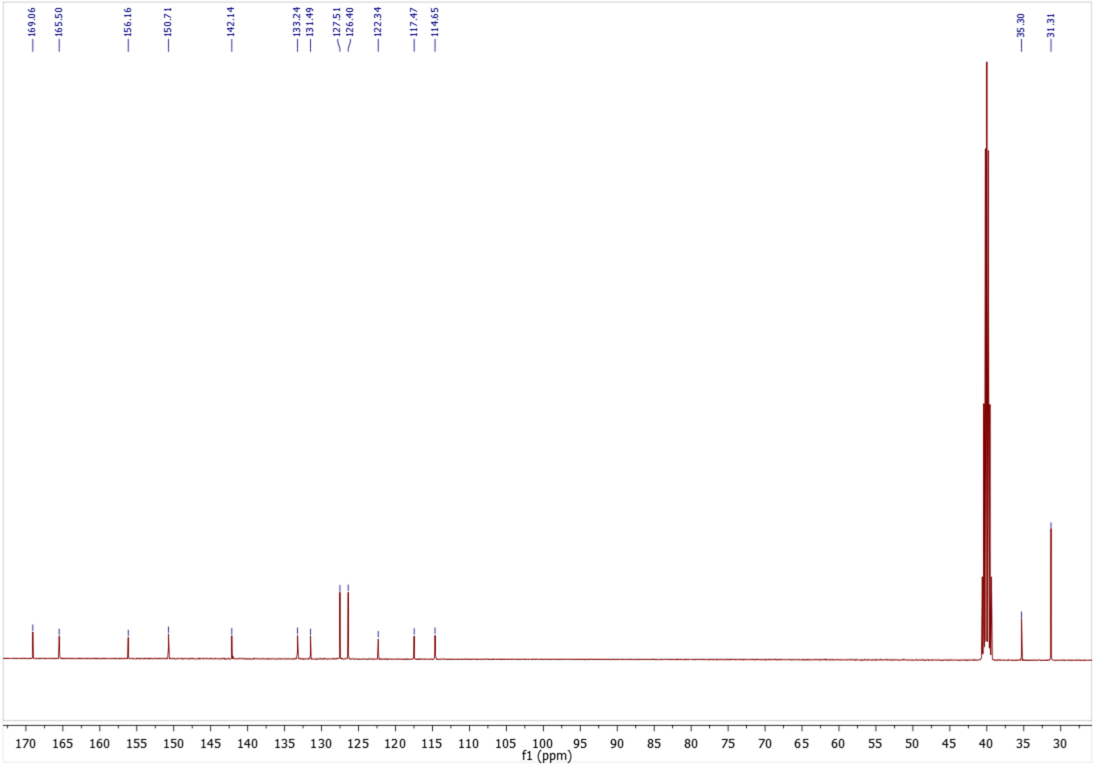

**Figure S61:** $^{13}$C NMR of compound **55**

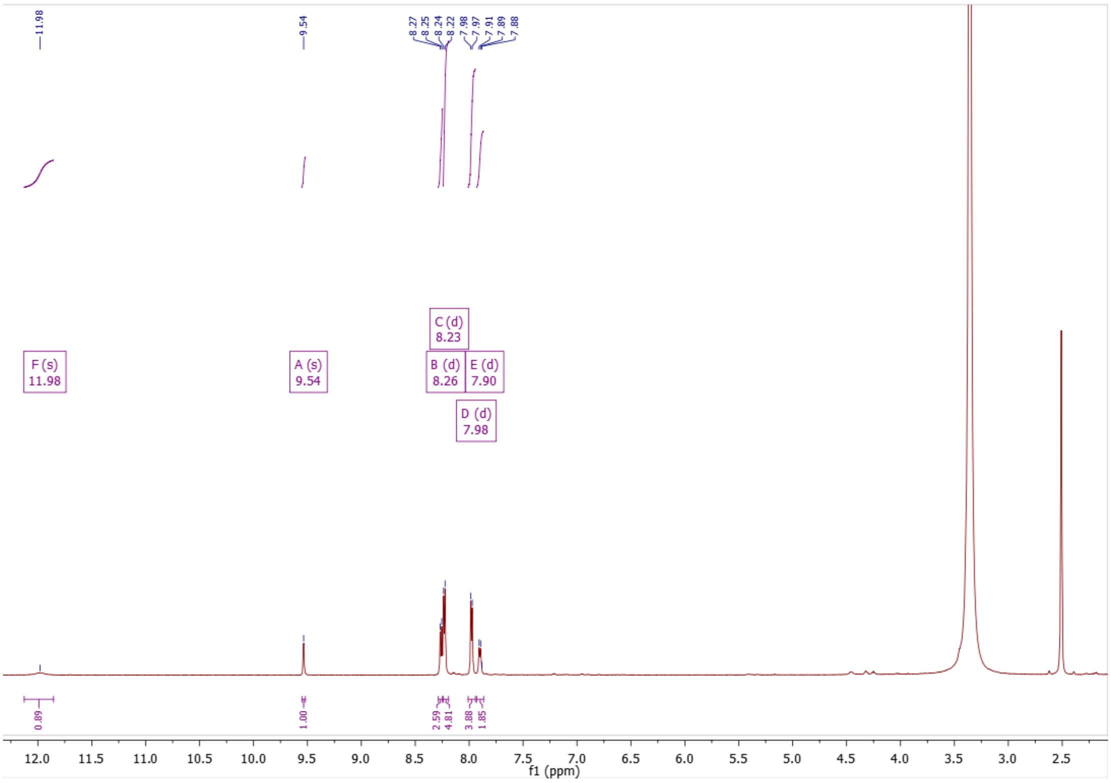

**Figure S62:** $^{1}H$ NMR of compound **56**

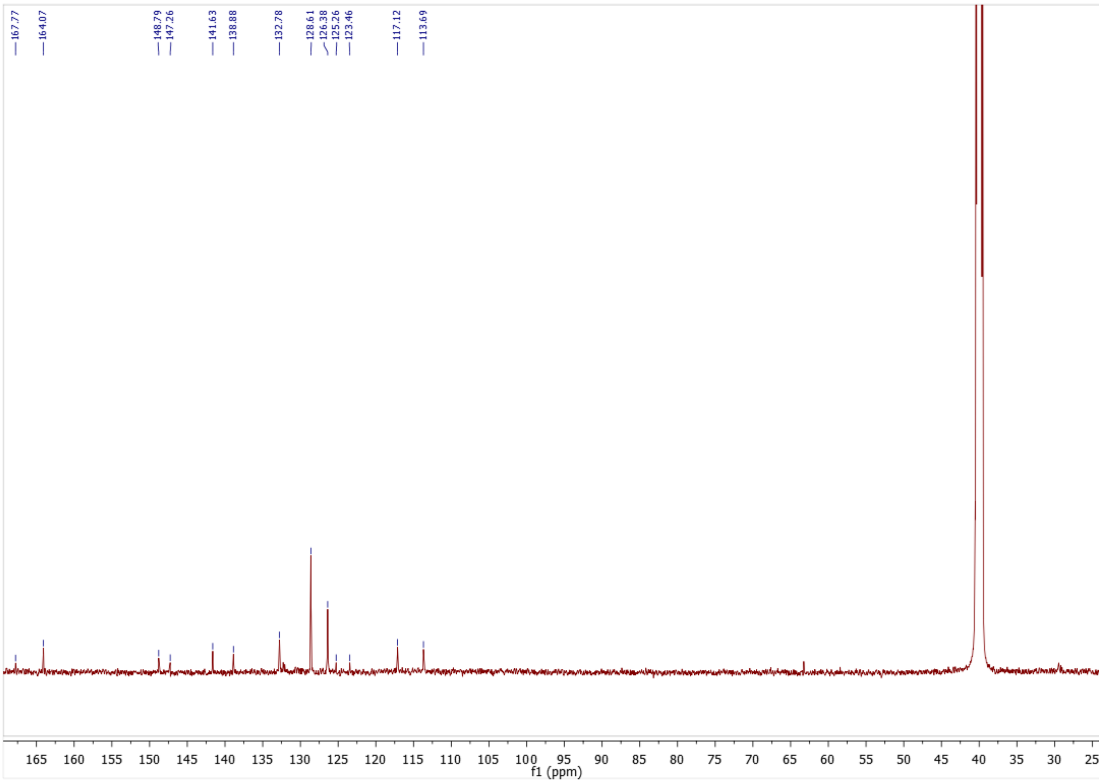

**Figure S63:** $^{13}C$ NMR of compound **56**

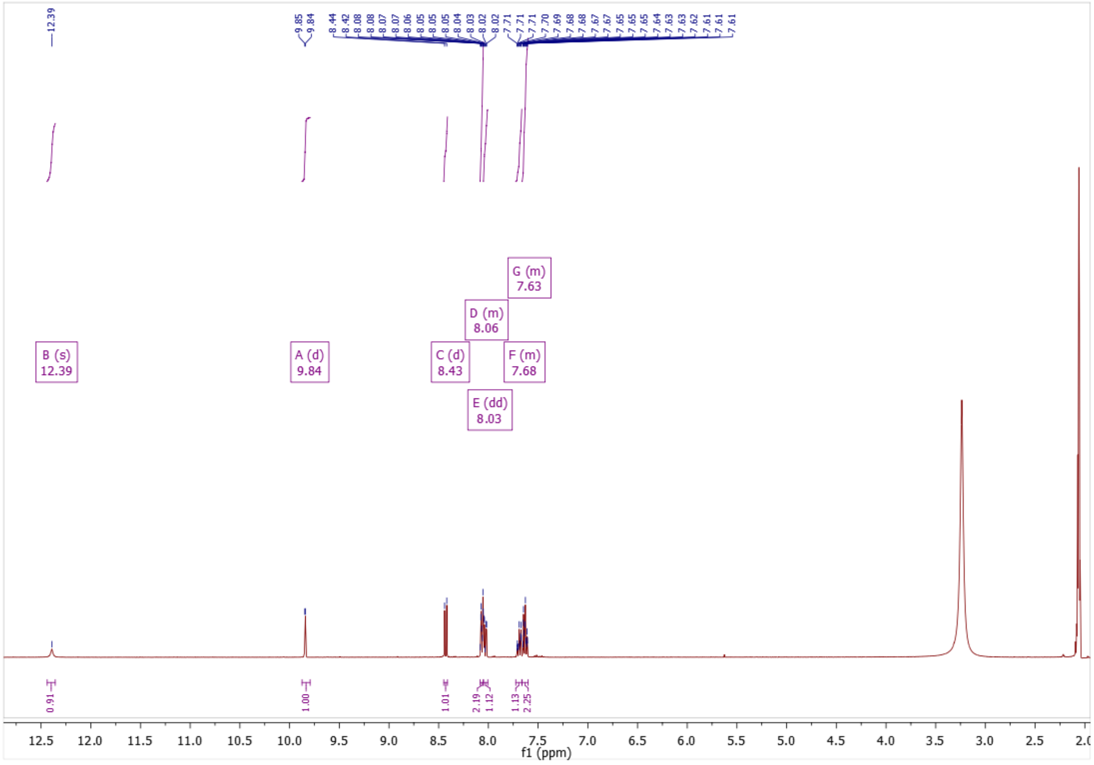

**Figure S64:** $^{1}$H NMR of compound **57**

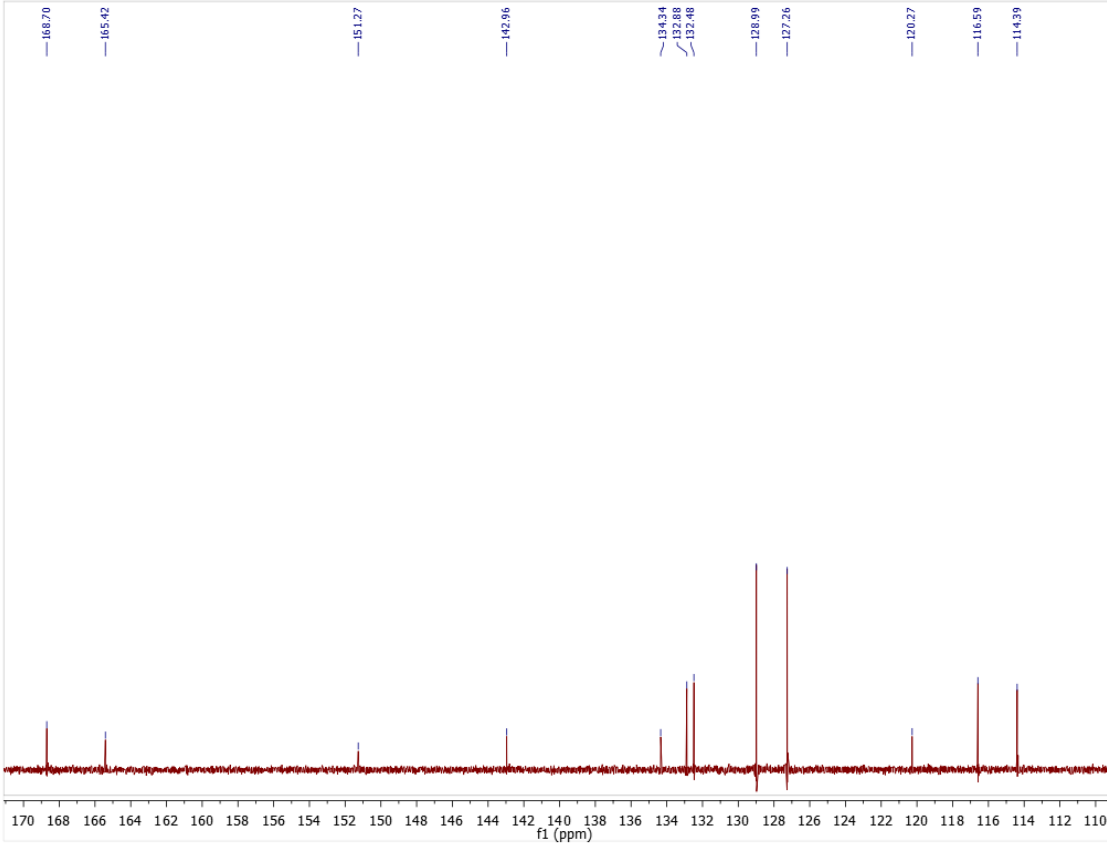

**Figure S65:** $^{13}$C NMR of compound **57**

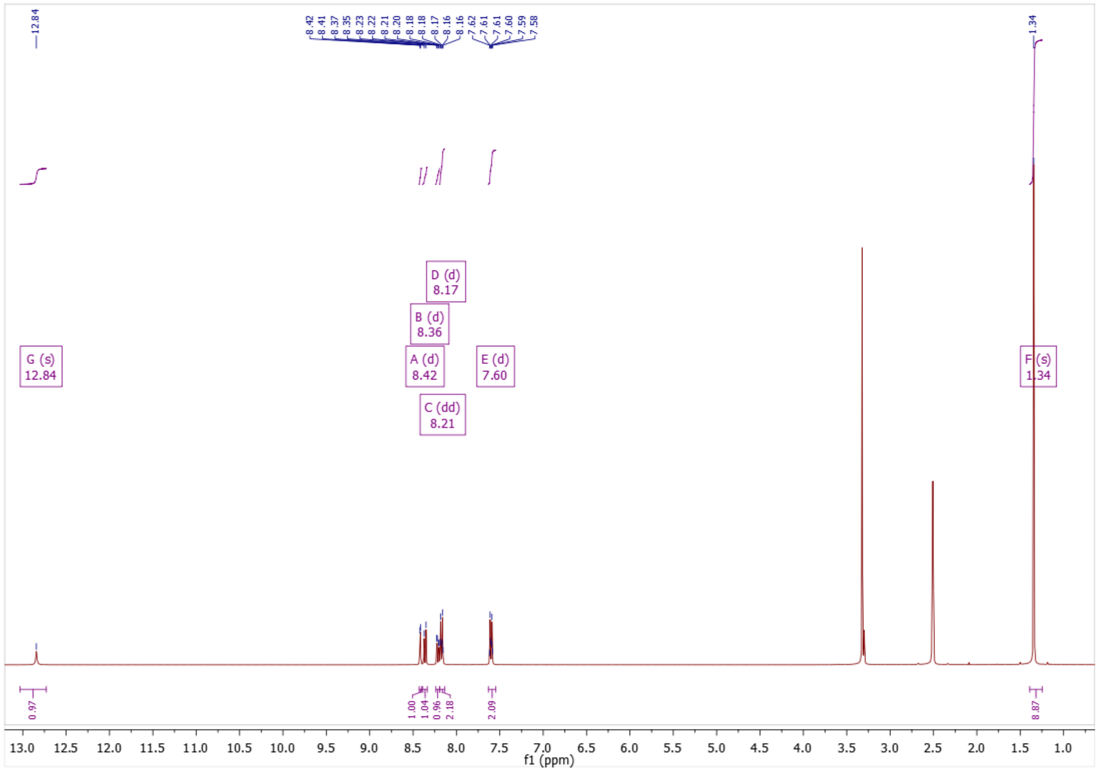

Figure S66: [1]H NMR of compound **64**

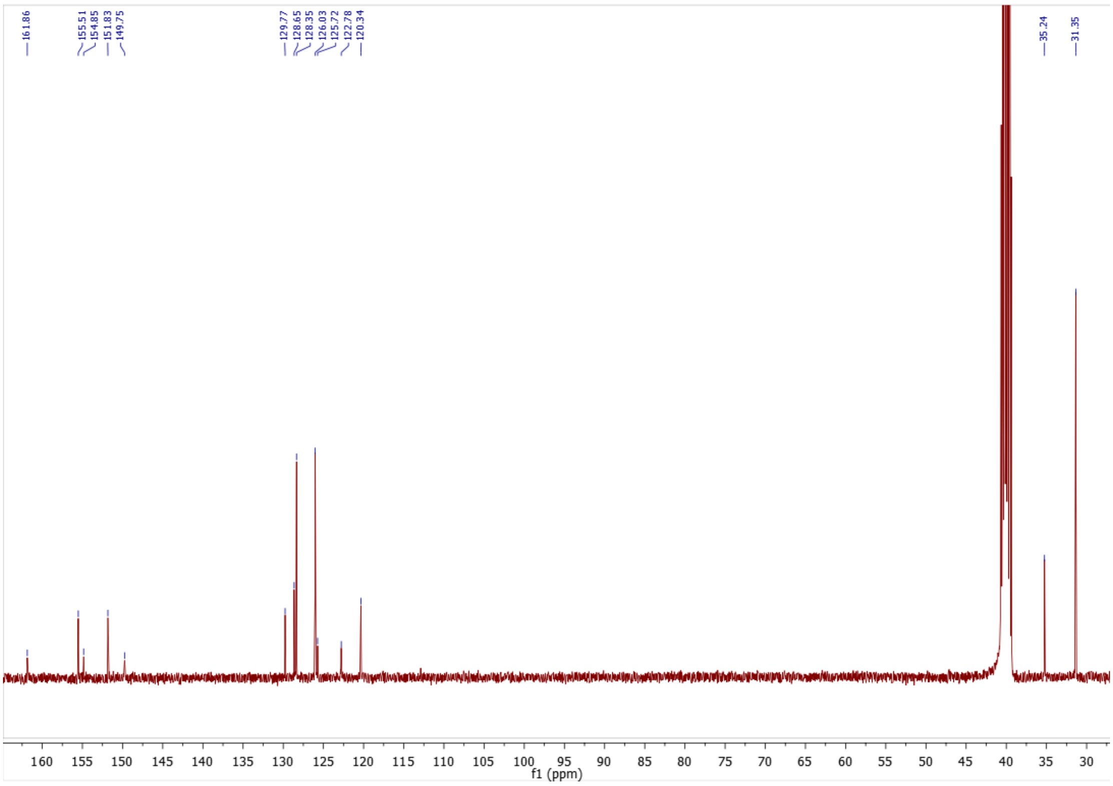

Figure S67: [13]C NMR of compound **64**

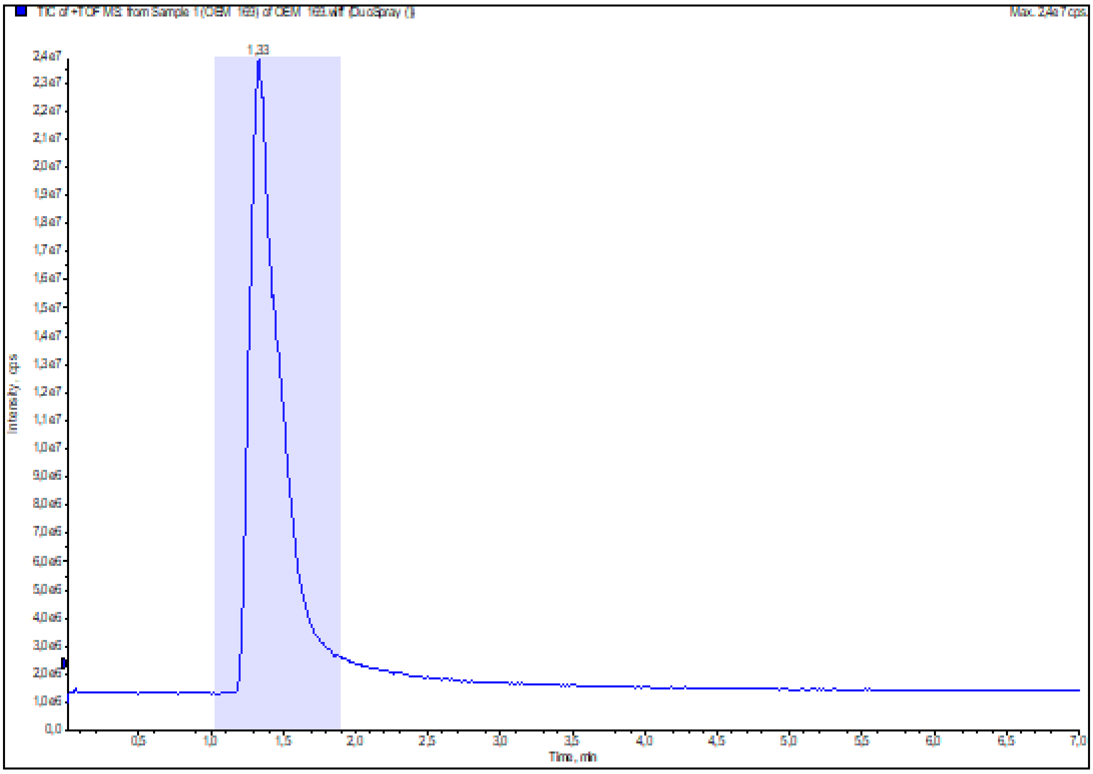

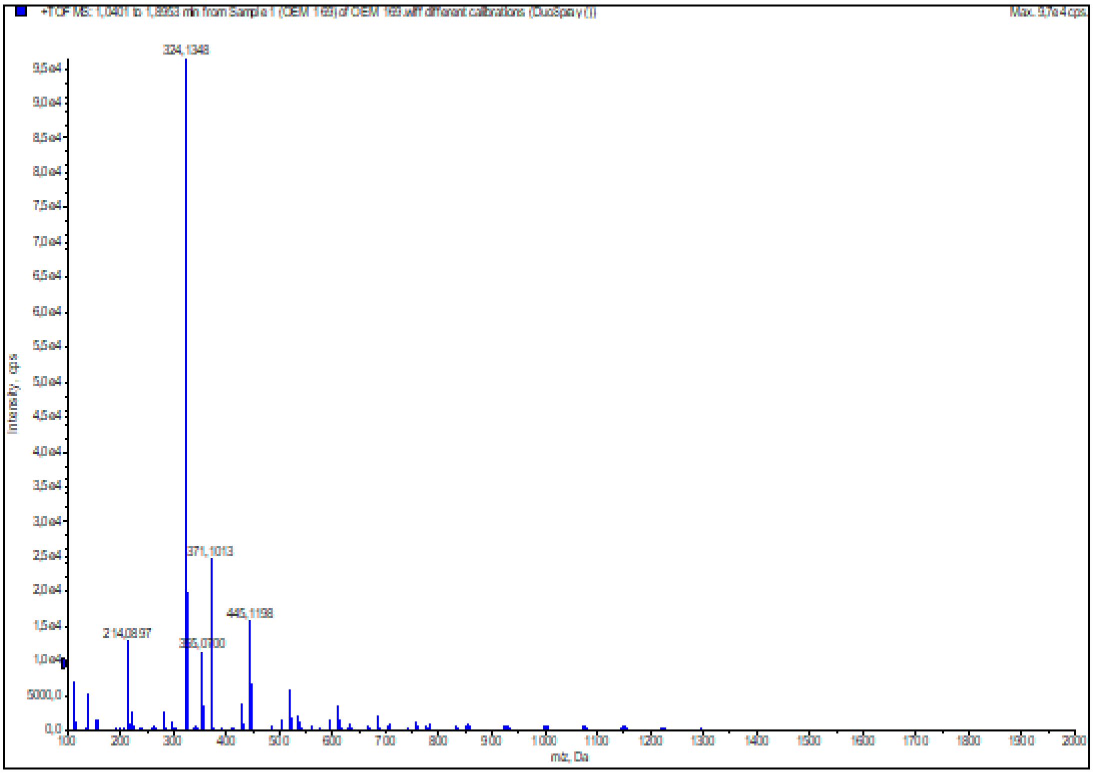

**Figure S68:** HRMS (ESI+) of compound **64**

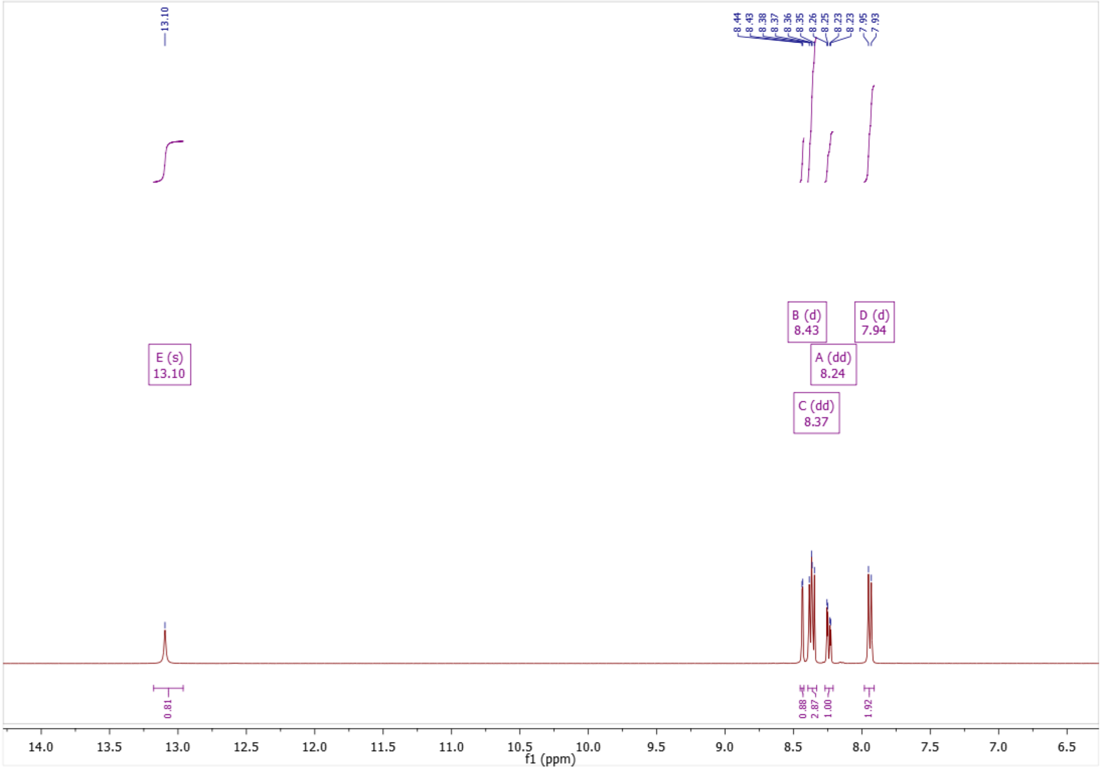

**Figure S69:** $^{1}$H NMR of compound **65**

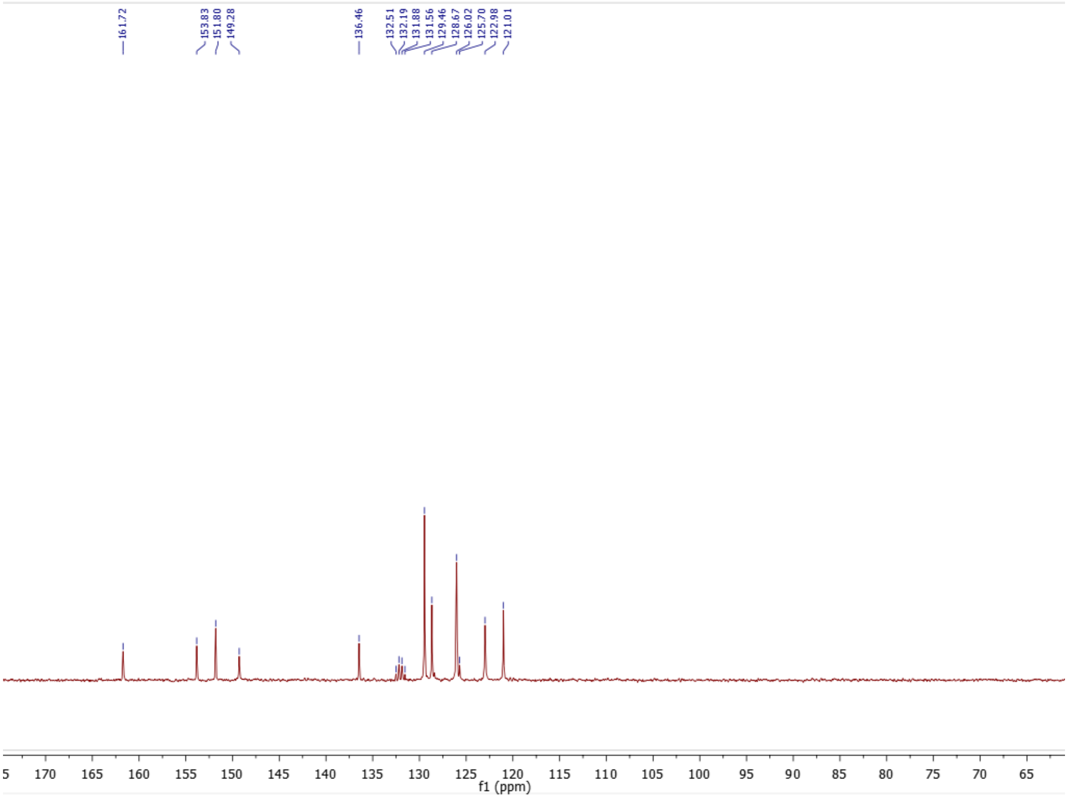

**Figure S70:** $^{13}$C NMR of compound **65**

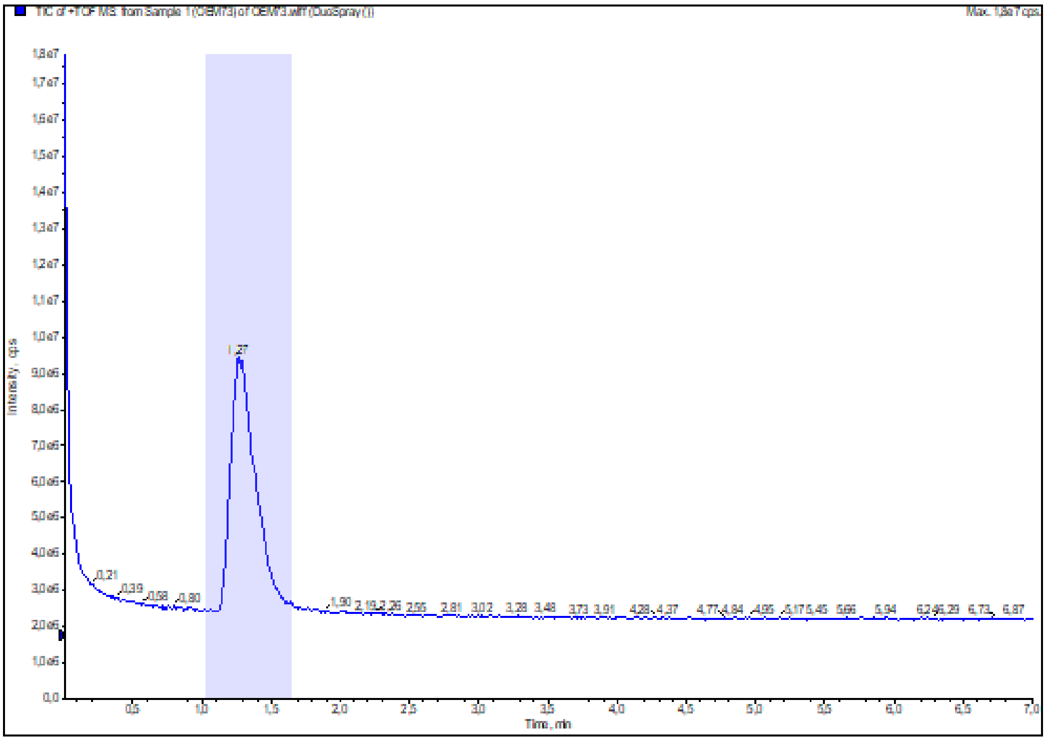

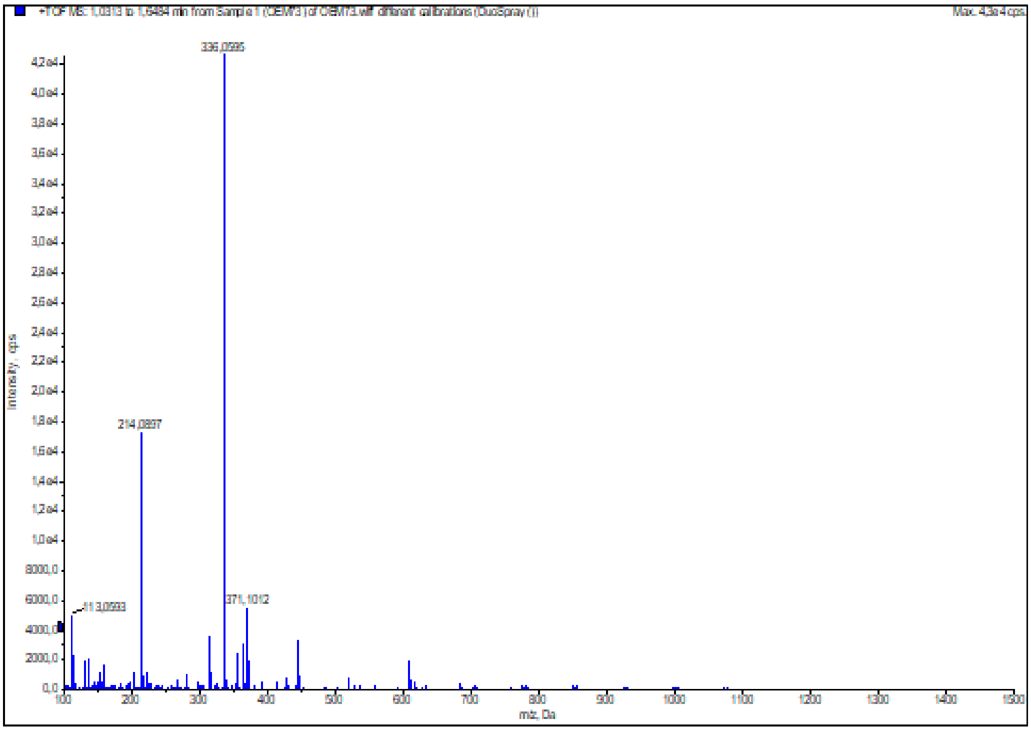

**Figure S71:** HRMS (ESI+) of compound **65**

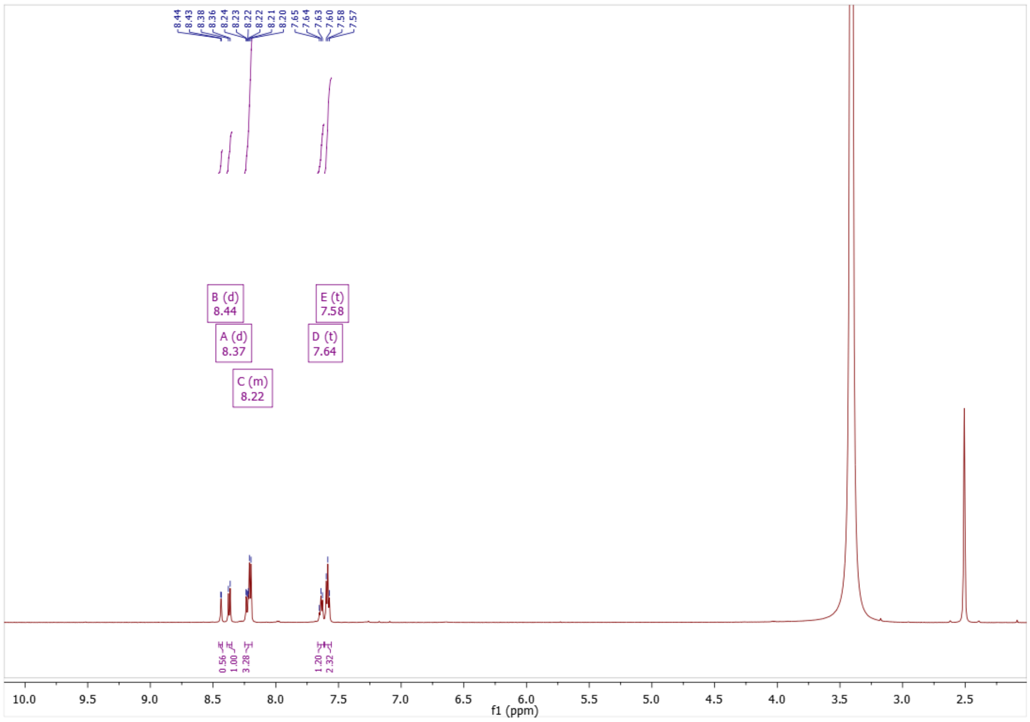

**Figure S72:** [1]H NMR of compound **66**

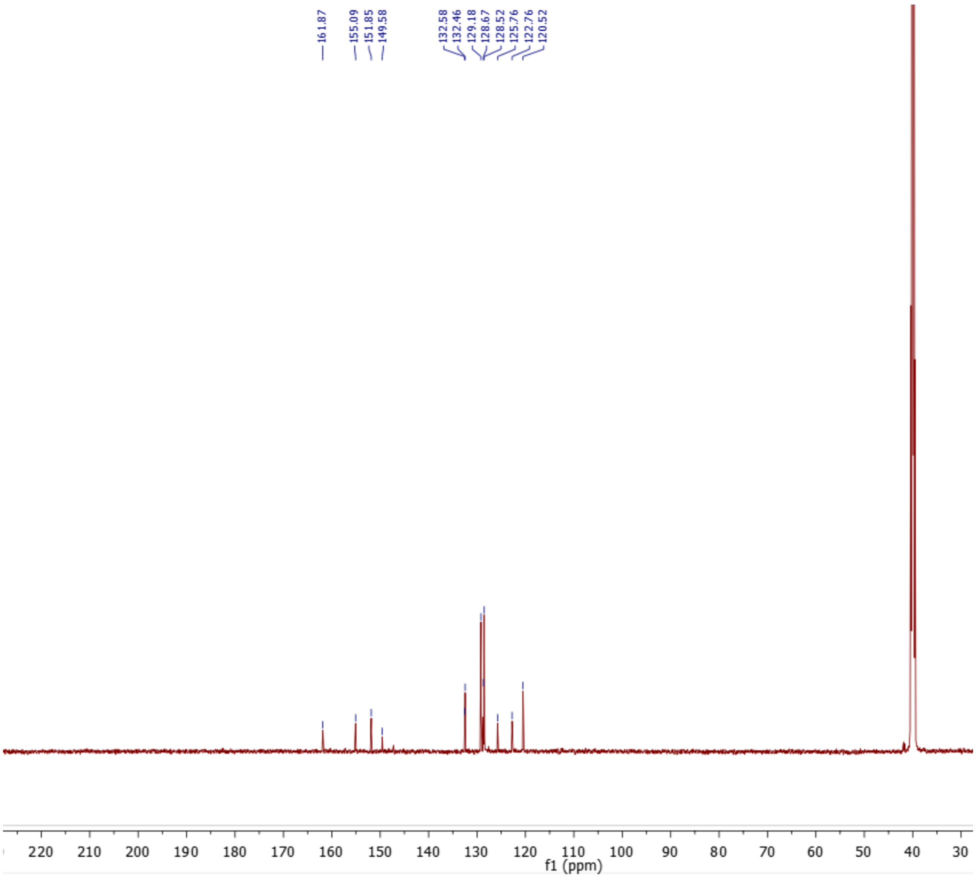

**Figure S73:** [13]C NMR of compound **66**

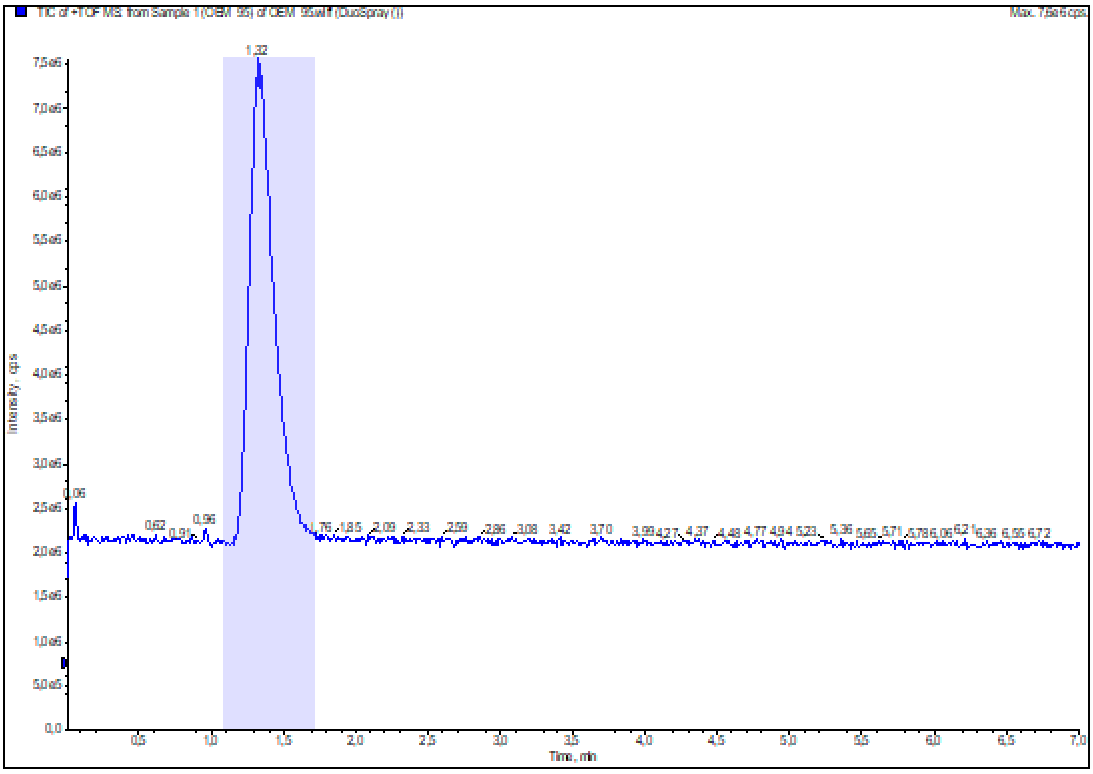

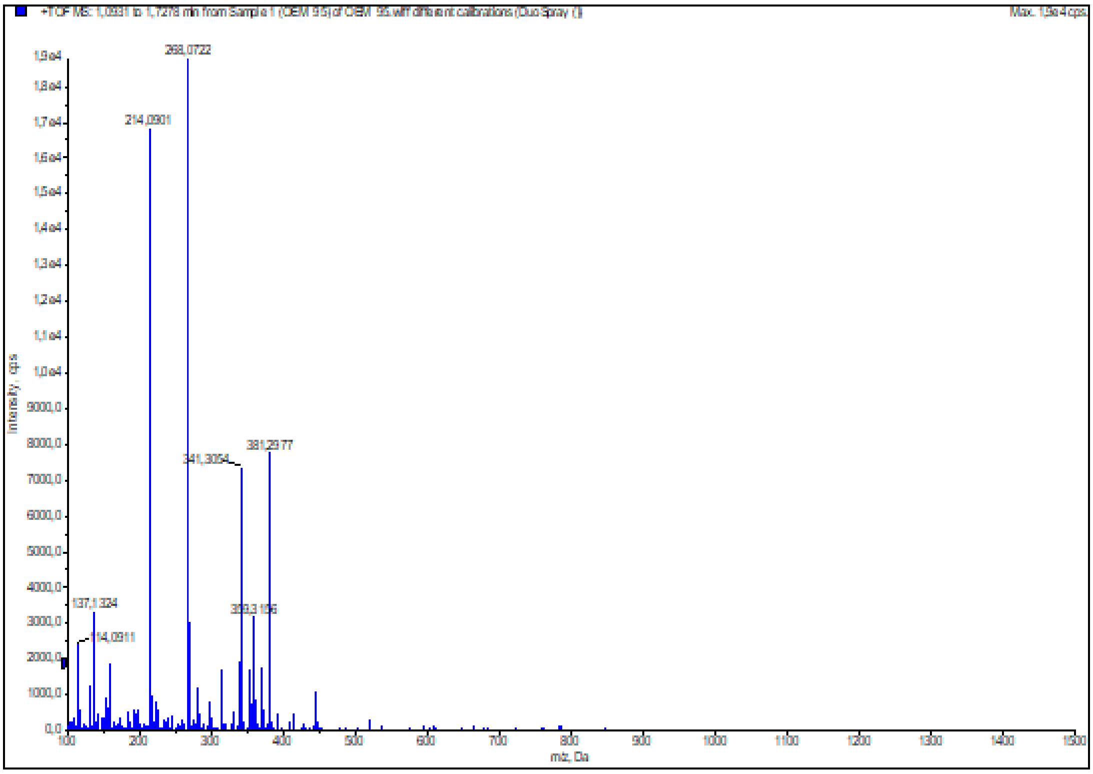

**Figure S74:** HRMS (ESI+) of compound **66**

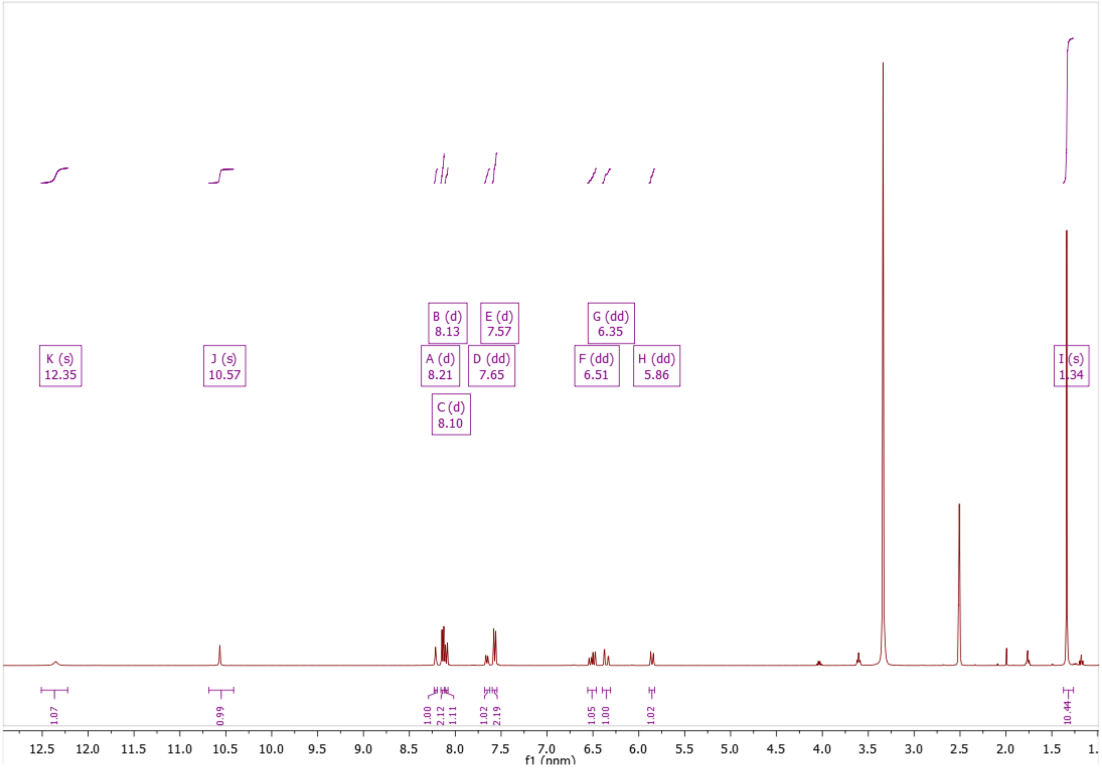

**Figure S75:** $^{1}$H NMR of compound **70**

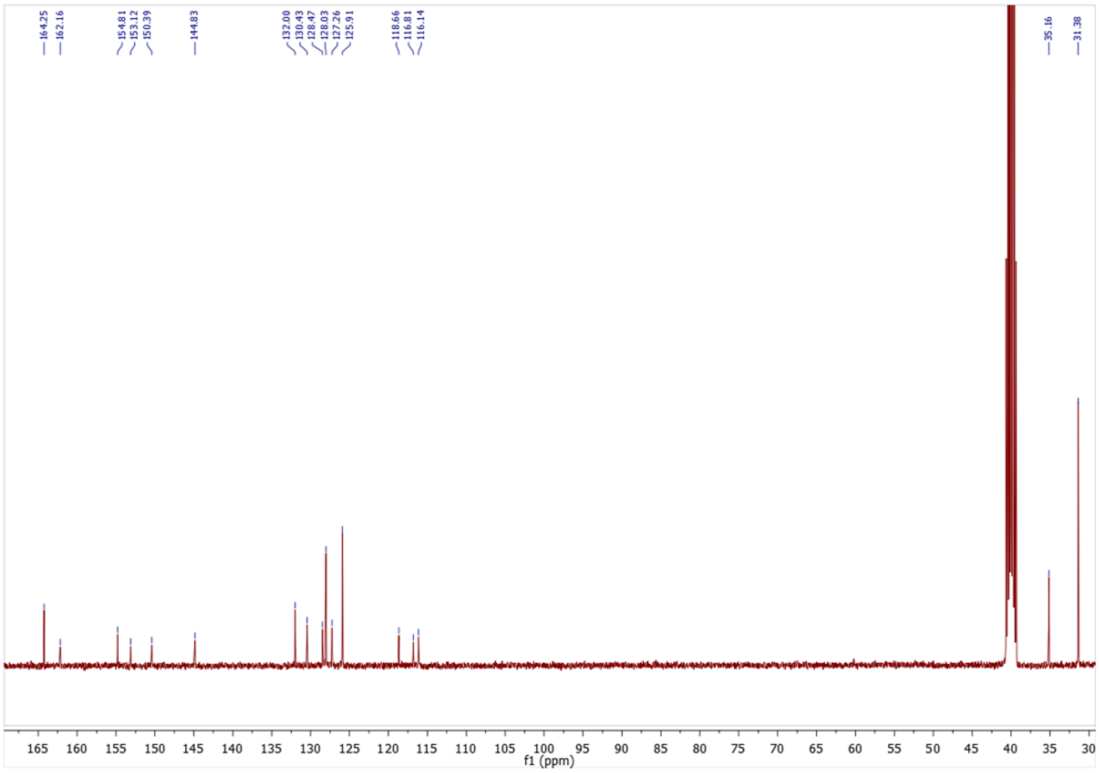

**Figure S76:** $^{13}$C NMR of compound **70**

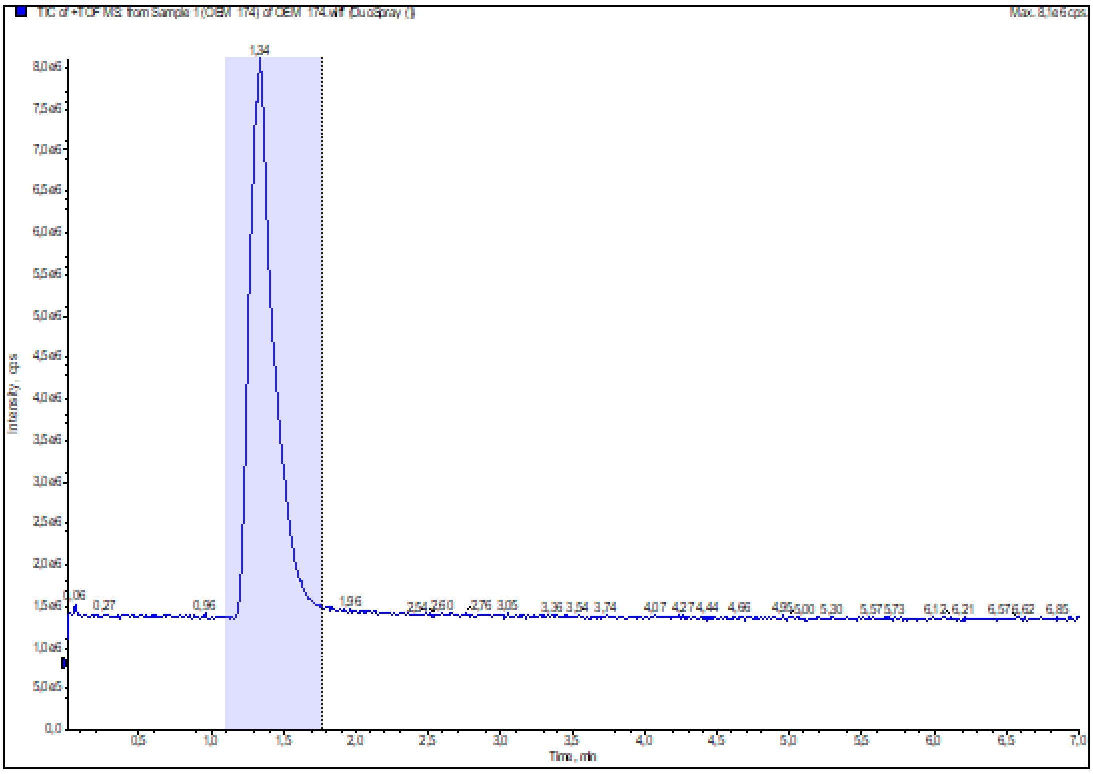

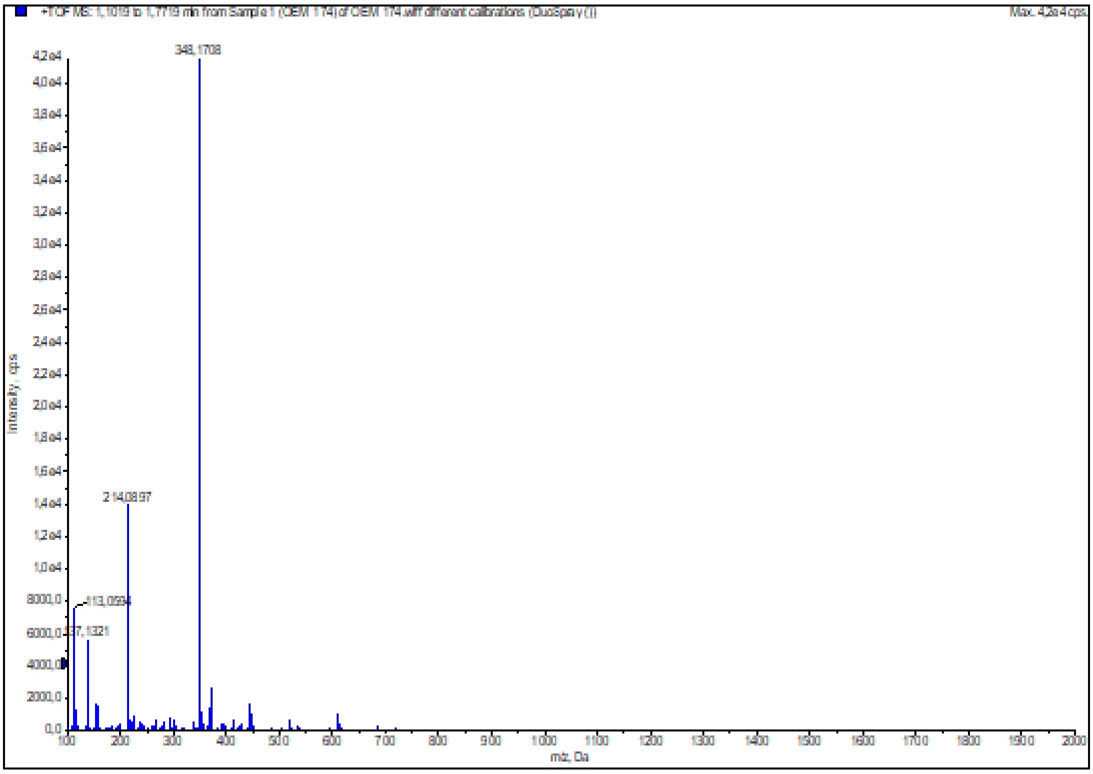

**Figure S77:** HRMS (ESI+) of compound **70**

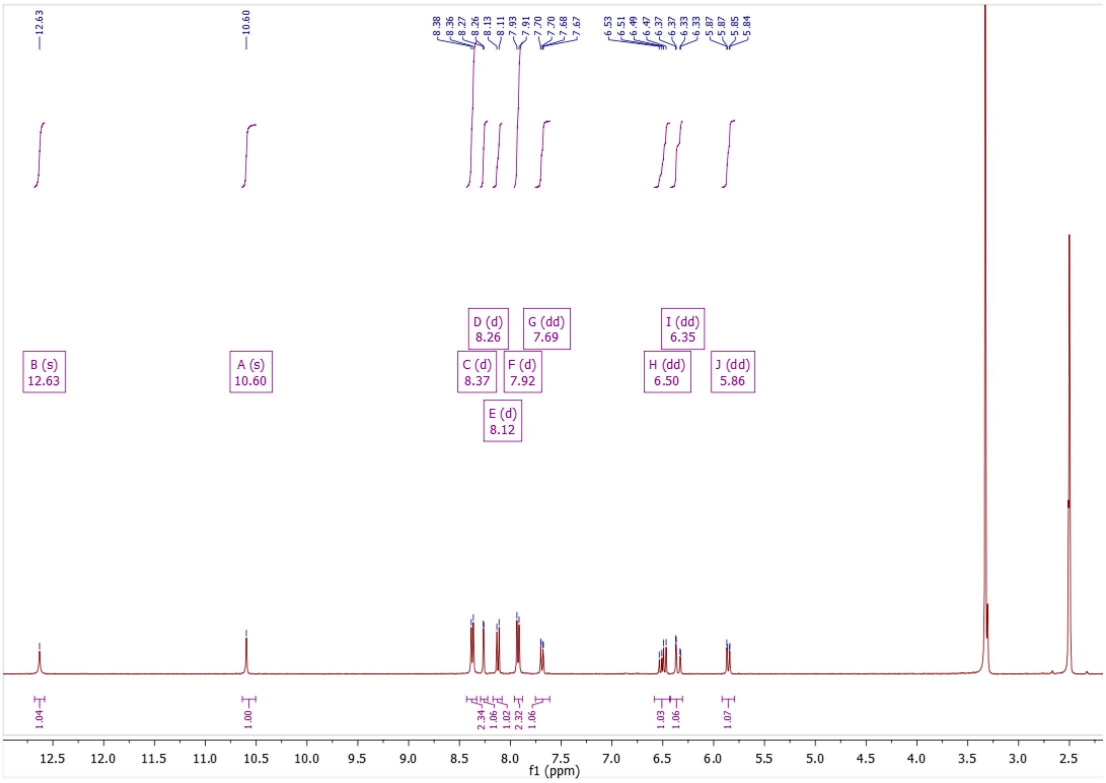

**Figure S78:** $^{1}$H NMR of compound **71**

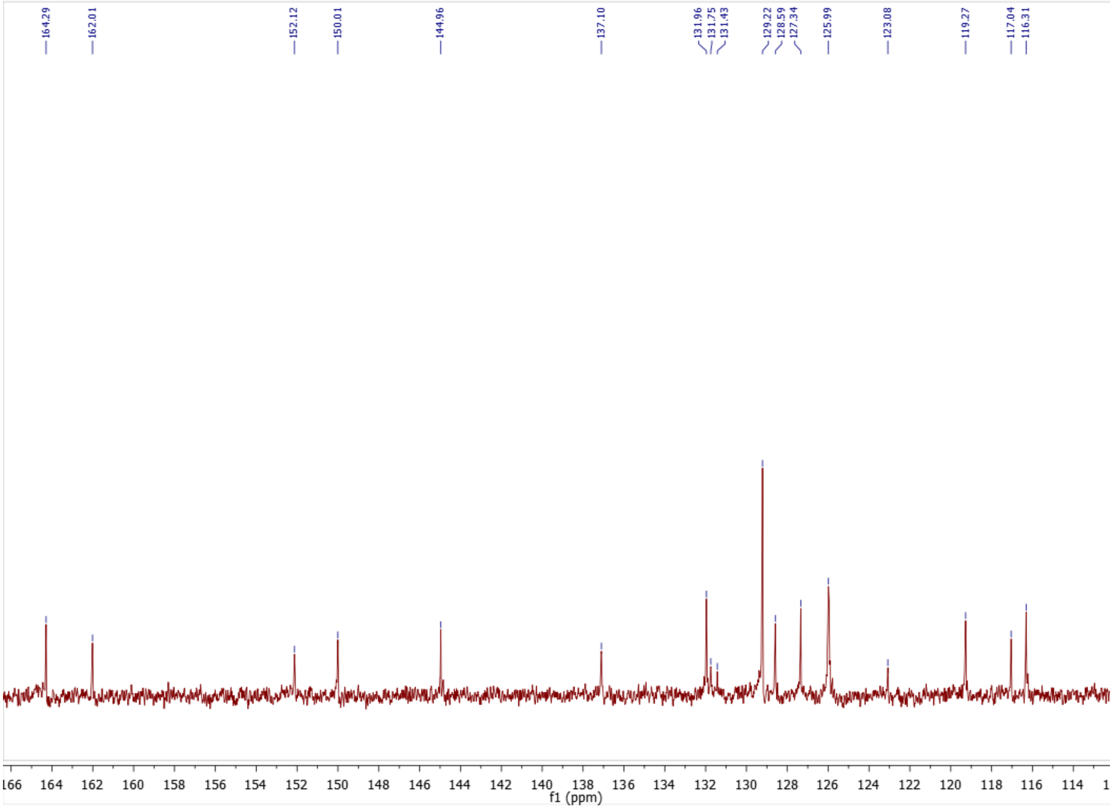

**Figure S79:** $^{13}$C NMR of compound **71**

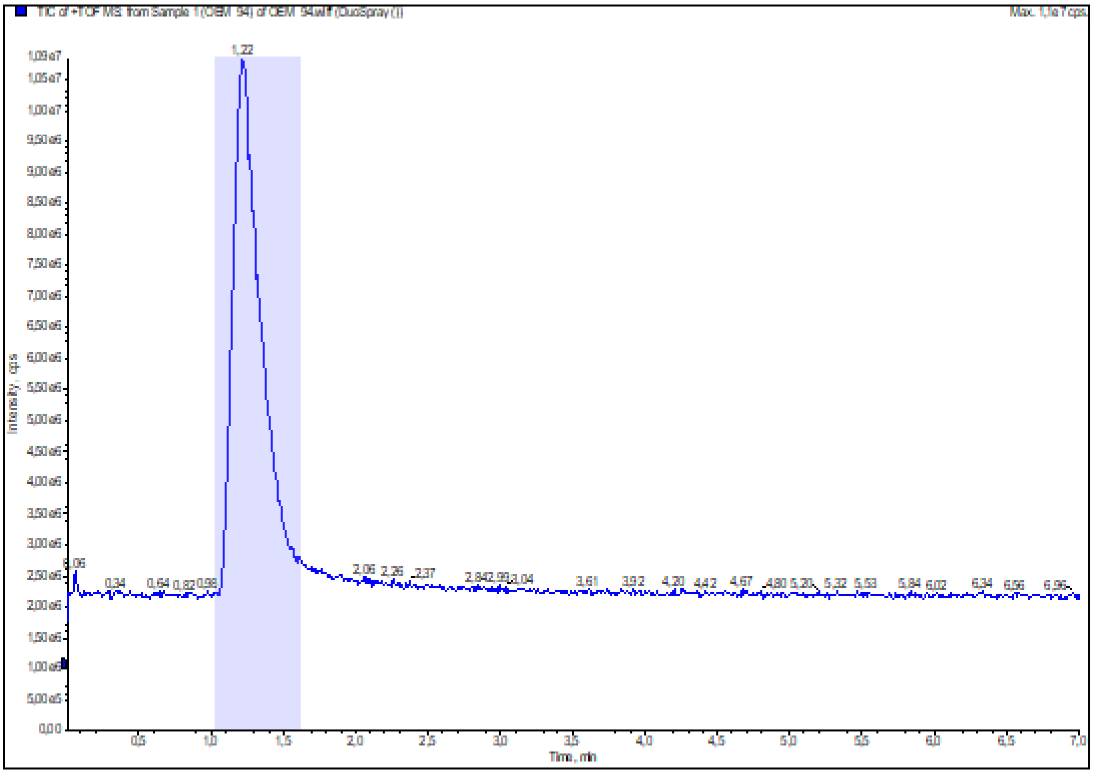

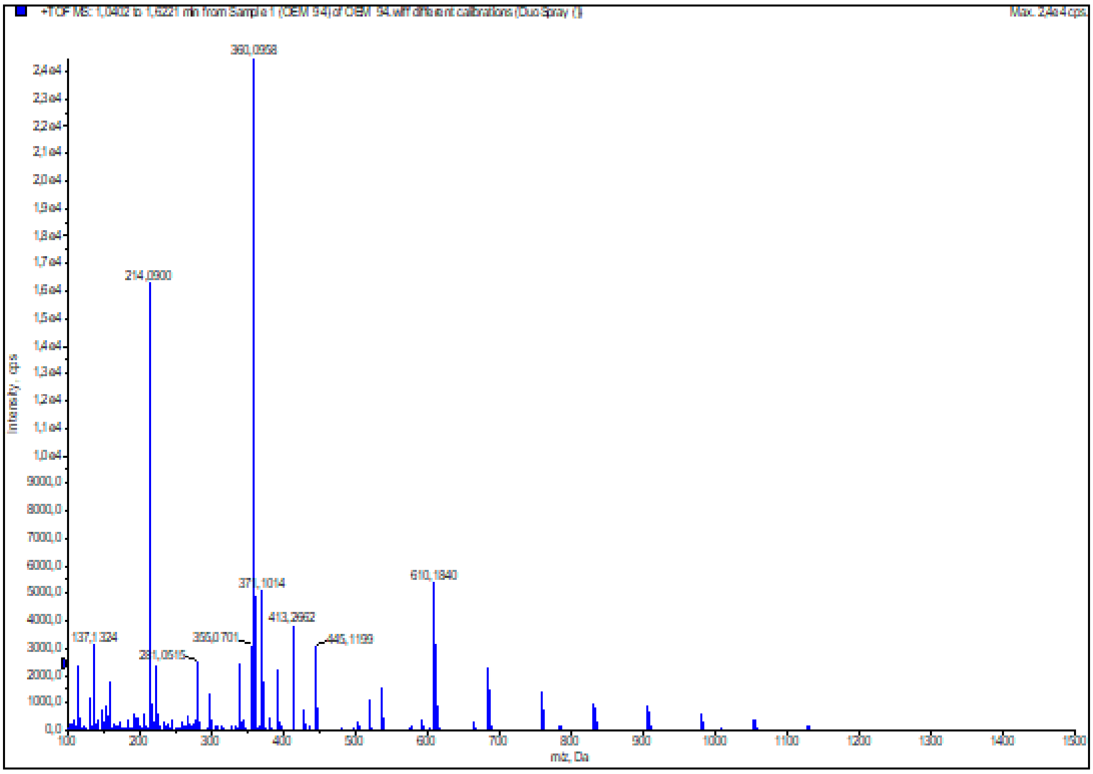

**Figure S80:** HRMS (ESI+) of compound **71**

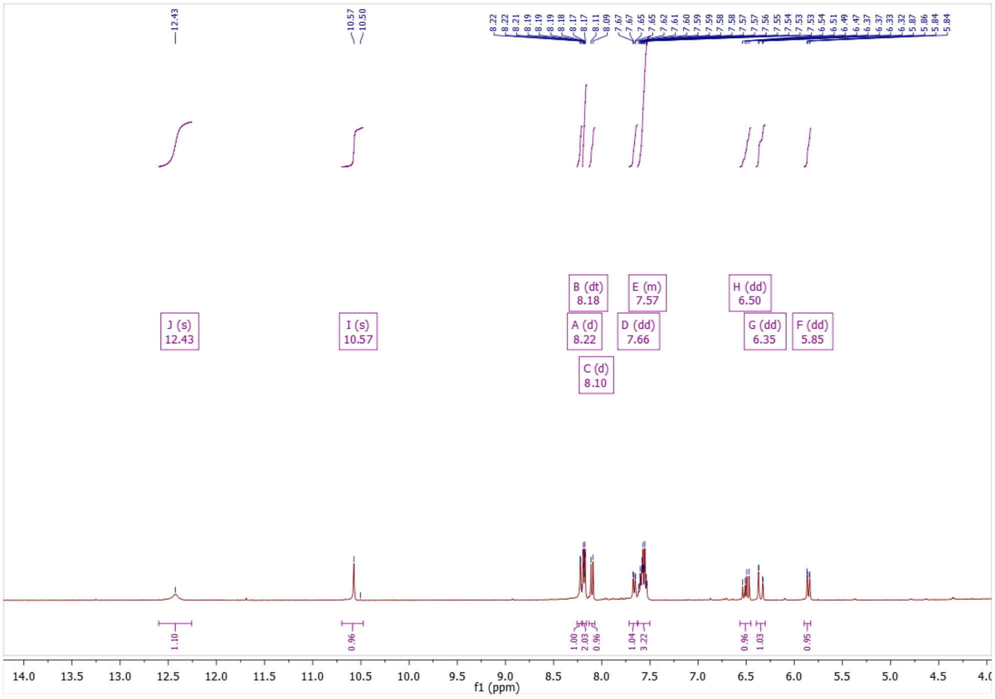

**Figure S81:** $^{1}$H NMR of compound **72**

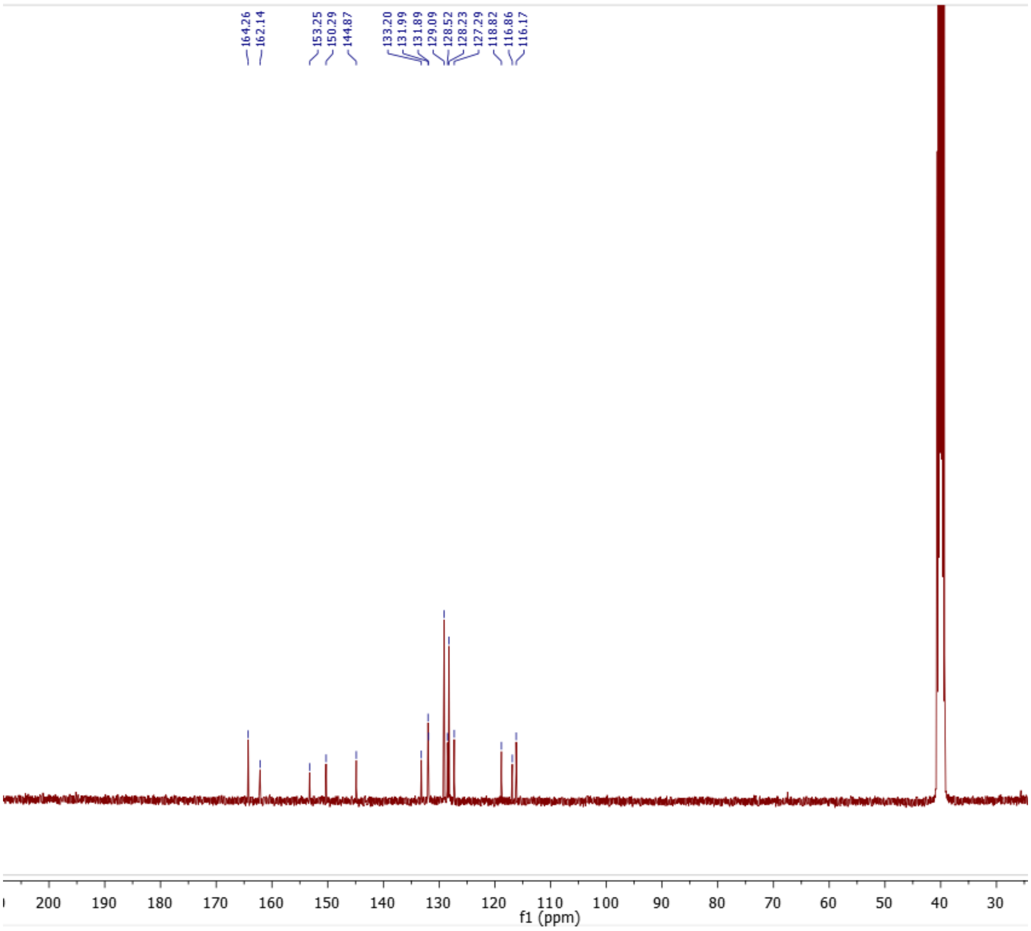

**Figure S82:** $^{13}$C NMR of compound **72**

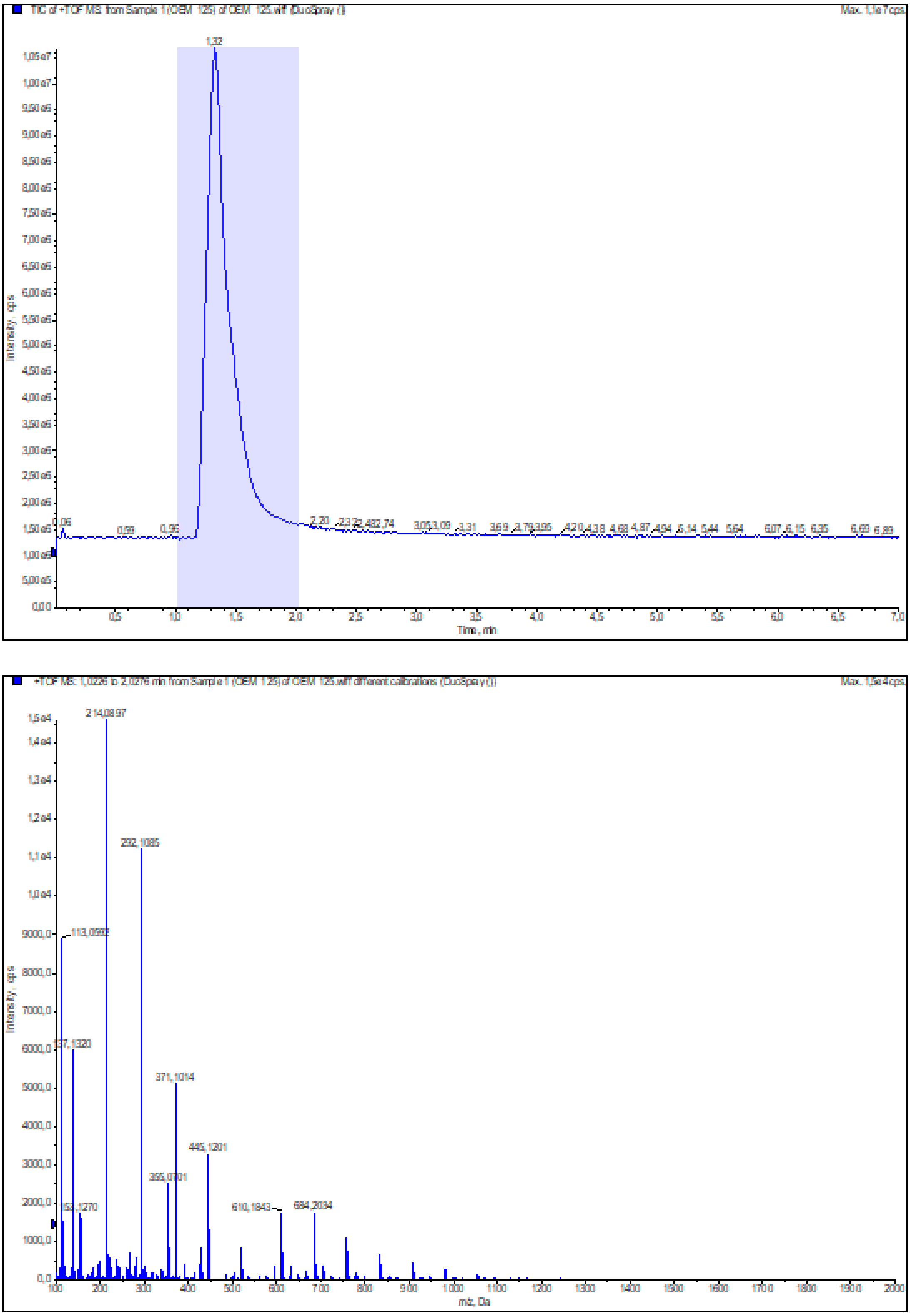

**Figure S83:** HRMS (ESI+) of compound **72**

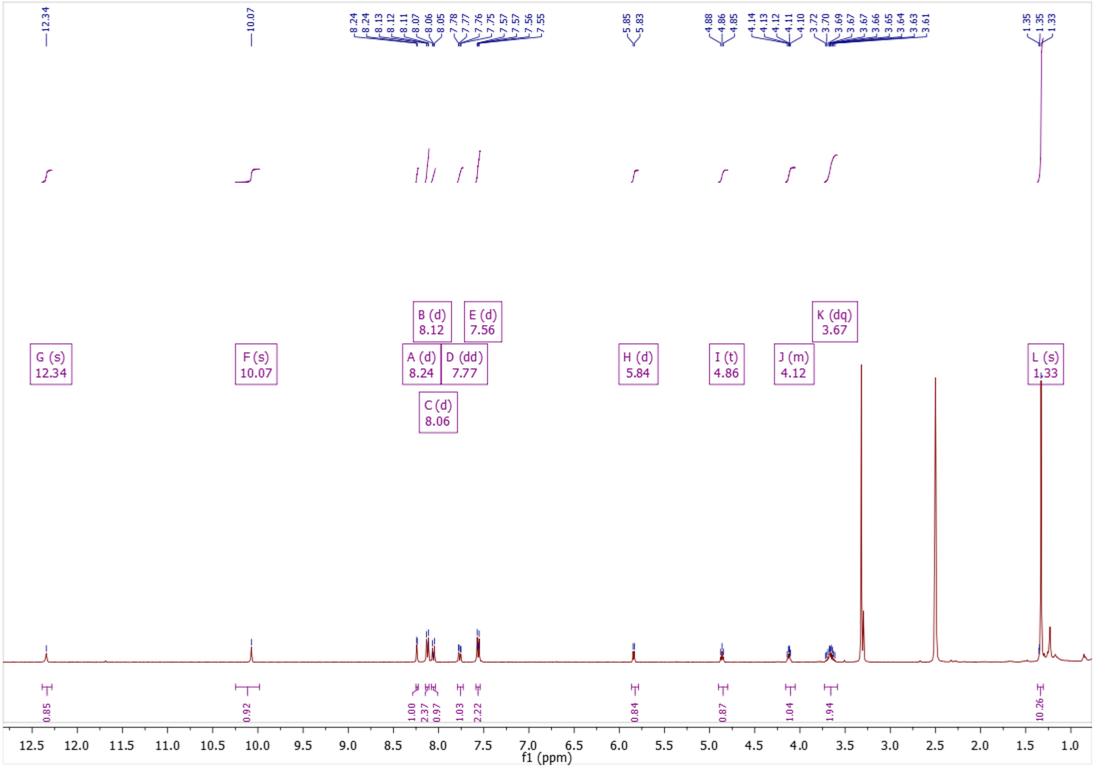

**Figure S84:** $^{1}$H NMR of compound **73**

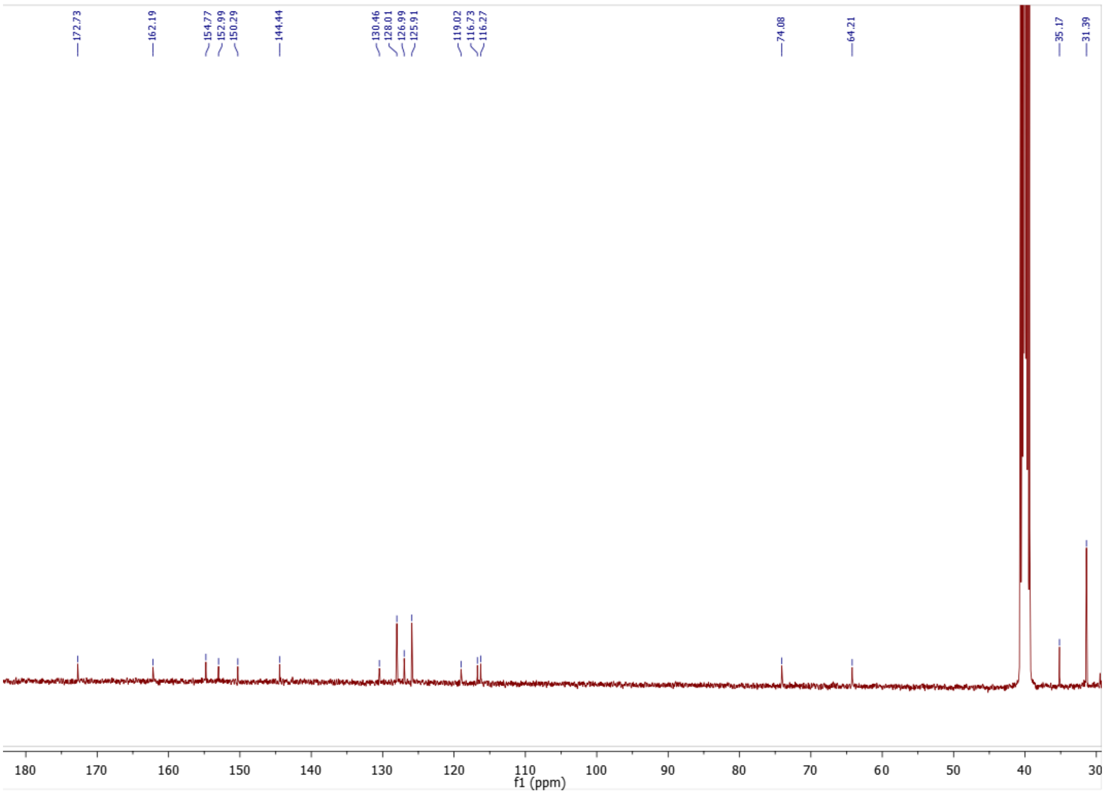

**Figure S85:** $^{13}$C NMR of compound **73**

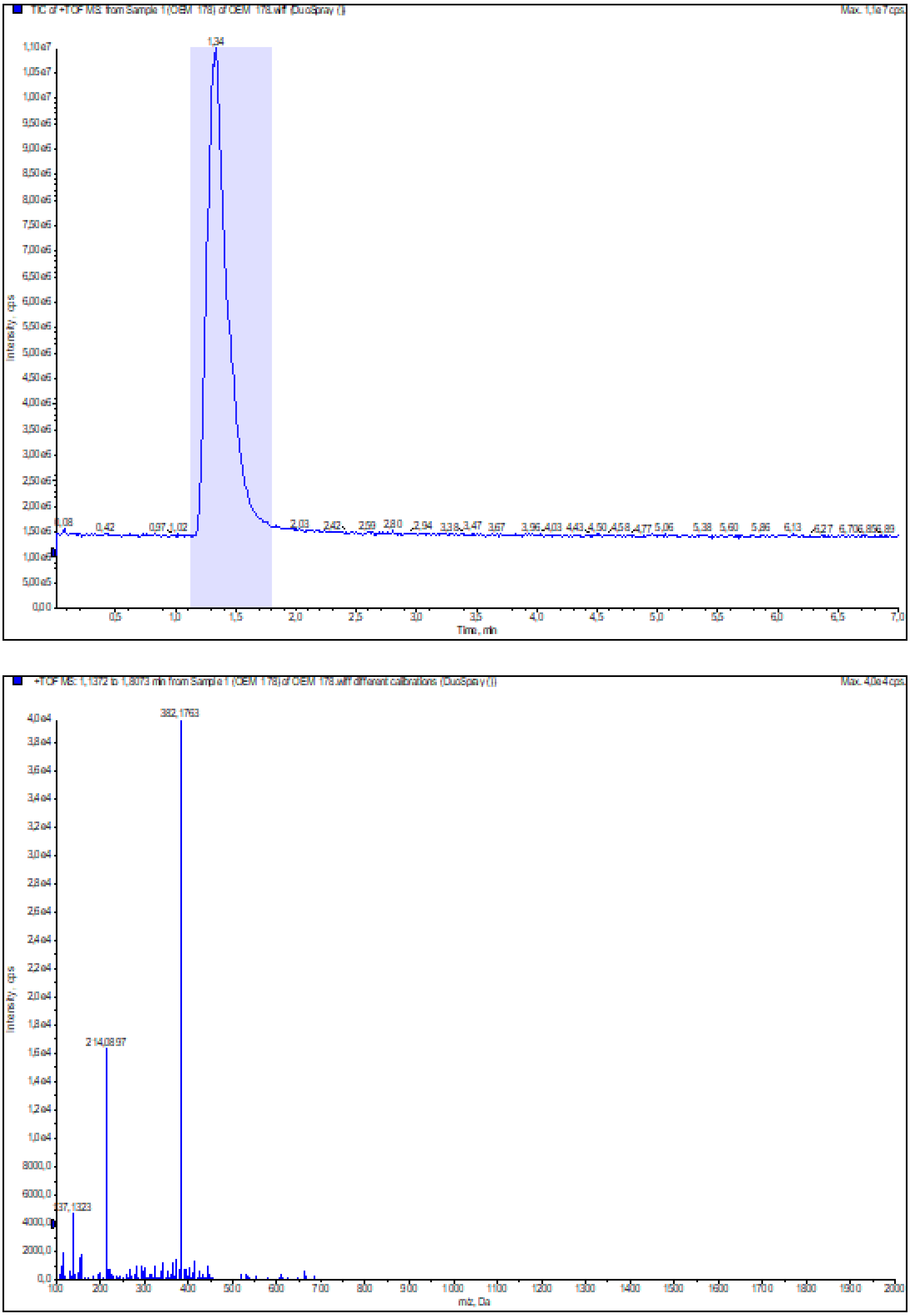

**Figure S86:** HRMS (ESI+) of compound **73**

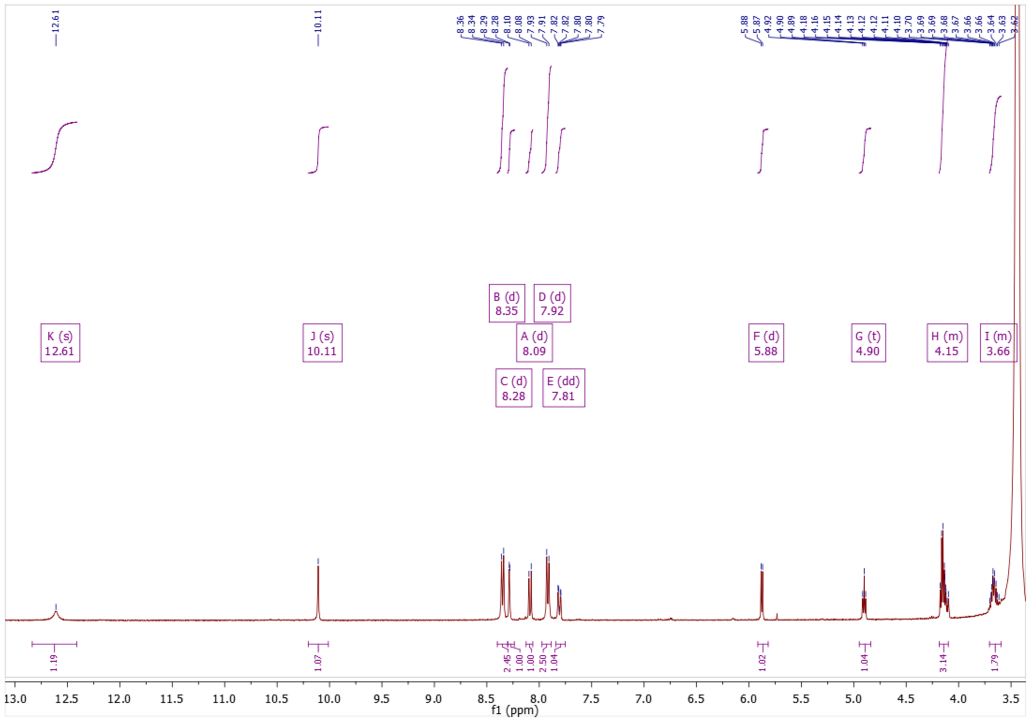

**Figure S87:** $^{1}$H NMR of compound **74**

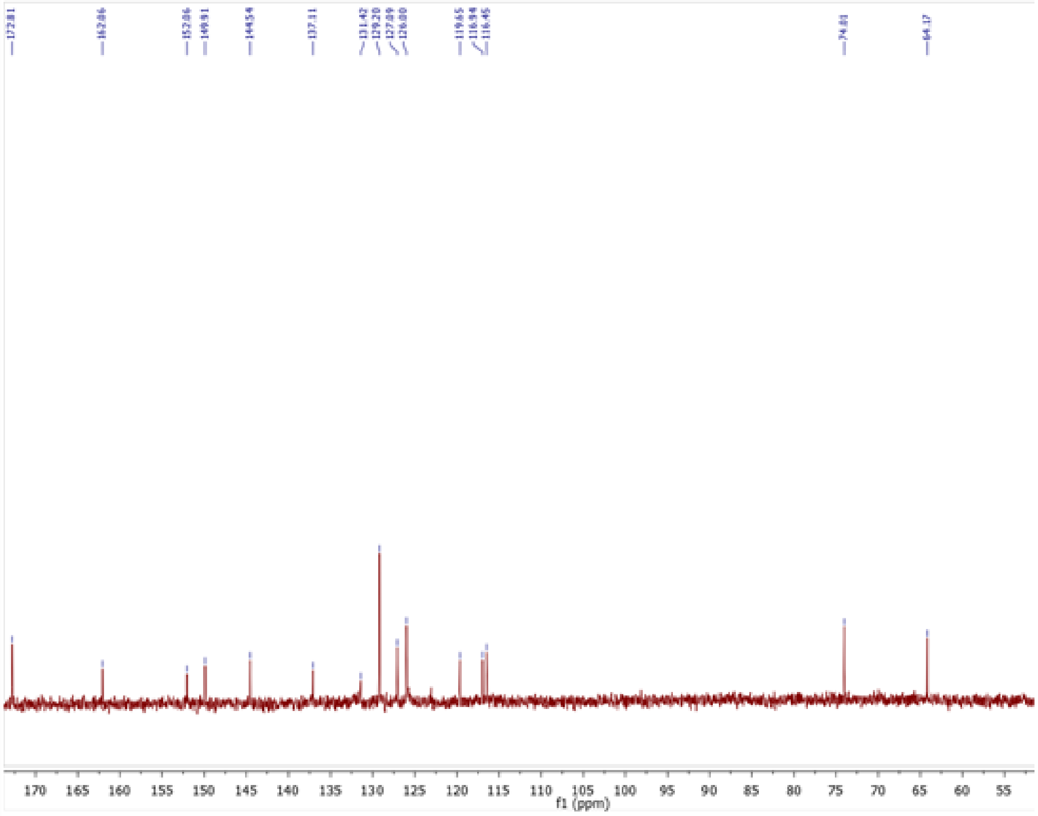

**Figure S88:** $^{13}$C NMR of compound **74**

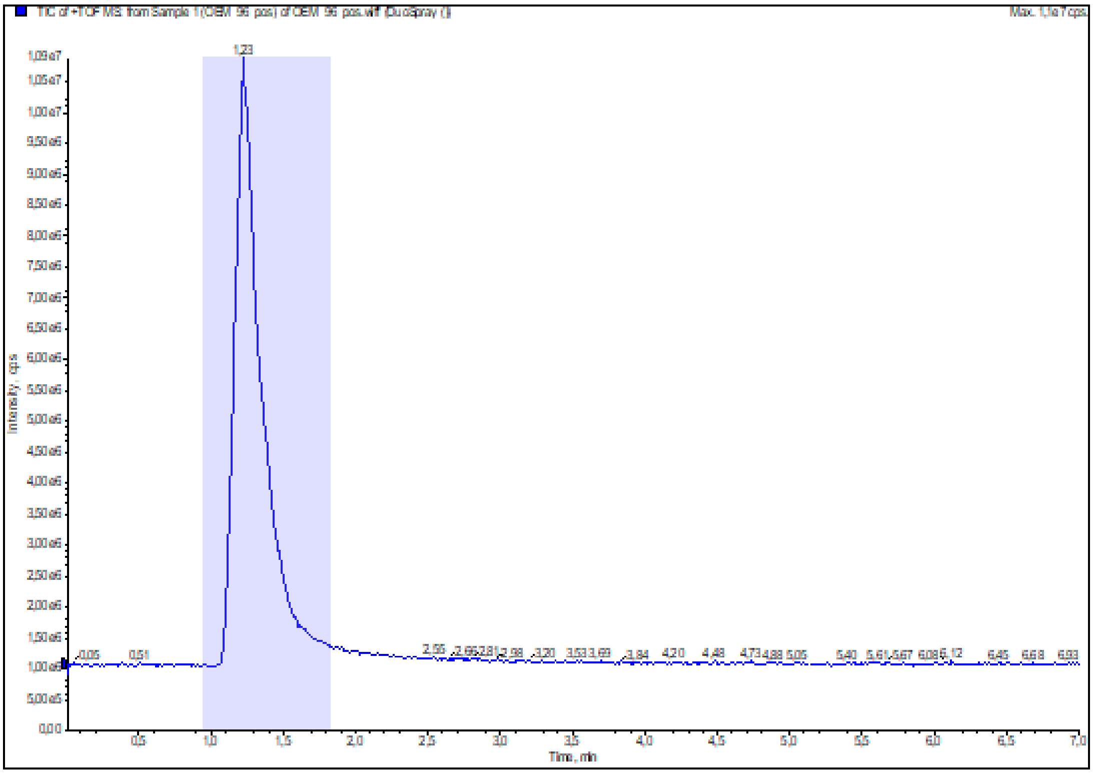

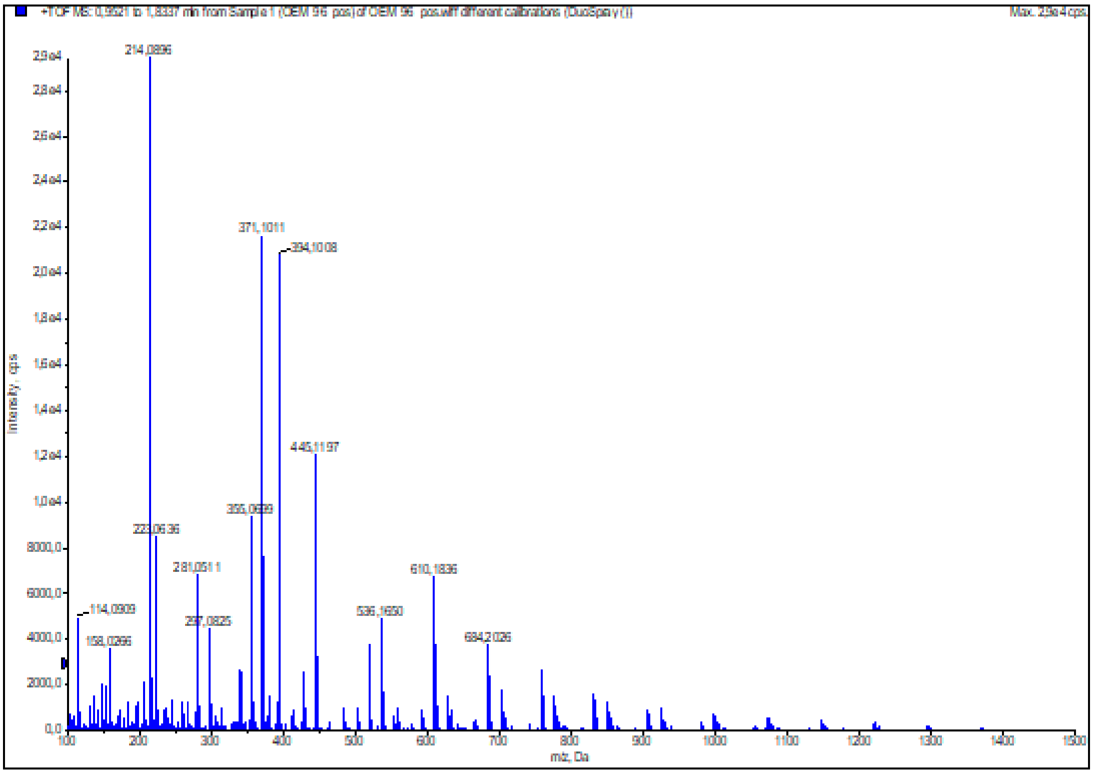

**Figure S89:** HRMS (ESI+) of compound **74**

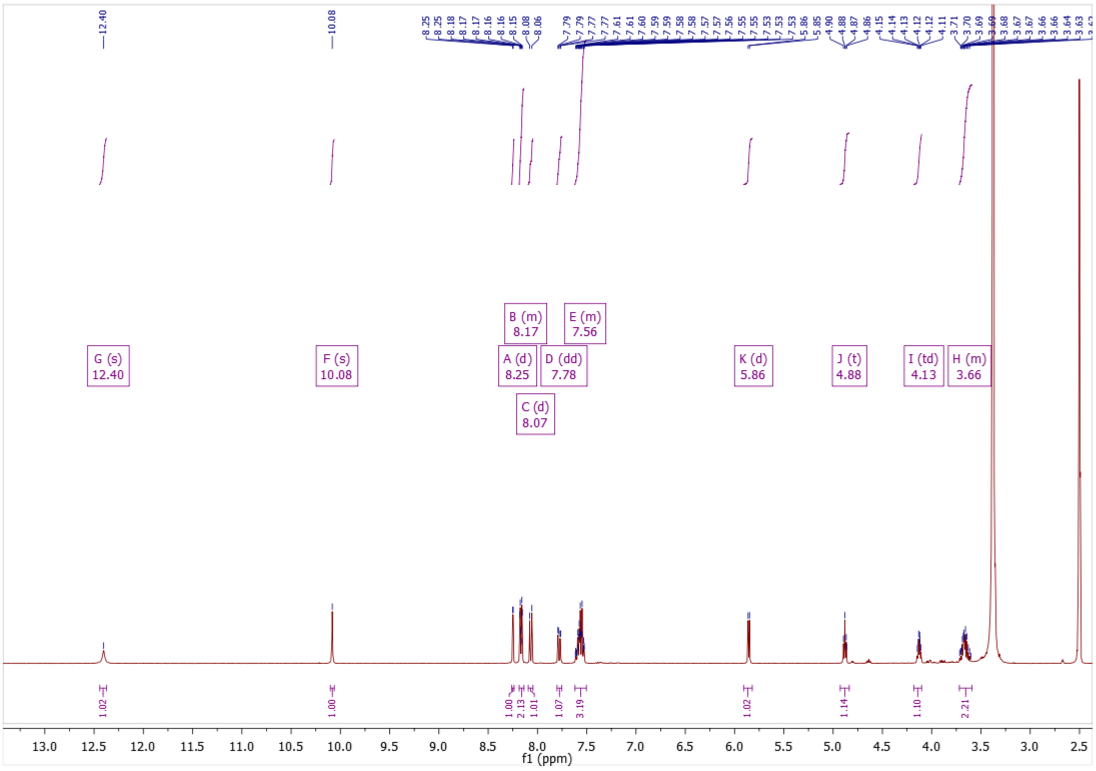

**Figure S90:** $^{1}$H NMR of compound **75**

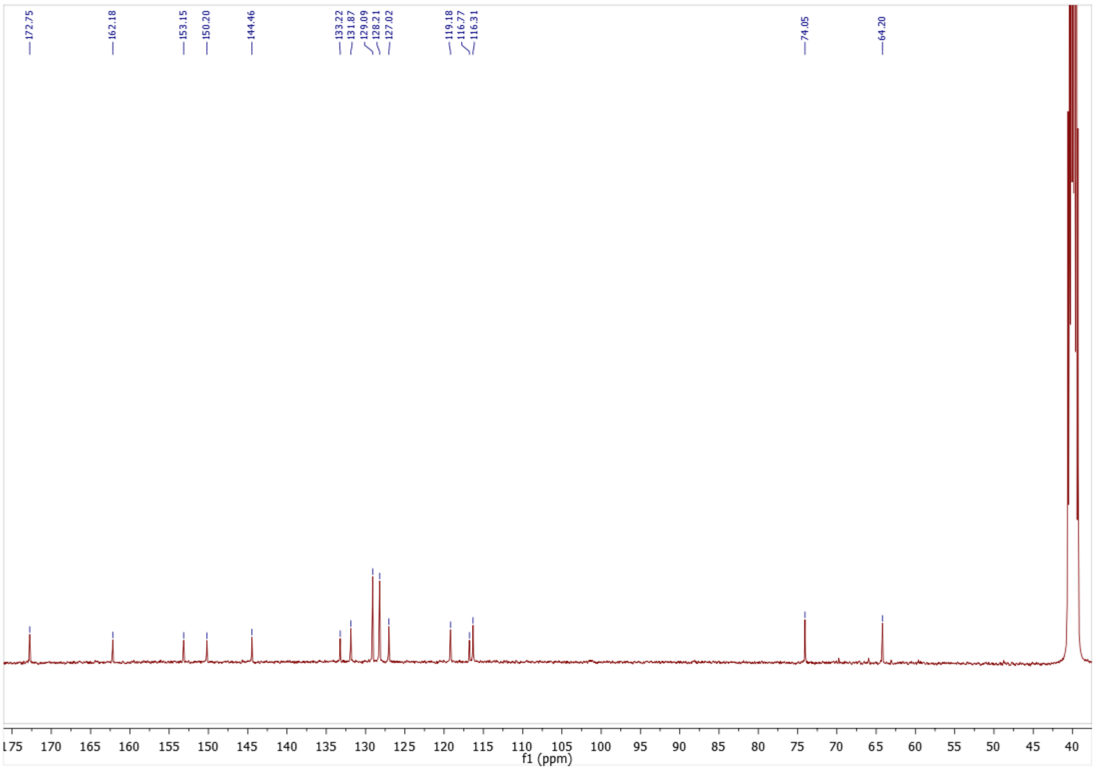

**Figure S91**: $^{13}$C NMR of compound **75**

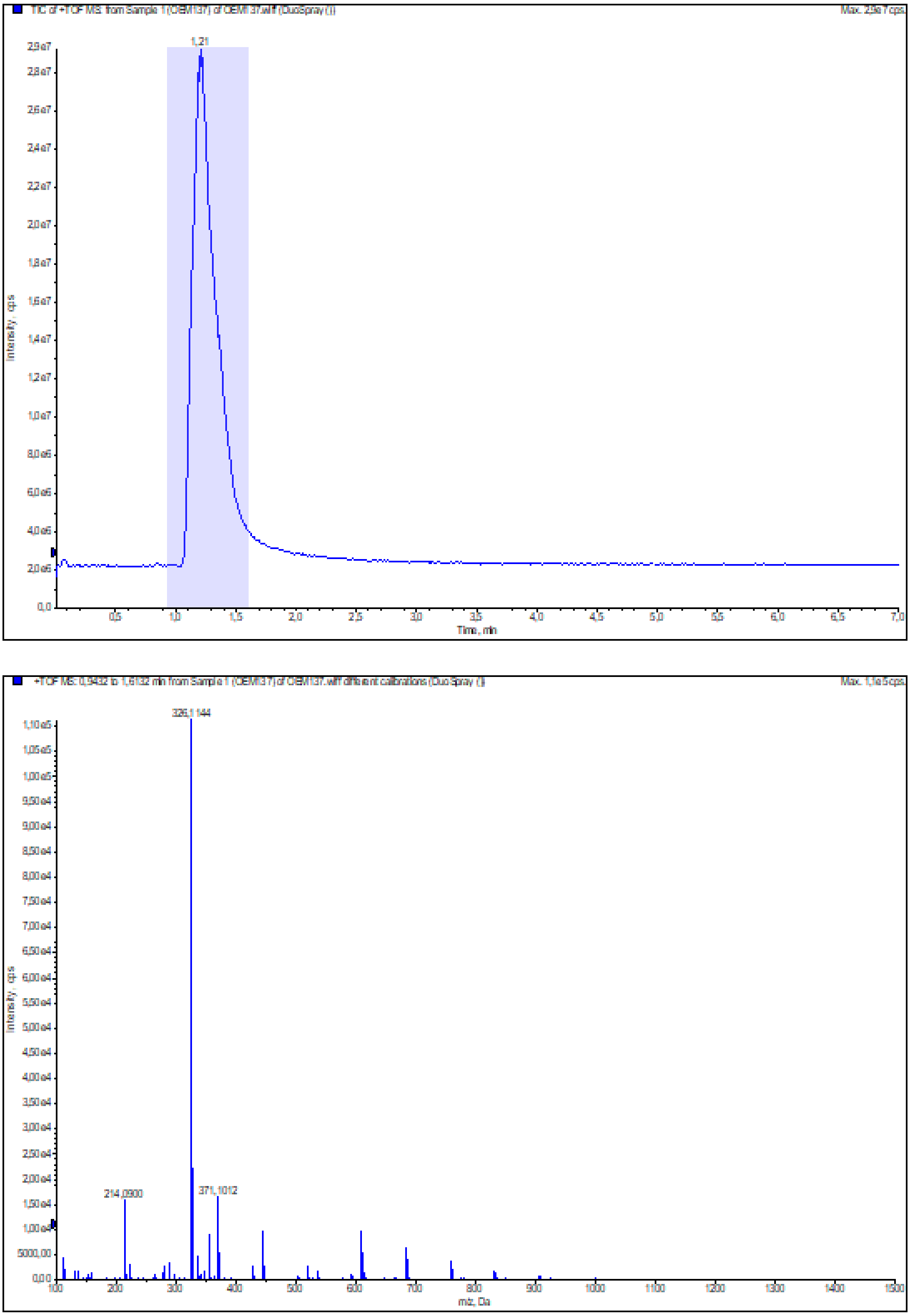

Figure S92: HRMS (ESI+) of compound **75**
